# m⁶A modification and prion-like domain proteins converge to dysregulate Neuronal RNA Granules in Alzheimer’s disease

**DOI:** 10.64898/2026.01.19.700388

**Authors:** Soukayna Boulaassafre, Hassan Ainani, Abdellatif El Khayari, Yuyu Song, Rachid El Fatimy

**Affiliations:** Faculty of Medical Sciences, UM6P Hospitals, Mohammed VI Polytechnic University, 43150, Ben-Guerir, Morocco; Department of Neurology, Massachusetts General Hospital, Charlestown, MA 02129, USA

**Keywords:** Neuronal RNA Granules (NRGS), N6-methyladenosine (m^6^A), Fragile X Mental Retardation Protein (FMRP), Prion-like domain proteins (PrLD), Synapses, Alzheimer’s disease (AD)

## Abstract

Alzheimer’s disease (AD) is a deadly neurodegenerative disorder with no cure. It is associated with several dysregulated pathways, including axonal transport. The latter supplies synapses with several essential components, including proteins and mRNAs. A proportion of RNAs in neurons is transported from the soma to neuronal extensions along microtubules in highly organized structures, known as Neuronal RNA Granules (NRGs). NRGs have a heterogeneous composition of coding and non-coding RNAs, RNA-binding proteins (RBPs), and components of translational machinery. In this study, we investigate the potential involvement of NRGs in AD pathogenesis, with a particular focus on the N6-methyladenosine (m⁶A), a key RNA modification, and prion-like domain (PrLD) proteins. Our *in-silico* analysis revealed that a significant portion of mRNAs in NRGs are likely to be highly methylated. Using transcriptomic data from AD brain, we identify dysregulation of key genes in the m⁶A-methylation pathway (*METTL3, FTO, YTHDF2/3, eIF3m*) as well as PrLD-containing proteins associated with NRGs (*STAU2, YBX1*). We further observe aberrant expression of m⁶A-methylated mRNAs within both NRGs and synapses. Gene Ontology analysis highlights disruptions in pathways related to NRGs and synaptic function. Together, our findings suggest that impaired NRGs homeostasis may represent a critical and previously underappreciated contributor to AD pathogenesis. By outlining the potential roles of m⁶A and PrLD proteins in regulating NRGs, this work offers a new conceptual framework to better understand AD and identify NRGs as a potential therapeutic target. Finally, we propose a working model illustrating how dysregulation of NRGs homeostasis may drive neurodegeneration in AD.

## Introduction

Alzheimer’s disease (AD) is a progressive neurodegenerative disorder primarily associated with memory loss and cognitive decline. It is characterized by several neuropathological hallmarks, including disruptions in axonal transport, which impair neuronal communication, contributing to the dying back neuropathy observed in AD^1–5^. Neuronal RNA Granules (NRGs) are membraneless subcellular structures that contain components of the translation machinery in a repressed state together with diverse RNA-binding proteins (RBPs) and a variety of RNA molecules, both coding and non-coding^6–8^. NRGs are transported from the soma to the neuronal extensions, where they facilitate localized protein synthesis within the synaptic compartments, allowing neurons to rapidly adapt to local functional demands^9,10^.

In contrast to other membraneless organelles, the molecular mechanisms governing NRGs assembly from the polyribosomes, the selection of their mRNA cargo formation, and their spatiotemporal dynamics remain poorly understood. A detailed understanding of how disrupted NRGs homeostasis contributes to axonal degeneration in neurodegenerative disorders, particularly AD, is therefore critical. We propose that, under physiological conditions, NRGs are actively transported along axons from the soma to distal neuronal compartments, thereby enabling localized protein synthesis at synapses. In AD, this transport process may be compromised, leading to aberrant subcellular distribution of NRGs. Such mislocalization is likely to perturb NRGs dynamics and homeostasis, ultimately contributing to synaptic dysfunction and axonal degeneration. In this study, we focus on two key elements: epitranscriptomic N⁶-methyladenosine (m⁶A) modifications and proteins containing disordered prion-like domains (PrLDs) and investigate how these factors may influence NRGs formation, assembly and potential dysregulation in the context of AD pathogenesis.

Epitranscriptomic modifications represent a recently decoded form of gene expression regulation in health and disease, including neurological related pathologies^11–14^. Currently, over 170 distinct RNA nucleoside modifications have been discovered^14^ including inosine^15^, 5-hydroxymethylcytidine^16^, N6,2′-O-dimethyladenosine^17,18^, N7-methylguanosine^19^, pseudouridine (Ψ)^20,21^, N1 -methyladenosine^20^ and N6-methyladenosine (m⁶A)^21^. Among these, m⁶A stands out as the sole extensively characterized modification in mRNA, accounting for approximately 80% of the known modifications^22^. It is a highly prevalent, evolutionarily conserved, and reversible post-transcriptional modification of RNA that has been identified on various RNAs, including mRNAs, ribosomal RNAs (rRNAs), transfer RNAs (tRNAs), microRNAs (miRNAs), and other non-coding RNAs (ncRNAs)^21,23–27^. This modification is regulated by specific m⁶A regulatory enzymatic complex composed of 1) methyltransferases, known as “writers,” which catalyze m⁶A such as the *METTL3/METTL14*; 2) demethylases known as “erasers” such as *FTO* and *ALKBH5* which remove this modification from methylated bases ; 3) specific m⁶A -binding proteins called “readers”, which can recognize the methylation such as the *YTHDF* and *YTHDC* family^28,29,30, 31,32^. m⁶A influences different aspects of RNA metabolism, including RNA splicing, translation, stability, nuclear export, and mRNA decay. Additionally, m⁶A plays key roles in essential biological processes, most notably neurogenesis, axon regeneration, and brain development^25,33–36^. Moreover, dysregulation of m⁶A has been implicated in a range of neurodevelopmental and neurodegenerative disorders^32,36–40^. Transcriptome-wide mapping using m⁶A-specific methylated RNA immunoprecipitation combined with next-generation sequencing has shown that m⁶A is enriched predominantly within the gene (94.8%)^24^. In the untranslated region (UTRs), m⁶A is largely concentrated near the stop codon^21,23,24^. At this level, this modification plays a significant role in post-transcriptional regulation of gene expression by influencing RNA stability^41,42^, and the efficiency of translation termination, potentially impacting the final protein product generated from the mRNAs^24,43–45,46^. m⁶A RNA modification exhibits significant tissue specificity, with the brain displaying the highest detected sites^49^. This modification plays a crucial role in various biological processes, including embryonic stem cell differentiation and brain development. We hypothesize that aberrant m⁶A-methylation homeostasis in AD compromises the machinery required for precise regulation of translation in neurons, potentially driving early synaptic and axonal pathology.

The PrLD are low-complexity AA sequences present in a subset of proteins^47^. Several studies have shown that proteins containing these domains can drive the formation and spread of pathological inclusions in several neurodegenerative diseases, including AD^48–50^. Prion-like mechanisms enable Tau protein to trans-seed and promote the formation of tau aggregates, supporting the propagation of tau pathology in AD^51^. Consistently, tau exhibits prion-like spreading behavior in both AD animal models^52^ and human brain tissue^53^. Beyond their role in disease-associated protein aggregation and propagation, PrLD contribute to the assembly of membraneless organelles, such as stress granules (SG), through direct interactions with components of the translational machinery^62,63^. Given these properties, we speculate that PrLD-containing proteins may play a role in NRGs formation by phase transition from polyribosome-associated particles to coalescent form as granules.

In this study, we propose that alterations in the processes linked to NRGs formation, composition, and transport could be associated with neurodevelopmental and neurodegenerative disorders like AD^8,20–33^. We further propose that m⁶A-modifications may influence the selective recruitment of RNAs into granules and modify their behavior. In addition, we suggest that the dynamic interplay of intrinsically disordered PrLD-containing proteins may be crucial for the transition of polyribosomes into condensed granules. To investigate these mechanisms, we analyzed gene expression profiles from both healthy and AD brains, focusing on m⁶A regulatory factors and PrLD-containing proteins implicated in NRGs biology. Overall, we propose that, beyond the potential mis-localization arising from axonal transport defects, dysregulation of m⁶A signaling, and PrLD within NRGs may disrupt synaptic RNA translation, thus contributing to the neurodegenerative processes observed in AD.

## Results

### 1. m⁶A-methylation enrichment in NRGs, nerve terminals, and FMRP targets

We evaluated the m⁶A-methylation status of mRNAs in NRGs and compared it to that of mouse hippocampal cultured neurons (MHCN), neuronal extensions, and rat neuropil (RN) by analyzing publicly available RNA-seq datasets. Specifically, we intersected the significantly expressed mRNAs in three models with the published m⁶A-CLIP datasets generated from mouse embryonic stem cells (mESC1)^59^, mESC2^60^, mouse neuronal stem cells (mNSC) during proliferation & differentiation^61^, mouse embryonic fibroblast cells (mEFC)^62^, and Human embryonic stem cells (hESC)^60^ (Supplementary table S1). Our findings indicate that NRGs contain the highest proportion of m⁶A-methylated mRNAs compared with RN and MHCN (Fig. 1A and Fig. S1), indicating a marked enrichment of methylated mRNAs within NRGs structures. Using the same analytical approach, we compared m⁶A-methylation with another RNA modification, pseudouridine modification (Ψ). We evaluated the presence of Ψ-modified mRNAs in NRGs, RN, and MHCN using data from a mouse brain Ψ-mapping study generated by N3-(N-cyclohexyl-N’-β-(4 methylmorpholinium) ethylcarbodiimide)–enriched Ψ-sequencing^63^. Our analysis showed that Ψ-modified mRNAs are markedly less abundant in these structures compared with m⁶A-methylation (Fig. 1A and Fig. S1). These findings indicate that m⁶A is a predominant RNA modification in these structures, suggesting a significant role played by this modification in mRNAs enriched in NRGs and axonal terminals relative to Ψ modification. To further strengthen our findings, we explored m⁶A-methylation within the mRNA targets of one of the highly enriched RBPs in NRGs, the fragile X mental retardation protein (*FMRP*)^7^. To do so, we analyze FMRP-targeted mRNAs identified in the adult mouse brain (MB FMRP)^64^, adult mouse brain polyribosome (MBP FMRP)^65^, and embryonic mouse cerebral cortex (EmCrtx FMRP)^66^, and intersected these datasets with m⁶A - methylated mRNAs derived from the studies described above following the previously outlined approach (Supplementary table S1 and Fig. S1). Interestingly, our results revealed that a large proportion of FMRP targets in EmCrtx, MB, and MBP were highly methylated (Fig. 1B; Fig. S1), indicating widespread modification of FMRP-associated transcripts. We next compared m⁶A with Ψ RNA modification within FMRP targets by overlapping MB FMRP targets with Ψ-modified mRNAs of the previously described dataset^63^. Together, these results indicate that the Ψ-modified mRNAs are considerably lower in FMRP targets when compared to m⁶A-methylation (Fig. 1B and Fig. S1) and highlight a pronounced enrichment of m⁶A within RBP-associated mRNAs enriched in granules such as FMRP.

**Figure 1:**
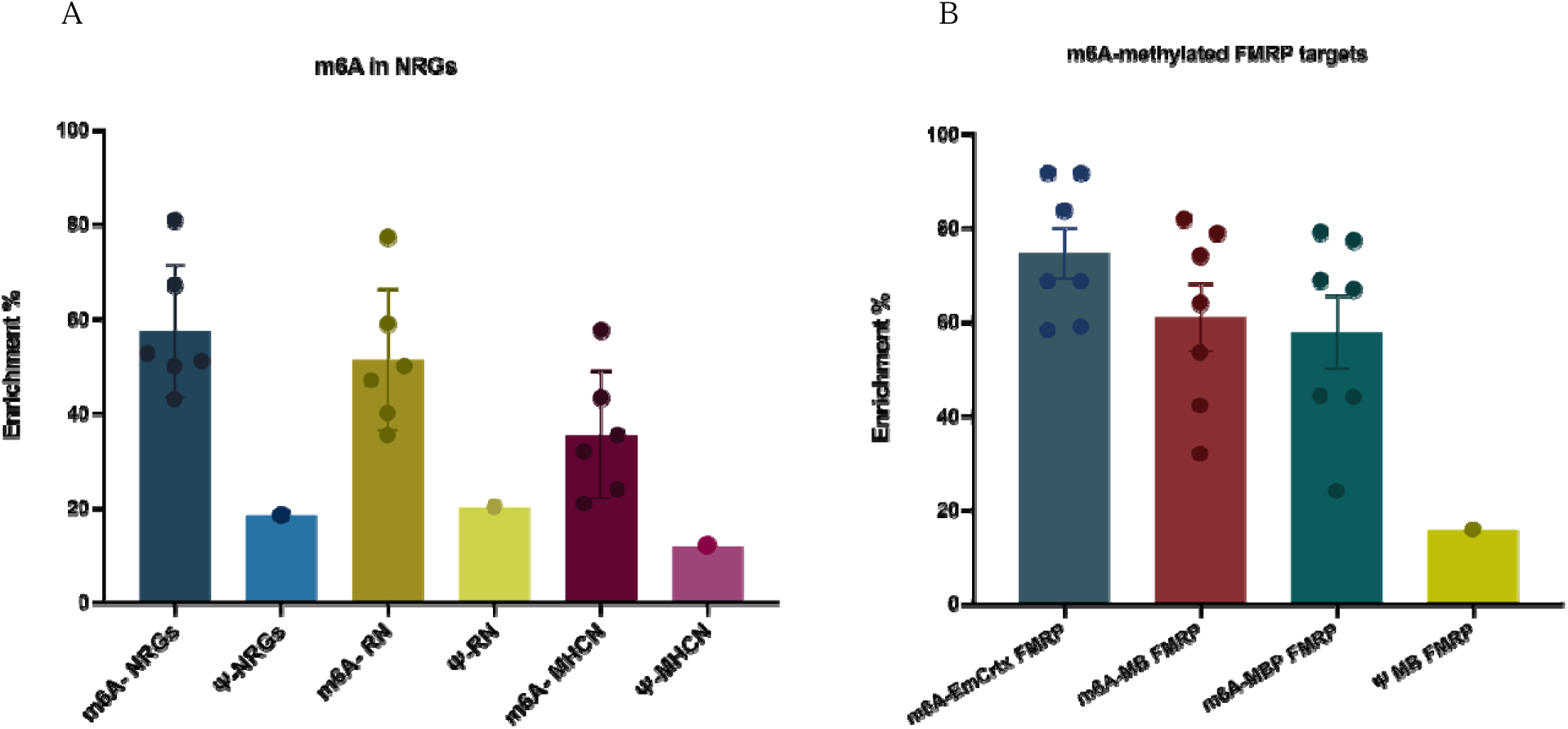
Relative enrichment of potentially m□A-methylated mRNAs detected in different structures. **(A):** Estimated proportions of m□A -methylated mRNAs obtained by overlapping lists of mRNAs from NRGs (n= 1798), RN (n=2550), and MHCN (n= 2160), with lists of experimentally validated m□A -methylated mRNAs obtained from: mESC1 (n=14559), mESC 2 (n=5213), mNSC Proliferation (n=9206), mNSC Differentiation (n=7336), mEFC (n=15453), hESC (n=7530) and Ψ modification (n=1372). (**B**): Percentage of m□A-methylated FMRP targets obtained after overlapping lists of FMRP targets from EmCrtx FMRP (n=856), MB FMRP (n=443), MBP FMRP (n=842), with lists of m□A-methylated mRNAs obtained from the previously mentioned studies.

We next examined the distribution of m⁶A sites within the NRGs transcriptome using a published MeRIP-Seq dataset, a technique that utilizes methylated RNA immunoprecipitation, combined with next-generation sequencing (MeRIP-Seq) from total mouse brain^26^, which we overlapped with our transcriptomic data from mouse granules. This approach allowed us to determine the positional enrichment of m⁶A-modification sites within NRGs-associated mRNAs. Our results showed that the majority of m⁶A sites within the NRGs mRNAs occur in the coding sequences (CDS; 66.83%) followed by 3’ untranslated regions (3’UTR; 56%), with relatively few peaks mapping to introns (22.3%) and 5’untranslated regions (5’UTR; 8.7%) (Fig. 2A). When overlapped to the set of filtered m⁶A peaks, which is the total list of m⁶A peaks identified in all MeRIP-Seq replicates (Fig. 2A). Similar results emerged when considering high-confidence sites, with enrichment primary in CDS (41%) followed by 3’UTR (32%). However, the peaks are less abundant in the introns and 5’UTR (2.27% and 2.3%, respectively) (Fig. 2B). Together, these findings highlight a strong positional preference for m⁶A within CDS and 3′UTRs of NRGs-associated transcripts.

**Figure 2:**
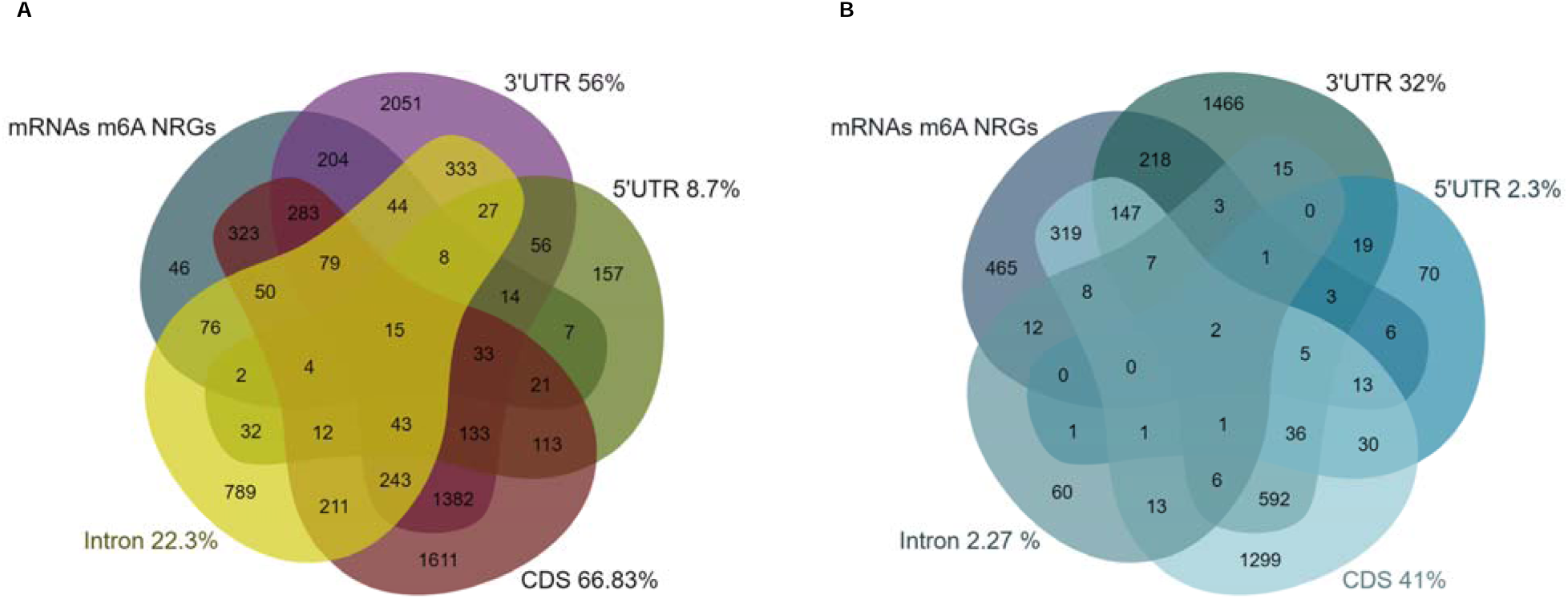
Distribution of m□A sites across NRG mRNAs. **(A)** Estimated proportion of potentially methylated regions within NRGs mRNAs in filtered set of peaks , obtained by intersecting the NRGs transcript list (n = 1798) with m□A peaks identified in a published mouse brain MeRIP-seq dataset across distinct transcript regions: 3′UTR (n = 11245), 5′UTR (n = 1077), CDS (n = 15332), and introns (n = 4259). (**B**) Equivalent analysis using a high-confidence set of mouse brain m□A peaks: 3′UTR (n = 4672), 5′UTR (n = 307), CDS (n = 6079), and introns (n = 263).

### 2. m⁶A-methylation regulatory genes and methylated mRNAs in NRGs and synapses in AD

To conduct a more comprehensive analysis of m⁶A dynamics, we examined the components of the m⁶A regulatory enzymatic complex within the granules. The mRNA content in NRGs was found to include m⁶A writers, including *METTL3* and *ZC3H13*. Furthermore, NRGs exhibited several readers, including *YTHDF2* and *YTHDF3*, factors belonging to the eukaryotic initiation factor *eIF3* family (*eIF3a, eIF3c, eIF3k, eIF3m, eIF2a, eIF2b3, and eIF2s3x*), along with *Prrc2a* and *HNRNPk* mRNAs. At the protein level, two *eIF3* subunits *eIF3b* and *eIF3c*, and four *HNRNP* family proteins *HNRNPc*, *HNRNPr*, *HNRNPm*, and *HNRNPu* were identified as m⁶A readers within NRGs. However, no m⁶A erasers were detected in the granules (Table 1).

**Table 1:**
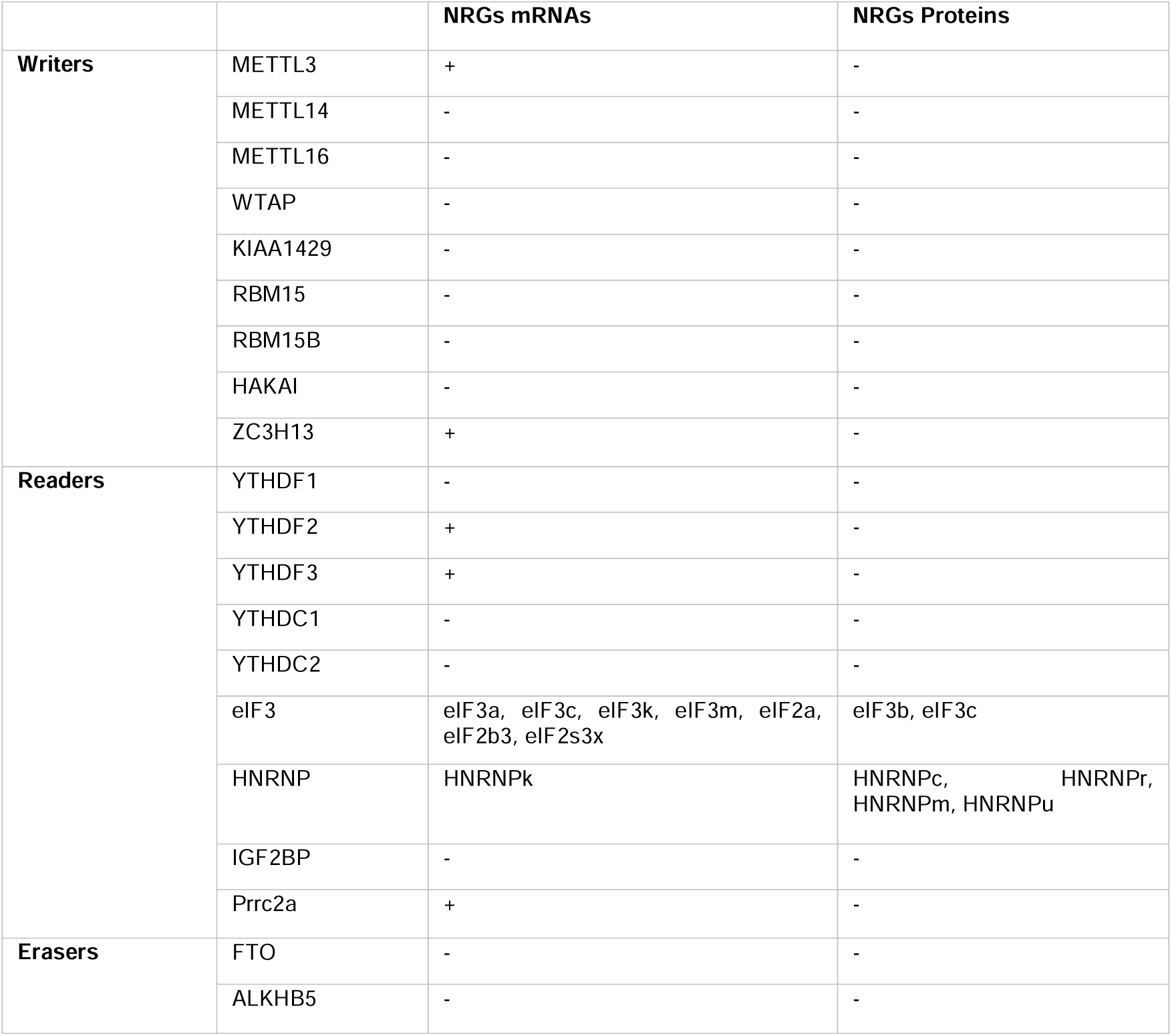
Summary of m□A regulatory complex in both mRNAs and proteins found in mouse NRGs. METTL3/14/16: Methyltransferase like 3/14/16, WTAP: Wilms tumor 1-associating protein, KIAA1429: VIRMA, vir-Like m□A methyltransferase associated, RBM15/15B: RNA binding motifs protein 15/15B, HAKAI: E3 ubiquitin-protein ligase Hakai, ZC3H13: zinc finger CCCH-type containing 13, YTHDF1, 2 and 3: YTH domain-containing family protein 1, 2 and 3, YTHDC1 and 2; YTH Domain Containing 1 and 2. eIF3 (a, b, c, k, and m) : eukaryotic initiation factor 3 subunit a,b,c,k and m, eIF2: eukaryotic initiation factor 2 subunit 2a,2b3, and 2s3x, HNRNP (k, c, r, m, u): heterogeneous nuclear ribonucleoprotein subunit k, c, r, m and u, IGF2BP: Insulin-like growth factor 2 mRNA-binding protein, Prrc2a: proline-rich coiled-coil 2A, FTO: fat mass and obesity-associated protein, ALKBH5: alkB homolog 1. 5. "+": Present, "-" : Absent

**Table 2:** Datasets used in the analysis. Human and mouse profiles derived from RNA-seq and microarray datasets were included in the DEG analysis. GEO accession numbers correspond to publicly available studies archived in the NCBI Gene Expression Omnibus.

|  | RNA-seq | Microarrays |
| --- | --- | --- |
| **Human**<br>**datasets** | GSE174367 | GSE5281 |
|  | GSE159699 | GSE48350 |
|  | GSE163877 | GSE39420 |
|  | GSE125050 | GSE36980 |
|  | GSE118553 |  |
| **Mice**<br>**datasets** | GSE161904 | GSE25926 |
|  | GSE131659 | GSE85162 |
|  | GSE140286 | GSE104249 |
|  | GSE145907 |  |

To further investigate the potential implication of this complex in granules, we subsequently focused on the key components of m⁶A-methylation system (writers, readers, and erasers) and assessed whether their expression might potentially impact NRGs imbalance in AD. We evaluated the expression of these genes using 16 publicly accessible microarray and RNA-seq datasets from both AD patients and mouse models. This comparative analysis enabled us to identify genes that were dysregulated in AD brain relative to controls. Interestingly, our analysis showed that several core m⁶A regulatory genes were consistently altered in 11 human RNA-seq and microarray datasets, as well as in 2 mouse RNA-seq datasets. From the set of dysregulated genes, we selected those most frequently affected across datasets and visualized them graphically (Fig. 3A, B). Our results revealed that *METTL3* is up-regulated, whereas *YTHDF2, YTHDF3, EIF3m*, and the demethylase *FTO* are down-regulated in AD (Fig. 3A, B; Fig. S2, S3; Supplementary Table S2).

**Figure 3:**
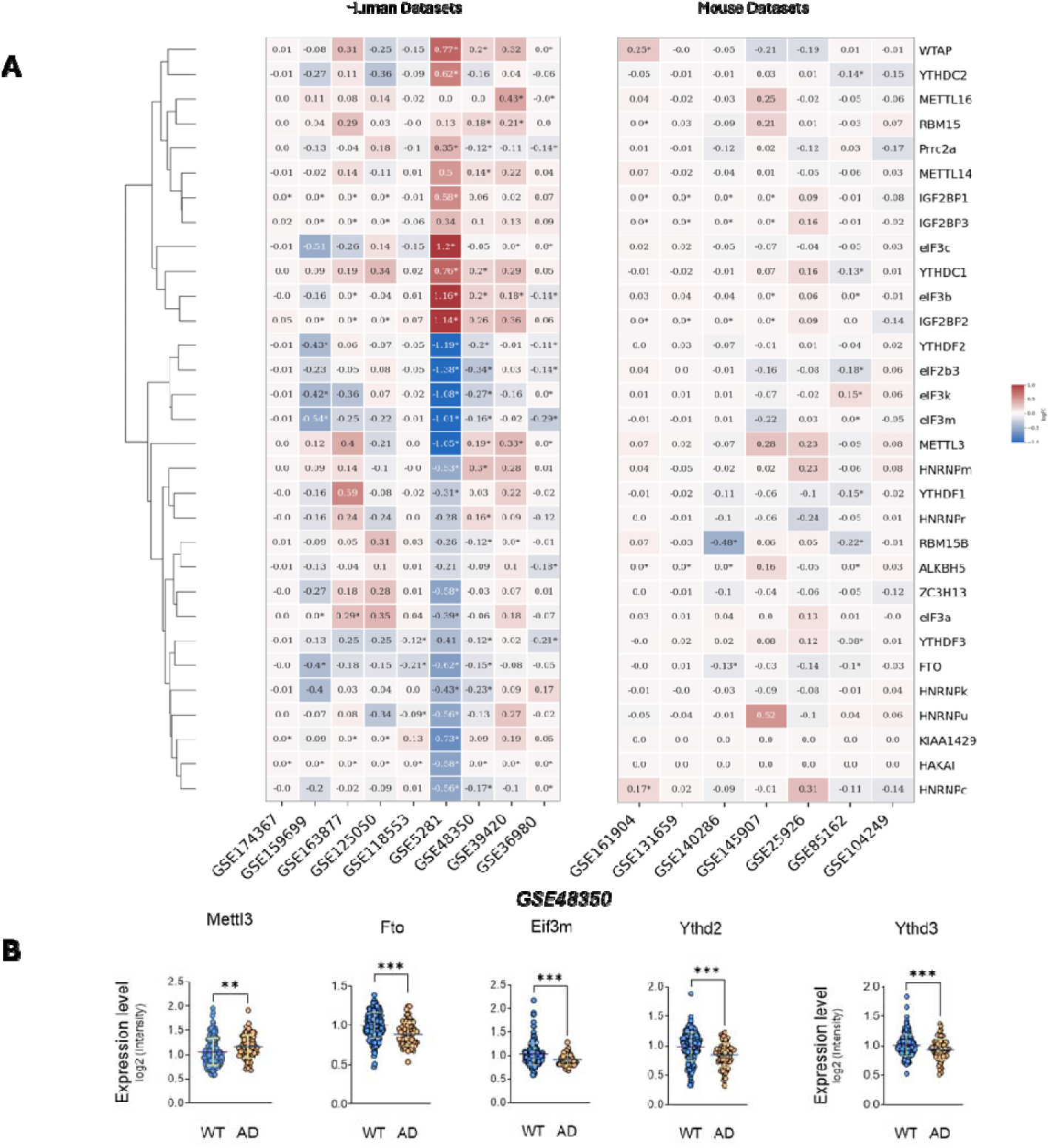
Methylated and m□A regulatory genes in AD. **A)** Heatmap of m□A regulatory genes from human and mice RNA-Seq and microarrays datasets. B) Boxplots of the selected m□A regulatory genes showing their dysregulation in a representative dataset (GSE 48350). Gene expression data shown as log2 (TPM+1) and log2 (intensity) for RNASeq and microarray datasets, respectively. Statistical comparison between AD vs control patients performed by limma workflow (moderated t-statistics). Corrected P-values for multiple testing using the Benjamini-Hochberg (BH) false discovery rate. The asterisks denote statistical significance (* adjP ≤ 0,05; ** adjP ≤ 0.01; *** adjP ≤ 0.001). C and D) UpSet Plots illustrating quantitative intersection of the sets of the most repeated differentially expressed genes between the different datasets in NRGs (C), and Synapses (E) with upregulated and downregulated genes in red and blue, respectively. The numbers above the bars show the number of common genes between the groups of datasets marked below the bars. Top 30 gene ontology (GO) enrichment analysis for genes identified in C and D under biological processes in Granules (D), and Synapses (F). Pathways are first filtered based on FDR cutoff < 0.05, then the top 30 most significant pathways were sorted by Fold Enrichment. ’Remove redundant pathway’ is selected and similar pathways sharing 95% of genes are represented by the most significant pathway. Pathways that are too big or too small are excluded from analysis using the Pathway Size limits.

Next, we wanted to pinpoint methylated genes present within granules or synaptic transcriptomes that are also dysregulated in AD. To accomplish this, we first compiled a list of m⁶A-methylated mRNAs from publicly available datasets ^24,71–74^, yielding 14,448 genes commonly marked by m⁶A. These methylated transcripts were then intersected with differentially expressed genes (DEG) identified across AD datasets and subsequently overlapped with mRNAs detected in NRGs and synapses^70,71^. Through this multi-level integration, we identified 22 dysregulated genes in AD that could be m⁶A - methylated and present in NRGs. The upset plot highlights the genes that were dysregulated in NRGs in at least two different datasets, revealing 13 up-regulated genes and 1 down-regulated gene (Fig. 3C). Following the same approach to analyze synaptic transcriptomes, we identified 39 potentially m⁶A-methylated and dysregulated genes in synapses. Among these, the most recurrent genes included 6 up-regulated and 14 down-regulated transcripts (Fig. 3E). The biological relevance of these dysregulated, methylated transcripts was investigated using gene ontology (GO) analysis^72,73^ . In NRGs, enriched pathways spanned a wide functional range, with prominent terms including hippocampal neuron apoptotic processes and multiple metabolic processes (Fig. 3D, and Supplementary table S3, GO Granules). In the synapses, the most prominently dysregulated m⁶A-modified transcripts were enriched in process central to synaptic physiology, most prominently regulation of synaptic vesicles process, neurotransmitter secretion and transport (Fig. 3F and Supplementary table S3, GO Synapses). Functional enrichment analysis revealed a more detailed biological function spectrum of the methylated dysregulated genes. Among the GO terms overrepresented in granules, hippocampal neuron apoptotic process (FDR< 0.05 and fold enrichment = 364.75), driven by the upregulated transcript Cx3cr1, emerged as the most strongly overrepresented category (Fig. 3C, D, and Supplementary table S3, GO Granules). Additionally, other enriched pathways linked to microglial cell activation process, regulation of protein stability, and immune and metabolic process were found (Fig. 3D), (Supplementary table S3, GO Granules). In the synapse, key ranked dysregulated genes were associated with regulatory pathways such as regulation of synaptic vesicles process (*Cyfip1 Apba2, Brsk1, Tbc1d24* and *Grm7),* neurotransmitter secretion and transport (*Apba2, Brsk1 and Grm7*), as well as associative learning and cognition processes (*Cyfip1, Brsk1, Gabra5 and Grm7)*. Furthermore, other genes were found to be implicated in the regulation of dendrite development (*Stau2, Cyfip1 and Tbc1d24*) (Fig. 3E, F and Supplementary table S3, GO Synapses). Together, these results reveal that m⁶A-modified transcripts dysregulated in AD converge on pathways essential for neuronal survival, synaptic transmission, and higher-order cognitive functions, suggesting that disrupted epitranscriptomic regulation may represent a previously underappreciated mechanism contributing to granule imbalance and synaptic vulnerability in AD.

### 3. NRGs proteins with low complexity and Prion-like domain dysregulation in AD

We next investigated whether NRGs contain proteins with Low-Complexity Domain (LCD) or PrLD, given their known roles in RNA granule assembly and neurodegenerative pathology. By overlapping NRG-associated proteins with previously reported LCD- and PrLD-containing proteins (Supplementary Table S1), we identified a substantial subset of NRG components containing LCDs and/or PrLDs (Additional File 1, and Fig. S4).

To further validate the presence of PrLD-containing proteins within NRGs, we applied the PLAAC algorithm, which confirmed that approximately 9.37% of all NRGs proteins contain PrLD sequences (Fig. 4A; Fig. S5). These included *Caprin1, EIF3c, Elavl2, Elavl4, Hnrnpr, Hnrnpu, ILf3, Npm1, Stau2, Ubap21, Upf1, and Ybx1.* Building on our differential expression analysis, we summarized expression changes of PrLD-containing NRGs proteins in a heatmap (Fig. 4B; Supplementary Table S2). We then highlighted the most recurrently altered genes across datasets, identifying *STAU2* as consistently downregulated and *YBX1* as upregulated in AD brain compared with controls (Fig. 4C).

**Figure 4:**
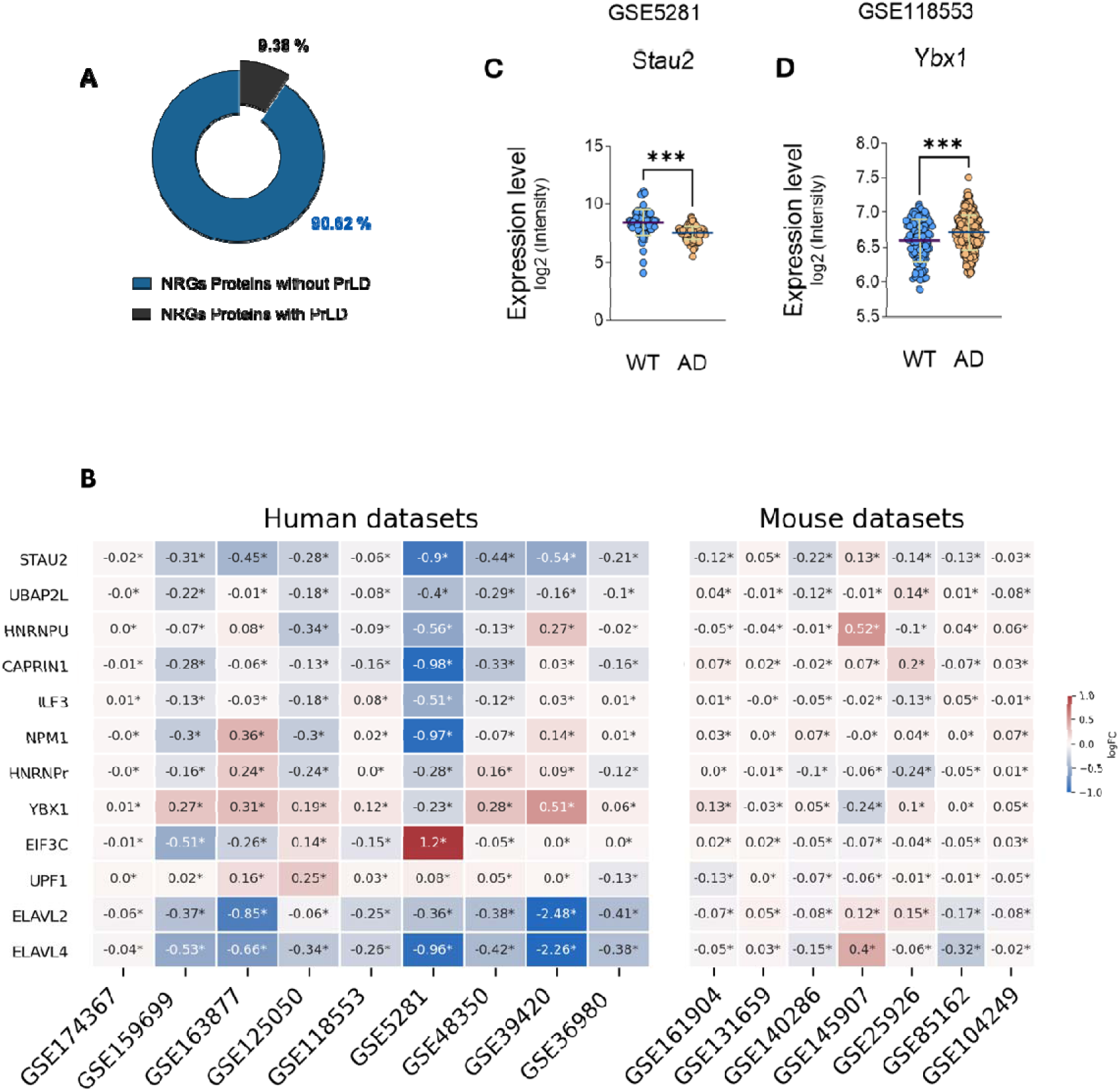
Proteins with PrLD detected in NRGs and their dysregulated gene expression across the studied datasets: **A**) Percentage of proteins containing PrLD in NRGs. **B**) Heatmap of differentiall expressed genes of proteins with PrLD from human and mouse RNA-Seq and microarrays datasets. asterisks denote statistical significance (* adjP ≤ 0,05). **C**) Boxplots of downregulated gene expression of Stau2 in four datasets, and upregulated gene expression of YBX1 in two datasets. Genes’ expression data shown as log2 (TPM+1) and log2(intensity) for RNASeq and microarray datasets, respectively. Statistical comparison between AD and WT (control) samples was performed by limma workflow (moderated t-statistics). P-values were corrected for multiple testing using the Benjamini-Hochberg (BH) false discovery rate. The asterisks denote statistical significance (* adjP ≤ 0,05; ** adjP ≤ 0.01; *** adjP ≤ 0.001).

## Discussion

NRGs are heterogeneous assemblies of RNAs, RBPs, and translation factors that travel as translationally repressed cargo to synapses, where specific cues reactivate translation^7,43^. However, the mechanisms determining which mRNAs are selectively packed into these granules remain poorly understood. To investigate this, we studied RNA modifications within NRGs, focusing on m⁶A as the most abundant internal RNA modification in eukaryotes implicated in regulating neuronal growth, memory, and synaptic plasticity ^34,39,74–79^, and its dysregulation is associated with neurodegenerative diseases, particularly AD ^23,69,80–83^. Here, we identify m⁶A-modified mRNAs in NRGs and synapses that show altered regulation in AD, suggesting that disrupted m⁶A homeostasis ^41,42,50,51,69,84^ may serve as a biomarker of AD pathology. Our findings show that m⁶A is a dominant modification in NRG-associated mRNAs, supporting its role in selective mRNAs recruitment during granule formation.

The identified methylated and dysregulated genes are involved in several biological processes ranging from cell death control, immune system modulation, synaptic remodeling, vesicles trafficking, neurotransmission ^144 145 146 147–150 151–153 154,154–159^, neuronal proliferation, dynamic plasticity ^160–164^, and pathways influencing Aβ and aggregation in AD brains^165–171^. Dysregulated m⁶A homeostasis across brain regions ^34,106–110^, may therefore track of AD progression. Our findings indicate that m⁶A-methylation stands out as a predominant modification in the mRNAs present in NRGs, highlighting the possibility that this methylation may play a crucial role in the selection process of mRNAs that form NRGs transported to the synapses. Prior studies have shown that m⁶A within coding sequences can slow cognate tRNA selection during translation. In Parallel, SG, another class of RNA granule implicated in AD have been reported to preferentially accumulate m⁶A-modified transcripts^45,111,112^

m_6_A not only promotes SG enrichment, but also influences their dynamics via m6A readers such as YTHDF2, correlates with reduced translation efficiency and the preferential inclusion of long transcripts^113–115^. Together, these observations underscore the critical role of m_6_A in shaping RNA granule composition. Consistent with this, we observed elevated m_6_A density within the CDS, and 3′UTR associated with NRGs. m_6_A sites proximal to the stop codon appear particularly relevant , as the 3′UTR is a key regulatory hub controlling alternative polyadenylation, mRNA localization, stability, and translation^26,116,117^. Our data suggest that m_6_A deposition within the 3′UTR, particularly near the termination codon, may contribute to ribosome stalling and thereby promote NRGs formation, as m_6_A sites near the stop codon may have regulatory significance, by regulating alternative polyadenylation, localization, stability, and translation^44,116,118^.

In this study, we demonstrate a disruption of m_6_A pathway homeostasis in AD by analyzing expression changes in key enzymes and effectors involved in this modification. Specifically, we observed upregulation of the m_6_A methyltransferase METTL3, concomitant with downregulation of the demethylase FTO, together with a marked reduction in the m_6_A readers YTHDF2, YTHDF3, and eIF3m. Collectively, these alterations are predicted to increase global m_6_A methylation levels, consistent with previous reports in AD patient samples and APP/PS1 mouse models^108,119–121^. Indeed, in these mouse models, elevated m_6_A levels correlate with increased METTL3 and reduced FTO expression in the cortex and hippocampus, supporting a contributory role for m_6_A dysregulation in AD pathogenesis.

The downregulation of YTHDF2 and YTHDF3 may further exacerbate this imbalance by impairing the clearance and translational regulation of m⁶A-modified transcripts. Loss of YTHDF proteins has been shown to result in cytosolic accumulation of hypermethylated mRNAs, elevated global m_6_A levels, and reduced translation efficiency^122–124^,. Under physiological conditions, YTHDF family members facilitate the turnover of m⁶A-marked mRNAs, while YTHDF3 additionally promotes cap-dependent translation, potentially through interactions with 3′UTR methylation^125,126^. Notably, YTHDF2 is also essential for synapse formation, as YTHDF2-deficient neurons fail to establish normal synaptic structures^127,128^. Recent studies showed that, unlike non-methylated transcripts, polymethylated mRNAs promote SG formation via liquid–liquid phase separation (LLPS) through interactions with *YTHDF* proteins. *YTHDF2’s* ability to undergo LLPS depends on its low-complexity domain, and deletion of this domain abolishes LLPS^59,114,129^. Polymethylated transcripts are enriched in SG and P-bodies, and transcripts containing four or more m_6_A sites exhibit reduced translation efficiency in wild-type cells compared to m⁶A-deficient cells^59,114,129,130^, indicating selective translational repression linked to phase-separation behavior.

In parallel, the observed downregulation of eIF3m may have important consequences for neuronal proteostasis and synaptic integrity. eIF3m is an integral subunit of the eIF3 complex, which plays a central role in translation initiation and ribosome recruitment to mRNAs ^131–133^. Given the dependence of neurons on precisely regulated protein synthesis, particularly for synaptic maintenance and plasticity, reduced eIF3m may compromise the stability or efficiency of the eIF3 complex, thereby aggravating the translational deficits widely reported in AD brains^134^. In this context, decreased eIF3m expression may limit the pool of functional ribosomes available for local protein synthesis. Together, the coordinated downregulation of m_6_A readers, including YTHDF2, YTHDF3, and eIF3m, suggests a convergent mechanism by which altered m_6_A recognition, impaired mRNA turnover, and defective translation initiation contribute to RNA granule dysregulation, synaptic dysfunction, and neuronal vulnerability in AD.

Altogether, the upregulation of *METTL3* combined with *FTO* downregulation confirmed across multiple AD datasets likely drives increased mRNA polymethylation, including transcripts transported within granules. This may contribute to impaired translation and enhanced LLPS, potentially exacerbating AD-related cellular dysfunction. In addition, our analysis showed downregulation of *eIF3m* in AD brains, suggesting reduced translation efficiency and local protein synthesis within NRGs. m_6_A within the CDS has been shown to induce ribosome pausing, slowing elongation and altering translation dynamics^36,111,135^. *METTL3* enhances translation independently of its catalytic activity by recruiting *eIF3* to the initiation complex^127,136^, likely at synapses, and also co-regulates splicing and miRNA processing with *HNRNPA2B1* and *ZC3H13*, which anchors the methyltransferase complex in the nucleus. m⁶A’s effects on translation are context-dependent, with both enhancement and repression^137,138^. Given that m⁶A’s impact varies by transcript region, further studies are needed to explore how m_6_A influences localized translation in NRGs. Other study report that m_6_A can either enhance or repress translation efficiency and protein synthesis^139–141^, and because methylation at different transcript regions may exert distinct effects, it is essential to investigate the regional impact of m_6_A on translation, particularly within NRGs in neurites and synapses.

*FMRP*, an RBP enriched in NRGs widely used as a key granule marker, was recently identified as a potential m_6_A reader. It is known to suppress translation of its target mRNAs by blocking ribosome translocation^142–144^. Recent studies further show that *FMRP* binds m_6_A sites to promote the export of m⁶A-modified transcripts from the nucleus, and that *FMRP* depletion increases nuclear m_6_A levels^142,145^. Consistent with these findings, our analysis indicates that many *FMRP* targets within NRGs are highly m⁶A-methylated. Indeed, more than 95% of *FMRP*-associated mRNAs in the cerebellum and cortex carry m_6_A modifications^67^, and FMRP shows a strong preference for m⁶A-modified transcripts. PAR-CLIP and m⁶A-IP studies further reveal that FMRP-binding peaks at m_6_A sites are enriched within the CDS and 3′UTR^142^. m_6_A is mainly deposited at the RRm⁶ACH consensus sequence, closely matching FMRP-binding motifs^146,147^. This observation suggests that many FMRP-targeted mRNAs carry m⁶A-modified, highlighting their potential roles in NRGs formation and transport as well as synaptic protein synthesis. Dysregulation of these processes may, in turn, contribute to the synaptic dysfunction and neuronal degeneration characteristic of AD ^148,149^.Our DEG analysis highlights the dysregulation of PrLD-containing proteins, notably *STAU2* and *YBX1*. *STAU2*, another key NRGs marker, regulates dendritic RNA localization and synaptic plasticity by interacting with over 1,200 RNAs^150^, and is essential for neurogenesis and neuronal differentiation^151–153^. Loss of *STAU2* alters the neuronal transcriptome, reducing short 3′UTR transcripts while increasing long 3′UTR transcripts, particularly those linked to synaptic function and memory, and induces actin cytoskeleton reorganization, fewer mature dendritic spines, and decreased synapse numbers^98,150^. These findings suggest *STAU2’s* critical role in spatial gene regulation at synapses and possibly in NRGs, aligning with synaptic deficits observed in AD. *YBX1*, a DNA/RNA-binding protein involved in transcription, translation, mRNA processing, and DNA repair,^154^ has been shown to improve AD phenotypes in 5XFAD mice^155^, it is also implicated in liquid-liquid phase separation and SG assembly^156,157^. Our analysis shows *YBX1* upregulation in AD, suggesting a role in granule formation and PrLD-mediated RNA-protein interactions within NRGs. Such dysregulated interactions may promote aggregation, impair axonal transport, and exacerbate AD pathology by compromising neuronal function. Although our findings indicate clear dysregulation of PrLD-containing proteins in NRGs, their precise role within these granules remains unresolved and requires further experimental investigation. Overall, our findings support a model in which dysregulated m⁶A-modified transcripts and PrLD-containing proteins in NRGs could contribute to AD pathogenesis (Fig. 5). The strong conservation of these genes across humans, mice, and rats (Fig. S6) suggests shared functional roles across species, yet experimental studies will be necessary to determine their precise contributions to disease onset and progression.

**Figure 5:**
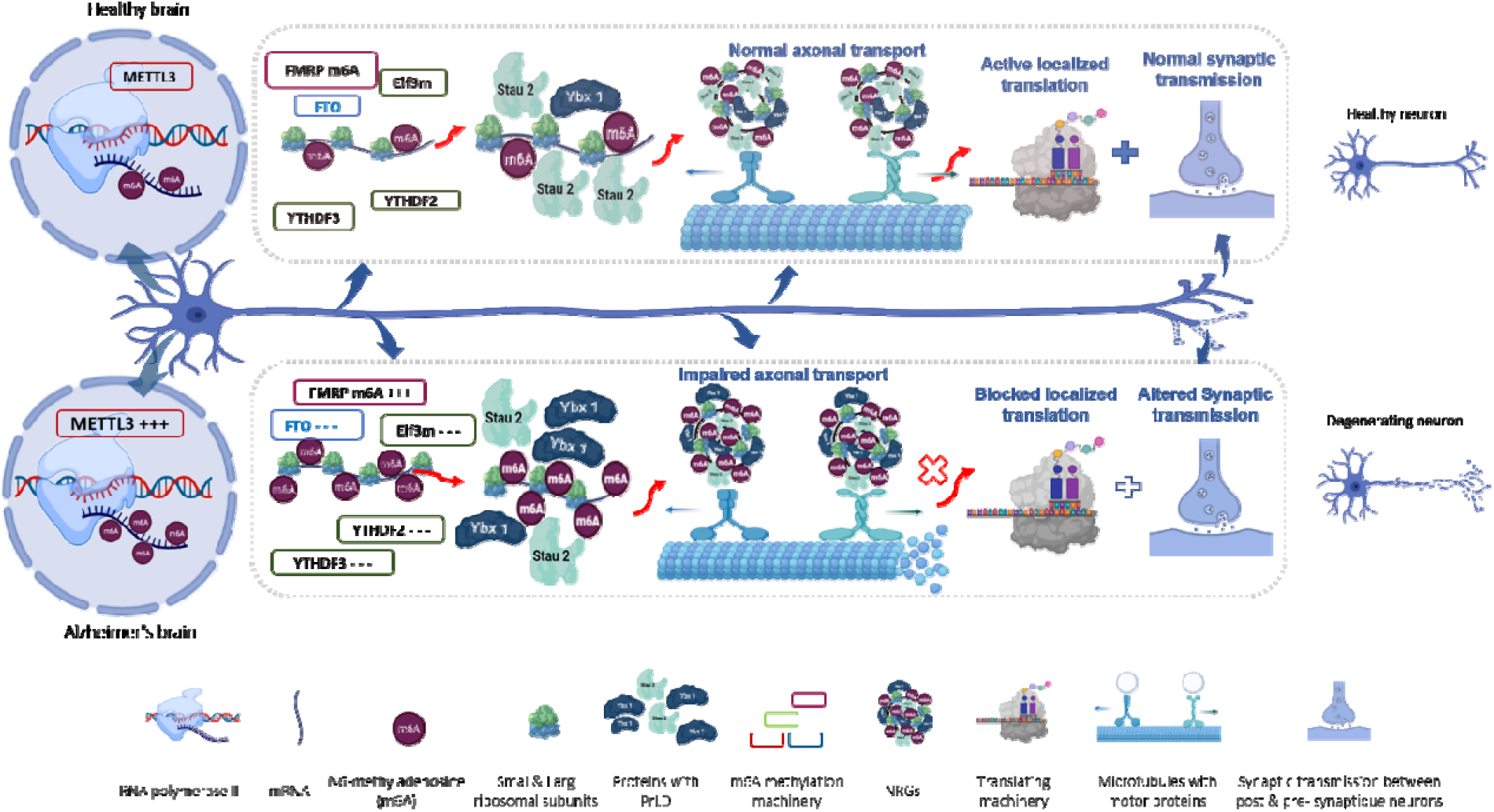
Hypothetical model for NRGs dysregulation in AD. Top: In healthy neurons, balanced m□A methylation machinery (METTL3, FTO, FMRP, YTHDF2/3, eIF3m), together with PrLD-containing RNA-binding proteins such as STAU2, and YBX1, ensures proper NRGs function, axonal transport, and activity-dependent local translation, thereby supporting normal synaptic function. Bottom: In AD, increased METTL3 and reduced FTO, YTHDF2/3, and eIF3m levels lead to hypermethylated mRNAs, altered translation and RBP interactions, and aberrant NRGs composition. These changes impair axonal transport and local translation, resulting in synaptic dysfunction and neuronal degeneration.

## Conclusion

AD, the most common form of dementia, is characterized by β-amyloid and tau accumulation, accompanied by profound defects in axonal transport and synaptic functions. NRGs are microscopic membrane-less structures composed of RBPs, RNAs, and components of the translation machinery, thereby supporting neural development, maintenance, and synaptic plasticity. In this study, we propose that impaired NRGs formation, assembly and transport represent a previously underappreciated mechanism contributing to AD. Although NRGs are structurally heterogeneous and their assembly remains poorly understood, our findings highlight the key role of m⁶A-methylation and PrLD-containing proteins as central regulators of NRGs formation and dynamics. Mechanistically, METTL3 catalyzes the co-transcriptional deposition of m_6_A on mRNA transcripts in the nucleus in association with RNA polymerase II. Increased METTL3 expression combined with downregulation of the m_6_A demethylase FTO results in global mRNA hypermethylation. m⁶A-modified transcripts are exported to the cytoplasm through interactions with FMRP. Reduced expression of YTHDF2, YTHDF3, and eIF3m impairs m⁶A-dependent mRNA decay and translation, leading to cytosolic accumulation of hypermethylated transcripts and their translational repression. Through interactions with FMRP and additional m_6_A readers, these transcripts become sequestered within NRGs, forming enlarged granules that fail to undergo efficient distal axonal transport. In parallel, dysregulation of proteins containing PrLD, including YBX1 and STAU2, further disrupts granule dynamics. Downregulation of STAU2 impairs RNA granule trafficking to distal neuronal compartments, while upregulation of YBX1, together with the somatic accumulation of hypermethylated transcripts, promotes stalled granule formation.

Collectively, impaired granule formation and transport and altered local translation compromise synaptic maintenance and function, ultimately contributing to synaptic dysfunction and the development of AD pathology (Fig.5). Overall, our findings point to m⁶A-associated NRGs and PrLD dysregulation as a potential contributing factor in AD and emphasize the need for further experimental investigations to clarify their mechanistic and therapeutic relevance.

## Methods

### mRNA m⁶A-methylation enrichment in different structures

As there is a lack of studies characterizing m⁶A-methylation within NRGs, to obtain the estimation of percentages of m⁶A-modified transcripts in these granules in comparison with other structures and other organelles, we performed an overlap-based comparative analysis using published mRNAs lists. We leveraged lists of transcripts from mouse granules (NRGs, n=1798)^7^, rat dendritic/axonic synaptic neuropil (RN, n=2550)^71^, mouse hippocampal neurons (MHCN, n= 2160)^158^, and we overlapped them with lists of mRNAs datasets of experimentally validated m_6_A -methylated mRNA from several studies: mouse embryonic stem cells, study1 (m_6_A mESC1, n=1455)^59^, and study 2 (mESC 2, n=5213)^60^, Mouse adult neuronal stem cells in proliferation and differentiation phases: (mNSC Proliferation, n=9206 and mNSC Differentiation, n=7336)^61^, mouse embryonic fibroblast cells (mEFC, n=15453)^62^, and human embryonic stem cells (hESC , n=7530)^60^. Overlap results were then quantified and visualized using Venn diagrams (Supplementary table S1). From these overlaps, percentage of methylated mRNAs in each structure was estimated and plotted. As a comparison to m⁶A, pseudouridine-modified transcripts from mouse brain (Ψ MB, n= 1313) ^63^ were similarly overlapped with NRGs, RN and MHCN mRNAs datasets (Fig. S1).

The same analytical workflow was followed to obtain the percentage of methylated mRNAs in FMRP targets. FMRP mRNAs from adult mouse brain polyribosomes (MBP FMRP, n=842)^64^, mouse brain (MB FMRP, n=443)^159^, and embryonic mouse cerebral cortex (EmCrtx FMRP, n=856)^66^, were overlapped with m_6_A methylated mRNAs derived from mESC1, mESC 2, mNSC Proliferation, mNSC Differentiation, mEFC, hESC and lists of FMRP m_6_A methylated mRNAs from mouse cerebral cortex (m_6_A mCrtx, n=811) ^61^. Rat and Human datasets used in this analysis were converted to their mouse orthologs to ensure cross-species comparison.

### m_6_A peaks localization and gene ontology analysis

To determine the localization of m_6_A peaks within NRG-associated mRNAs, we intersected the list of this latter with list from a study ^49^ that identify regions that contains m_6_A Peaks localization across the whole transcriptome. Two separate analyses were conducted using two different lists of genes as background for the mouse brain data set:

1. the list of genes expressed in all MeRIP, and non-IP samples combined and 2) a list of random genes taken from the mouse transcriptome.

Gene ontology analysis of the genes coding for NRGs mRNAs was performed using the bioinformatic database ShinyGO^160,161^ to detect involved biological process, cellular components and molecular function filtered based on the false discovery rate (FDR) cutoff < 0.05, the pathways were sorted by fold enrichment. All GO results are summarized in Supplementary table S3. Analysis for DEGs and m_6_A methylated genes present in granules and synapses was performed using the same ShinyGO^160,161^ to detect involved biological process. FDR cutoff (FDR < 0.05), the pathways were sorted by fold enrichment. All GO results are summarized in Supplementary table S3. GO Analysis for cellular component, biological process, and molecular function were performed at default settings.

To identify mRNAs that are m⁶A-methylated in granules and synapses, 5 data sets of m_6_A -IP in the brain were overlapped to obtain the genes that are m⁶A-methylated in the brain. These genes are overlapped again with the lists of mRNAs present in the granules and synapses, to obtain the subsets of transcripts that are m⁶A-methylated in the granules and synapses (Supplementary table S3). These mRNAs were run against DEGs results (AD vs Control) to pinpoint m_6_A -methylated mRNAs in NRGS and synapses that are consistently dysregulated in AD patients and animal models. The upset plots show the intersections of these overlaps between at least 2 studies. Plots were generated using UpsetR package. The total size of each set is represented on the left barplot. Every possible intersection is represented by the bottom plot, and their occurrence is shown on the top barplot.

#### Datasets collection and differentially expressed genes (DEG) analysis

Datasets used for differential expression analysis were obtained from NCBI Gene Expression Omnibus (GEO) repository. We included Human and mouse datasets containing AD and control samples, derived from both RNAseq and microarray plateforms. The datasets are summarized in the table below.

The [LIMMA](https://academic.oup.com/nar/article/43/7/e47/2414268) ^162^, and [EdgeR](https://academic.oup.com/bioinformatics/article/26/1/139/182458?login=true)^164^ packages were used for filtering and identification of differentially expressed genes (DEGs) in Micro-array and RNA-seq datasets, respectively. R package “[ggplot2](https://pmc.ncbi.nlm.nih.gov/articles/PMC6513982/#B50)” ^163^, and [VolcanoseR](https://pubmed.ncbi.nlm.nih.gov/33239692/) ^165^were used to visualize the results. DEGs in each dataset were visualized through Volcano-plots depicting the upregulated and downregulated expressed m⁶A-methylation genes and proteins with PrLD in human (Fig. S2), and mouse (Fig. S3). The significant DEGs were selected using an adjusted P value < 0.05 (Supplementary table S2).

### Proteins with PrLD sequences detection

To detect proteins with PrLD present in NRGs, the web application prion-like amino acid composition (PLAAC) based Hidden Markov Model (HMM) algorithm was used^166^. Each NRGs protein sequence was submitted to PLAAC in FASTA format (File 2, supplementary datas).

### Sequences alignment

The Homo sapiens, Mus musculus, and Rattus norvegicus sequences of the genes of interest are obtained first using NCBI for each gene. The sequences were aligned to obtain multiple protein alignment. Coverage and percentage identity values are indicated in the “cov” and “pid” columns respectively (fig. S6).

## Supporting information

Supplementary datas

## Acknowledgment

This work was supported by UM6P University and the OCP Foundation. We extend our gratitude to our colleagues at the UM6P Faculty of Medical Sciences for their valuable contributions, especially Drs. Soukaina Hakam and Taha Mohamed Moutaouafik.

## Declaration of Competing Interest

The authors declare that they have no known competing financial interests or personal relationships that could have appeared to influence the work reported in this paper.

## Declaration of Generative AI and AI-assisted technologies in the writing process

During the preparation of this work, the authors used ChatGPT in order to correct spelling/grammar and improve readability and language. After using this tool/service, the authors reviewed and edited the content as needed and took full responsibility for the content of the publication.

## Authorship Contribution Statement

REL conceived and oversaw the project. SB, HA, AE, and REL collected and/or critically analyzed the data. SB drafted the initial version of the manuscript and prepared the figures with the input from all authors. YS provided intellectual contribution to the manuscript. All authors read, reviewed, and approved the final manuscript.

## Ethics declaration

Not applicable

## Notes

### Competing Interest Statement

The authors have declared no competing interest.

## References

1. DeTure, M.A., and Dickson, D.W. (2019). The neuropathological diagnosis of Alzheimer’s disease. Molecular Neurodegeneration 14, 32. 10.1186/s13024-019-0333-5.

2. Knopman, D.S., Amieva, H., Petersen, R.C., Chételat, G., Holtzman, D.M., Hyman, B.T., Nixon, R.A., and Jones, D.T. (2021). Alzheimer disease. Nat Rev Dis Primers 7, 1–21. 10.1038/s41572-021-00269-y.

3. Golde, T.E. (2022). Alzheimer’s disease – the journey of a healthy brain into organ failure. Molecular Neurodegeneration 17, 18. 10.1186/s13024-022-00523-1.

4. Long, J.M., and Holtzman, D.M. (2019). Alzheimer Disease: An Update on Pathobiology and Treatment Strategies. Cell 179, 312–339. 10.1016/j.cell.2019.09.001.

5. Salvadores, N., Gerónimo-Olvera, C., and Court, F.A. (2020). Axonal Degeneration in AD: The Contribution of Aβ and Tau. Frontiers in Aging Neuroscience 12.

6. Shiina, N., Shinkura, K., and Tokunaga, M. (2005). A Novel RNA-Binding Protein in Neuronal RNA Granules: Regulatory Machinery for Local Translation. J Neurosci 25, 4420–4434. 10.1523/JNEUROSCI.0382-05.2005.

7. El Fatimy, R., Davidovic, L., Tremblay, S., Jaglin, X., Dury, A., Robert, C., De Koninck, P., and Khandjian, E.W. (2016). Tracking the fragile X mental retardation protein in a highly ordered neuronal ribonucleoparticles population: a link between stalled polyribosomes and RNA granules. PLoS genetics 12, e1006192.

8. Khandjian, E., Dury, A., Koninck, P., El fatimy, R., and Davidovic, L. (2015). RNA Granules: Functions Within Presynaptic Terminals and Postsynaptic Spines. In Reference Module in Biomedical Sciences 10.1016/B978-0-12-801238-3.04756-5.

9. Ainger, K., Avossa, D., Morgan, F., Hill, S.J., Barry, C., Barbarese, E., and Carson, J.H. (1993). Transport and localization of exogenous myelin basic protein mRNA microinjected into oligodendrocytes. J Cell Biol 123, 431–441. 10.1083/jcb.123.2.431.

10. Kipper, K., Mansour, A., and Pulk, A. (2022). Neuronal RNA granules are ribosome complexes stalled at the pre-translocation state. Journal of Molecular Biology 434, 167801. 10.1016/j.jmb.2022.167801.

11. Costa Cruz, P.H., and Kawahara, Y. (2021). RNA Editing in Neurological and Neurodegenerative Disorders. Methods Mol Biol 2181, 309–330. 10.1007/978-1-0716-0787-9_18.

12. Jonkhout, N., Tran, J., Smith, M.A., Schonrock, N., Mattick, J.S., and Novoa, E.M. (2017). The RNA modification landscape in human disease. RNA 23, 1754–1769. 10.1261/rna.063503.117.

13. Noack, F., and Calegari, F. (2018). Epitranscriptomics: A New Regulatory Mechanism of Brain Development and Function. Front Neurosci 12, 85. 10.3389/fnins.2018.00085.

14. Wang, D.O. (2020). RNA Modifications in the Central Nervous System. In The Oxford Handbook of Neuronal Protein Synthesis.

15. Levanon, E.Y., Eisenberg, E., Yelin, R., Nemzer, S., Hallegger, M., Shemesh, R., Fligelman, Z.Y., Shoshan, A., Pollock, S.R., and Sztybel, D. (2004). Systematic identification of abundant A-to-I editing sites in the human transcriptome. Nature biotechnology 22, 1001–1005.

16. Delatte, B., Wang, F., Ngoc, L.V., Collignon, E., Bonvin, E., Deplus, R., Calonne, E., Hassabi, B., Putmans, P., and Awe, S. (2016). Transcriptome-wide distribution and function of RNA hydroxymethylcytosine. Science 351, 282–285.

17. Akichika, S., Hirano, S., Shichino, Y., Suzuki, T., Nishimasu, H., Ishitani, R., Sugita, A., Hirose, Y., Iwasaki, S., and Nureki, O. (2019). Cap-specific terminal N6-methylation of RNA by an RNA polymerase II–associated methyltransferase. Science 363.

18. Sun, H., Zhang, M., Li, K., Bai, D., and Yi, C. (2019). Cap-specific, terminal N 6-methylation by a mammalian m 6 Am methyltransferase. Cell research 29, 80–82.

19. Zhang, L.-S., Liu, C., Ma, H., Dai, Q., Sun, H.-L., Luo, G., Zhang, Z., Zhang, L., Hu, L., and Dong, X. (2019). Transcriptome-wide mapping of internal N7-methylguanosine methylome in mammalian mRNA. Molecular cell 74, 1304–1316.

20. Carlile, T.M., Rojas-Duran, M.F., Zinshteyn, B., Shin, H., Bartoli, K.M., and Gilbert, W.V. (2014). Pseudouridine profiling reveals regulated mRNA pseudouridylation in yeast and human cells. Nature 515, 143–146.

21. Jung, Y., and Goldman, D. (2018). Role of RNA modifications in brain and behavior. Genes, Brain and Behavior 17, e12444.

22. Zhang, Y., Lu, L., and Li, X. (2022). Detection technologies for RNA modifications. Exp Mol Med 54, 1601–1616. 10.1038/s12276-022-00821-0.

23. Dominissini, D., Moshitch-Moshkovitz, S., Schwartz, S., Salmon-Divon, M., Ungar, L., Osenberg, S., Cesarkas, K., Jacob-Hirsch, J., Amariglio, N., and Kupiec, M. (2012). Topology of the human and mouse m 6 A RNA methylomes revealed by m 6 A-seq. Nature 485, 201–206.

24. Desrosiers, R., Friderici, K., and Rottman, F. (1974). Identification of methylated nucleosides in messenger RNA from Novikoff hepatoma cells. Proc Natl Acad Sci U S A 71, 3971–3975. 10.1073/pnas.71.10.3971.

25. Chatterjee, B., Shen, C.-K.J., and Majumder, P. (2021). RNA Modifications and RNA Metabolism in Neurological Disease Pathogenesis. Int J Mol Sci 22, 11870. 10.3390/ijms222111870.

26. Meyer, K.D., Saletore, Y., Zumbo, P., Elemento, O., Mason, C.E., and Jaffrey, S.R. (2012). Comprehensive Analysis of mRNA Methylation Reveals Enrichment in 3′ UTRs and near Stop Codons. Cell 149, 1635–1646. 10.1016/j.cell.2012.05.003.

27. Patil, D.P., Chen, C.-K., Pickering, B.F., Chow, A., Jackson, C., Guttman, M., and Jaffrey, S.R. (2016). m 6 A RNA methylation promotes XIST-mediated transcriptional repression. Nature 537, 369–373.

28. Roundtree, I.A., Evans, M.E., Pan, T., and He, C. (2017). Dynamic RNA modifications in gene expression regulation. Cell 169, 1187–1200. 10.1016/j.cell.2017.05.045.

29. Yue, Y., Liu, J., and He, C. (2015). RNA N6-methyladenosine methylation in post-transcriptional gene expression regulation. Genes & development 29, 1343–1355.

30. Jia, G., Fu, Y., Zhao, X., Dai, Q., Zheng, G., Yang, Y., Yi, C., Lindahl, T., Pan, T., and Yang, Y.-G. (2011). N 6-methyladenosine in nuclear RNA is a major substrate of the obesity-associated FTO. Nature chemical biology 7, 885–887.

31. Rana, A.K., and Ankri, S. (2016). Reviving the RNA World: An Insight into the Appearance of RNA Methyltransferases. Front Genet 7, 99. 10.3389/fgene.2016.00099.

32. Berlivet, S., Scutenaire, J., Deragon, J.-M., and Bousquet-Antonelli, C. (2019). Readers of the m6A epitranscriptomic code. Biochimica et Biophysica Acta (BBA) - Gene Regulatory Mechanisms 1862, 329–342. 10.1016/j.bbagrm.2018.12.008.

33. Yang, Y., Hsu, P.J., Chen, Y.-S., and Yang, Y.-G. (2018). Dynamic transcriptomic m6A decoration: writers, erasers, readers and functions in RNA metabolism. Cell Res 28, 616–624. 10.1038/s41422-018-0040-8.

34. Jiang, X., Liu, B., Nie, Z., Duan, L., Xiong, Q., Jin, Z., Yang, C., and Chen, Y. (2021). The role of m6A modification in the biological functions and diseases. Sig Transduct Target Ther 6, 1–16. 10.1038/s41392-020-00450-x.

35. Liu, Z.-X., Li, L.-M., Sun, H.-L., and Liu, S.-M. (2018). Link Between m6A Modification and Cancers. Front. Bioeng. Biotechnol. 6. 10.3389/fbioe.2018.00089.

36. Meyer, K.D., Patil, D.P., Zhou, J., Zinoviev, A., Skabkin, M.A., Elemento, O., Pestova, T.V., Qian, S.-B., and Jaffrey, S.R. (2015). 5′ UTR m6A promotes cap-independent translation. Cell 163, 999–1010.

37. Xiao, W., Adhikari, S., Dahal, U., Chen, Y.-S., Hao, Y.-J., Sun, B.-F., Sun, H.-Y., Li, A., Ping, X.-L., and Lai, W.-Y. (2016). Nuclear m6A reader YTHDC1 regulates mRNA splicing. Molecular cell 61, 507–519.

38. Yen, Y.-P., and Chen, J.-A. (2021). The m6A epitranscriptome on neural development and degeneration. J Biomed Sci 28, 40. 10.1186/s12929-021-00734-6.

39. Deng, J., Chen, X., Chen, A., and Zheng, X. (2022). m6A RNA methylation in brain injury and neurodegenerative disease. Front Neurol 13, 995747. 10.3389/fneur.2022.995747.

40. Zhang, Y., Zhang, S., Shi, M., Li, M., Zeng, J., and He, J. (2022). Roles of m6A modification in neurological diseases. Zhong Nan Da Xue Xue Bao Yi Xue Ban 47, 109–115. 10.11817/j.issn.1672-7347.2022.200990.

41. Xiong, X., Hou, L., Park, Y., Molinie, B., Gregory, R.I., and Kellis, M. (2021). Genetic drivers of m6A methylation in human brain, lung, heart and muscle. Nat Genet 53, 1156–1165. 10.1038/s41588-021-00890-3.

42. Fan, Y., Lv, X., Chen, Z., Peng, Y., and Zhang, M. (2023). m6A methylation: Critical roles in aging and neurological diseases. Frontiers in Molecular Neuroscience 16.

43. Bushkin, G.G., Pincus, D., Morgan, J.T., Richardson, K., Lewis, C., Chan, S.H., Bartel, D.P., and Fink, G.R. (2019). m6A modification of a 3′ UTR site reduces RME1 mRNA levels to promote meiosis. Nat Commun 10, 3414. 10.1038/s41467-019-11232-7.

44. Yue, Y., Liu, J., Cui, X., Cao, J., Luo, G., Zhang, Z., Cheng, T., Gao, M., Shu, X., and Ma, H. (2018). VIRMA mediates preferential m 6 A mRNA methylation in 3′ UTR and near stop codon and associates with alternative polyadenylation. Cell discovery 4, 1–17.

45. Jain, S., Koziej, L., Poulis, P., Kaczmarczyk, I., Gaik, M., Rawski, M., Ranjan, N., Glatt, S., and Rodnina, M.V. (2023). Modulation of translational decoding by m6A modification of mRNA. Nat Commun 14, 4784. 10.1038/s41467-023-40422-7.

46. Mao, Y., Dong, L., Liu, X.-M., Guo, J., Ma, H., Shen, B., and Qian, S.-B. (2019). m6A in mRNA coding regions promotes translation via the RNA helicase-containing YTHDC2. Nat Commun 10, 5332. 10.1038/s41467-019-13317-9.

47. Meyer, K.D. (2019). m6A-Mediated Translation Regulation. Biochim Biophys Acta Gene Regul Mech 1862, 301–309. 10.1016/j.bbagrm.2018.10.006.

48. Ke, S., Alemu, E.A., Mertens, C., Gantman, E.C., Fak, J.J., Mele, A., Haripal, B., Zucker-Scharff, I., Moore, M.J., and Park, C.Y. (2015). A majority of m6A residues are in the last exons, allowing the potential for 3′ UTR regulation. Genes & development 29, 2037–2053.

49. Meyer, K.D., Saletore, Y., Zumbo, P., Elemento, O., Mason, C.E., and Jaffrey, S.R. (2012). Comprehensive Analysis of mRNA Methylation Reveals Enrichment in 3′ UTRs and near Stop Codons. Cell 149, 1635–1646. 10.1016/j.cell.2012.05.003.

50. King, O.D., Gitler, A.D., and Shorter, J. (2012). The tip of the iceberg: RNA-binding proteins with prion-like domains in neurodegenerative disease. Brain Research 1462, 61–80. 10.1016/j.brainres.2012.01.016.

51. Walker, L.C. (2018). Prion-like mechanisms in Alzheimer disease. Handb Clin Neurol 153, 303–319. 10.1016/B978-0-444-63945-5.00016-7.

52. Goedert, M., Clavaguera, F., and Tolnay, M. (2010). The propagation of prion-like protein inclusions in neurodegenerative diseases. Trends Neurosci 33, 317–325. 10.1016/j.tins.2010.04.003.

53. Kellett, K.A., and Hooper, N.M. (2009). Prion protein and Alzheimer disease. Prion 3, 190–194.

54. Flach, M., Leu, C., Martinisi, A., Skachokova, Z., Frank, S., Tolnay, M., Stahlberg, H., and Winkler, D.T. (2022). Trans-seeding of Alzheimer-related tau protein by a yeast prion. Alzheimers Dement 18, 2481–2492. 10.1002/alz.12581.

55. Clavaguera, F., Tolnay, M., and Goedert, M. (2017). The Prion-Like Behavior of Assembled Tau in Transgenic Mice. Cold Spring Harb Perspect Med 7, a024372. 10.1101/cshperspect.a024372.

56. Lasagna-Reeves, C.A., Castillo-Carranza, D.L., Sengupta, U., Guerrero-Munoz, M.J., Kiritoshi, T., Neugebauer, V., Jackson, G.R., and Kayed, R. (2012). Alzheimer brain-derived tau oligomers propagate pathology from endogenous tau. Sci Rep 2, 700. 10.1038/srep00700.

57. Fomicheva, A., and Ross, E.D. (2021). From Prions to Stress Granules: Defining the Compositional Features of Prion-Like Domains That Promote Different Types of Assemblies. International Journal of Molecular Sciences 22, 1251. 10.3390/ijms22031251.

58. Sprunger, M.L., and Jackrel, M.E. (2021). Prion-Like Proteins in Phase Separation and Their Link to Disease. Biomolecules 11, 1014. 10.3390/biom11071014.

59. Ries, R.J., Zaccara, S., Klein, P., Olarerin-George, A., Namkoong, S., Pickering, B.F., Patil, D.P., Kwak, H., Lee, J.H., and Jaffrey, S.R. (2019). m6A enhances the phase separation potential of mRNA. Nature 571, 424–428.

60. Batista, P.J., Molinie, B., Wang, J., Qu, K., Zhang, J., Li, L., Bouley, D.M., Lujan, E., Haddad, B., Daneshvar, K., et al. (2014). m(6)A RNA modification controls cell fate transition in mammalian embryonic stem cells. Cell Stem Cell 15, 707–719. 10.1016/j.stem.2014.09.019.

61. Chen, J., Zhang, Y.-C., Huang, C., Shen, H., Sun, B., Cheng, X., Zhang, Y.-J., Yang, Y.-G., Shu, Q., Yang, Y., et al. (2019). m6A Regulates Neurogenesis and Neuronal Development by Modulating Histone Methyltransferase Ezh2. Genomics Proteomics Bioinformatics 17, 154–168. 10.1016/j.gpb.2018.12.007.

62. Zhou, J., Wan, J., Gao, X., Zhang, X., and Qian, S.-B. (2015). Dynamic m6A mRNA methylation directs translational control of heat shock response. Nature 526, 591–594. 10.1038/nature15377.

63. Li, X., Zhu, P., Ma, S., Song, J., Bai, J., Sun, F., and Yi, C. (2015). Chemical pulldown reveals dynamic pseudouridylation of the mammalian transcriptome. Nat Chem Biol 11, 592–597. 10.1038/nchembio.1836.

64. Darnell, J.C., Jensen, K.B., Jin, P., Brown, V., Warren, S.T., and Darnell, R.B. (2001). Fragile X Mental Retardation Protein Targets G Quartet mRNAs Important for Neuronal Function. Cell 107, 489–499. 10.1016/S0092-8674(01)00566-9.

65. Darnell, J.C., Van Driesche, S.J., Zhang, C., Hung, K.Y.S., Mele, A., Fraser, C.E., Stone, E.F., Chen, C., Fak, J.J., Chi, S.W., et al. (2011). FMRP stalls ribosomal translocation on mRNAs linked to synaptic function and autism. Cell 146, 247–261. 10.1016/j.cell.2011.06.013.

66. Casingal, C.R., Kikkawa, T., Inada, H., Sasaki, Y., and Osumi, N. (2020). Identification of FMRP target mRNAs in the developmental brain: FMRP might coordinate Ras/MAPK, Wnt/β-catenin, and mTOR signaling during corticogenesis. Mol Brain 13, 167. 10.1186/s13041-020-00706-1.

67. Chang, M., Lv, H., Zhang, W., Ma, C., He, X., Zhao, S., Zhang, Z.-W., Zeng, Y.-X., Song, S., and Niu, Y. (2017). Region-specific RNA m6A methylation represents a new layer of control in the gene regulatory network in the mouse brain. Open biology 7, 170166.

68. Engel, M., Eggert, C., Kaplick, P.M., Eder, M., Röh, S., Tietze, L., Namendorf, C., Arloth, J., Weber, P., Rex-Haffner, M., et al. (2018). The Role of m6A/m-RNA Methylation in Stress Response Regulation. Neuron 99, 389–403.e9. 10.1016/j.neuron.2018.07.009.

69. Liu, J., Li, K., Cai, J., Zhang, M., Zhang, X., Xiong, X., Meng, H., Xu, X., Huang, Z., Peng, J., et al. (2020). Landscape and Regulation of m6A and m6Am Methylome across Human and Mouse Tissues. Molecular Cell 77, 426–440.e6. 10.1016/j.molcel.2019.09.032.

70. Shafik, A.M., Zhang, F., Guo, Z., Dai, Q., Pajdzik, K., Li, Y., Kang, Y., Yao, B., Wu, H., He, C., et al. (2021). N6-methyladenosine dynamics in neurodevelopment and aging, and its potential role in Alzheimer’s disease. Genome Biol 22, 17. 10.1186/s13059-020-02249-z.

71. Cajigas, I.J., Tushev, G., Will, T.J., tom Dieck, S., Fuerst, N., and Schuman, E.M. (2012). The local transcriptome in the synaptic neuropil revealed by deep sequencing and high-resolution imaging. Neuron 74, 453–466. 10.1016/j.neuron.2012.02.036.

72. The Gene Ontology Consortium (2019). The Gene Ontology Resource: 20 years and still GOing strong. Nucleic Acids Research 47, D330–D338. 10.1093/nar/gky1055.

73. Ashburner, M., Ball, C.A., Blake, J.A., Botstein, D., Butler, H., Cherry, J.M., Davis, A.P., Dolinski, K., Dwight, S.S., Eppig, J.T., et al. (2000). Gene Ontology: tool for the unification of biology. Nat Genet 25, 25–29. 10.1038/75556.

74. Hernandez, C.M., McQuail, J.A., Schwabe, M.R., Burke, S.N., Setlow, B., and Bizon, J.L. (2018). Age-Related Declines in Prefrontal Cortical Expression of Metabotropic Glutamate Receptors that Support Working Memory. eNeuro 5, ENEURO.0164-18.2018. 10.1523/ENEURO.0164-18.2018.

75. Budgett, R.F., Bakker, G., Sergeev, E., Bennett, K.A., and Bradley, S.J. (2022). Targeting the Type 5 Metabotropic Glutamate Receptor: A Potential Therapeutic Strategy for Neurodegenerative Diseases? Frontiers in Pharmacology 13.

76. Fisher, N.M., Seto, M., Lindsley, C.W., and Niswender, C.M. (2018). Metabotropic Glutamate Receptor 7: A New Therapeutic Target in Neurodevelopmental Disorders. Front Mol Neurosci 11, 387. 10.3389/fnmol.2018.00387.

77. Eden, S., Rohatgi, R., Podtelejnikov, A.V., Mann, M., and Kirschner, M.W. (2002). Mechanism of regulation of WAVE1-induced actin nucleation by Rac1 and Nck. Nature 418, 790–793. 10.1038/nature00859.

78. Rubeis, S.D., Pasciuto, E., Li, K.W., Fernández, E., Marino, D.D., Buzzi, A., Ostroff, L.E., Klann, E., Zwartkruis, F.J.T., Komiyama, N.H., et al. (2013). CYFIP1 Coordinates mRNA Translation and Cytoskeleton Remodeling to Ensure Proper Dendritic Spine Formation. Neuron 79, 1169–1182. 10.1016/j.neuron.2013.06.039.

79. Cory, G.O.C., and Ridley, A.J. (2002). Cell motility - Braking WAVEs. NATURE 418, 732–733. 10.1038/418732a.

80. Domínguez-Iturza, N., Lo, A.C., Shah, D., Armendáriz, M., Vannelli, A., Mercaldo, V., Trusel, M., Li, K.W., Gastaldo, D., Santos, A.R., et al. (2019). The autism- and schizophrenia-associated protein CYFIP1 regulates bilateral brain connectivity and behaviour. Nat Commun 10, 3454. 10.1038/s41467-019-11203-y.

81. Zheng, G., Dahl, J.A., Niu, Y., Fedorcsak, P., Huang, C.-M., Li, C.J., Vågbø, C.B., Shi, Y., Wang, W.-L., Song, S.-H., et al. (2013). ALKBH5 is a mammalian RNA demethylase that impacts RNA metabolism and mouse fertility. Mol Cell 49, 18–29. 10.1016/j.molcel.2012.10.015.

82. Schapira, M. (2016). Structural chemistry of human RNA methyltransferases. ACS chemical biology 11, 575–582.

83. Agarwala, S.D., Blitzblau, H.G., Hochwagen, A., and Fink, G.R. (2012). RNA methylation by the MIS complex regulates a cell fate decision in yeast. PLoS Genet 8, e1002732.

84. Sharma, V.K., Mehta, V., and Singh, T.G. (2020). Alzheimer’s Disorder: Epigenetic Connection and Associated Risk Factors. Curr Neuropharmacol 18, 740–753. 10.2174/1570159X18666200128125641.

85. Chen, Z., Borek, D., Padrick, S.B., Gomez, T.S., Metlagel, Z., Ismail, A., Umetani, J., Billadeau, D.D., Otwinowski, Z., and Rosen, M.K. (2010). Structure and Control of the Actin Regulatory WAVE Complex. Nature 468, 533–538. 10.1038/nature09623.

86. Bonnycastle, K., Davenport, E.C., and Cousin, M.A. (2021). Presynaptic dysfunction in neurodevelopmental disorders: Insights from the synaptic vesicle life cycle. Journal of Neurochemistry 157, 179–207. 10.1111/jnc.15035.

87. Hsiao, K., Harony-Nicolas, H., Buxbaum, J.D., Bozdagi-Gunal, O., and Benson, D.L. (2016). Cyfip1 Regulates Presynaptic Activity during Development. J Neurosci 36, 1564–1576. 10.1523/JNEUROSCI.0511-15.2016.

88. Allaire, P.D., Marat, A.L., Dall’Armi, C., Di Paolo, G., McPherson, P.S., and Ritter, B. (2010). The Connecdenn DENN Domain: A GEF for Rab35 Mediating Cargo-Specific Exit from Early Endosomes. Molecular Cell 37, 370–382. 10.1016/j.molcel.2009.12.037.

89. Klinkert, K., and Echard, A. (2016). Rab35 GTPase: A Central Regulator of Phosphoinositides and F-actin in Endocytic Recycling and Beyond. Traffic 17, 1063–1077. 10.1111/tra.12422.

90. Sheehan, P., Zhu, M., Beskow, A., Vollmer, C., and Waites, C.L. (2016). Activity-Dependent Degradation of Synaptic Vesicle Proteins Requires Rab35 and the ESCRT Pathway. J. Neurosci. 36, 8668–8686. 10.1523/JNEUROSCI.0725-16.2016.

91. Uytterhoeven, V., Kuenen, S., Kasprowicz, J., Miskiewicz, K., and Verstreken, P. (2011). Loss of Skywalker Reveals Synaptic Endosomes as Sorting Stations for Synaptic Vesicle Proteins. Cell 145, 117–132. 10.1016/j.cell.2011.02.039.

92. Campeau, P.M., Kasperaviciute, D., Lu, J.T., Burrage, L.C., Kim, C., Hori, M., Powell, B.R., Stewart, F., Félix, T.M., Ende, J. van den, et al. (2014). The genetic basis of DOORS syndrome: an exome-sequencing study. The Lancet Neurology 13, 44–58. 10.1016/S1474-4422(13)70265-5.

93. Lüthy, K., Mei, D., Fischer, B., De Fusco, M., Swerts, J., Paesmans, J., Parrini, E., Lubarr, N., Meijer, I.A., Mackenzie, K.M., et al. (2019). TBC1D24-TLDc-related epilepsy exercise-induced dystonia: rescue by antioxidants in a disease model. Brain 142, 2319–2335. 10.1093/brain/awz175.

94. Duchaîne, T.F., Hemraj, I., Furic, L., Deitinghoff, A., Kiebler, M.A., and DesGroseillers, L. (2002). Staufen2 isoforms localize to the somatodendritic domain of neurons and interact with different organelles. J Cell Sci 115, 3285–3295. 10.1242/jcs.115.16.3285.

95. Vessey, J.P., Macchi, P., Stein, J.M., Mikl, M., Hawker, K.N., Vogelsang, P., Wieczorek, K., Vendra, G., Riefler, J., Tübing, F., et al. (2008). A loss of function allele for murine Staufen1 leads to impairment of dendritic Staufen1-RNP delivery and dendritic spine morphogenesis. Proc Natl Acad Sci U S A 105, 16374–16379. 10.1073/pnas.0804583105.

96. Wu, K.Y., Hengst, U., Cox, L.J., Macosko, E.Z., Jeromin, A., Urquhart, E.R., and Jaffrey, S.R. (2005). Local translation of RhoA regulates growth cone collapse. Nature 436, 1020–1024. 10.1038/nature03885.

97. Heraud-Farlow, J.E., Sharangdhar, T., Li, X., Pfeifer, P., Tauber, S., Orozco, D., Hörmann, A., Thomas, S., Bakosova, A., Farlow, A.R., et al. (2013). Staufen2 Regulates Neuronal Target RNAs. Cell Reports 5, 1511–1518. 10.1016/j.celrep.2013.11.039.

98. Goetze, B., Tuebing, F., Xie, Y., Dorostkar, M.M., Thomas, S., Pehl, U., Boehm, S., Macchi, P., and Kiebler, M.A. (2006). The brain-specific double-stranded RNA-binding protein Staufen2 is required for dendritic spine morphogenesis. J Cell Biol 172, 221–231. 10.1083/jcb.200509035.

99. Borg, J.-P., Yang, Y., De Taddéo-Borg, M., Margolis, B., and Turner, R.S. (1998). The X11α Protein Slows Cellular Amyloid Precursor Protein Processing and Reduces Aβ40 and Aβ42 Secretion*. Journal of Biological Chemistry 273, 14761–14766. 10.1074/jbc.273.24.14761.

100. Ho, C.S., Marinescu, V., Steinhilb, M.L., Gaut, J.R., Turner, R.S., and Stuenkel, E.L. (2002). Synergistic Effects of Munc18a and X11 Proteins on Amyloid Precursor Protein Metabolism *. Journal of Biological Chemistry 277, 27021–27028. 10.1074/jbc.M201823200.

101. Mueller, H.T., Borg, J.-P., Margolis, B., and Turner, R.S. (2000). Modulation of Amyloid Precursor Protein Metabolism by X11α/Mint-1: A DELETION ANALYSIS OF PROTEIN-PROTEIN INTERACTION DOMAINS *. Journal of Biological Chemistry 275, 39302–39306. 10.1074/jbc.M008453200.

102. O’Brien, R.J., and Wong, P.C. (2011). Amyloid Precursor Protein Processing and Alzheimer’s Disease. Annu Rev Neurosci 34, 185–204. 10.1146/annurev-neuro-061010-113613.

103. Sastre, M., Turner, R.S., and Levy, E. (1998). X11 Interaction with β-Amyloid Precursor Protein Modulates Its Cellular Stabilization and Reduces Amyloid β-Protein Secretion *. Journal of Biological Chemistry 273, 22351–22357. 10.1074/jbc.273.35.22351.

104. Shrivastava-Ranjan, P., Faundez, V., Fang, G., Rees, H., Lah, J.J., Levey, A.I., and Kahn, R.A. (2008). Mint3/X11γ Is an ADP-Ribosylation Factor-dependent Adaptor that Regulates the Traffic of the Alzheimer’s Precursor Protein from the Trans-Golgi Network. MBoC 19, 51–64. 10.1091/mbc.e07-05-0465.

105. Tomita, S., Ozaki, T., Taru, H., Oguchi, S., Takeda, S., Yagi, Y., Sakiyama, S., Kirino, Y., and Suzuki, T. (1999). Interaction of a Neuron-specific Protein Containing PDZ Domains with Alzheimer’s Amyloid Precursor Protein *. Journal of Biological Chemistry 274, 2243–2254. 10.1074/jbc.274.4.2243.

106. Zhao, F., Xu, Y., Gao, S., Qin, L., Austria, Q., Siedlak, S.L., Pajdzik, K., Dai, Q., He, C., Wang, W., et al. (2021). METTL3-dependent RNA m6A dysregulation contributes to neurodegeneration in Alzheimer’s disease through aberrant cell cycle events. Molecular Neurodegeneration 16, 70. 10.1186/s13024-021-00484-x.

107. Xia, L., Zhang, F., Li, Y., Mo, Y., Zhang, L., Li, Q., Luo, M., Hou, X., Du, Z., Deng, J., et al. (2023). A new perspective on Alzheimer’s disease: m6A modification. Frontiers in Genetics 14.

108. Liu, Z., Xia, Q., Zhao, X., Zheng, F., Xiao, J., Ge, F., Wang, D., and Gao, X. (2023). The Landscape of m6A Regulators in Multiple Brain Regions of Alzheimer’s Disease. Mol Neurobiol 60, 5184–5198. 10.1007/s12035-023-03409-5.

109. Han, M., Liu, Z., Xu, Y., Liu, X., Wang, D., Li, F., Wang, Y., and Bi, J. (2020). Abnormality of m6A mRNA Methylation Is Involved in Alzheimer’s Disease. Front Neurosci 14, 98. 10.3389/fnins.2020.00098.

110. Baumann, K. (2021). Tau oligomers are linked to m6A-RNA. Nat Rev Mol Cell Biol 22, 650–650. 10.1038/s41580-021-00419-w.

111. Zhou, Y., Ćorović, M., Hoch-Kraft, P., Meiser, N., Mesitov, M., Körtel, N., Back, H., Vries, I.S.N., Katti, K., Obrdlík, A., et al. (2024). m6A sites in the coding region trigger translation-dependent mRNA decay. Molecular Cell 84, 4576–4593.e12. 10.1016/j.molcel.2024.10.033.

112. Khong, A., Matheny, T., Huynh, T.N., Babl, V., and Parker, R. (2022). Limited effects of m6A modification on mRNA partitioning into stress granules. Nat Commun 13, 3735. 10.1038/s41467-022-31358-5.

113. Murakami, S., Olarerin-George, A.O., Liu, J.F., Zaccara, S., Hawley, B., and Jaffrey, S.R. (2025). m6A alters ribosome dynamics to initiate mRNA degradation. Cell 188, 3728–3743.e20. 10.1016/j.cell.2025.04.020.

114. Wang, J., Wang, L., Diao, J., Shi, Y.G., Shi, Y., Ma, H., and Shen, H. (2020). Binding to m6A RNA promotes YTHDF2-mediated phase separation. Protein & Cell 11, 304–307. 10.1007/s13238-019-00660-2.

115. Fu, Y., and Zhuang, X. (2020). m6A-binding YTHDF proteins promote stress granule formation. Nat Chem Biol 16, 955–963. 10.1038/s41589-020-0524-y.

116. Loedige, I., Baranovskii, A., Mendonsa, S., Dantsuji, S., Popitsch, N., Breimann, L., Zerna, N., Cherepanov, V., Milek, M., Ameres, S., et al. (2023). mRNA stability and m6A are major determinants of subcellular mRNA localization in neurons. Molecular Cell 83, 2709–2725.e10. 10.1016/j.molcel.2023.06.021.

117. Zhang, M., Zhai, Y., Zhang, S., Dai, X., and Li, Z. (2020). Roles of N6-Methyladenosine (m6A) in Stem Cell Fate Decisions and Early Embryonic Development in Mammals. Front. Cell Dev. Biol. 8. 10.3389/fcell.2020.00782.

118. Wang, S., Lv, W., Li, T., Zhang, S., Wang, H., Li, X., Wang, L., Ma, D., Zang, Y., Shen, J., et al. (2022). Dynamic regulation and functions of mRNA m6A modification. Cancer Cell International 22, 48. 10.1186/s12935-022-02452-x.

119. Zhao, F., Xu, Y., Gao, S., Qin, L., Austria, Q., Siedlak, S.L., Pajdzik, K., Dai, Q., He, C., Wang, W., et al. (2021). METTL3-dependent RNA m6A dysregulation contributes to neurodegeneration in Alzheimer’s disease through aberrant cell cycle events. Molecular Neurodegeneration 16, 70. 10.1186/s13024-021-00484-x.

120. Qiao, Y., Mei, Y., Xia, M., Luo, D., and Gao, L. (2024). The role of m6A modification in the risk prediction and Notch1 pathway of Alzheimer’s disease. iScience 27. 10.1016/j.isci.2024.110235.

121. Huang, H., Camats-Perna, J., Medeiros, R., Anggono, V., and Widagdo, J. (2020). Altered Expression of the m6A Methyltransferase METTL3 in Alzheimer’s Disease. eNeuro 7, ENEURO.0125-20.2020. 10.1523/ENEURO.0125-20.2020.

122. Zou, Z., and He, C. (2024). The YTHDF proteins display distinct cellular functions on m6A-modified RNA. Trends in Biochemical Sciences 49, 611–621. 10.1016/j.tibs.2024.04.001.

123. Zaccara, S., and Jaffrey, S.R. (2020). A Unified Model for the Function of YTHDF Proteins in Regulating m6A-Modified mRNA. Cell 181, 1582–1595.e18. 10.1016/j.cell.2020.05.012.

124. Du, H., Zhao, Y., He, J., Zhang, Y., Xi, H., Liu, M., Ma, J., and Wu, L. (2016). YTHDF2 destabilizes m(6)A-containing RNA through direct recruitment of the CCR4-NOT deadenylase complex. Nat Commun 7, 12626. 10.1038/ncomms12626.

125. Shi, H., Wang, X., Lu, Z., Zhao, B.S., Ma, H., Hsu, P.J., Liu, C., and He, C. (2017). YTHDF3 facilitates translation and decay of N 6-methyladenosine-modified RNA. Cell research 27, 315–328.

126. Chang, G., Shi, L., Ye, Y., Shi, H., Zeng, L., Tiwary, S., Huse, J.T., Huo, L., Ma, L., Ma, Y., et al. (2020). YTHDF3 Induces the Translation of m6A-Enriched Gene Transcripts to Promote Breast Cancer Brain Metastasis. Cancer Cell 38, 857–871.e7. 10.1016/j.ccell.2020.10.004.

127. Lin, S., Choe, J., Du, P., Triboulet, R., and Gregory, R.I. (2016). The m6A Methyltransferase METTL3 Promotes Translation in Human Cancer Cells. Molecular Cell 62, 335–345. 10.1016/j.molcel.2016.03.021.

128. Zaccara, S., and Jaffrey, S.R. (2020). A unified model for the function of YTHDF proteins in regulating m6A-modified mRNA. Cell 181, 1582–1595.

129. Fu, Y., and Zhuang, X. (2020). m6A-binding YTHDF proteins promote stress granule formation. Nat Chem Biol 16, 955–963. 10.1038/s41589-020-0524-y.

130. Riggs, C.L., Kedersha, N., Ivanov, P., and Anderson, P. (2020). Mammalian stress granules and P bodies at a glance. J Cell Sci 133, jcs242487. 10.1242/jcs.242487.

131. Valášek, L.S., Zeman, J., Wagner, S., Beznosková, P., Pavlíková, Z., Mohammad, M.P., Hronová, V., Herrmannová, A., Hashem, Y., and Gunišová, S. (2017). Embraced by eIF3: structural and functional insights into the roles of eIF3 across the translation cycle. Nucleic Acids Res 45, 10948–10968. 10.1093/nar/gkx805.

132. Erzberger, J.P., Stengel, F., Pellarin, R., Zhang, S., Schaefer, T., Aylett, C.H.S., Cimermančič, P., Boehringer, D., Sali, A., Aebersold, R., et al. (2014). Molecular Architecture of the 40S⋅eIF1⋅eIF3 Translation Initiation Complex. Cell 158, 1123–1135. 10.1016/j.cell.2014.07.044.

133. Hinnebusch, A.G. (2014). The scanning mechanism of eukaryotic translation initiation. Annu Rev Biochem 83, 779–812. 10.1146/annurev-biochem-060713-035802.

134. Hernández-Ortega, K., Garcia-Esparcia, P., Gil, L., Lucas, J.J., and Ferrer, I. (2016). Altered Machinery of Protein Synthesis in Alzheimer’s: From the Nucleolus to the Ribosome. Brain Pathol 26, 593–605. 10.1111/bpa.12335.

135. Jain, S., Koziej, L., Poulis, P., Kaczmarczyk, I., Gaik, M., Rawski, M., Ranjan, N., Glatt, S., and Rodnina, M.V. (2023). Modulation of translational decoding by m6A modification of mRNA. Nat Commun 14, 4784. 10.1038/s41467-023-40422-7.

136. Choe, J., Lin, S., Zhang, W., Liu, Q., Wang, L., Ramirez-Moya, J., Du, P., Kim, W., Tang, S., Sliz, P., et al. (2018). mRNA circularization by METTL3-eIF3h enhances translation and promotes oncogenesis. Nature 561, 556–560. 10.1038/s41586-018-0538-8.

137. Alarcón, C.R., Goodarzi, H., Lee, H., Liu, X., Tavazoie, S., and Tavazoie, S.F. (2015). HNRNPA2B1 Is a Mediator of m6A-Dependent Nuclear RNA Processing Events. Cell 162, 1299–1308. 10.1016/j.cell.2015.08.011.

138. Knuckles, P., Lence, T., Haussmann, I.U., Jacob, D., Kreim, N., Carl, S.H., Masiello, I., Hares, T., Villaseñor, R., and Hess, D. (2018). Zc3h13/Flacc is required for adenosine methylation by bridging the mRNA-binding factor Rbm15/Spenito to the m6A machinery component Wtap/Fl (2) d. Genes & development 32, 415–429.

139. Widagdo, J., Wong, J.J.-L., and Anggono, V. (2022). The m6A-epitranscriptome in brain plasticity, learning and memory. Semin Cell Dev Biol 125, 110–121. 10.1016/j.semcdb.2021.05.023.

140. Wang, X., Zhao, B.S., Roundtree, I.A., Lu, Z., Han, D., Ma, H., Weng, X., Chen, K., Shi, H., and He, C. (2015). N6-methyladenosine Modulates Messenger RNA Translation Efficiency. Cell 161, 1388–1399. 10.1016/j.cell.2015.05.014.

141. Shi, H., Wei, J., and He, C. (2019). Where, when, and how: context-dependent functions of RNA methylation writers, readers, and erasers. Molecular cell 74, 640–650.

142. Hsu, P.J., Shi, H., Zhu, A.C., Lu, Z., Miller, N., Edens, B.M., Ma, Y.C., and He, C. (2019). The RNA-binding protein FMRP facilitates the nuclear export of N6-methyladenosine–containing mRNAs. J Biol Chem 294, 19889–19895. 10.1074/jbc.AC119.010078.

143. Lai, A., Valdez-Sinon, A.N., and Bassell, G.J. (2020). Regulation of RNA granules by FMRP and implications for neurological diseases. Traffic 21, 454–462. 10.1111/tra.12733.

144. Zhang, F., Kang, Y., Wang, M., Li, Y., Xu, T., Yang, W., Song, H., Wu, H., Shu, Q., and Jin, P. (2018). Fragile X mental retardation protein modulates the stability of its m6A-marked messenger RNA targets. Human molecular genetics 27. 10.1093/hmg/ddy292.

145. Edupuganti, R.R., Geiger, S., Lindeboom, R.G.H., Shi, H., Hsu, P.J., Lu, Z., Wang, S.-Y., Baltissen, M.P.A., Jansen, P.W.T.C., Rossa, M., et al. (2017). N 6 - methyladenosine (m 6 A) recruits and repels proteins to regulate mRNA homeostasis. Nature Structural & Molecular Biology 24, 870–878. 10.1038/nsmb.3462.

146. Deng, X., Su, R., Weng, H., Huang, H., Li, Z., and Chen, J. (2018). RNA N 6-methyladenosine modification in cancers: current status and perspectives. Cell research 28, 507–517.

147. Narayan, P., Ludwiczak, R.L., Goodwin, E.C., and Rottman, F.M. (1994). Context effects on N6-adenosine methylation sites in prolactin mRNA. Nucleic Acids Res 22, 419–426. 10.1093/nar/22.3.419.

148. Thelen, M.P., and Kye, M.J. (2020). The Role of RNA Binding Proteins for Local mRNA Translation: Implications in Neurological Disorders. Front. Mol. Biosci. 6. 10.3389/fmolb.2019.00161.

149. Jishi, A., Qi, X., and Miranda, H. (2020). Implications of mRNA translation dysregulation for neurological disorders. Seminars in Cell and Developmental Biology 114. 10.1016/j.semcdb.2020.09.005.

150. Heraud-Farlow, J.E., Sharangdhar, T., Li, X., Pfeifer, P., Tauber, S., Orozco, D., Hörmann, A., Thomas, S., Bakosova, A., Farlow, A.R., et al. (2013). Staufen2 Regulates Neuronal Target RNAs. Cell Reports 5, 1511–1518. 10.1016/j.celrep.2013.11.039.

151. Kusek, G., Campbell, M., Doyle, F., Tenenbaum, S.A., Kiebler, M., and Temple, S. (2012). Asymmetric Segregation of the Double-Stranded RNA Binding Protein Staufen2 during Mammalian Neural Stem Cell Divisions Promotes Lineage Progression. Cell Stem Cell 11, 505–516. 10.1016/j.stem.2012.06.006.

152. Mallardo, M., Deitinghoff, A., Müller, J., Goetze, B., Macchi, P., Peters, C., and Kiebler, M.A. (2003). Isolation and characterization of Staufen-containing ribonucleoprotein particles from rat brain. PNAS 100, 2100–2105. 10.1073/pnas.0334355100.

153. Kiebler, M.A., Hemraj, I., Verkade, P., Köhrmann, M., Fortes, P., Marión, R.M., Ortín, J., and Dotti, C.G. (1999). The mammalian staufen protein localizes to the somatodendritic domain of cultured hippocampal neurons: implications for its involvement in mRNA transport. Journal of Neuroscience 19, 288–297.

154. Zheng, X., Zeng, F., Lei, Y., Li, Y., Deng, J., Luo, G., He, Q., and Zhou, Y. (2025). YBX1: an RNA/DNA-binding protein that affects disease progression. Front. Oncol. 15. 10.3389/fonc.2025.1635209.

155. Bobkova, N.V., Lyabin, D.N., Medvinskaya, N.I., Samokhin, A.N., Nekrasov, P.V., Nesterova, I.V., Aleksandrova, I.Y., Tatarnikova, O.G., Bobylev, A.G., and Vikhlyantsev, I.M. (2015). The Y-box binding protein 1 suppresses Alzheimer’s disease progression in two animal models. PLoS One 10, e0138867.

156. Suresh, P.S., Tsutsumi, R., and Venkatesh, T. (2018). YBX1 at the crossroads of non-coding transcriptome, exosomal, and cytoplasmic granular signaling. European Journal of Cell Biology 97, 163–167. 10.1016/j.ejcb.2018.02.003.

157. Tang, J., Liu, J., Nie, J., Pei, H., and Zhou, G. (2024). YBX1 Underwent Phase Separation into Stress Granules Stimulated by Ionizing Radiation. Radiat Res 201, 215–223. 10.1667/rade-23-00113.1.

158. Raveendra, B.L., Swarnkar, S., Avchalumov, Y., Liu, X.-A., Grinman, E., Badal, K., Reich, A., Pascal, B.D., and Puthanveettil, S.V. (2018). Long noncoding RNA GM12371 acts as a transcriptional regulator of synapse function. Proc Natl Acad Sci U S A 115, E10197–E10205. 10.1073/pnas.1722587115.

159. Brown, V., Jin, P., Ceman, S., Darnell, J.C., O’Donnell, W.T., Tenenbaum, S.A., Jin, X., Feng, Y., Wilkinson, K.D., Keene, J.D., et al. (2001). Microarray identification of FMRP-associated brain mRNAs and altered mRNA translational profiles in fragile X syndrome. Cell 107, 477–487. 10.1016/s0092-8674(01)00568-2.

160. Ge, S.X., Jung, D., and Yao, R. (2020). ShinyGO: a graphical gene-set enrichment tool for animals and plants. Bioinformatics 36, 2628–2629. 10.1093/bioinformatics/btz931.

161. Ge, S.X., and Jung, D. ShinyGO: a web application for in-depth analysis of gene sets.

162. Ritchie, M.E., Phipson, B., Wu, D., Hu, Y., Law, C.W., Shi, W., and Smyth, G.K. (2015). limma powers differential expression analyses for RNA-sequencing and microarray studies. Nucleic Acids Research 43, e47. 10.1093/nar/gkv007.

163. Wickham, H. (2011). ggplot2. WIREs Computational Statistics 3, 180–185. 10.1002/wics.147.

164. Robinson, M.D., McCarthy, D.J., and Smyth, G.K. (2010). edgeR: a Bioconductor package for differential expression analysis of digital gene expression data. Bioinformatics 26, 139–140. 10.1093/bioinformatics/btp616.

165. Goedhart, J., and Luijsterburg, M.S. (2020). VolcaNoseR is a web app for creating, exploring, labeling and sharing volcano plots. Sci Rep 10, 20560. 10.1038/s41598-020-76603-3.

166. Lancaster, A.K., Nutter-Upham, A., Lindquist, S., and King, O.D. (2014). PLAAC: a web and command-line application to identify proteins with prion-like amino acid composition. Bioinformatics 30, 2501–2502. 10.1093/bioinformatics/btu310.

