## Supplementary datas for "m⁶A modification and prion-like domain proteins converge to dysregulate Neuronal RNA Granules in Alzheimer’s disease": Additional File 1; PrLDs Proteins detected by PLAAC.pdf

### NP_058085.2 heterogeneous nuclear ribonucleoprotein U [Mus musculus]

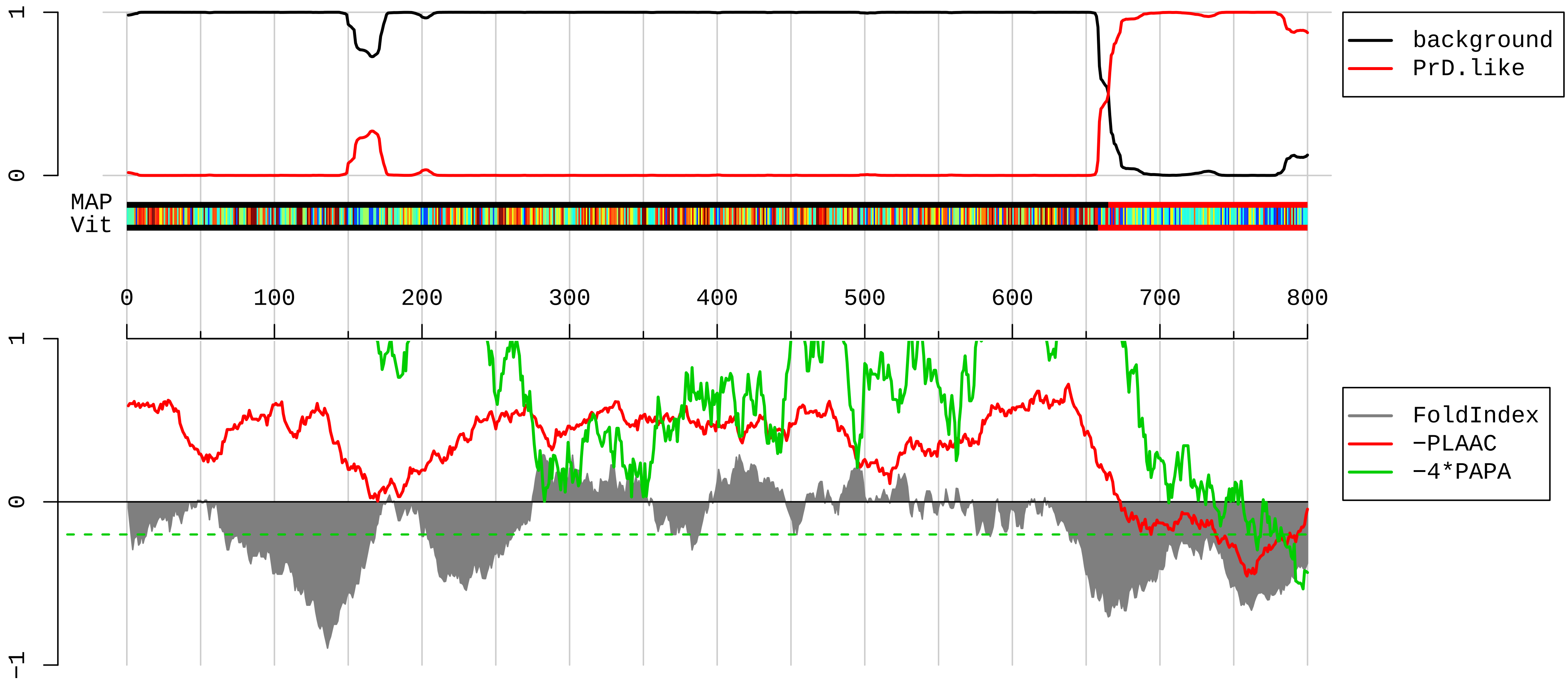


1 MSSSPVNVKKLKVSELKEELKKRRLSDKGLKADLMDRLQAALDNEAGGRPAMEPGNGSLDLGGDAAGRSGAGLEQEAAAGAEDDEEEEGIAALDGDQMEL
101 GEENGAAGAADAGAMEEEEAASEDENGDDQGFQEGEDELGDEEEGAGDENGHGEQQSQPPAAAAQQQPSQQRGAGKEAAGKSSGPTSLFAVTVAPPGARQ
201 GQQQAGGDGKTEQKGGDKKRGVKRPREDHGRGYFEYIEENKYSRAKSPQPPVEEEDEHFDDTVVCLDTYNCDLHFKISRDRLSASSLTMESFAFLWAGGR
301 ASYGVSKGKVCFEMKVTEKIPVRHLYTKDIDIHEVRIGWSLTTSGMLLGEEEFSYGYSLKGIKTCNCETEDYGEKFDENDVITCFANFETDEVELSYAKN
401 GQDLGVAFKISKEVLADRPLFPHVLCHNCAVEFNFGQKEKPYFPIPEDCTFIQNVPLEDRVRGPKGPEEKKDCEVVMMIGLPGAGKTTWVTKHAAENPGK
501 YNILGTNTIMDKMMVAGFKKQMADTGKLNTLLQRAPQCLGKFIEIAARKKRNFILDQTNVSAAAQRRKMCLFAGFQRKAVVVCPKDEDYKQRTQKKAEVE
601 GKDLPEHAVLKMKGNFTLPEVAECFDEITYVELQKEEAQKLLEQYKEESKKALPPEKKQNTGSKKSNKNKSGKNQFNRGGGHRGRGGFNMRGGNFRGGAP
701 GNRGGYNRRGNMPQRGGGGGSGGIGYPYPRGPVFPGRGGYSNRGNYNRGGMPNRGNYNQNFRGRGNNRGYKNQSQGYNQWQQGQFWGQKPWSQHYHQGYY
801

#### AAI27031.1 Caprin1 protein, partial [Mus musculus]

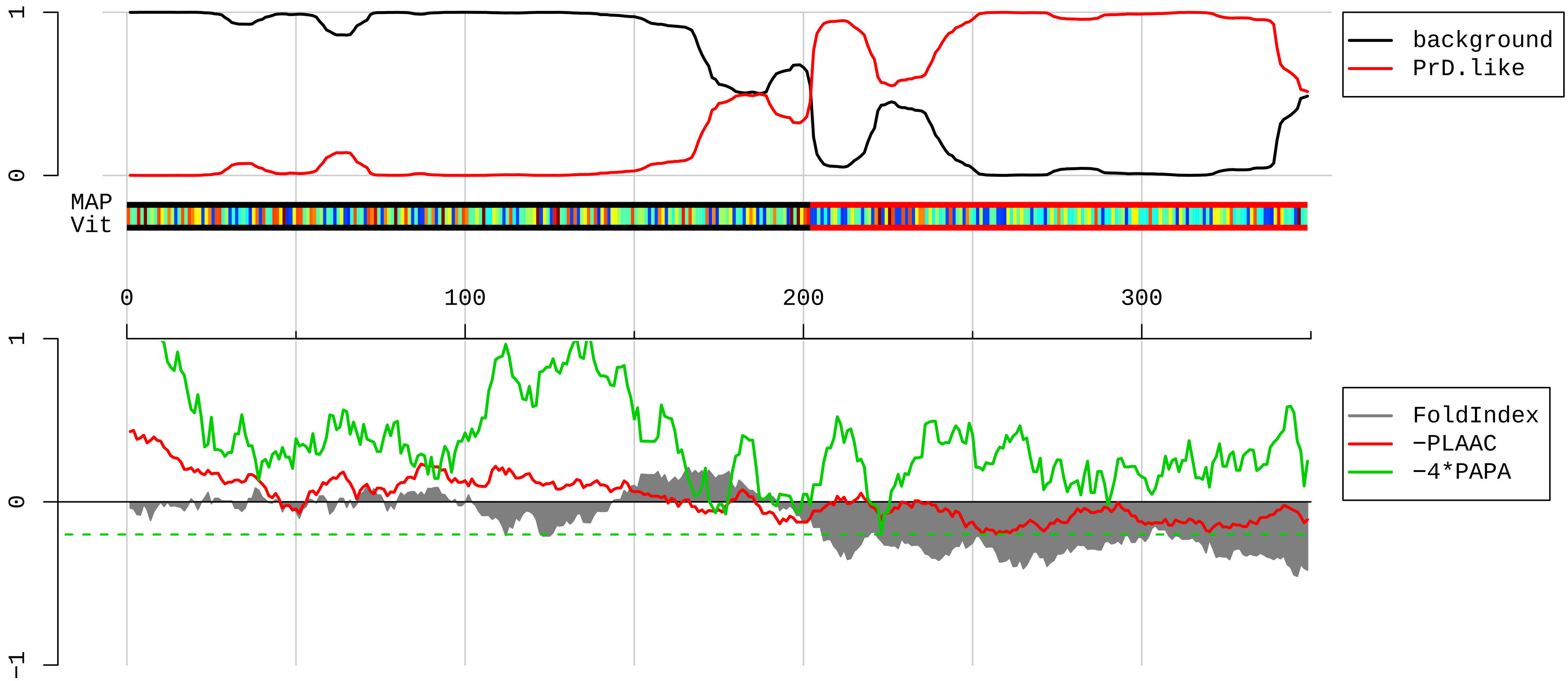


1 DPSCPEPHSLTPVAQSDPLVRRQRVQDLMAQMQGPYNFIQDSMLDFENQTLDPAIVSAQPMNPTQNMDMPQLVCPQVHSESRLAQSNQVPVQPEATQVPL
101 VSSTSEGYTASQPLYQPSHATEQRPQKEPMDQIQATISLNTDQTTASSSLPAASQPQVFQAGTSKPLHSSGINVNAAPFQSMQTVFNMNAPVPPANEPET
201 LKQQSQYQATYNQSFSSQPHQVEQTELQQDQLQTVVGTYHGSQDQPHQVPGNHQQPPQQNTGFPRSSQPYYNSRGVSRGGSRGARGLMNGYRGPANGFRG
301 GYDGYRPSFSNTPNSGYSQSQFTAPRDYSGYQRDGYQQNFKRGSGQEGS

AAH56442.1 Upf1 protein [Mus musculus]

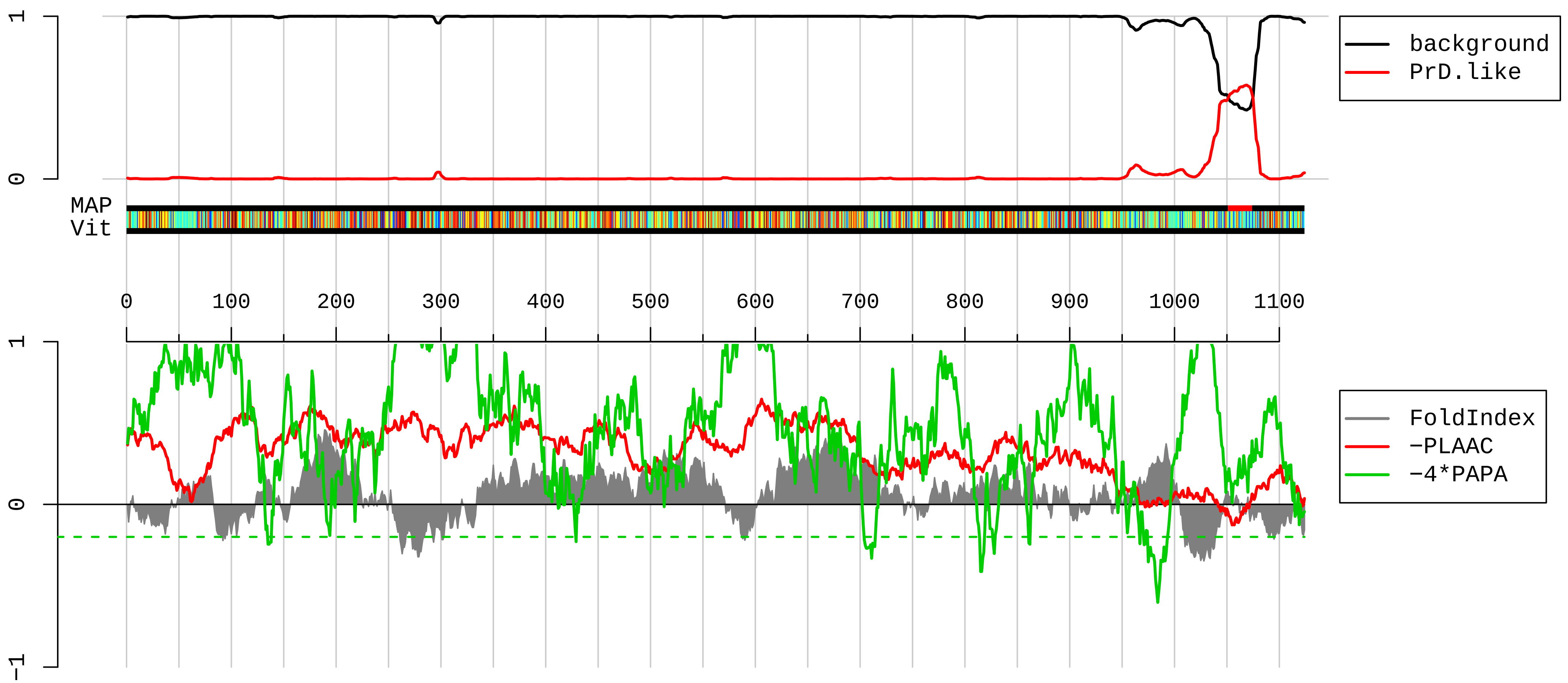


1 MSVEAYGPSSQTLTFLDTEEAELLGADTQGSEFEFTDFTLPSQTQTPPGGPGGAGGPGGAGAGGAAGQLDAQVGPEGILQNGAVDDSVAKTSQLLAELNF
101 EEDEEDTYYTKDLPVHACSYCGIHDPACVVYCNTSKKWFCNGRGNTSGSHIVNHLVRAKCKEVTLHKDGPLGETVLECYNCGCRNVFLLGFIPAKADSVV
201 VLLCRQPCASQSSLKDINWDSSQWQPLIQDRCFLSWLVKIPSEQEQLRARQITAQQINKLEELWKENPSATLEDLEKPGVDEEPQHVLLRYEDAYQYQNI
301 FGPLVKLEADYDKKLKESQTQDNITVRWDLGLNKKRIAFFTLPKTDSGNEDLVIIWLRDMRLMQGDEICLRYKGDLAPLWKGIGHVIKVPDNYGDEIAIE
401 LRSSVGAPVEVTHNFQVDFVWKSTSFDRMQSALKTFAVDETSVSGYIYHKLLGHEVEDVVIKCQLPKRFTAQGLPDLNHSQVYAVKTVLQRPLSLIQGPP
501 GTGKTVTSATIVYHLARQGNGPVLVCAPSNIAVDQLTEKIHQTGLKVVRLCAKSREAIDSPVSFLALHNQIRNMDSMPELQKLQQLKDETGELSSADEKR
601 YRALKRTAERELLMNADVICCTCVGAGDPRLAKMQFRSILIDESTQATEPECMVPVVLGAKQLILVGDHCQLGPVVMCKKAAKAGLSQSLFERLVVLGIR
701 PIRLQVQYRMHPALSAFPSNIFYEGSLQNGVTAADRVKKGFDFQWPQPDKPMFFYVTQGQEEIASSGTSYLNRTEAANVEKITTKLLKAGAKPDQIGIIT
801 PYEGQRSYLVQYMQFSGSLHTKLYQEVEIASVDAFQGREKDFIILSCVRANEHQGIGFLNDPRRLNVALTRARYGVIIVGNPKALSKQPLWNHLLSYYKE
901 QKALVEGPLNNLRESLMQFSKPRKLVNTVNPGARFMTTAMYDAREAIIPGSVYDRSSQGRPSNMYFQTHDQISMISAGPSHVAAMNIPIPFNLVMPPMPP
1001 PGYFGQANGPAAGRGTPKTKTGRGGRQKNRFGLPGPSQTTLPNSQASQDVASQPFSQGALTQGYVSMSQPSQMSQPGLSQPELSQDSYLGDEFKSQIDVA
1101 LSQDSTYQGERAYQHGGVTGLSQY

**AAH48159.1 Elavl4 protein [Mus musculus]**

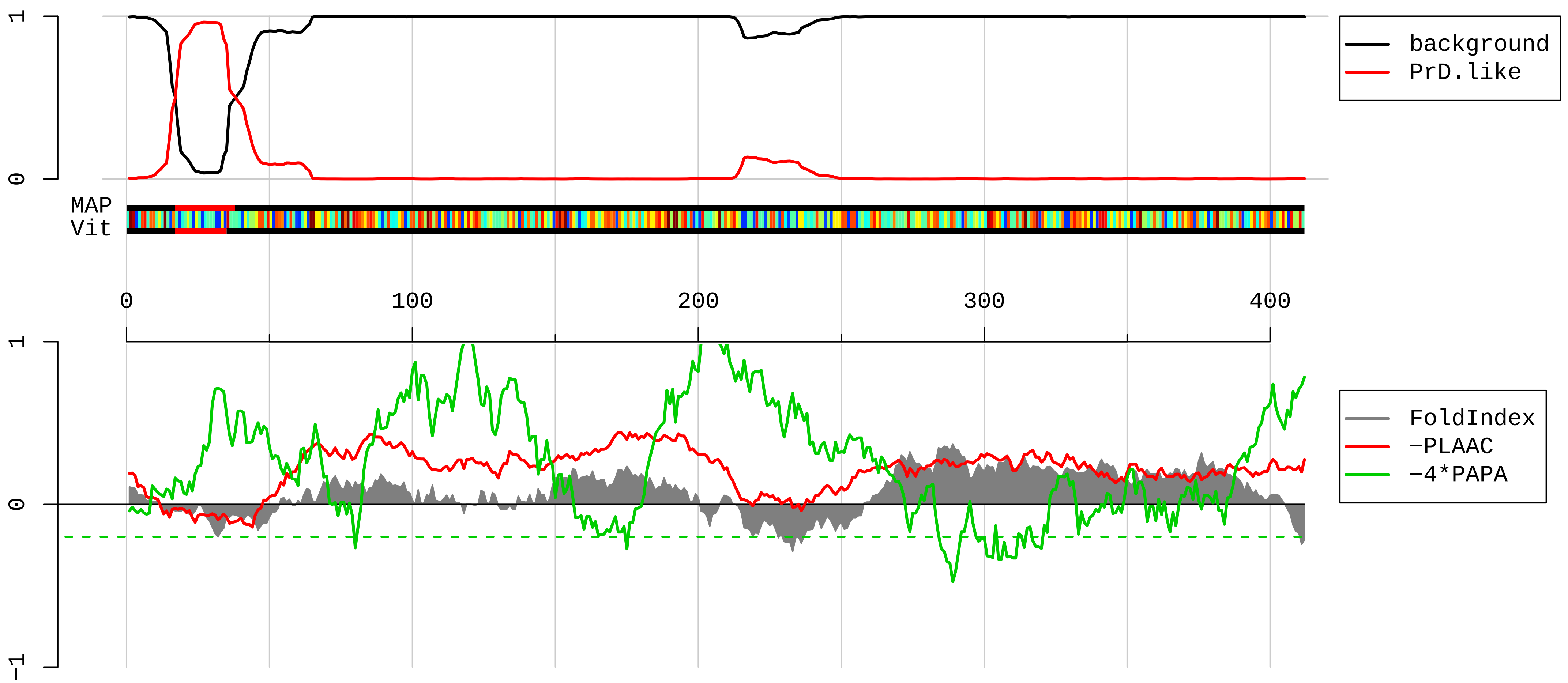


```
  1  MEWNGLKMIISTMEPQVSNGPTSNTSNGPSSNNRNCPSPMQTGAATDDSKTNLIVNYLPQNMTQEEFRSLFGSIGEIESCKLVRDKITGQSLGYGFVNYI
101  DPKDAEKAINTLNGLRLQTKTIKVSYARPSSASIRDANLYVSGLPKTMTQKELEQLFSQYGRIITSRILVDQVTGVSRGVGFIRFDKRIEAEEAIKGLNG
201  QKPSGATEPITVKFANNPSQKSSQALLSQLYQSPNRRYPGPLHHQAQRFRLDNLLNMAYGVKRLMSGPVPPSACPPRFSPITIDGMTSLVGMNIPGHTGT
301  GWCIFVYNLSPDSDESVLWQLFGPFGAVNNVKVIRDFNTNKCKGFGFVTMTNYDEAAMAIASLNGYRLGDRVLQVSMTNYDEAAMAIASLNGYRLGDRVL
401  QVSFKTNKAHKS
```

**EDL08818.1 poly A binding protein, cytoplasmic 1 [Mus musculus]**

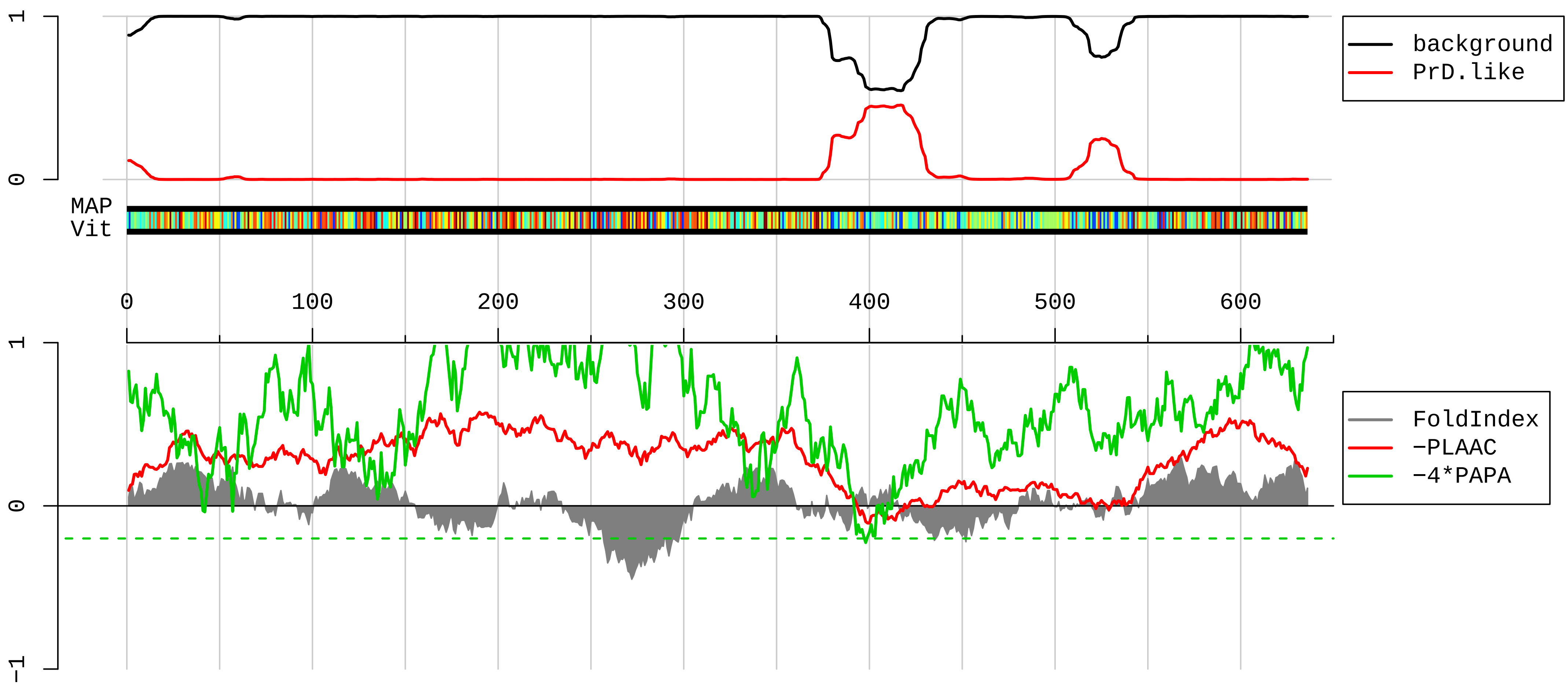


```
1    MNPSAPSYPMASLYVGDLHPDVTEAMLYEKFSPAGPILSIRVCRDMITRRSLGYAYVNFQQPADAERALDTMNFDVIKGKPVRIMWSQRDPSLRKSGVGN
101  IFIKNLDKSIDNKALYDTFSAFGNILSCKVVCDENGSKGYGFVHFETQEAAERAIEKMNGMLLNDRKVFVGRFKSRKEREAELGARAKEFTNVYIKNFGE
201  DMDDERLKELFGKFGPALSVKVMTDESGKSKGFGFVSFERHEDAQKAVDEMNGKELNGKQIYVGRAQKKVERQTELKRKFEQMKQDRITRYQGVNLYVKN
301  LDDGIDDERLRKEFSPFGTITSAKVMMEGGRSKGFGFVCFSSPEEATKAVTEMNGRIVATKPLYVALAQRKEERQAHLTNQYMQRMASVRAVPNPVINPY
401  QPAPPSGYFMAAIPQTQNRAAYYPPSQIAQLRPSPRWTAQGARPHPFQNMPGAIRPAAPRPPFSTMRPASSQVPRVMSTQRVANTSTQTMGPRPAAAAAA
501  ATPAVRTVPQYKYAAGVRNPQQHLNAQPQVTMQQPAVHVQGQEPLTASMLASAPPQEQKQMLGERLFPLIQAMHPSLAGKITGMLLEIDNSELLHMLESP
601  ESLRSKVDEAVAVLQAHQAKEAAQKAVNSATGVPTV
```

**NP_035862.2 Y-box-binding protein 1 [Mus musculus]**

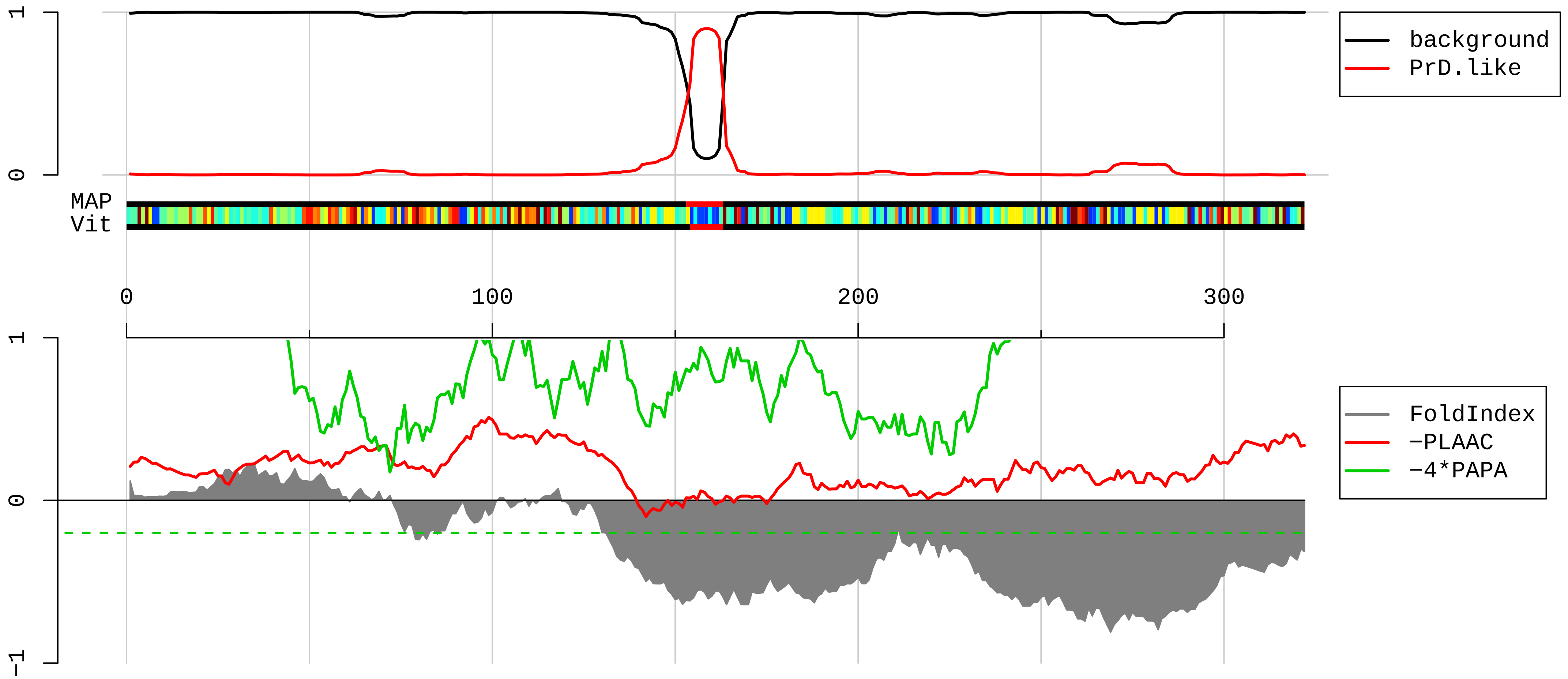


1 MSSEAETQQPPAAPAAALSAADTKPGSTGSGAGSGGPGGLTSAAPAGGDKKVIATKVLGTVKWFNVRNGYGFINRNDTKEDVFVHQTAIKKNNPRKYLRS
101 VGDGETVEFDVVEGEKGAEAANVTGPGGVPVQGSKYAADRNHYRRYPRRRGPPRNYQQNYQNSESGEKNEGSESAPEGQAQQRRPYRRRRFPPYYMRRPY
201 ARRPQYSNPPVQGEVMEGADNQGAGEQGRPVRQNMYRGYRPRFRRGPPRQRQPREDGNEEDKENQGDETQGQQPPQRRYRRNFNYRRRRPENPKPQDGKE
301 TKAADPPAENSSAPEAEQGGAE

**NP_997568.1 ELAV-like protein 2 isoform 1 [Mus musculus]**

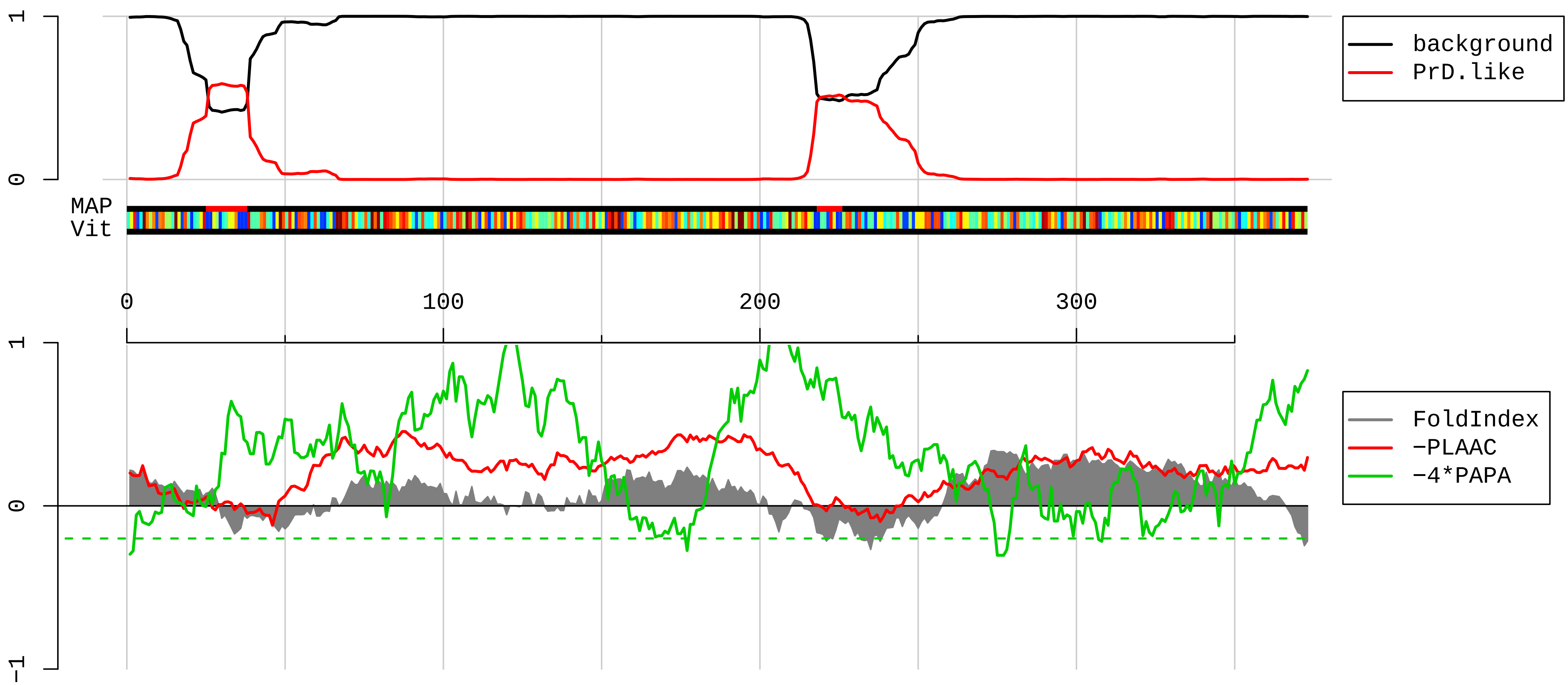


```
1   MTKQYEVAVQVIAAMETQLSNGPTCNNTANGPTTVNNNCSSPVDSGNTEDSKTNLIVNYLPQNMTQEELKSLFGSIGEIESCKLVRDKITGQSLGYGFVN
101 YIDPKDAEKAINTLNGLRLQTKTIKVSYARPSSASIRDANLYVSGLPKTMTQKELEQLFSQYGRIITSRILVDQVTGISRGVGFIRFDKRIEAEEAIKGL
201 NGQKPPGATEPITVKFANNPSQKTNQAILSQLYQSPNRRYPGPLAQQAQRFRLDNLLNMAYGVKRFSPMTIDGMTSLAGINIPGHPGTGWCIFVYNLAPD
301 ADESILWQMFGPFGAVTNVKVIRDFNTNKCKGFGFVTMTNYDEAAMAIASLNGYRLGDRVLQVSFKTNKTHKA
```

**AAH25118.1 Stau2 protein [Mus musculus]**

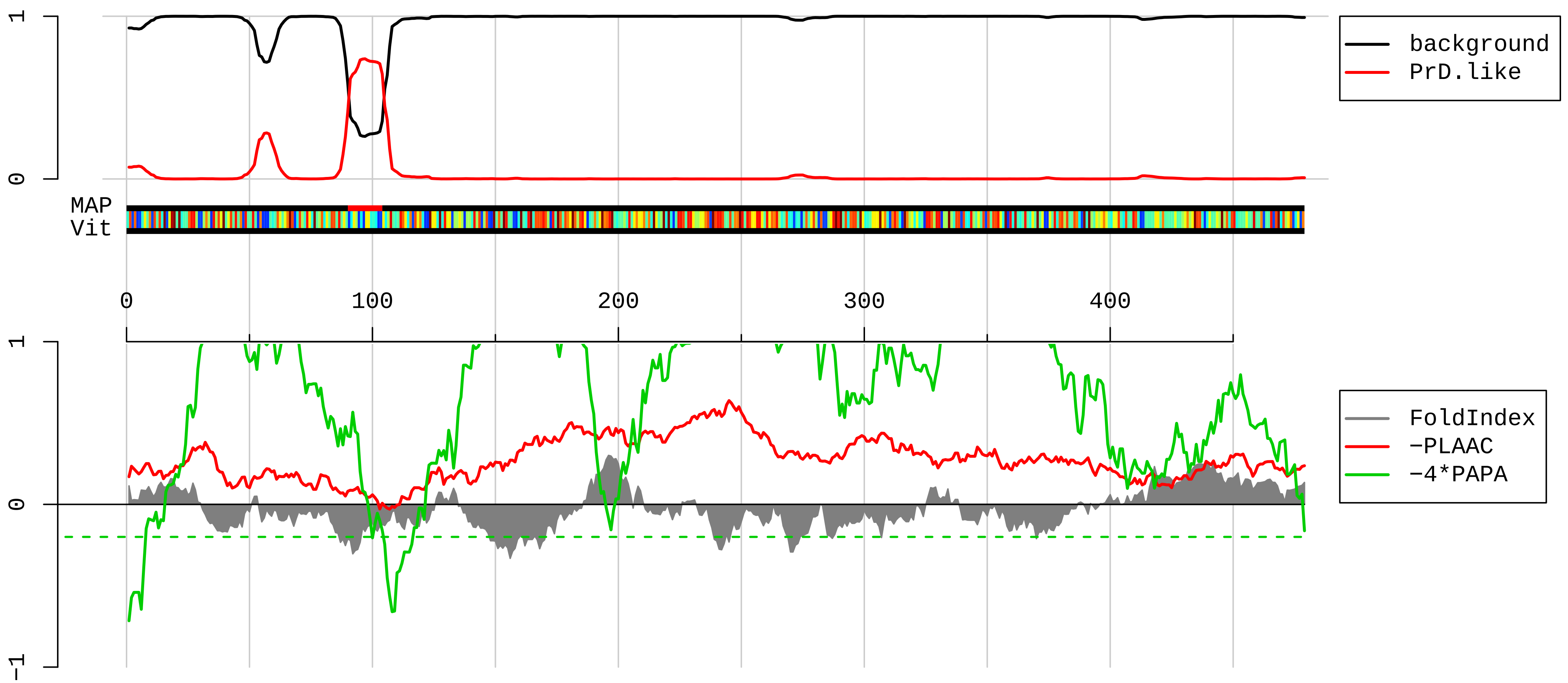


1 MLQINQMFSVQLSLGEQTWESEGSSIKKAQQAVANKALTESTLPKPVQKPPKSNVNNNPGSITPTVELNGLAMKRGEPAIYRPLDPKPFPNYRANYNFRG
101 MYNQRYHCPMPKIFYVQLTVGNNEFFGEGKTRQAARHNAAMKALQALQNEPIPEKSPQNGESGKEMDDDKDANKSEISLVFEIALKRNMPVSFEVIKESG
201 PPHMKSFVTRVSVGEFSAEGEGNSKKLSKKRAATTVLQELKKLPPLPVVEKPKLFFKKRPKTIVKAGPDYGQGMNPISRLAQIQQARKEKEPDYILLSER
301 GMPRRREFVMQVKVGNEVATGTGPNKKIAKKNAAEAMLLQLGYKASTSLQDPLDKTGENKGWSGPKPGFPEPTNNTPKGILHLSPDVYQEMEASRHRVTS
401 GTTLSYLSPKDMNQPSSSFFSVSPSSTSSATVARELLMNGTSPTAEAIGLKGSSPTSPCSSVQPSKQLEYLARIQGFQV

**AAH62971.1 Fragile X mental retardation, autosomal homolog 2 [Mus musculus]**

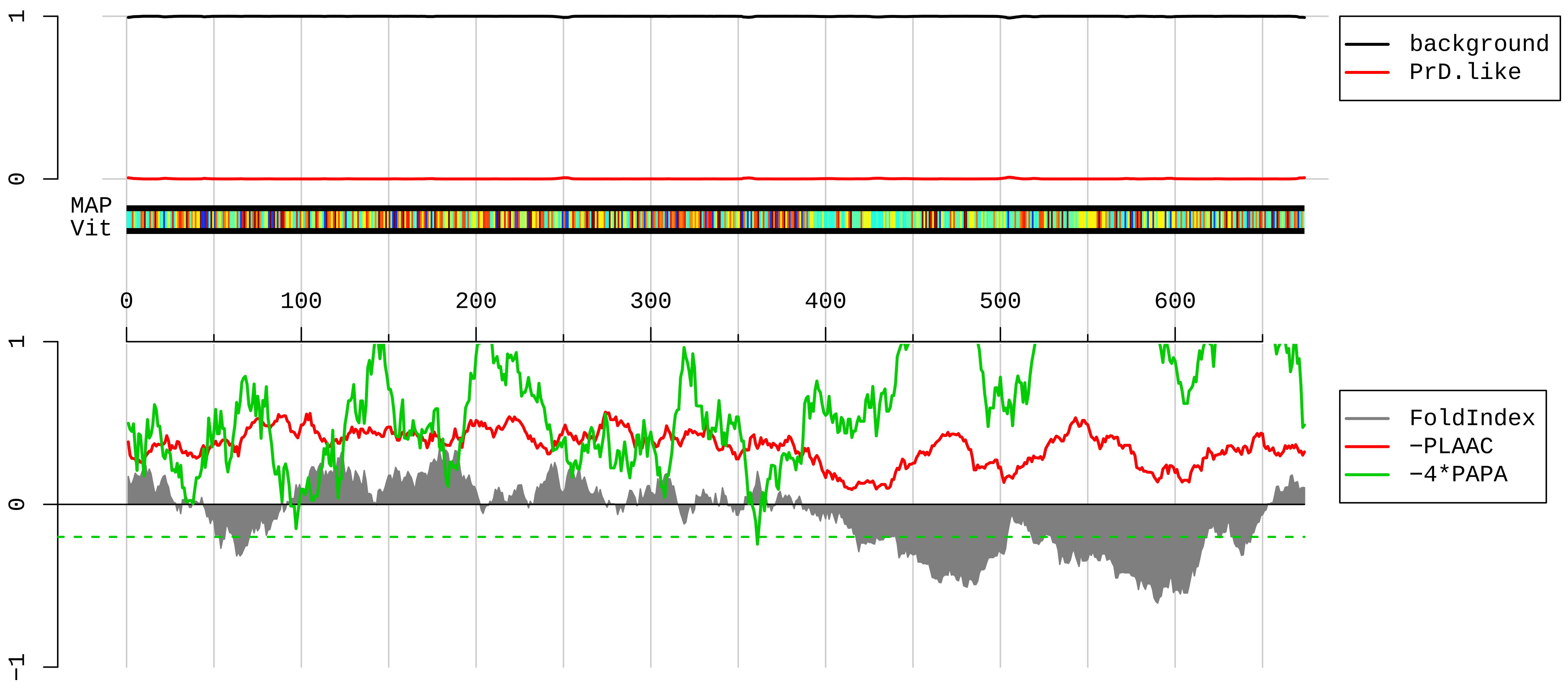


1 MGGLASGGDVEPGLPVEVRGSNGAFYKGFVKDVHEDSVTIFFENNWQSERQIPFGDVRLPPPADYNKEITEGDEVEVYSRANEQEPCGWWLARVRMMKGD
101 FYVIEYAACDATYNEIVTLERLRPVNPSPLATKGSFFKVTMAVPEDLREACSNENVHKEFKKALGANCIFLNITNSELFILSTTEAPVKRASLLGDMHFR
201 SLRTKLLLMSRNEEATKHLETSKQLAAAFQEEFTVREDLMGLAIGTHGANIQQARKVPGVTAIELGEETCTFRIYGETPEACRQARSYLEFSEDSVQVPR
301 NLVGKVIGKNGKVIQEIVDKSGVVRVRVEGDNDKKNPREEGMVPFIFVGTRENISNAQALLEYHLSYLQEVEQLRLERLQIDEQLRQIGLGFRPPGSGRG
401 GSGGGSDKAGYTTDESSSSSLHTTRTYGGSYGGRGRGRRTGGPAYGPSSDPSTASETESEKREESNRAGPGDRDPPSRGEESRRRPIGGRGRGPPPVPRP
501 TSRYNSSSISSVLKDPDSNPYSLLDTSEPEPPVDSEPGEPPPASARRRRSRRRRTDEDRTVMDGGLESDGPNMTENGLEDESRPQRRNRSRRRRNRGNRT
601 DGSISGDRQPVTVADYISRAESQSRQRPLERTKPSEDSLSGQKGDSVSKLPKGPSENGELSAPLELGSLVNGVS

**AAH55441.1 Tubulin, beta 2A [Mus musculus]**

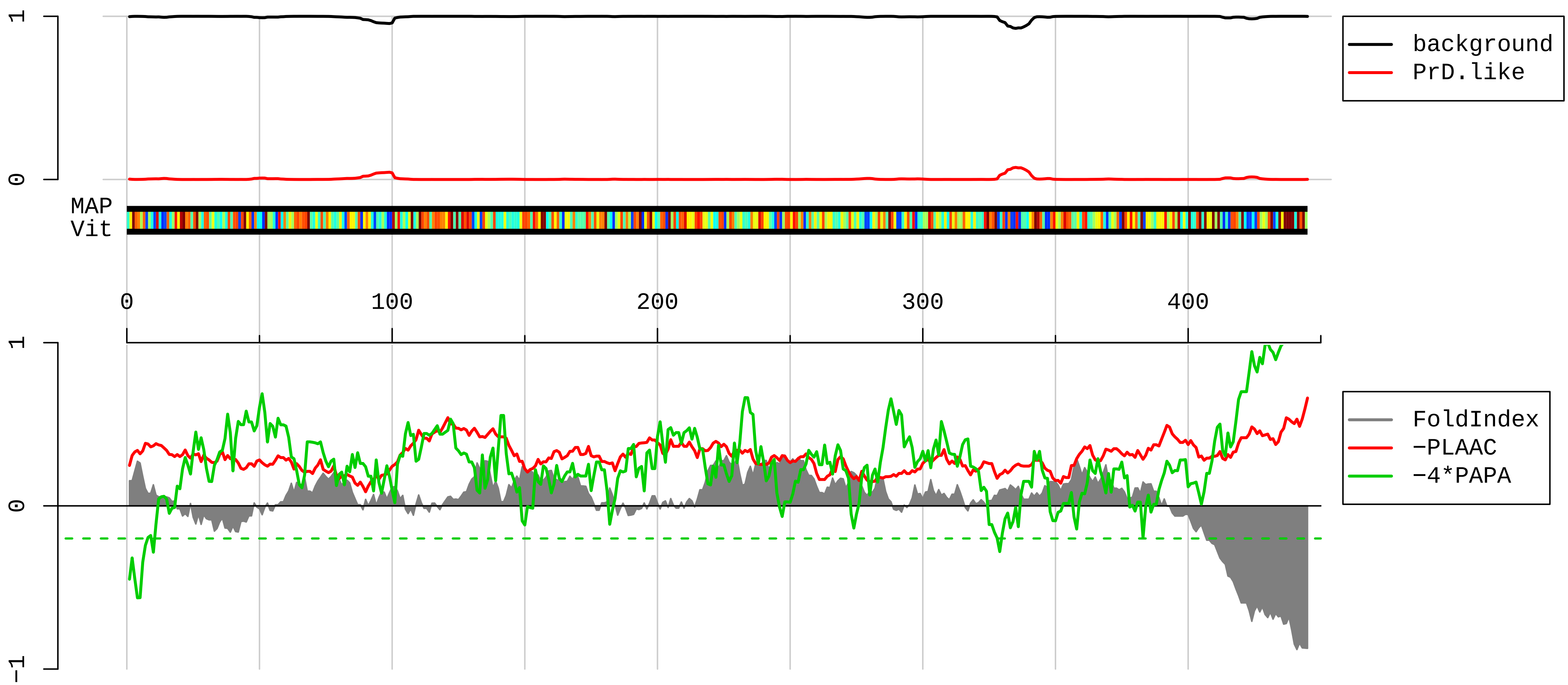


```
  1  MREIVHIQAGQCGNQIGAKFWEVISDEHGIDPTGSYHGDSDLQLERINVYYNEAAGNKYVPRAILVDLEPGTMDSVRSGPFGQIFRPDNFVFGQSGAGNN
101  WAKGHYTEGAELVDSVLDVVRKESESCDCLQGFQLTHSLGGGTGSGMGTLLISKIREEYPDRIMNTFSVMPSPKVSDTVVEPYNATLSVHQLVENTDETY
201  SIDNEALYDICFRTLKLTTPTYGDLNHLVSATMSGVTTCLRFPGQLNADLRKLAVNMVPFPRLHFFMPGFAPLTSRGSQQYRALTVPELTQQMFDSKNMM
301  AACDPRHGRYLTVAAIFRGRMSMKEVDEQMLNVQNKNSSYFVEWIPNNVKTAVCDIPPRGLKMSATFIGNSTAIQELFKRISEQFTAMFRRKAFLHWYTG
401  EGMDEMEFTEAESNMNDLVSEYQQYQDATADEQGEFEEEEGEDEA
```

**AAI32322.1 Ribosomal protein L4 [Mus musculus]**

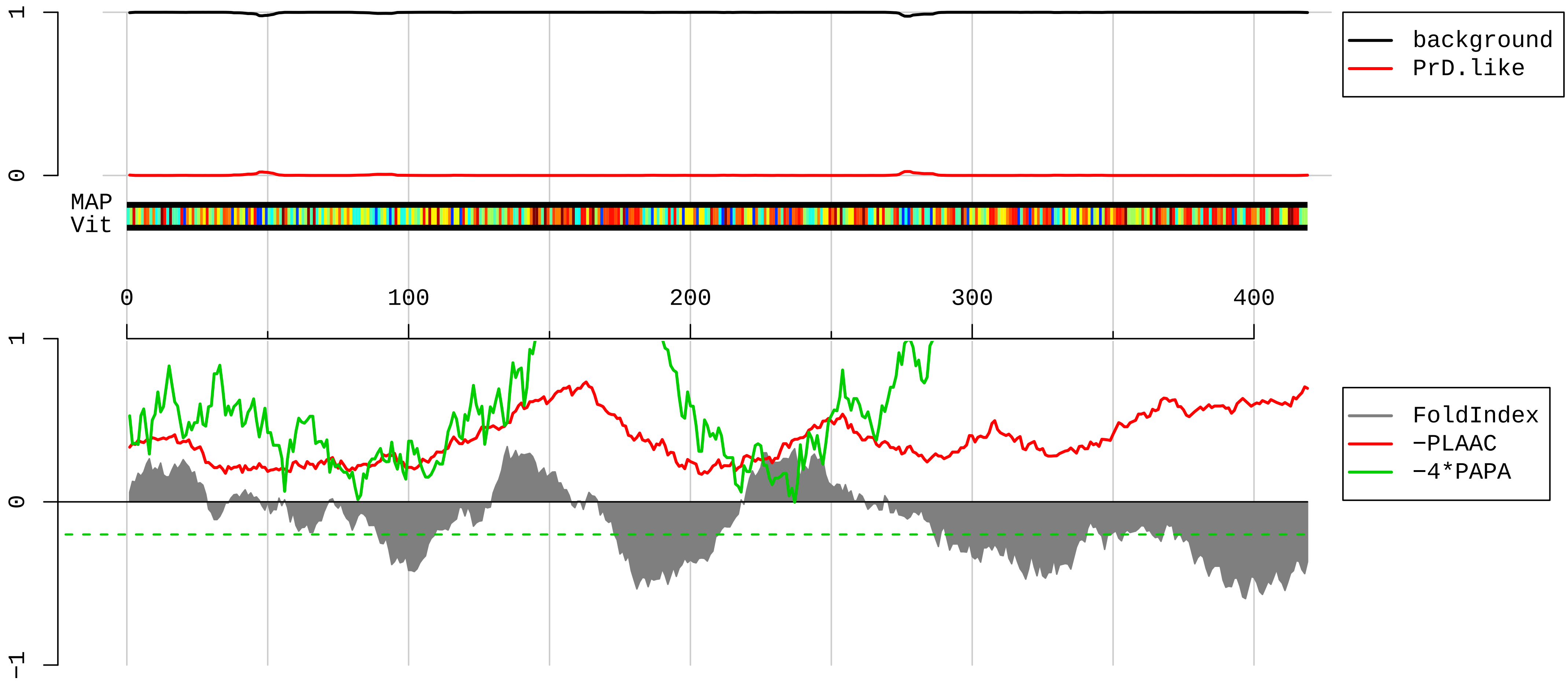


```
  1  MACARPLISVYSEKGESSGKNVTLPAVFKAPIRPDIVNFVHTNLRKNNRQPYAVSELAGHQTSAESWGTGRAVARIPRVRGGGTHRSGQGAFGNMCRGGR
101  MFAPTKTWRRWHRRVNTTQKRYAICSALAASALPALVMSKGHRIEEVPELPLVVEDKVEGYKKTKEAVQLLKKLKAWNDIKKVYASQRMRAGKGKMRNRR
201  RIQRRGPCIIYNEDNGIIKAFRNIPGITLLNVSKLNILKLAPGGHVGRFCIWTESAFRKLDELYGTWRKAASLKSNYNLPMHKMMNTDLSRILKSPEIQR
301  ALRAPRKKIHRRVLKKNPLKNLRIMLKLNPYAKTMRRNTILRQARNHKLRVKKLEAAATALATKSEKVVPEKGTADKKPAVGKKGKKVDAKKQKPAGKKV
401  VAKKPAEKKPTTEEKKPAA
```

**NP_035785.1 tubulin beta-5 chain [Mus musculus]**

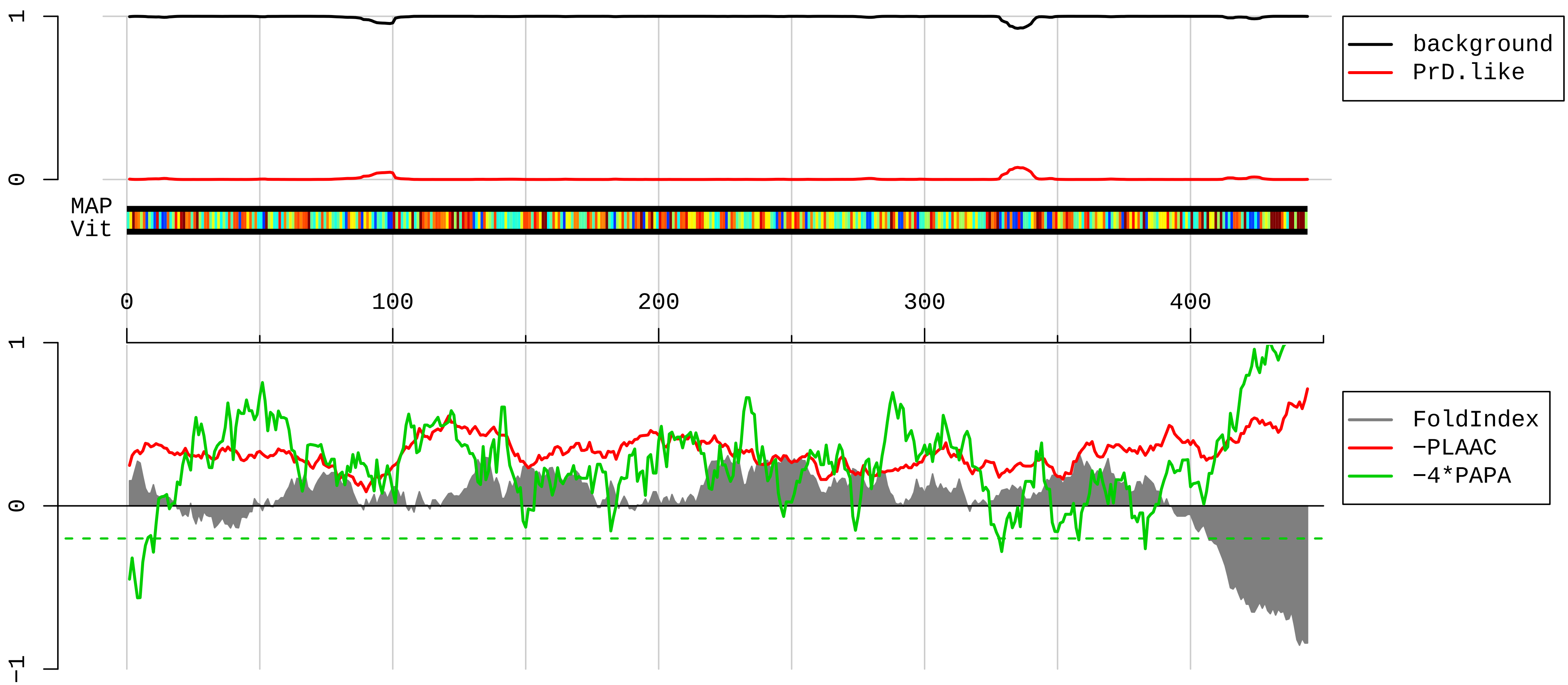


```
  1  MREIVHIQAGQCGNQIGAKFWEVISDEHGIDPTGTYHGDSDLQLDRISVYYNEATGGKYVPRAILVDLEPGTMDSVRSGPFGQIFRPDNFVFGQSGAGNN
101  WAKGHYTEGAELVDSVLDVVRKEAESCDCLQGFQLTHSLGGGTGSGMGTLLISKIREEYPDRIMNTFSVVPSPKVSDTVVEPYNATLSVHQLVENTDETY
201  CIDNEALYDICFRTLKLTTPTYGDLNHLVSATMSGVTTCLRFPGQLNADLRKLAVNMVPFPRLHFFMPGFAPLTSRGSQQYRALTVPELTQQVFDAKNMM
301  AACDPRHGRYLTVAAVFRGRMSMKEVDEQMLNVQNKNSSYFVEWIPNNVKTAVCDIPPRGLKMAVTFIGNSTAIQELFKRISEQFTAMFRRKAFLHWYTG
401  EGMDEMEFTEAESNMNDLVSEYQQYQDATAEEEEDFGEEAEEEA
```

**EDL26305.1 myosin Va, isoform CRA_a, partial [Mus musculus]**

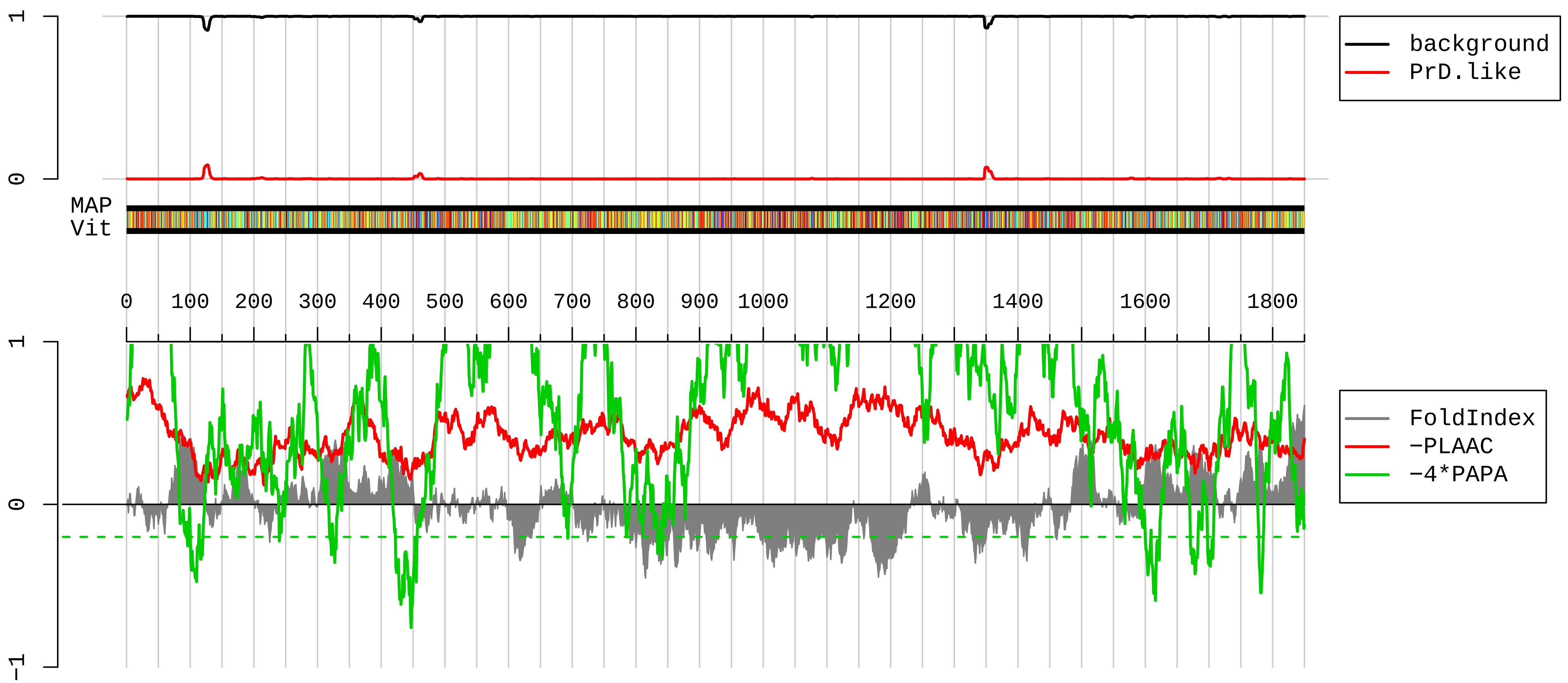


```
   1 SELYTKFARVWIPDPEEVWKSAELLKDYKPGDKVLLLHLEEGKDLEYRLDPKTGELPHLRNPDILVGENDLTALSYLHEPAVLHNLRVRFIDSKLIYTYC
 101 GIVLVAINPYEQLPIYGEDIINAYSGQNMGDMDPHIFAVAEEAYKQMARDERNQSIIVSGESGAGKTVSAKYAMRYFATVSGSASEANVEEKVLASNPIM
 201 ESIGNAKTTRNDNSSRFGKYIEIGFDKRYRIIGANMRTYLLEKSRVVFQAEEERNYHIFYQLCASAKLPEFKMLRLGNADSFHYTKQGGSPMIEGVDDAK
 301 EMAHTRQACTLLGISESYQMGIFRILAGILHLGNVGFASRDSDSCTIPPKHEPLTIFCDLMGVDYEEMCHWLCHRKLATATETYIKPISKLQATNARDAL
 401 AKHIYAKLFNWIVDHVNQALHSAVKQHSFIGVLDIYGFETFEINSFEQFCINYANEKLQQQFNMHVFKLEQEEYMKEQIPWTLIDFYDNQPCINLIESKL
 501 GILDLLDEECKMPKGTDDTWAQKLYNTHLNKCALFEKPRMSNKAFIIKHFADKVEYQCEGFLEKNKDTVFEEQIKVLKSSKFKMLPELFQDDEKAISPTS
 601 ATSSGRTPLTRVPVKPTKGRPGQTAKEHKKTVGHQFRNSLHLLMETLNATTPHYVRCIKPNDFKFPFTFDEKRAVQQLRACGVLETIRISAAGFPSRWTY
 701 QEFFSRYRVLMKQKDVLGDRKQTCKNVLEKLILDKDKYQFGKTKIFFRAGQVAYLEKLRADKLRAACIRIQKTIRGWLLRKRYLCMQRAAITVQRYVRGY
 801 QARCYAKFLRRTKAATTIQKYWRMYVVRRRYKIRRAATIVIQSYLRGYLTRNRYRKILREYKAVIIQKRVRGWLARTHYKRTMKAIVYLQCCFRRMMAKR
 901 ELKKLKIEARSVERYKKLHIGMENKIMQLQRKVDEQNKDYKCLMEKLTNLEGVYNSETEKLRNDVERLQLSEEEAKVATGRVLSLQEEIAKLRKDLEQTR
1001 SEKKSIEERADKYKQETDQLVSNLKEENTLLKQEKETLNHRIVEQAKEMTETMERKLVEETKQLELDLNDERLRYQNLLNEFSRLEERYDDLKEEMTLML
1101 NVPKPGHKRTDSTHSSNESEYTFSSEFAETEDIAPRTEEPIEKKVPLDMSLFLKLQKRVTELEQEKQLMQDELDRKEEQVFRSKAKEEERPQIRGAELEY
1201 ESLKRQELESENKKLKNELNELRKALSEKSAPEVTAPGAPAYRVLMEQLTSVSEELDVRKEEVLILRSQLVSQKEAIQPKDDKNTMTDSTILLEDVQKMK
1301 DKGEIAQAYIGLKETNRLLESQLQSQKRSHENEAEALRGEIQSLKEENNRQQQLLAQNLQLPPEARIEASLQHEITRLTNENLYFEELYADDPKKYQSYR
1401 ISLYKRMIDLMEQLEKQDKTVRKLKKQLKVFAKKIGELEVGQMENISPGQIIDEPIRPVNIPRKEKDFQGMLEYKREDEQKLVKNLILELKPRGVAVNLI
1501 PGLPAYILFMCVRHADYLNDDQKVRSLLTSTINSIKKVLKKRGDDFETVSFWLSNTCRFLHCLKQYSGEEGFMKHNTSRQNEHCLTNFDLAEYRQVLSDL
1601 AIQIYQQLVRVLENILQPMIVSGMLEHETIQGVSGVKPTGLRKRTSSIADEGTYTLDSILRQLNSFHSVMCQHGMDPELIKQVVKQMFYIVGAITLNNLL
1701 LRKDMCSWSKGMQIRYNVSQLEEWLRDKNLMNSGAKETLEPLIQAAQLLQVKKKTDDDAEAICSMCNALTTAQIVKVLNLYTPVNEFEERVSVSFIRTIQ
1801 MRLRDRKDSPQLLMDAKHIFPVTFPFNPSSLALETIQIPASLGLGFIARV
```

**AAI22884.1 GTPase activating protein (SH3 domain) binding protein 2 [Mus musculus]**

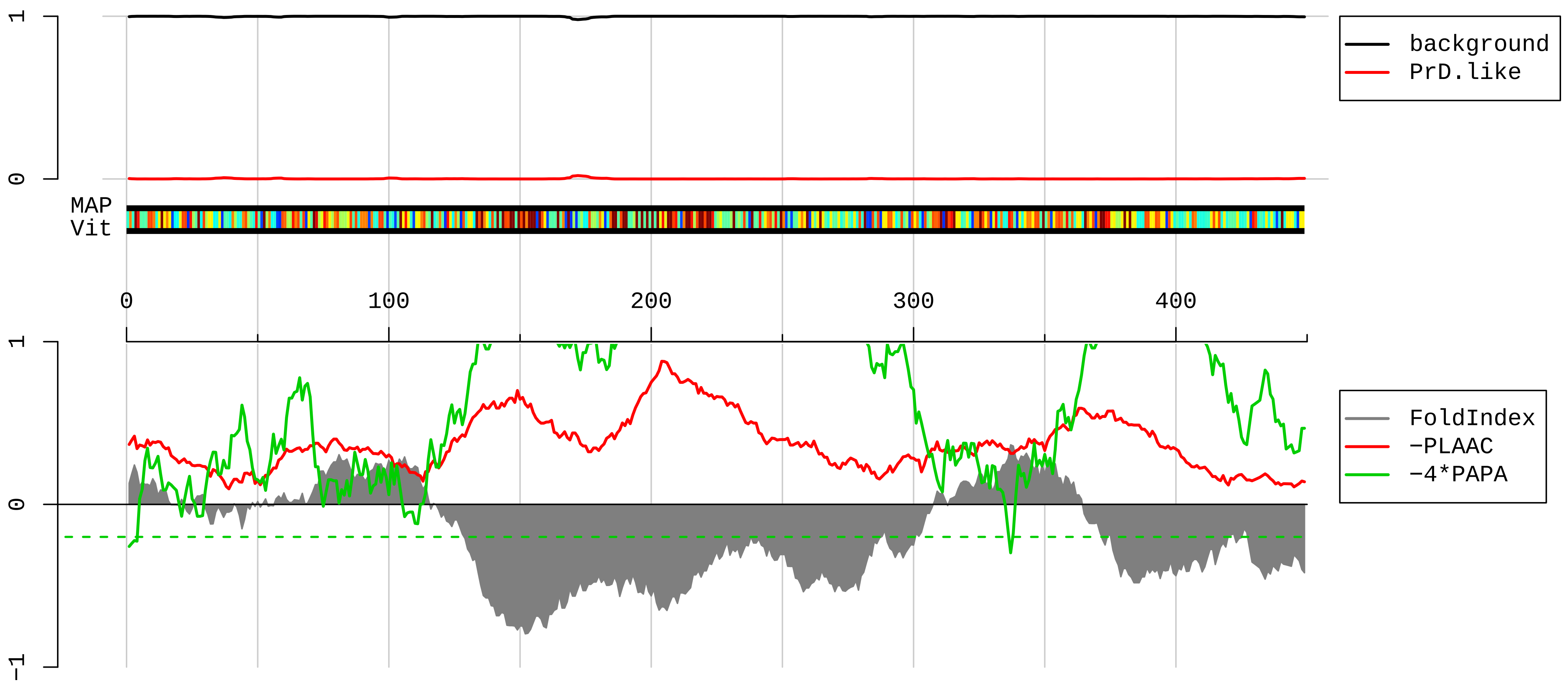


```
  1  MVMEKPSPLLVGREFVRQYYTLLNKAPEYLHRFYGRNSSYVHGGVDASGKPQEAVYGQNDIHHKVLSLNFSECHTKIRHVDAHATLSDGVVVQVMGLLSN
101  SGQPERKFMQTFVLAPEGSVPNKFYVHNDMFRYEDEVFGDSEPELDEESEDEVEEEQEDRQPSPEPVQENANSAYYDAHPVTNGIEEPLEESSHEPEPEP
201  ESETKTEELKPQVEEKHLEELEEKSATPPPAEPASLPQEPPKPRVDAKPEVQSQPPRVREQRPRERPGFPPRGPRPGRGDMEQNDSDNRRIIRYPDSHQL
301  FVGNLPHDIDENELKEFFMSFGNVVELRINTKGVGGKLPNFGFVVFDDSEPVQRILIAKPIMFRGEVRLNVEEKKTRAARERETRGGGDDRRDIRRNDRG
401  PGGPRGIVGGGMMRDRDGRGPPPRGGMTQKLGSGRGTGQMEGRFTGQRR
```

**AAH99371.1 Actin, gamma, cytoplasmic 1 [Mus musculus]**

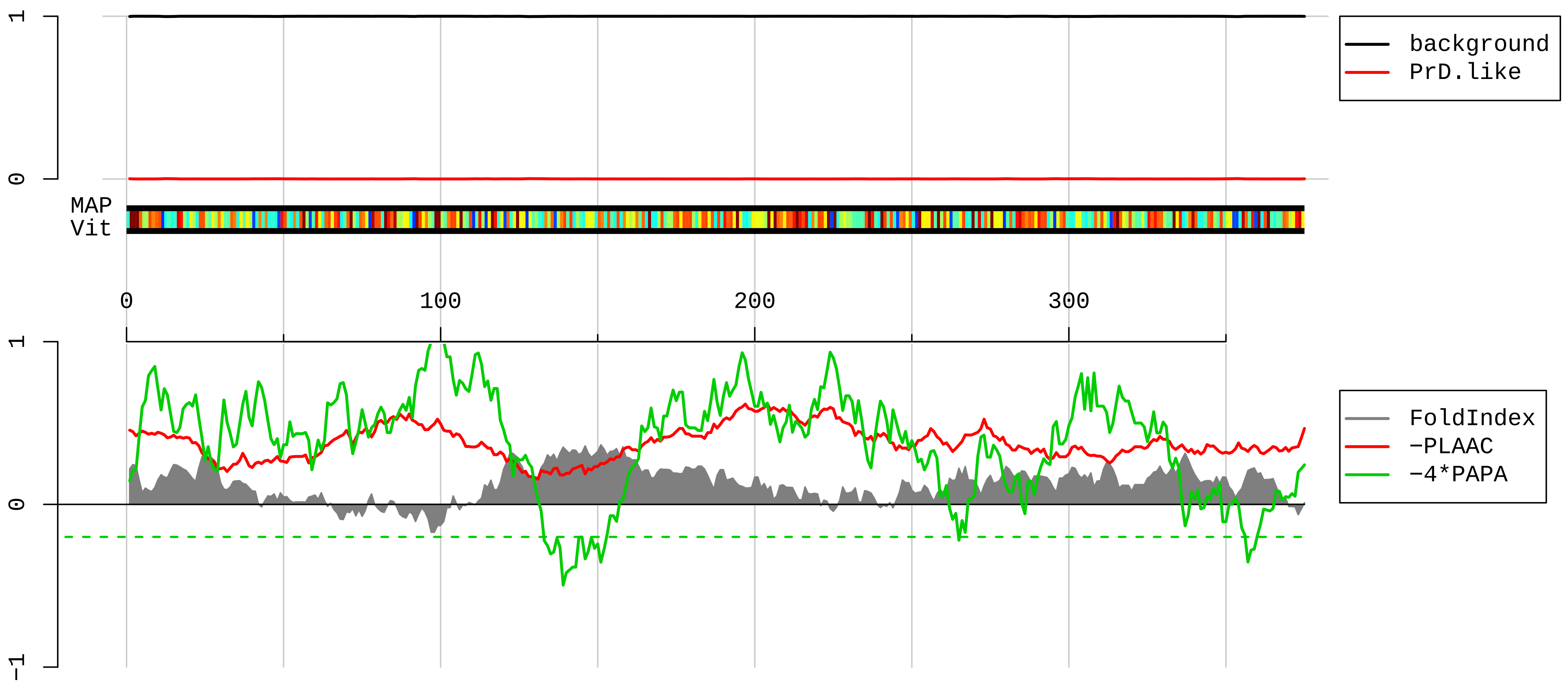


1 MEEEIAALVIDNGSGMCKAGFAGDDAPRAVFPSIVGRPRHQGVMVGMGQKDSYVGDEAQSKRGILTLKYPIEHGIVTNWDDMEKIWHHTFYNELRVAPEE
101 HPVLLTEAPLNPKANREKMTQIMFETFNTPAMYVAIQAVLSLYASGRTTGIVMDSGDGVTHTVPIYEGYALPHAILRLDLAGRDLTDYLMKILTERGYSF
201 TTTAEREIVRDIKEKLCYVALDFEQEMATAASSSSLEKSYELPDGQVITIGNERFRCPEALFQPSFLGMESCGIHETTFNSIMKCDVDIRKDLYANTVLS
301 GGTTMYPGIADRMQKEITALAPSTMKIKIIAPPERKYSVWIGGSILASLSTFQQMWISKQEYDESGPSIVHRKCF

**AAH94900.1 Hspa8 protein [Mus musculus]**

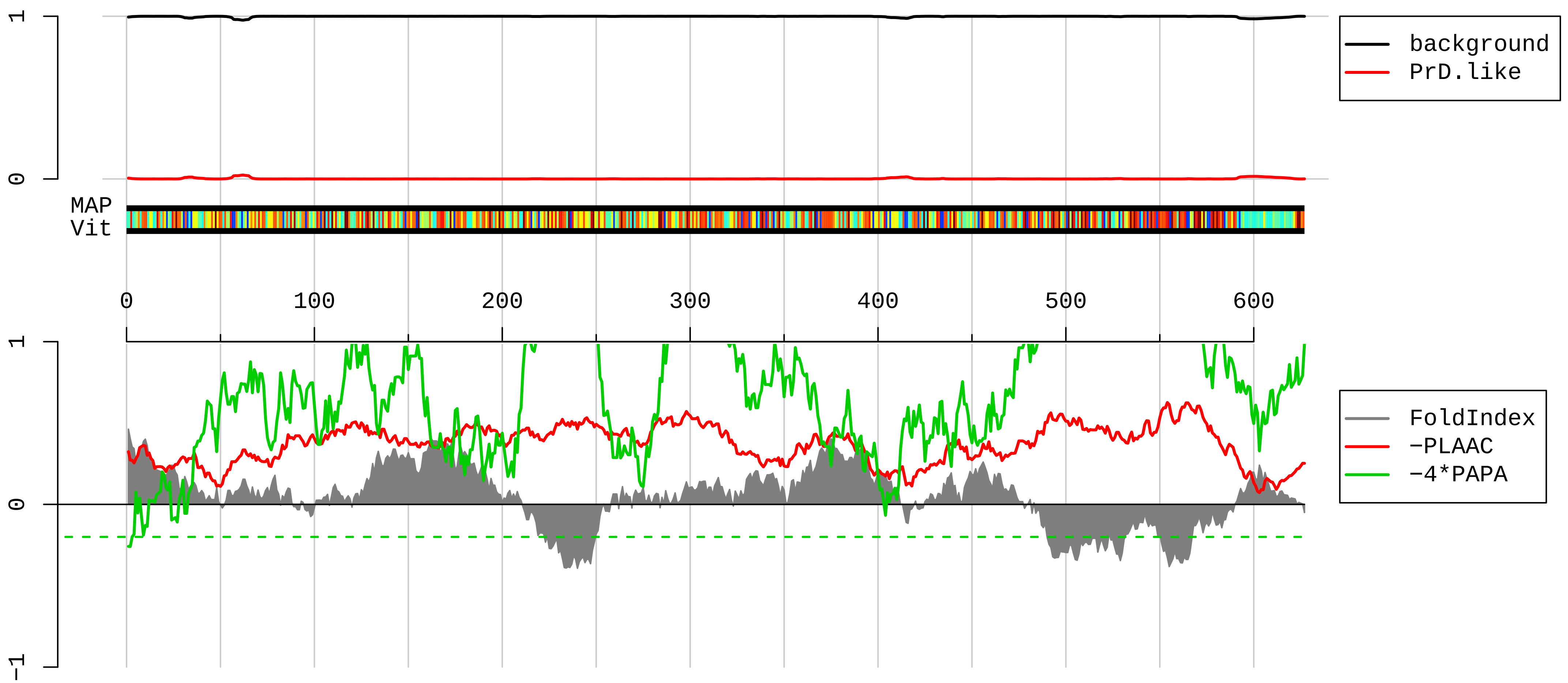


```
1   MSKGPAVGIDLGTTYSCVGVFQHGKVEIIANDQGNRTTPSYVAFTDTERLIGDAAKNQVAMNPTNTVFDAKRLIGRRFDDAVVQSDMKHWPFMVVNDAGR
101 PKVQVEYKGETKSFYPEEVSSMVLTKMKEIAEAYLGKATKDAGTIAGLNVLRIINEPTAAAIAYGLDKKVGAERNVLIFDLGGGTFDVSILTIEDGIFEV
201 KSTAGDTHLGGEDFDNRMVNHFIAEFKRKHKKDISENKRAVRRLRTACERAKRTLSSSTQASIEIDSLYEGIDFYTSITRARFEELNADLFRGTLDPVEK
301 ALRDAKLDKSQIHDIVLVGGSTRIPKIQKLLQDFFNGKELNKSINPDEAVAYGAAVQAAILSGDKSENVQDLLLLDVTPLSLGIETAGGVMTVLIKRNTT
401 IPTKQTQTFTTYSDNQPGVLIQVYEGERAMTKDNNLLGKFELTGIPPAPRGVPQIEVTFDIDANGILNVSAVDKSTGKENKITITNDKGRLSKEDIERMV
501 QEAEKYKAEDEKQRDKVSSKNSLESYAFNMKATVEDEKLQGKINDEDKQKILDKCNEIISWLDKNQTAEKEEFEHQQKELEKVCNPIITKLYQSAGGMPG
601 GMPGGFPGGGAPPSGGASSGPTIEEVD
```

**AAH94059.1 Ribosomal protein L3 [Mus musculus]**

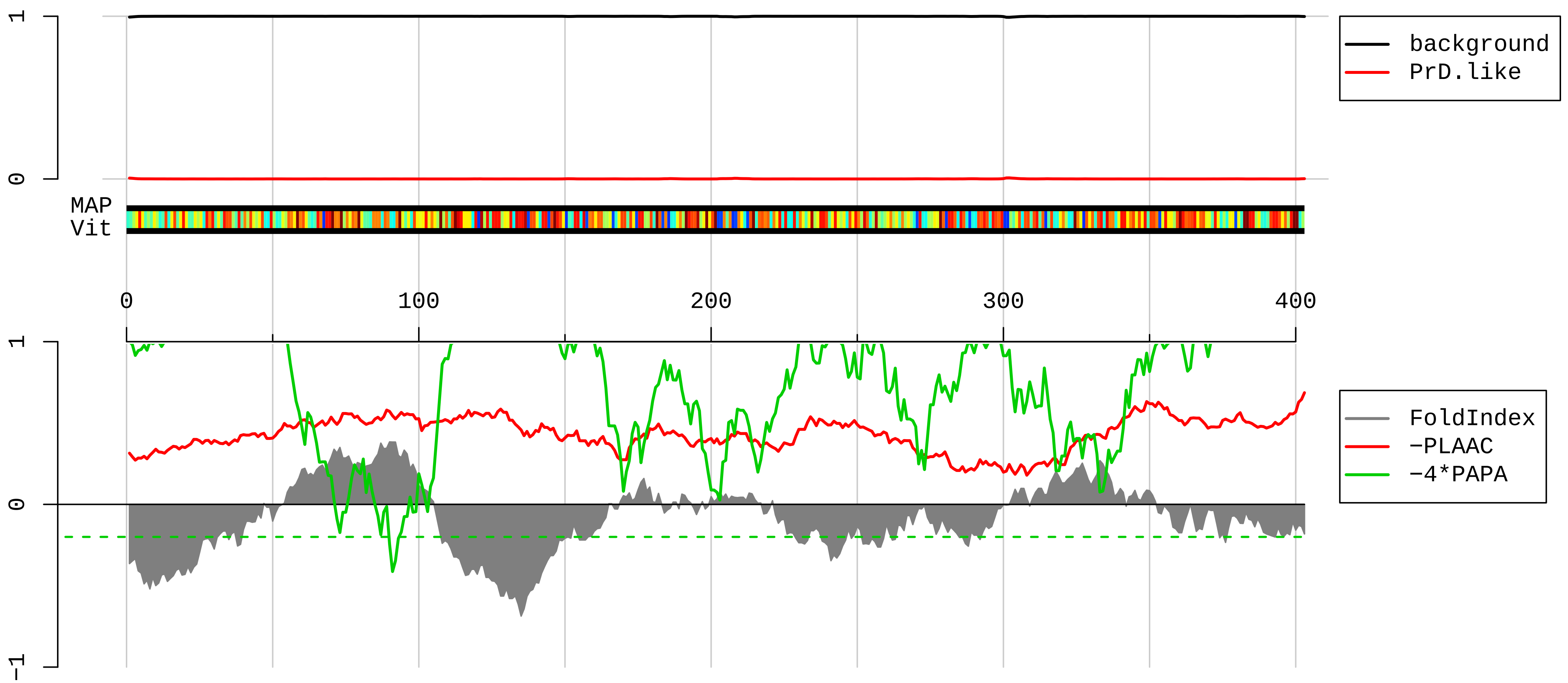


```
  1 MSHRKFSAPRHGSLGFLPRKRSSRHRGKVKSFPKDDASKPVHLTAFLGYKAGMTHIVREVDRPGSKVNKKEVVEAVTIVETPPMVVVGIVGYVETPRGLR
101 TFKTVFAEHISDECKRRFYKNWHKSKKKAFTKYCKKWQDDTGKKQLEKDFNSMKKYCQVIRIIAHTQMRLLPLRQKKAHLMEIQVNGGTVAEKLDWARER
201 LEQQVPVNQVFGQDEMIDVIGVTKGKGYKGVTSRWHTKKLPRKTHRGLRKVACIGAWHPARVAFSVARAGQKGYHHRTEINKKIYKIGQGYLIKDGKLIK
301 NNASTDYDLSDKSINPLGGFVHYGEVTNDFIMLKGCVVGTKKRVLTLRKSLLVQTKRRALEKIDLKFIDTTSKFGHGRFQTMEEKKAFMGPLKKDRIAKE
401 EGA
```

#### AAH09100.1 ribosomal protein S4, X-linked [synthetic construct]

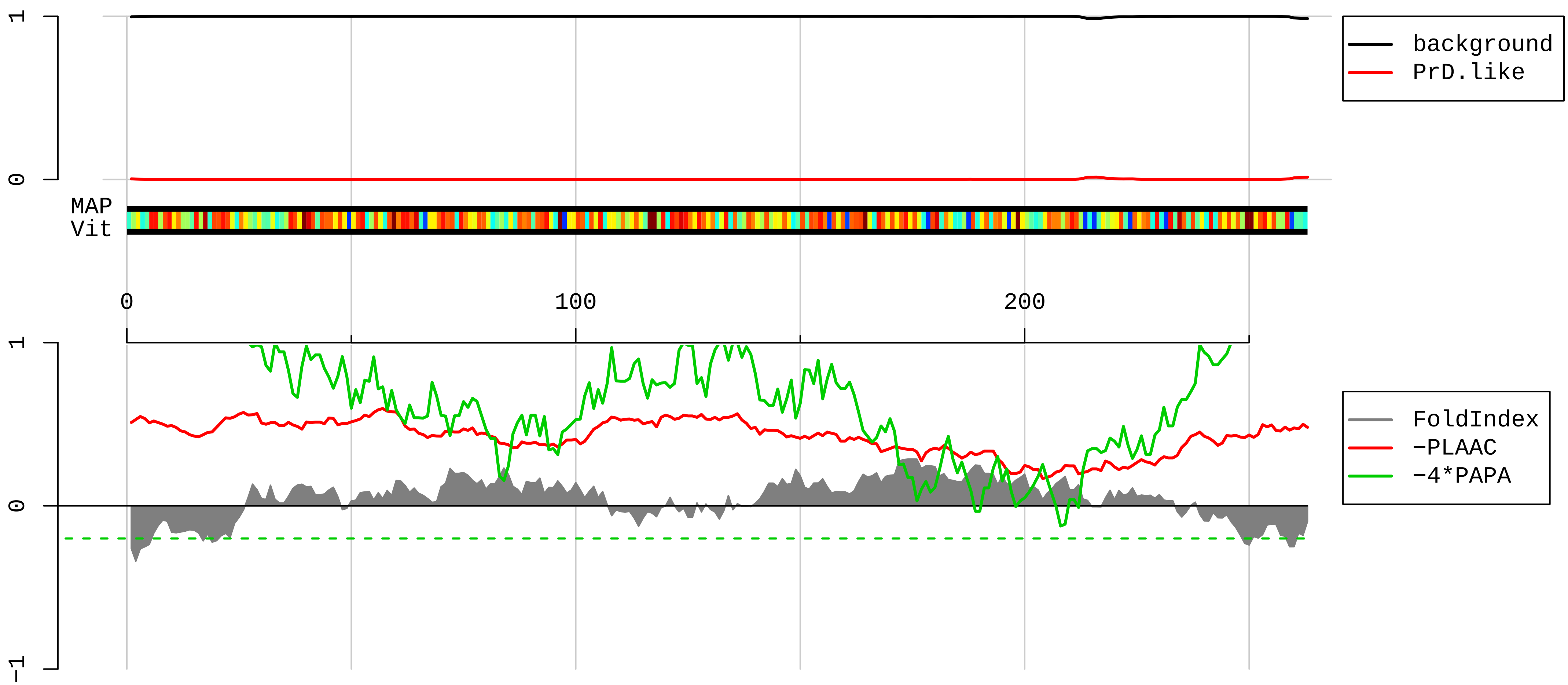


1 MARGPKKHLKRVAAPKHWML<u>DKLTGVFAPRPST</u>GPHKLRECLPLIIFLRNRLKYALTGDEVKKICMQRFIKIDGKVRTDITYPAGFMDVISIDKTGENFR
101 LIYDTKGRFAVHRITPE<u>EAKYKLCKVR</u>KIFVGTKGIPHLVTHDARTIRYPDPLIKVNDTIQIDLETGKITDFIKFDTGNLCMVTGGANLGRIGVITNRER
201 HPGSFDVVHVKDANGNSFATRLSNIFVIGK<u>GNKPWISLPRGKG</u>IRLTIAEERDKRLAAKQSSG

**AAH50927.1 Heat shock protein 5 [Mus musculus]**

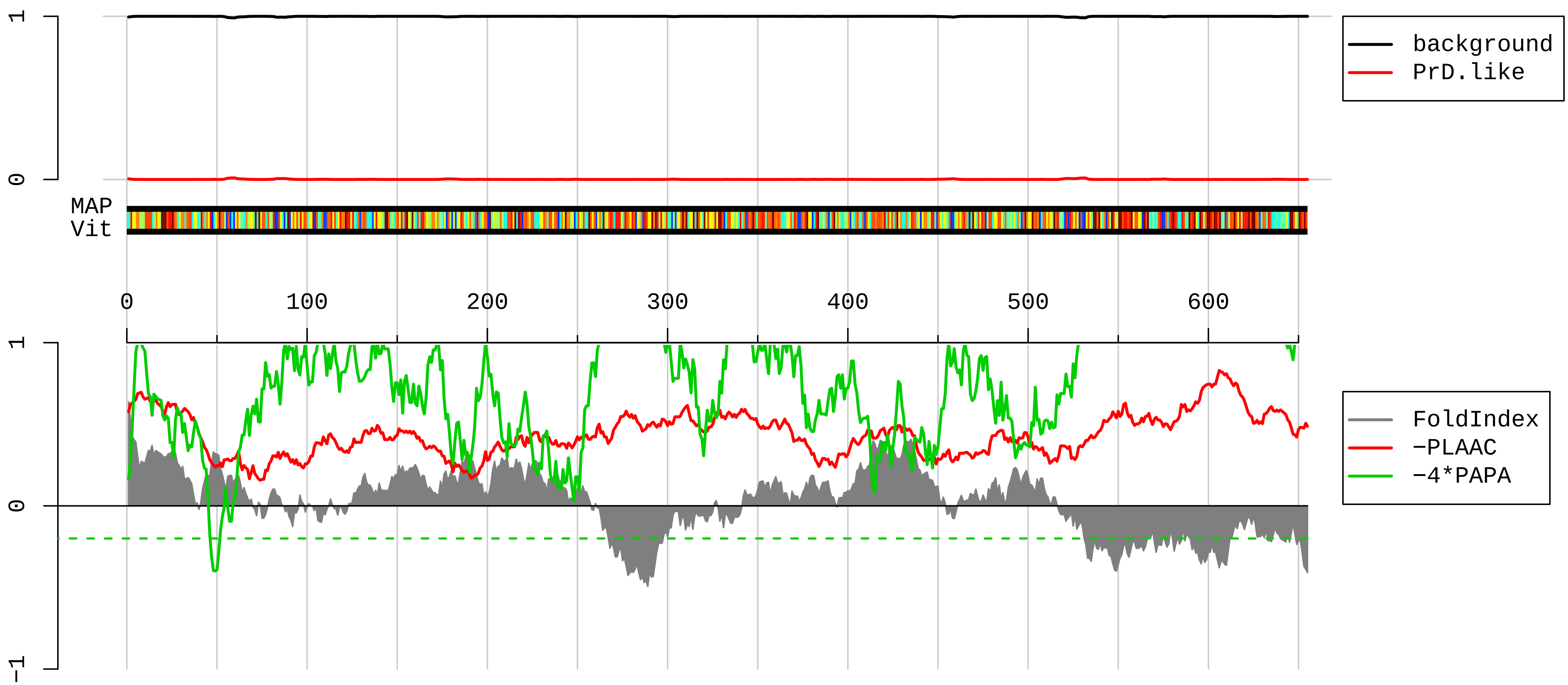


```
1   MMKFTVVAAALLLLGAVRAEEEDKKEDVGTVVGIDLGTTYSCVGVFKNGRVEIIANDQGNRITPSYVAFTPEGERLIGDAAKNQLTSNPENTVFDAKRLI
101 GRTWNDPSVQQDIKFLPFKVVEKKTKPYIQVDIGGGQTKTFAPEEISAMVLTKMKETAEAYLGKKVTHAVVTVPAYFNDAQRQATKDAGTIAGLNVMRII
201 NEPTAAAIAYGLDKREGEKNILVFDLGGGTFDVSLLTIDNGVFEVVATNGDTHLGGEDFDQRVMEHFIKLYKKKTGKDVRKDNRAVQKLRREVEKAKRAL
301 SSQHQARIEIESFFEGEDFSETLTRAKFEELNMDLFRSTMKPVQKVLEDSDLKKSDIDEIVLVGGSTRIPKIQQLVKEFFNGKEPSRGINPDEAVAYGAA
401 VQAGVLSGDQDTGDLVLLDVCPLTLGIETVGGVMTKLIPRNTVVPTKKSQIFSTASDNQPTVTIKVYEGERPLTKDNHLLGTFDLTGIPPAPRGVPQIEV
501 TFEIDVNGILRVTAEDKGTGNKNKITITNDQNRLTPEEIERMVNDAEKFAEEDKKLKERIDTRNELESYAYSLKNQIGDKEKLGGKLSSEDKETMEKAVE
601 EKIEWLESHQDADIEDFKAKKKELEEIVQPIISKLYGSGGPPPTGEEDTSEKDEL
```

**EDL42022.1 internexin neuronal intermediate filament protein, alpha [Mus musculus]**

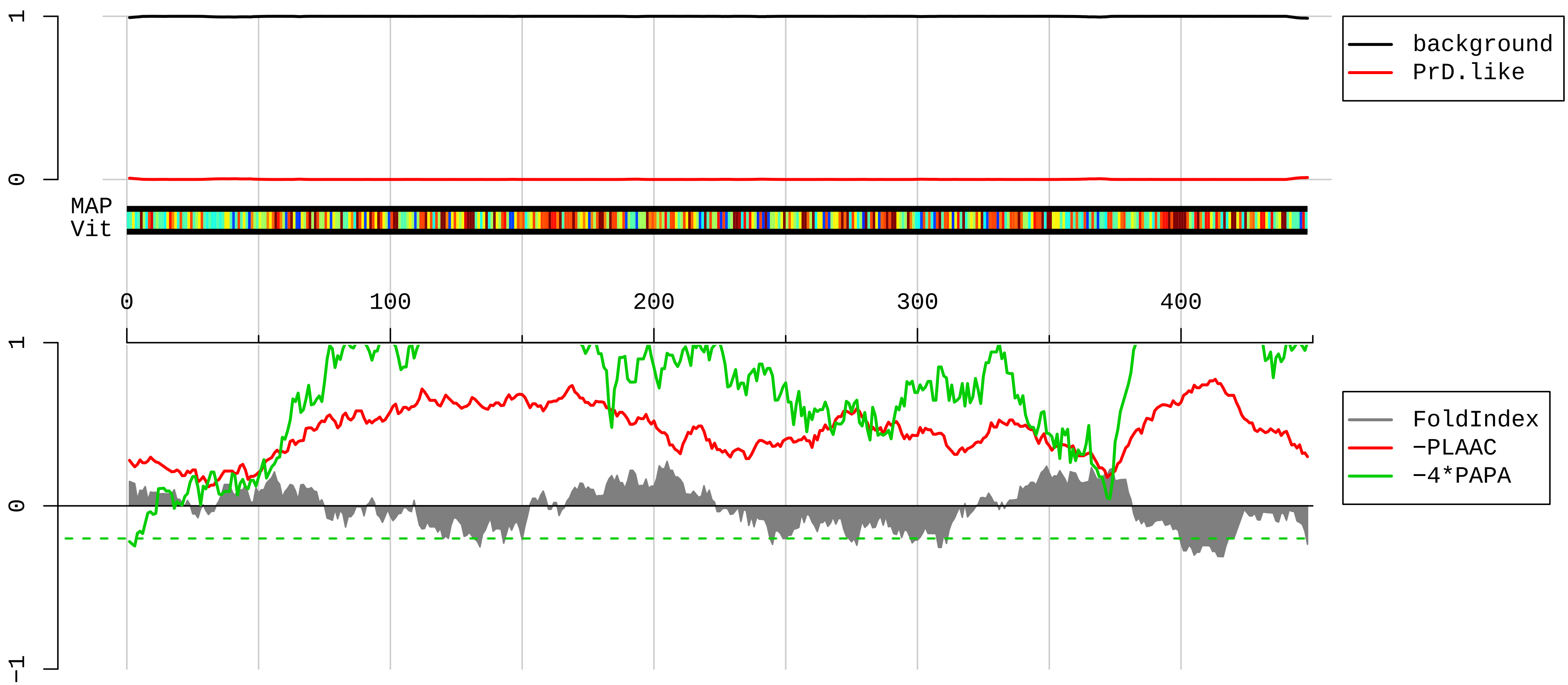


1 MSFGSEHYLCSASSYRKVFGDSSRLSARLSGPGGSGSFRSQSLSRSNVASTAAVFIEKVHQLETQNRALEAELAALRQRHAEPSRVGELFQRELRELRAQ

101 LEEASSARAQALLERDGLAEEVQRLRARCEEESRGREGAERALKAQQRDVDGATLARLDLEKKVESLLDELAFVRQVHDEEVAELLATLQASSQAAAEVD

201 VAVAKPDLTSALREIRAQYESLAAKNLQSAEEWYKSKFANLNEQAARSTEAIRASREEIHEYRRQLQARTIEIEGLRGANESLERQILELEERHSAEVAG

301 YQDSIGQLESDLRNTKSEMARHLREYQDLLNVKMALDIEIAAYRKLLEGEETRFSTGGLSISGLNPLPNPSYLLPPRILSSTASKVSSTGLSLKKEDEEE

401 EEDASKEVSKKTSKVGEGFEETLGEAVISTKKTGKSATEESTSSSQKM

**AAH99384.1 Ribosomal protein, large, P0 [Mus musculus]**

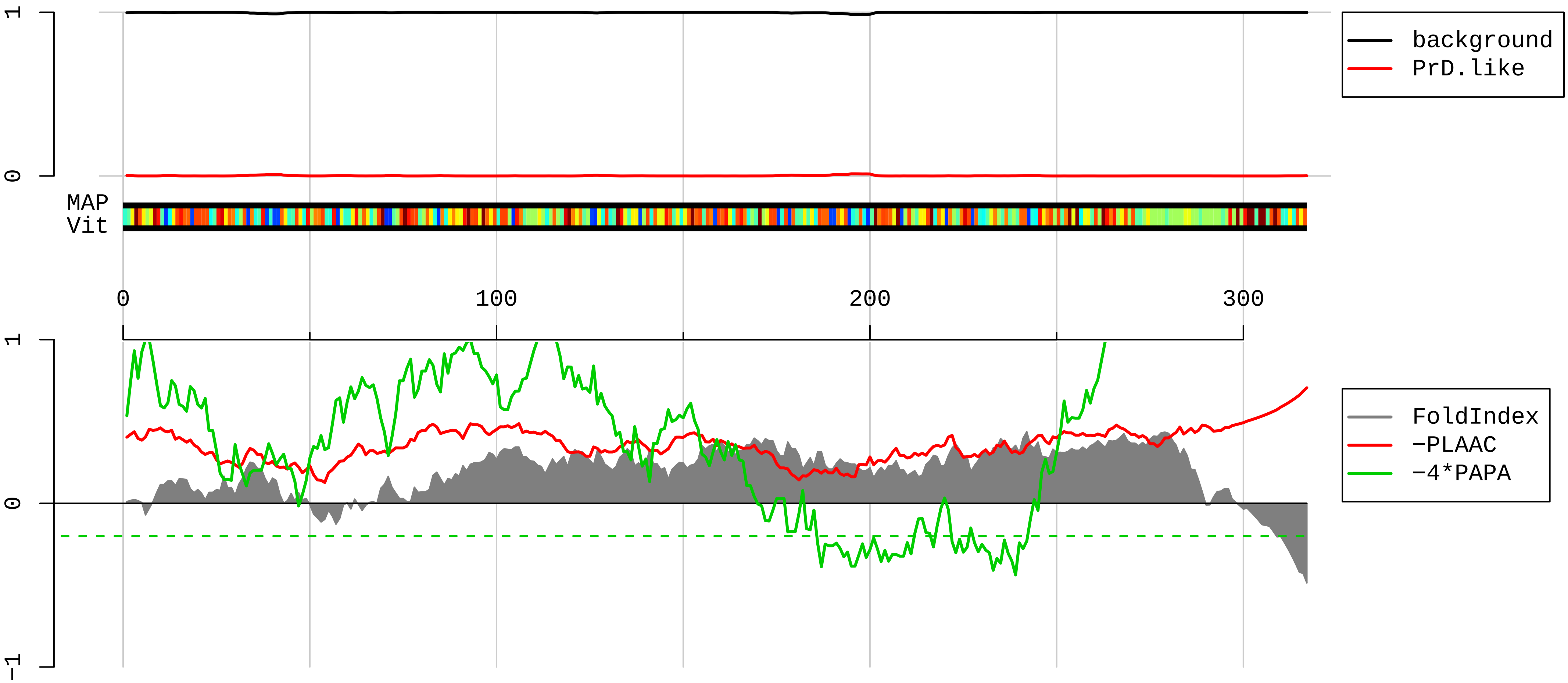


1 MPREDRATWKSNYFLKIIQLLDDYPKCFIVGADNVGSKQMQQIRMSLRGKAVVLMGKNTMMRKAIRGHLENNPALEKLLPHIRGNVGFVFTKEDLTEIRD
101 MLLANKVPAAARAGAIAPCEVTVPAQNTGLGPEKTSFFQALGITTKISRGTIEILSDVQLIKTGDKVGASEATLLNMLNISPFSFGLIIQQVFDNGSIYN
201 PEVLDITEQALHSRFLEGVRNVASVCLQIGYPTVASVPHSIINGYKRVLALSVETEYTFPLAEKVKAFLADPSAFAAAAPAAAATTAAPAAAAAPAKAEA
301 KEESEESDEDMGFGLFD

**AAH46760.1 Guanine nucleotide binding protein (G protein), beta polypeptide 2 like 1 [Mus musculus]**

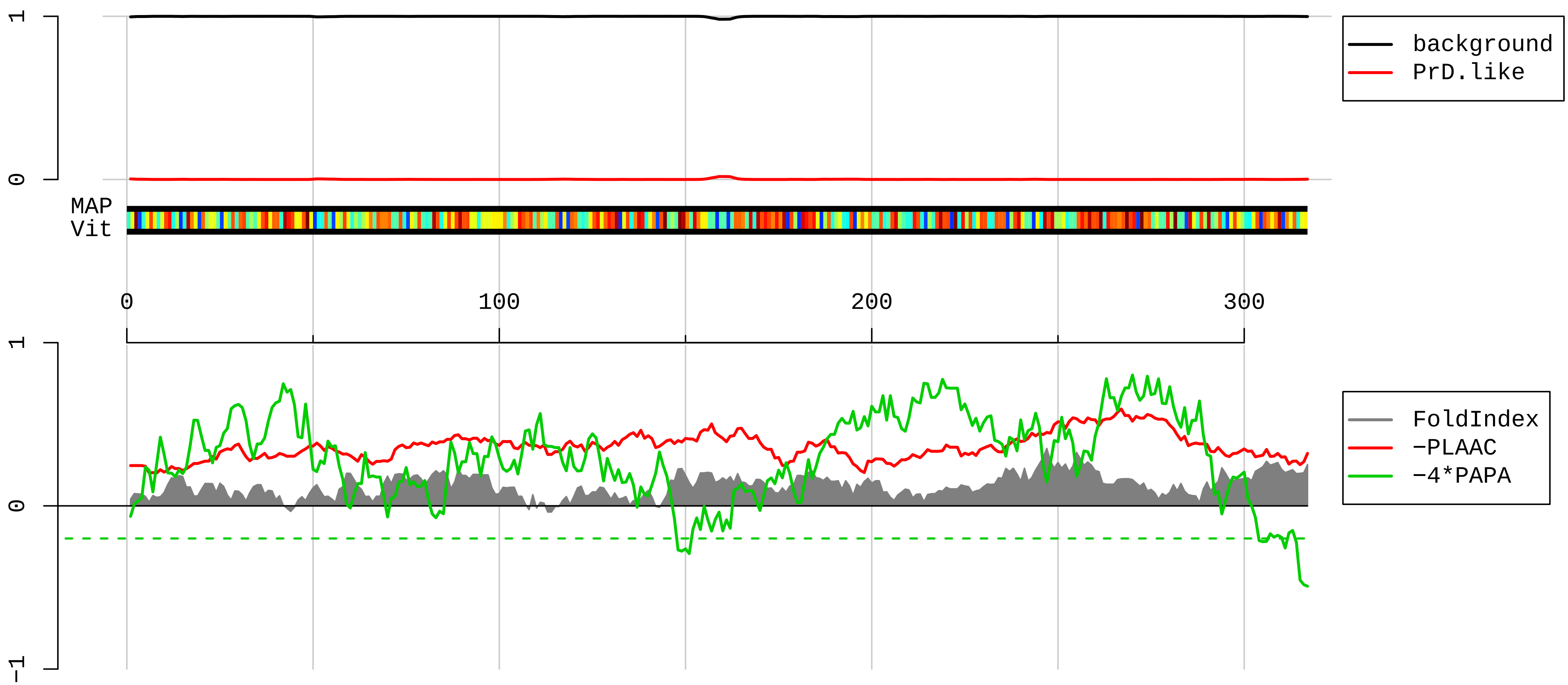


```
  1  MTEQMTLRGTLKGHNGWVTQIATTPQFPDMILSASRDKTIIMWKLTRDETNYGIPQRALRGHSHFVSDVVISSDGQFALSGSWDGTLRLWDLTTGTTTRR
101  FVGHTKDVLSVAFSSDNRQIVSGSRDKTIKLWNTLGVCKYTVQDESHSEWVSCVRFSPNSSNPIIVSCGWDKLVKVWNLANCKLKTNHIGHTGYLNTVTV
201  SPDGSLCASGGKDGQAMLWDLNEGKHLYTLDGGDIINALCFSPNRYWLCAATGPSIKIWDLEGKIIVDELKQEVISTSSKAEPPQCTSLAWSADGQTLFA
301  GYTDNLVRVWQVTIGTR
```

**AAH91752.1 Ribosomal protein L5 [Mus musculus]**

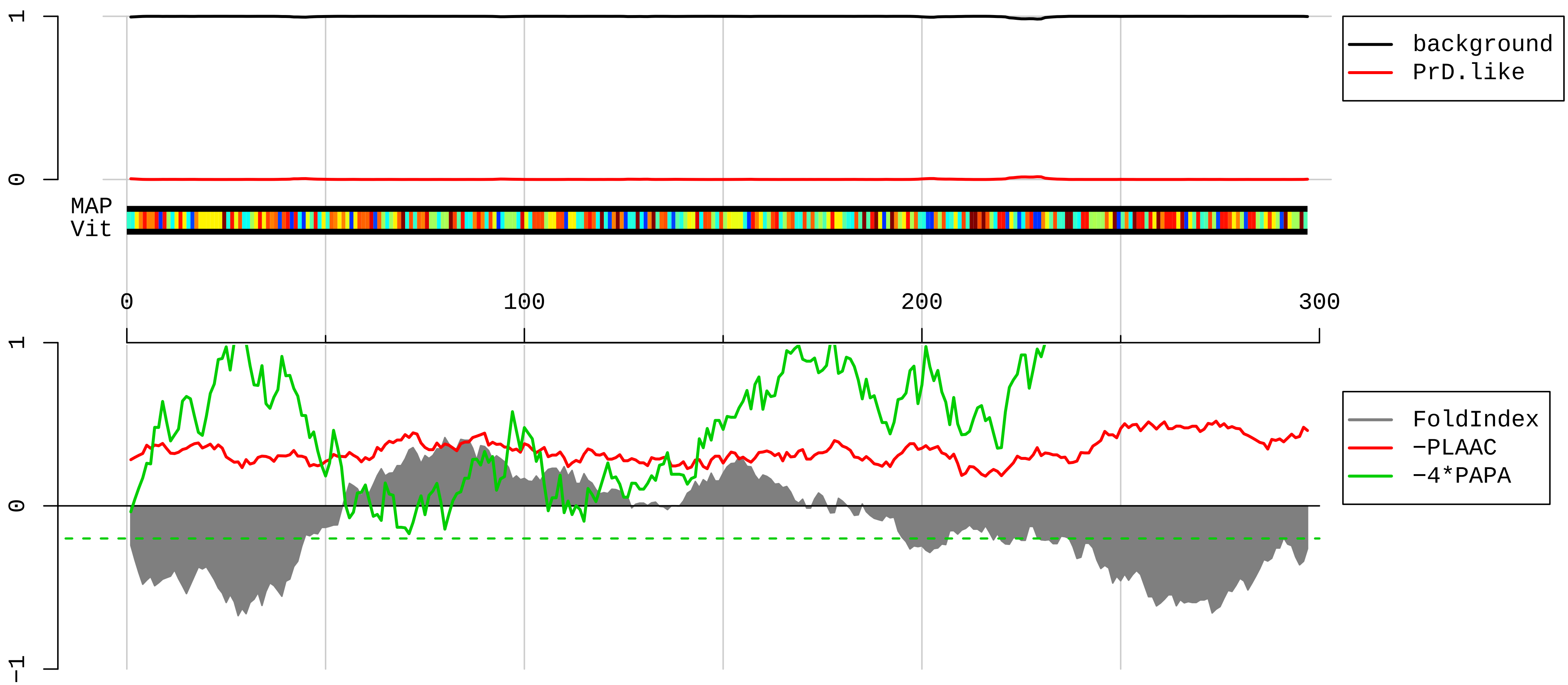


1 MGFVKVVKNKAYFKRYQVRFRRRREGKTDYYARKRLVIQDKNKYNTPKYRMIVRVTNRDIICQIAYARIEGDMIVCAAYAHELPKYGVKVGLTNYAAAYC
101 TGLLLARRLLNRFGMDKIYEGQVEVNGGEYNVESIDGQPGAFTCYLDAGLARTTTGNKVFGALKGAVDGGLSIPHSTKRFPGYDSESKEFNAEVHRKHIM
201 GQNVADYMRYLMEEDEDAYKKQFSQYIKNNVTPDMMEEMYKKAHAAIRENPVYEKKPKREVKKKRWNRPKMSLAQKKDRVAQKKASFLRAQERAAES

AAH92137.1 Ribosomal protein L30 [Mus musculus]

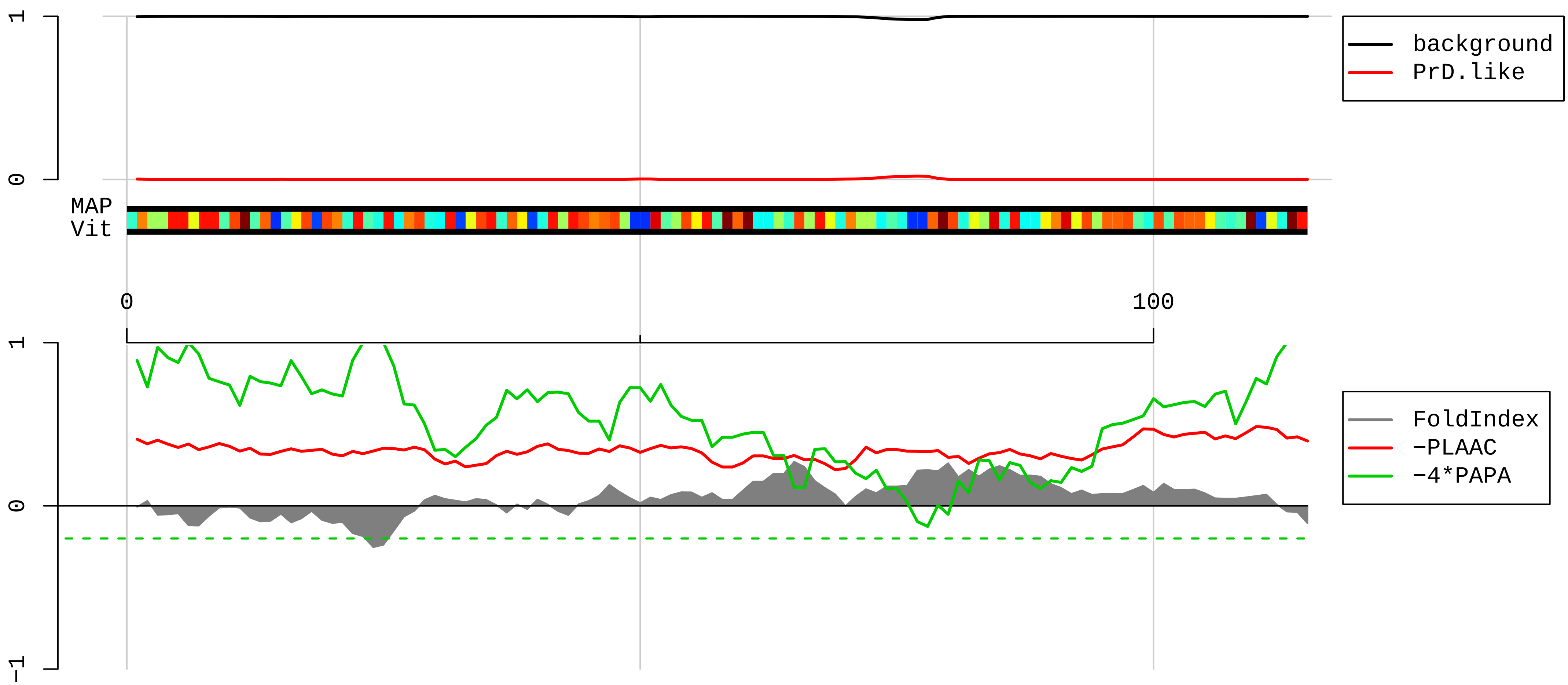


1 MVAAKKTKKSLESINSRLQLVMKSGKYVLGYKQTLKMIRQGKAKLVILANNCPALRKSEIEYYAMLAKTGVHHYSGNNIELGTACGKYYRVCTLAIIDPG
101 DSDIIRSMPEQTGEK

**AAH92249.1 Ribosomal protein L14 [Mus musculus]**

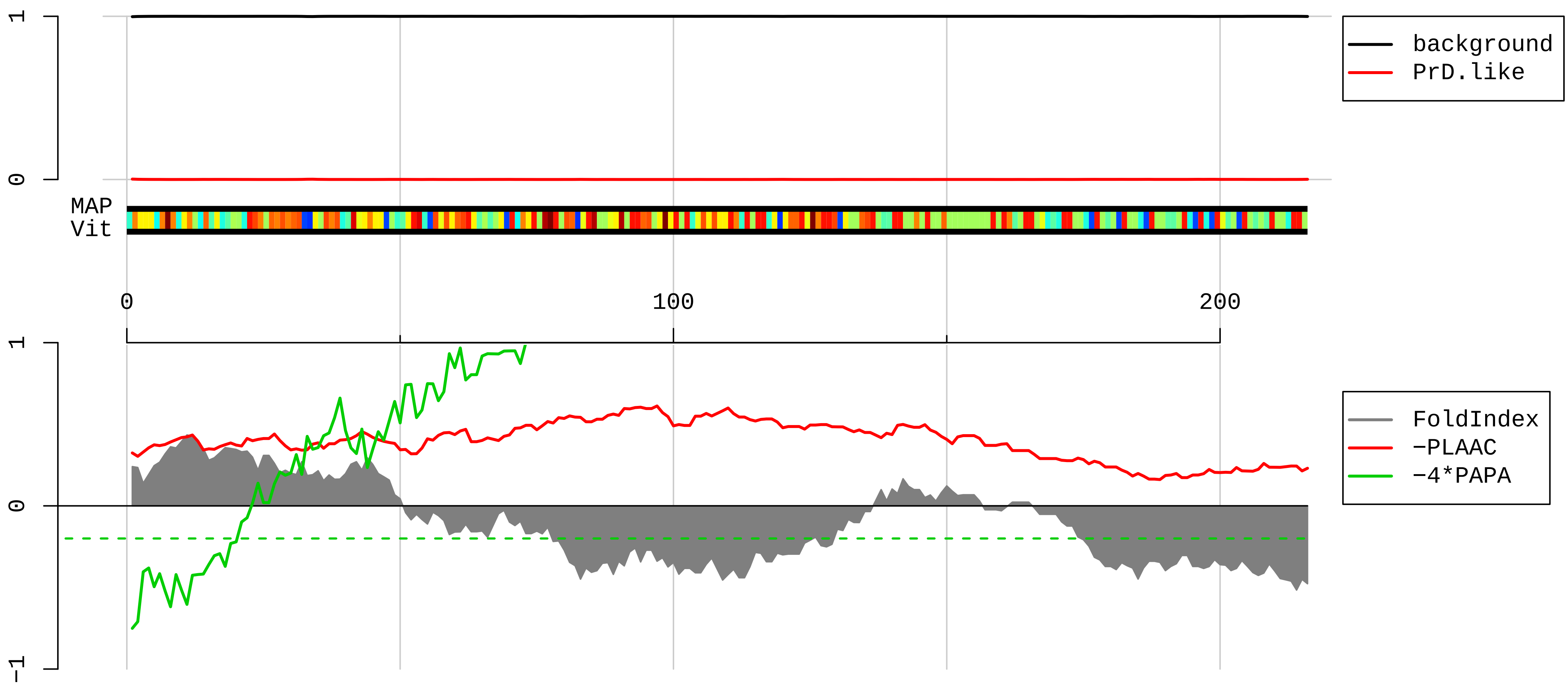


1 MVFRRYVEVGRVAYISFGPHAGKLVAIVDVIDQNRALVDGPCTRVRRQAMPFKCMQLTDFILKFPHSARQKYVRKAWEKADINTKWAATRWAKKIDARER
101 KAKMTDFDRFKVMKAKKMRNRIIKTEVKKLQRAAILKASPKKAAVAKAAIAAAAAAAAKAKVPAKKATGPGKKAAGQKAPAQKAAGQKAAPPAKGQKGQK
201 TPAQKAPAPKAAGKKA

**AAF88071.1 ribosomal protein L23 [Mus musculus]**

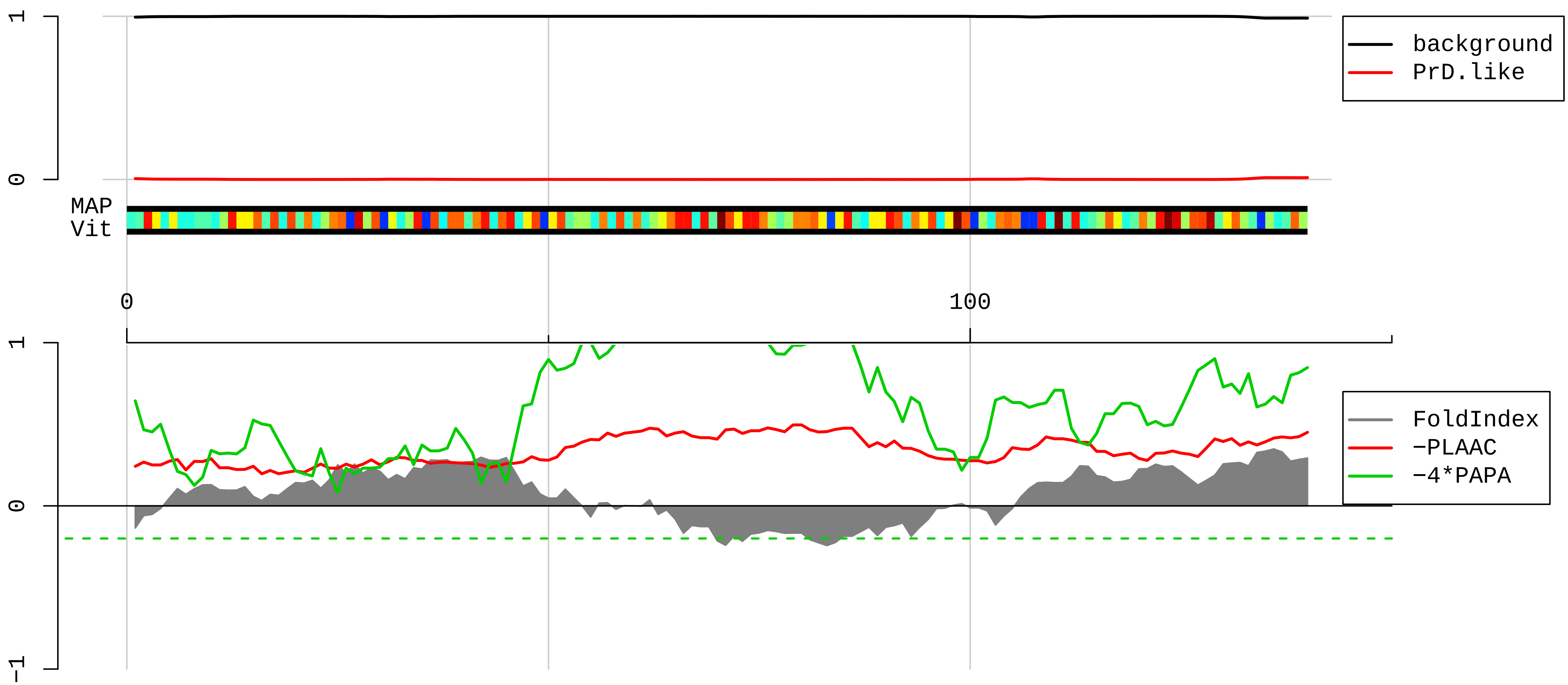


1 MSKRGRGGSSGAKFRISLGLPVGAVINCADNTGAKNLYIISVKGIKGRLNRLPAAGVGDMVMATVKKGKPELRKKVHPAVVIRQRKSYRRKDGVFLYFED
101 NAGVIVNNKGEMKGSAITGPVAKECADLWPRIASNAGSIA

**AAH86912.1 Ribosomal protein S21 [Mus musculus]**

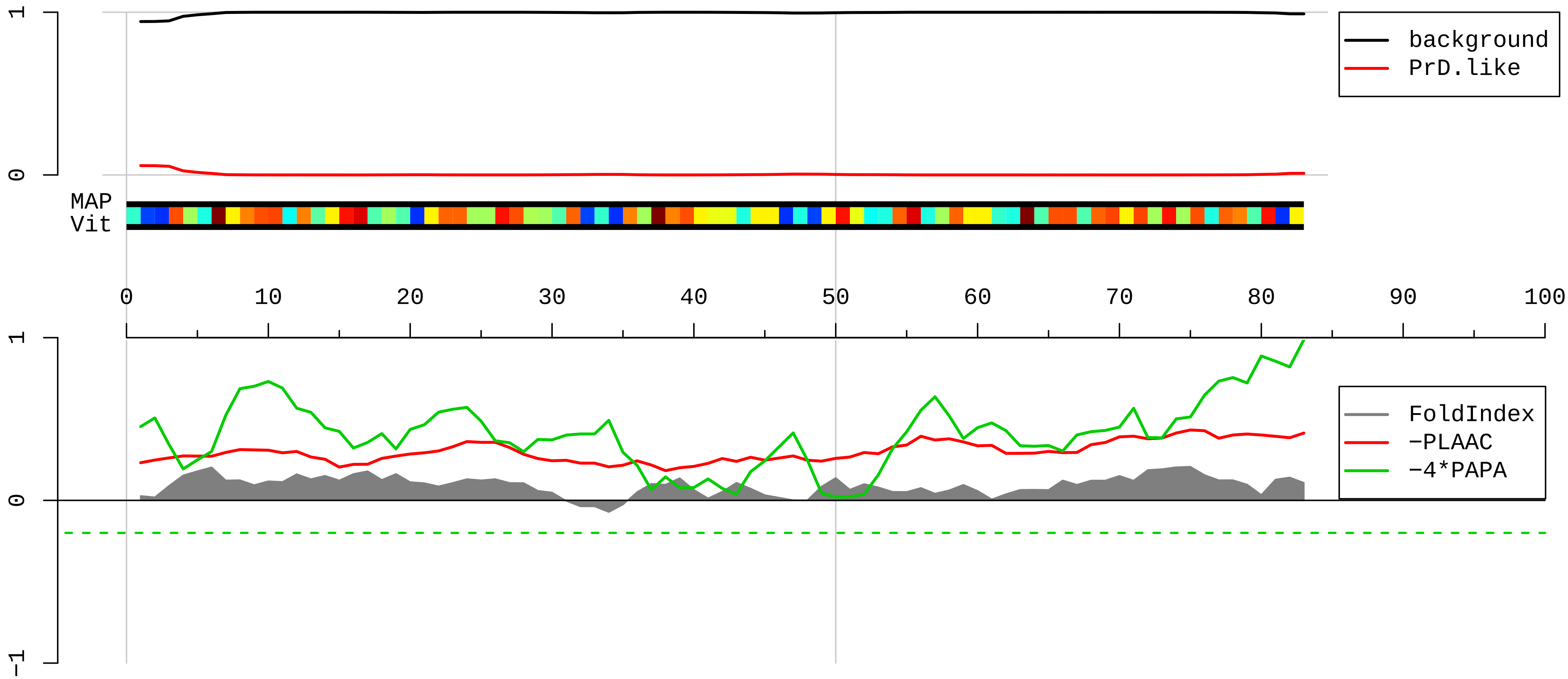


1 MQNDAGEFVDLYVPRKCSASNRIIAAKDHASIQMNVAEVDRTTGRFNGQFKTYGICGAIRRMGESDDSILRLAKADGIVSKNF

**AAH91769.1 Ribosomal protein L7A [Mus musculus]**

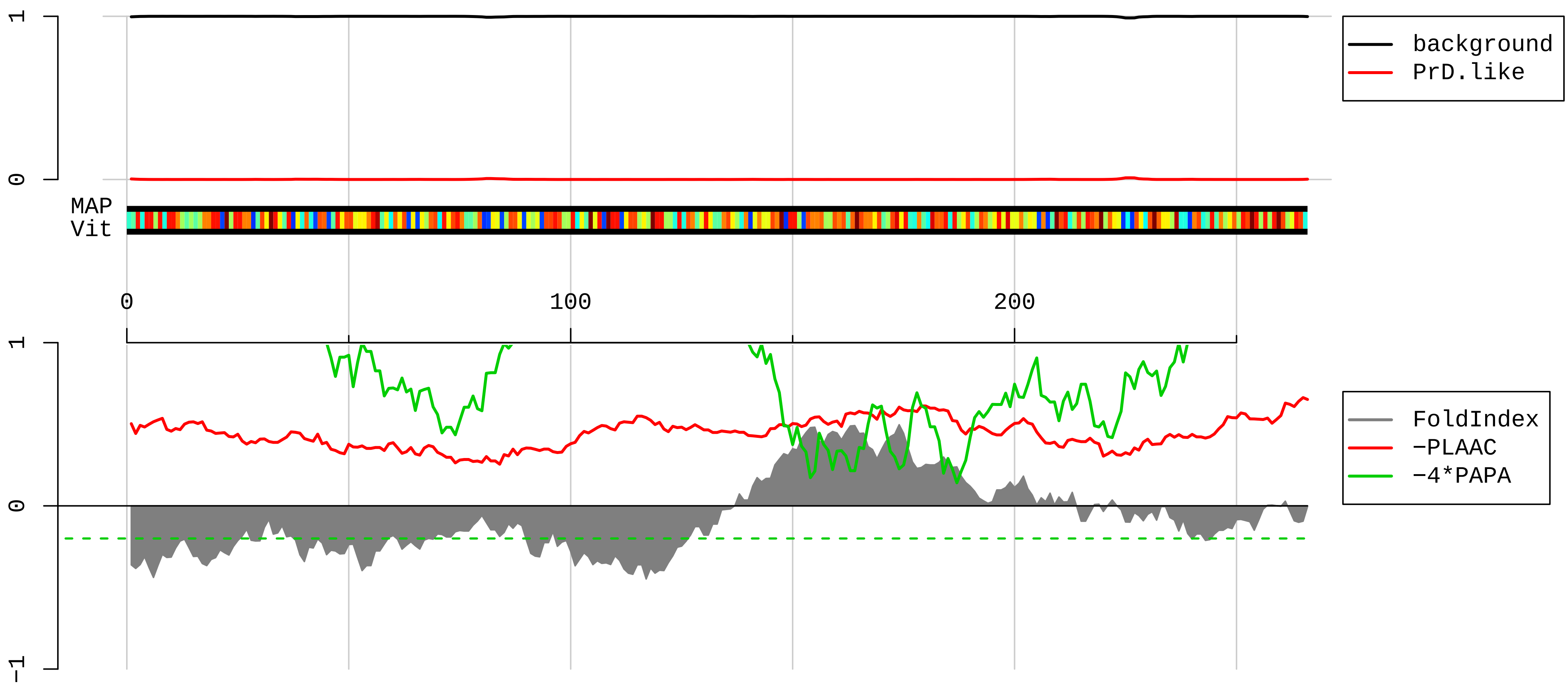


1 MPKGKKAKGKKVAPAPAVVKKQEAKKVVNPLFEKRPKNFGIGQDIQPKRDLTRFVKWPRYIRLQRQRAILYKRLKVPPAINQFTQALDRQTATQLLKLAH
101 KYRPETKQEKKQRLLARAEKKAAGKGDVPTKRPPVLRAGVNTVTTLVENKKAQLVVIAHDVDPIELVVFLPALCRKMGVPYCIIKGKARLGHLVHRKTCT
201 TVAFTQVNSEDKGALAKLVEAIRTNYNDRYDEIRRHWGGNVLGPKSVARIAKLEKAKAKELATKLG

**NP_443067.1 60S ribosomal protein L10 [Mus musculus]**

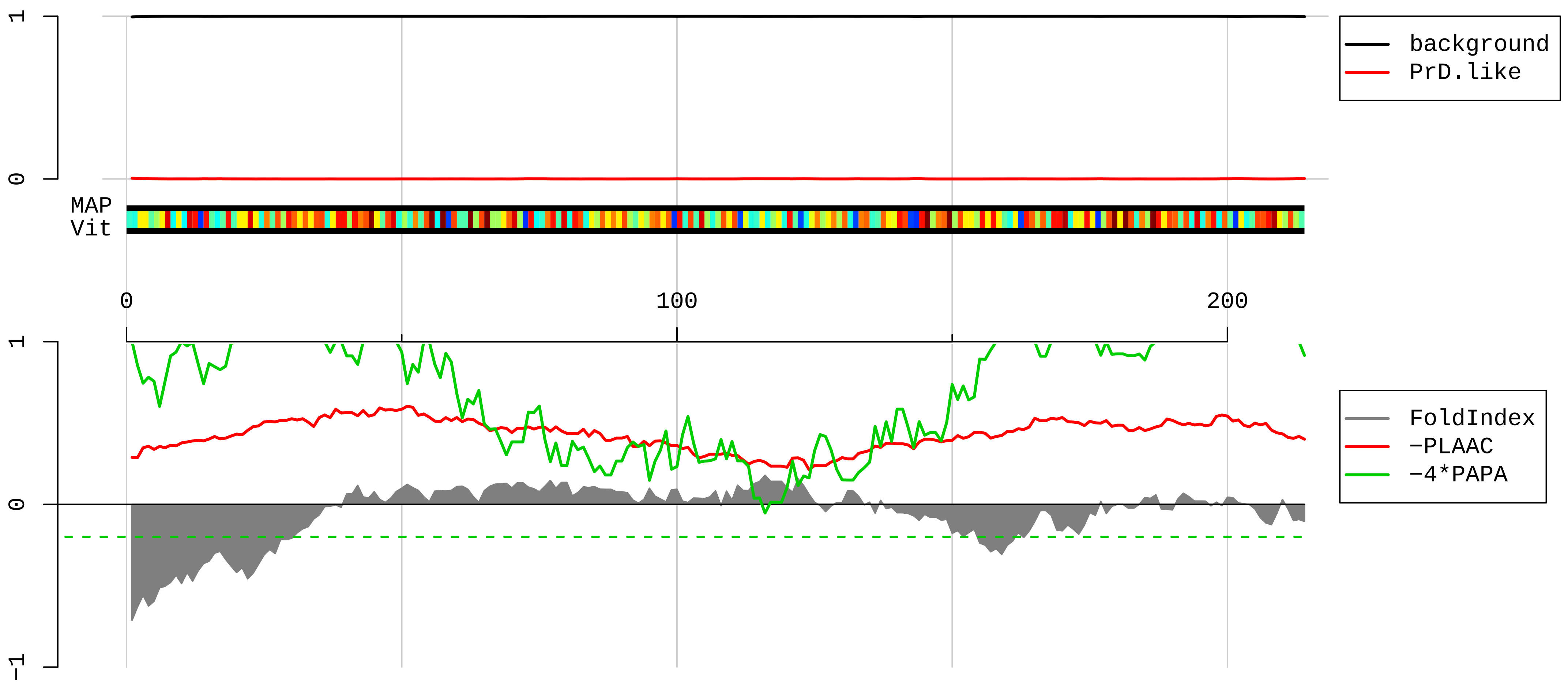


1 MGRRPARCYRYCKNKPYPKSRFCRGVPDAKIRIFDLGRKKAKVDEFPLCGHMVSDEYEQLSSEALEAARICANKYMVKSCGKDGFHIRVRLHPFHVIRIN
101 KMLSCAGADRLQTGMRGAFGKPQGTVARVHIGQVIMSIRTKLQNKEHVIEALRRAKFKFPGRQKIHISKKWGFTKFNADEFEDMVAEKRLIPDGCGVKYI
201 PNRGPLDKWRALHS

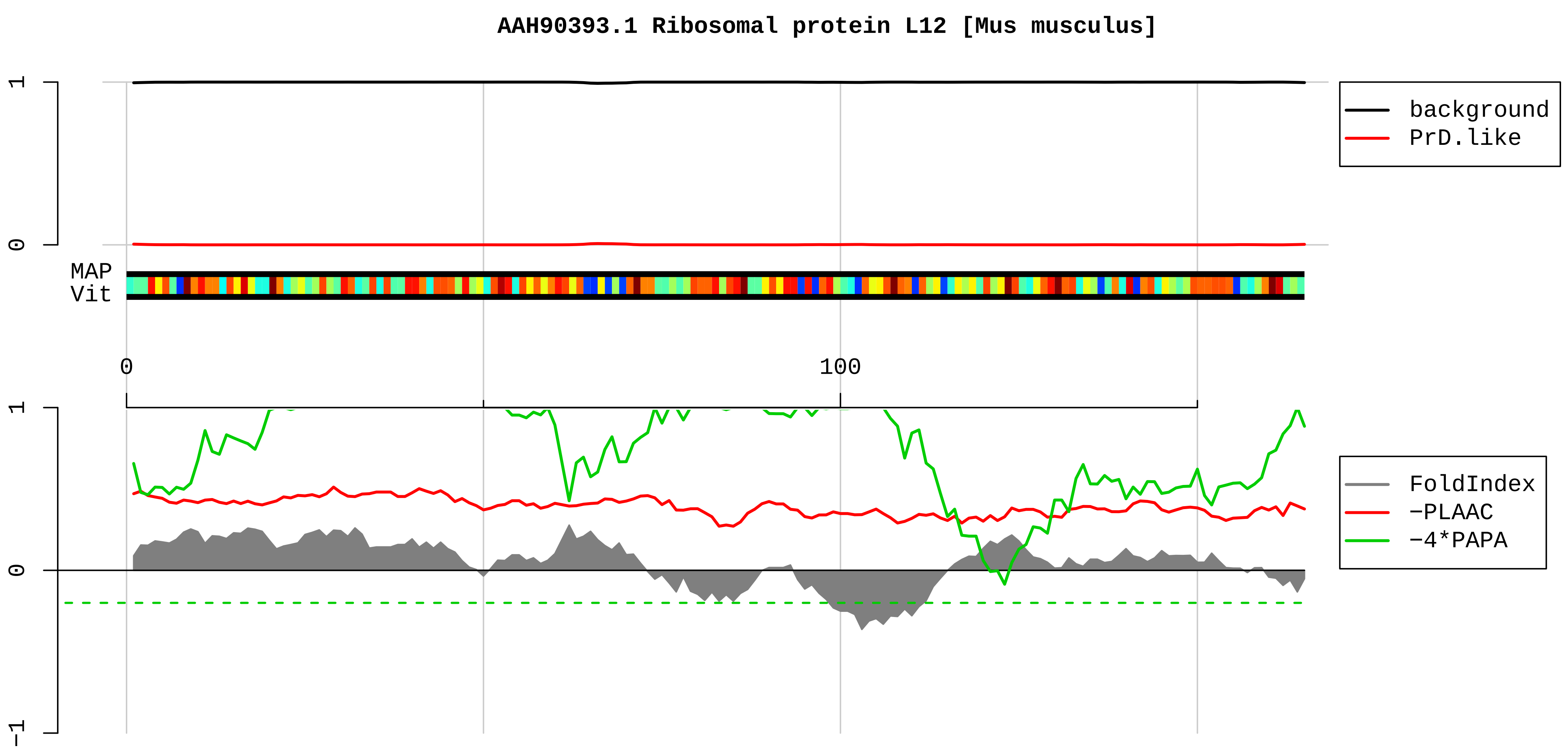


1 MPPKFDPNEVKVVYLRCTGGEVGATSALAPKIGPLGLSPKKVGDDIAKATGDWKGLRITVKLTIQNRQAQIEVVPSASALIIKALKEPPRDRKKQKNIKH
101 SGNITFDEIVNIARQMRHRSLARELSGTIKEILGTAQSVGCNVDGRHPHDIIDDINSGAVECPAS

**AAH98465.1 Myh10 protein, partial [Mus musculus]**

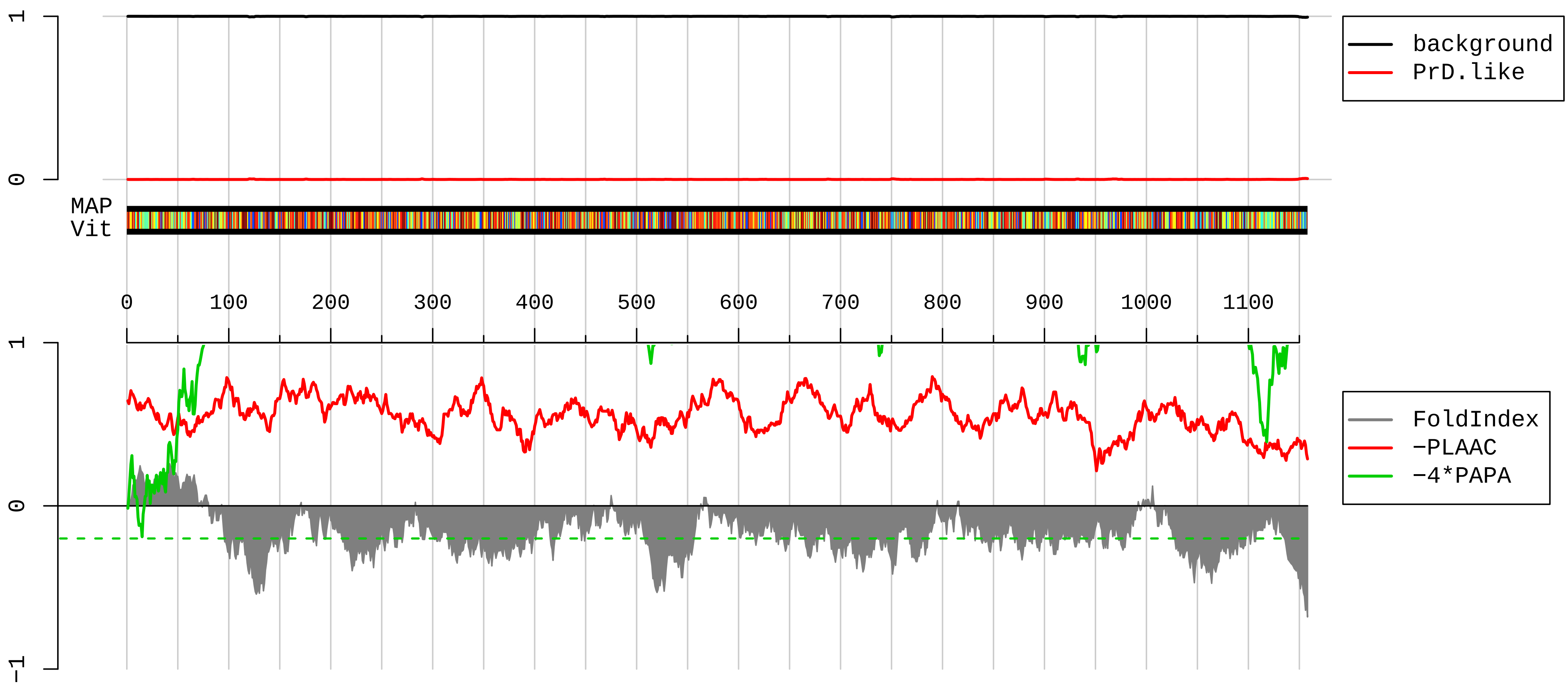


1 LELRRWICGRCWGRVPAPPSSAEEAITFGSFCLQWPRELDWRIPRGISLWTGLSSTTLPLKLTGQQLLEEKNILAEQLQAETELFAEAEEMRARLAAKKQ
101 ELEGILHDLESRVEEEEERNQILQNEKKKMQAHIQDLEEQLDEEEGARQKLQLEKVTAEAKIKKMEEEVLLLEDQNSKFIKEKKLMEDRIAECSSQLAEE
201 EEKAKNLAKIRNKQEVMISDLEERLKKEEKTRQELEKAKRKLDGETTDLQDQIAELQAQVDELKVQLTKKEEELQGALARGDDETLHKNNALKVARELQA
301 QIAELQEDFESEKASRNKAGKQKRDLSEELEALKTELEDTLDTTAAQQELRTKREQEVAELKKALEDETKNHEAQIQDMRQRHATALEELSEQLEQAKRF
401 KANLEKNKQGLETDNKELACEVKVLQQVKAESEHKRKKLDAQVQELHAKVSEGDRLRVELAEKANKLQNELDNVSTLLEEAEKKGIKFAKDAAGLESQLQ
501 DTQELLQEETRQKLNLSSRIRQLEEEKNSLQEQQEEEEARKNLEKQVLALQSQLADTKKKVDDDLGTIESLEEAKKKLLKDVEALSQRLEEKVLAYDKL
601 EKTKNRLQQELDDLTVDLDHQRQIVSNLEKKQKKFDQLLAEEKGISARYAEERDRAEAEAREKETKALSLARALEEALEAKEEFERQNKQLRADMEDLMS
701 SKDDVGKNVHELEKSKRALEQQVEEMRTQLEELEDELQATEDAKLRLEVNMQAMKAQFEGDLQTRDEQNEEKKRLLLKQVRELEAELEDERKQRALAVAS
801 KKKMEIDLKDLEAQIEAANKARDEVIKQLRKLQAQMKDYQRELEEARASRDEIFAQSKESEKKLKSLEAEILQLQEELASSERARRHAEQERDELADEIA
901 NSASGKSALLDEKRRLEARIAQLEEELEEEQSNMELLNDRFRKTTLQVDTLNTELAAERSAAQKSDNARQQLERQNKELKAKLQELEGAVKSKFKATISA
1001 LEAKIGQLEEQLEQEAKERAAANKLVRRTEKKLKEIFMQVEDERRHADQYKEQMEKANARMKQLKRQLEEAEEEATRANASRRKLQRELDDATEANEGLS
1101 REVSTLKNRLRRGGPISFSSSRSGRRQLHIEGASLELSDDDTESKTSDVNDTQPPQSE

**AAI06122.1 Ribosomal protein L6 [Mus musculus]**

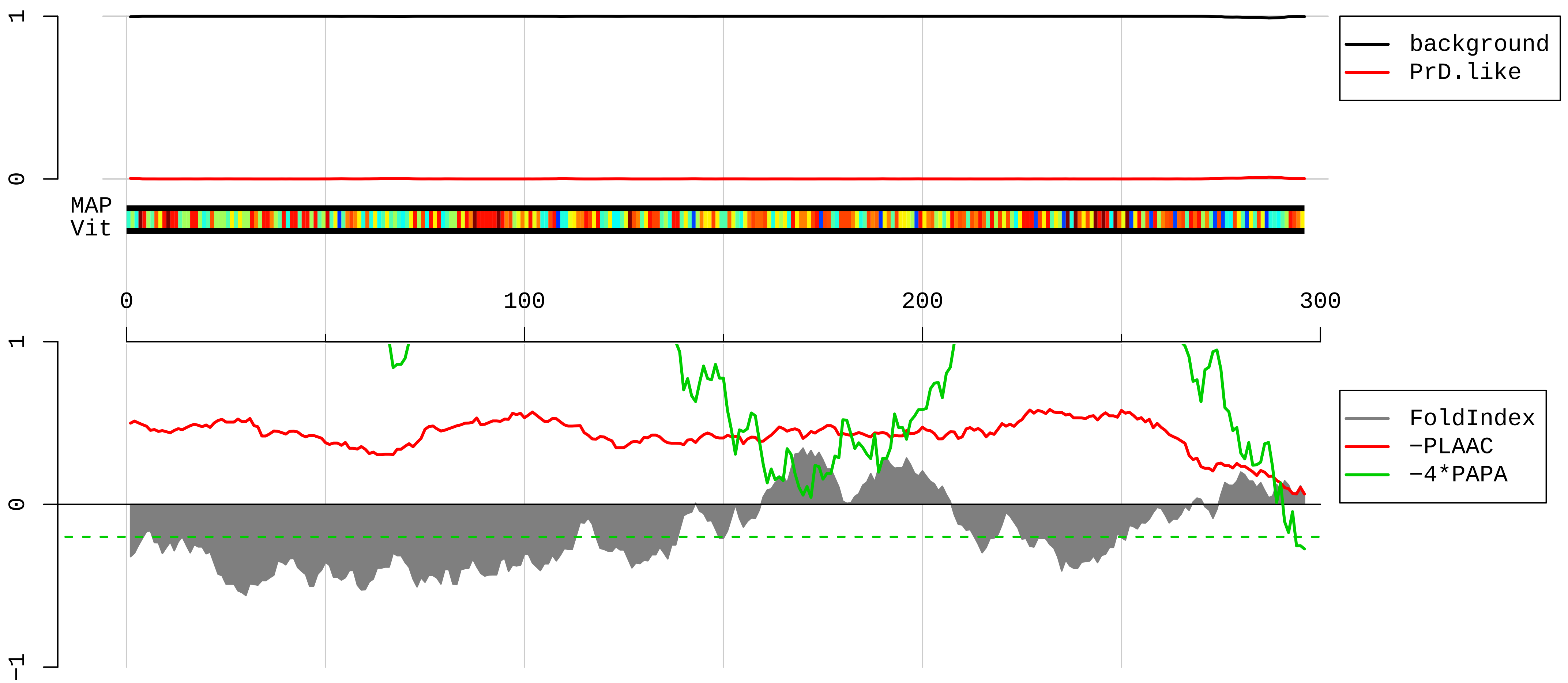


1 MAGEKAPDTKEKKPAAKKAGSDAAASRPRAAKVAKKVHPKGKKPKKAKPHCSRNPVLVRGIGRYSRSAMYSRKALYKRKYSAAKTKVEKKKKKEKVLATV
101 TKTVGGDKNGGTRVVKLRKMPRYYPTEDVPRKLLSHGKKPFSQHVRRLRSSITPGTVLIILTGRHRGKRVVFLKQLDSGLLLVTGPLVINRVPLRRTHQK
201 FVIATSTKVDISDVKIPKHLTDAYFKKKQLRKPRHQEGEIFDTEKEKYEITEQRKADQKAVDLQILPKIKAVPQLQGYLRSQFSLTNGMYPHKLVF

**AAH10721.1 Ribosomal protein S3 [Mus musculus]**

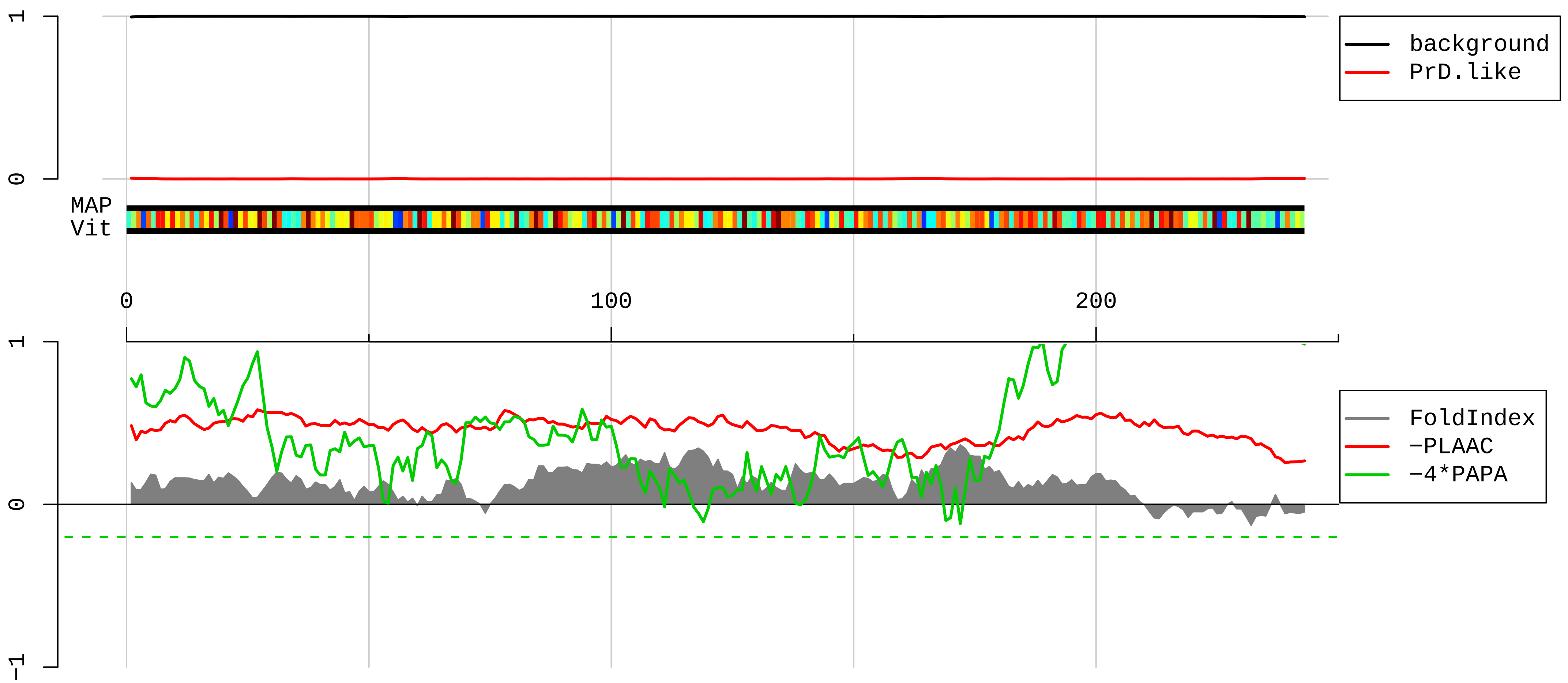


1 MAVQISKKRKFVADGIFKAELNEFLTRELAEDGYSGVEVRVTPTRTEIIILATRTQNVLGEKGRRIRELTAVVQKRFGFPEGSVELYAEKVATRGLCAIA
101 QAESLRYKLLGGLAVRRACYGVLRFIMESGAKGCEVVVSGKLRGQRAKSMKFVDGLMIHSGDPVNYYVDTAVRHVLLRQGVLGIKVKIMLPWDPSGKIGP
201 KKPLPDHVSIVEPKDEILPTTPISEQKGGKPEPPAMPQPVPTA

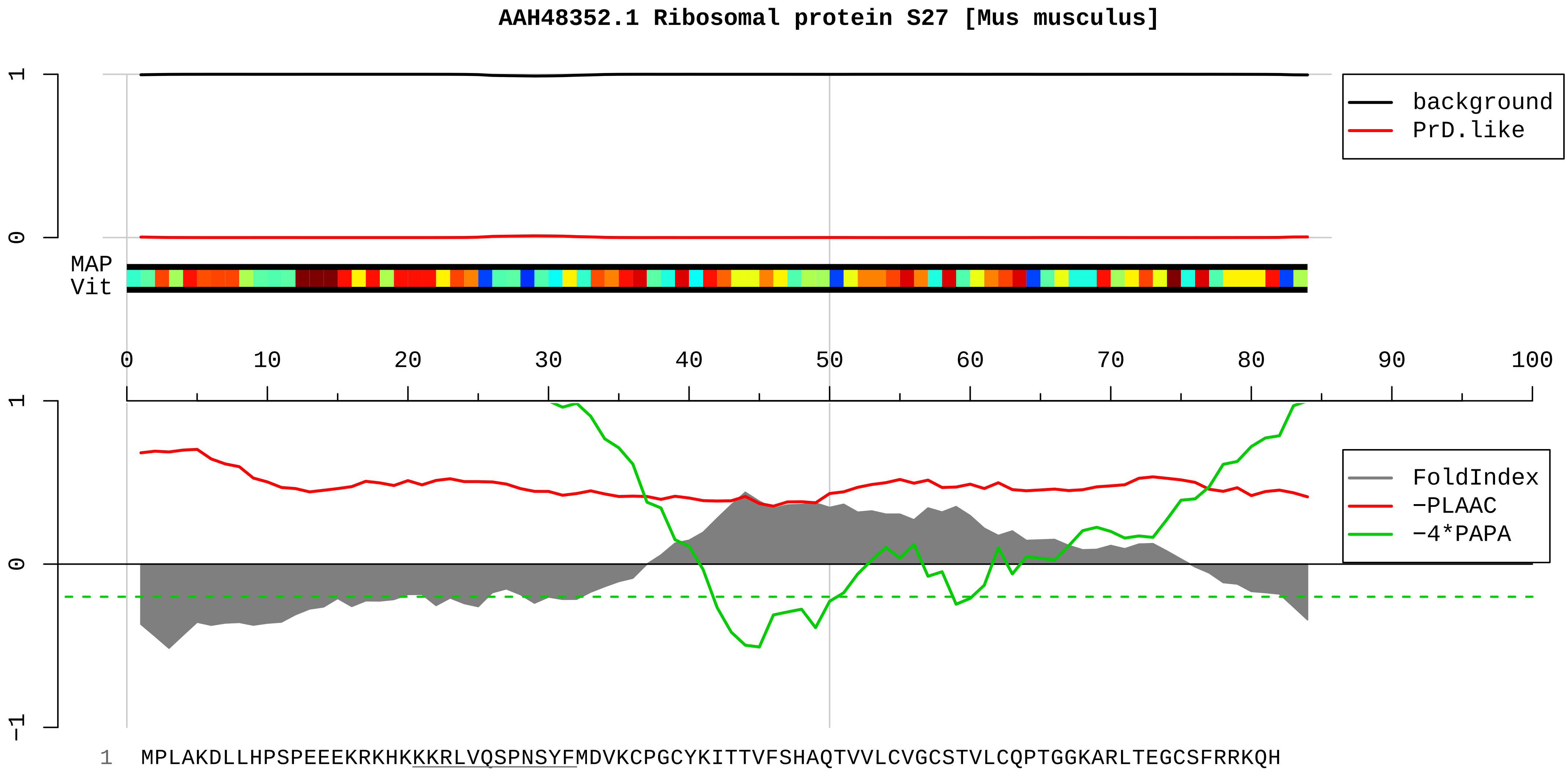

AAH48352.1 Ribosomal protein S27 [Mus musculus]
background
PrD.like
MAP
Vit
FoldIndex
−PLAAC
−4*PAPA
1 MPLAKDLLHPSPEEEKRKHKKKRLVQSPNSYFMDVKCPGCYKITTVFSHAQTVVLCVGCSTVLCQPTGGKARLTEGCSFRRKQH

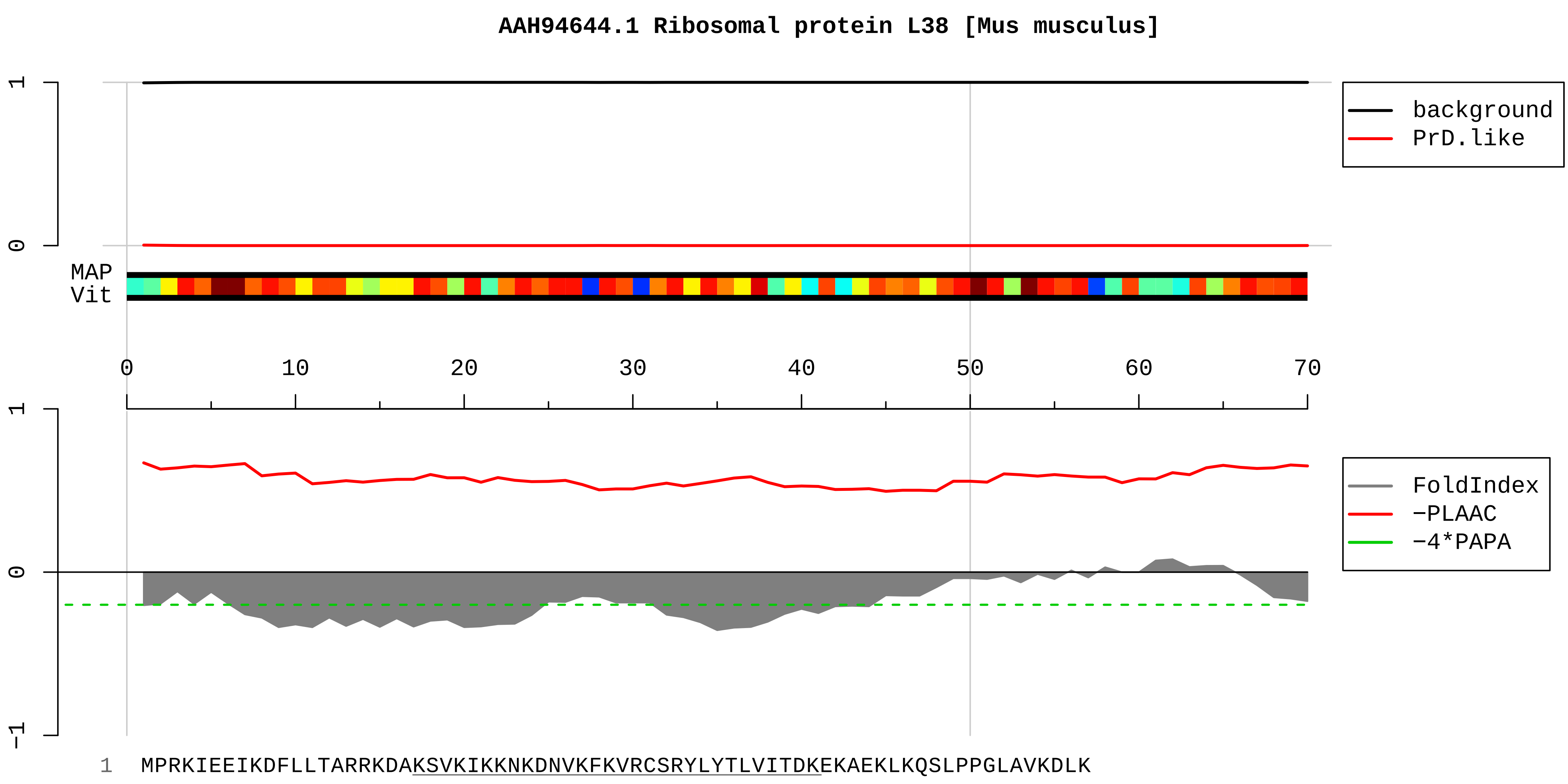
AAH94644.1 Ribosomal protein L38 [Mus musculus]
background
PrD.like
MAP
Vit
FoldIndex
−PLAAC
−4*PAPA
1 MPRKIEEIKDFLLTARRKDAKSVKIKKNKDNVKFKVRCSRYLYTLVITDKEKAEKLKQSLPPGLAVKDLK

**AAO23055.1 interleukin enhancer binding factor 3 [Mus musculus]**

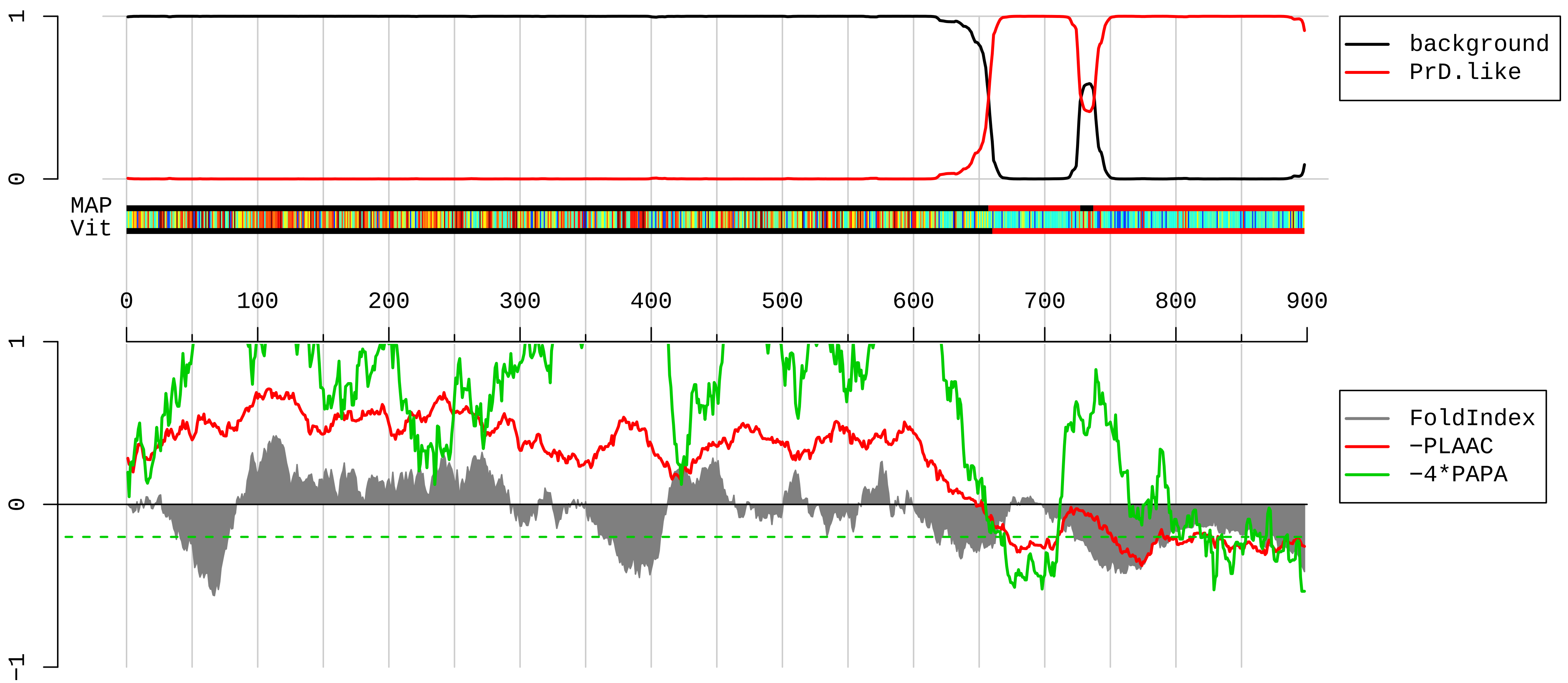


```
1   MRPMRIFVNDDRHVMAKHSSVYPTQEELEAVQNMVSHTERALKAVSDWIDEQEKGNSELSEAENMDTPPDDESKEGAGEQKAEHMTRTLRGVMRVGLVAK
101 GLLLKGDLDLELVLLCKEKPTTALLDKVADNLAIQLTTVTEDKYEILQSVDDAAIVIKNTKEPPLSLTIHLTSPVVREEMEKVLAGETLSVNDPPDVLDR
201 QKCLAALASLRHAKWFQARANGLKSCVIVIRVLRDLCTRVPTWGPLRGWPLELLCEKSIGTANRPMGAGEALRRVLECLASGIVMPDGSGIYDPCEKEAT
301 DAIGHLDRQQREDITQSAQHALRLAAFGQLHKVLGMDPLPSKMPKKPKNENPVDYTVQIPPSTTYAITPMKRPMEEDGEEKSPSKKKKKIQKKEEKADPP
401 QAMNALMRLNQLKPGLQYKLISQTGPVHAPIFTMSVEVDGSNFEASGPSKKTAKLHVAVKVLQDMGLPTGAEGRDSSKGEDSAEESDGKPAIVAPPPVVE
501 AVSNPSSVFPSDATTEQGPILTKHGKNPVMELNEKRRGLKYELISETGGSHDKRFVMEVEVDGQKFQGAGSNKKVAKAYAALAALEKLFPDTPLALEANK
601 KKRTPVPVRGGPKFAAKPHNPGFGMGGPMHNEVPPPPNIRGRGRGGNIRGRGRGRGFGGANHGGGYMNAGAGYGSYGYSSNSATAGYSQFYSNGGHSGNA
701 GGGGSGGGGGSSSYSSYYQGDSYNSPVPPKHAGKKPLHGGQQKASYSSGYQSHQGQQQPYNQSQYSSYGTPQGKQKGYGHGQGSYSSYSNSYNSPGGGGG
801 SDYSYDSKFNYSGSGGRSGGNSYGSSGSSSYNTGSHGGYGTGSGGSSSYQGKQGGYSSQSNYSSPGSSQSYSGPASSYQSSQGGYSRNTEHSMNYQYR
```

**AAH04679.1 Hnrpr protein, partial [Mus musculus]**

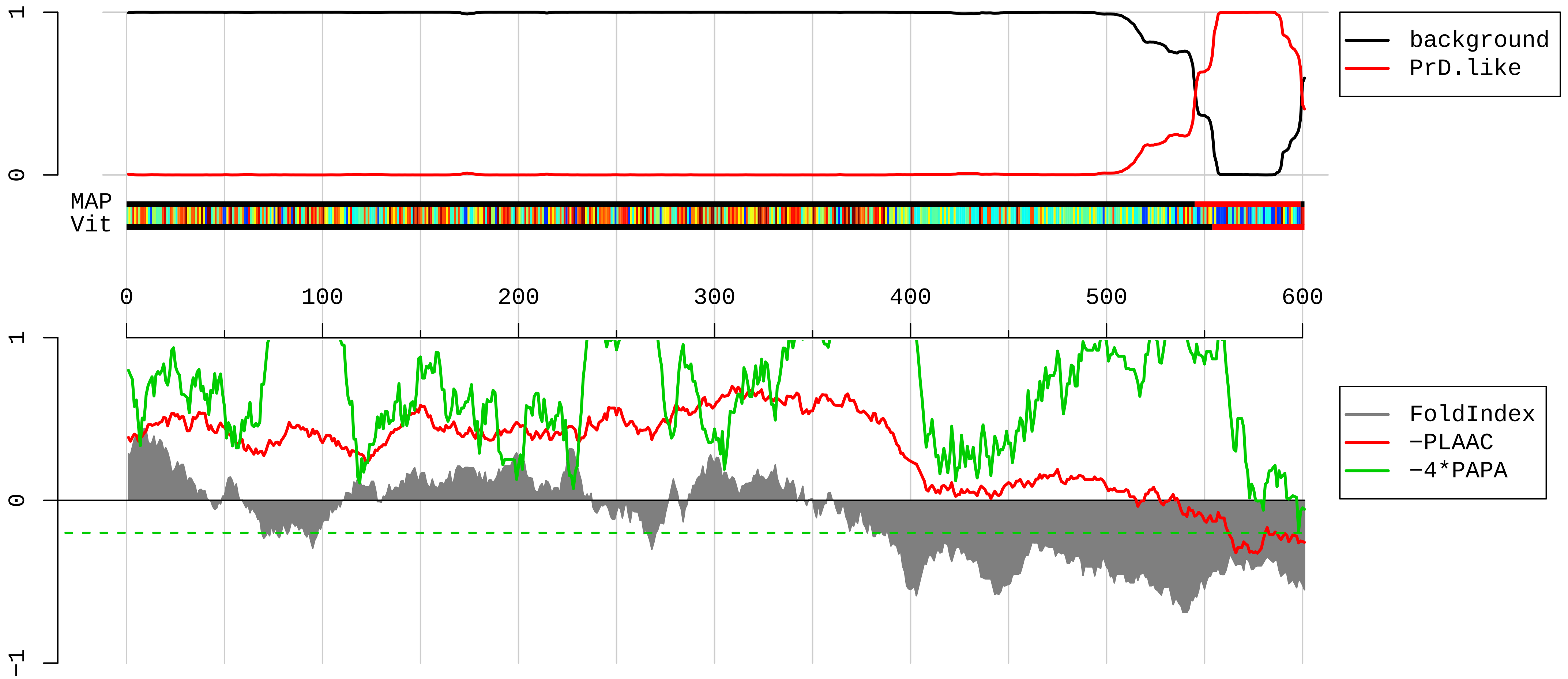


1 SFSKRLALLLTGQAASPPLCPGLVAYVDLDERAIDALREFNEEGALSVLQQFKESDLSHVQNKSAFLCGVMKTYRQREKQGSKVQESTKGPDEAKIKALL
101 ERTGYTLDVTTGQRKYGGPPPDSVYSGVQPGIGTEVFVGKIPRDLYEDELVPLFEKAGPIWDLRLMMDPLSGQNRGYAFITFCGKEAAQEAVKLCDSYEI
201 RPGKHLGVCISVANNRLFVGSIPKNKTKENILEEFSKVTEGLVDVILYHQPDDKKKNRGFCFLEYEDHKSAAQARRRLMSGKVKVWGNVVTVEWADPVEE
301 PDPEVMAKVKVLFVRNLATTVTEEILEKSFSEFGKLERVKKLKDYAFVHFEDRGAAVKAMDEMNGKEIEGEEIEIVLAKPPDKKRKERQAARQASRSTAY
401 EDYYYHPPPRMPPPMRGRGRGGRGGYGYPPDYYGYEDYYDDYYGYDYHDYRGGYEDPYYGYDDGYAVRGRGGGRGGRGAPPPPRGRGAPPPRGRAGYSQR
501 GAPLGPPRGSRGGRGGPAQQQRGRGSRGARGNRGGNVGGKRKADGYNQPDSKRRQTNNQQNWGSQPIAQQPLQQGGDYSGNYGYNNDNQEFYQDTYGQQW
601 K

**AAH03283.1 Poly(A) binding protein, cytoplasmic 4 [Mus musculus]**

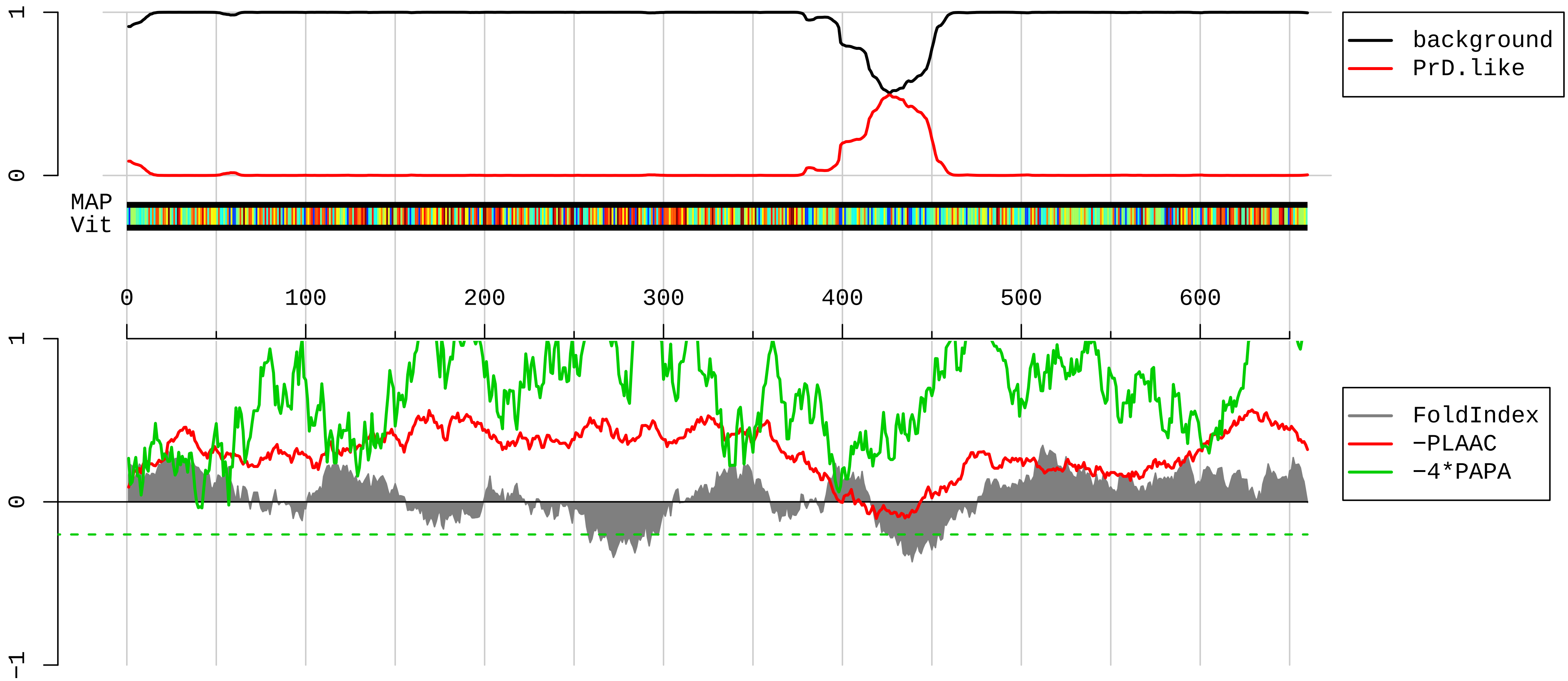


1 MNAAASSYPMASLYVGDLHSDVTEAMLYEKFSPAGPVLSIRVCRDMITRRSLGYAYVNFQQPADAERALDTMNFDVMKGKPIRIMWSQRDPSLRKSGVGN
101 VFIKNLDKSIDNKALYDTFSAFGNILSCKVVCDENGSKGYAFVHFETQEAADKAIEKMNGMLLNDRKVFVGRFKSRKEREAELGAKAKEFTNVYIKNFGE
201 EVDDGNLKELFSQFGKTLSVKVMRDSSGKSKGFGFVSYEKHEDANKAVEEMNGKEMSGKAIFVGRAQKKVERQAELKRKFEQLKQERISRYQGVNLYIKN
301 LDDTIDDEKLRKEFSPFGSITSAKVMLEDGRSKGFGFVCFSSPEEATKAVTEMNGRIVGSKPLYVALAQRKEERKAHLTNQYMQRVAGMRALPASAILNQ
401 FQPAAGGYFVPAVPQAQGRPPYYTPNQLAQMRPNPRWQQGGRPQGFQGMPSALRQSGPRPALRHLAPTGNAPASRGLPTTAQRVGSECPDRLAMDFGGAG
501 AAQQGLTDSCQSGGVPTAVPNLAPRAAVAAAAPRAVAPYKYASSVRSPHPAIQPLQAPQPAVHVQGQEPLTASMLAAAPPQEQKQMLGERLFPLIQTMHS
601 NLAGKITGMLLEIDNSELLHMLESPESLRSKVDEAVAVLQAHHAKKEAAQKVGTVAAATS

**AAI08364.2 Syncrip protein [Mus musculus]**

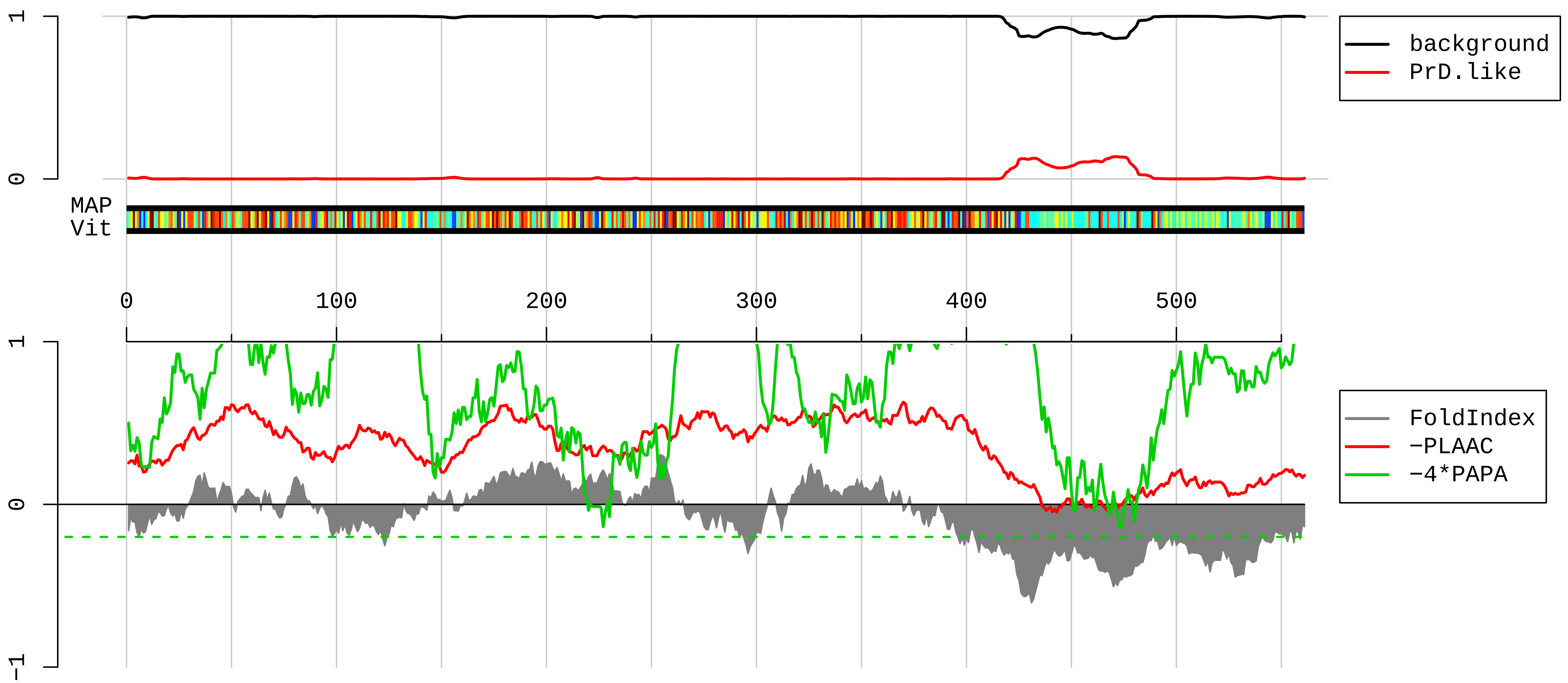


1 MATEHVNGNGTEEPMDTTSAVIHSENFQTLLDAGLPQKVAEKLDEIYVAGLVAHSDLDERAIEALKEFNEDGALAVLQQFKDSDLSHVQNKSAFLCGVMK
101 TYRQREKQGTKVADSSKGPDEAKIKALLERTGYTLDVTTGQRKYGGPPPDSVYSGQQPSVGTEIFVGKIPRDLFEDELVPLFEKAGPIWDLRLMMDPLTG
201 LNRGYAFVTFCTKEAAQEAVKLYNNHEIRSGKHIGVCISVANNRLFVGSIPKSKTKEQILEEFSKVTEGLTDVILYHQPDDKKKNRGFCFLEYEDHKTAA
301 QARRRLMSGKVKVWGNVGTVEWADPIEDPDPEVMAKVKVLFVRNLANTVTEEILEKSFSQFGKLERVKKLKDYAFIHFDERDGAVKAMEEMNGKDLEGEN
401 IEIVFAKPPDQKRKERKAQRQAAKNQMYDDYYYYGPPHMPPPTRGRGRGGRGGYGYPPDYYGYEDYYDYYGYDYHNYRGGYEDPYYGYEDFQVGARGRGG
501 RGARGAAPSRGRGAAPPRGRAGYSQRGGPGSARGVRGARGGAQQQRGRGGKGVEAGPDLLQ

**NP_035831.2 vimentin [Mus musculus]**

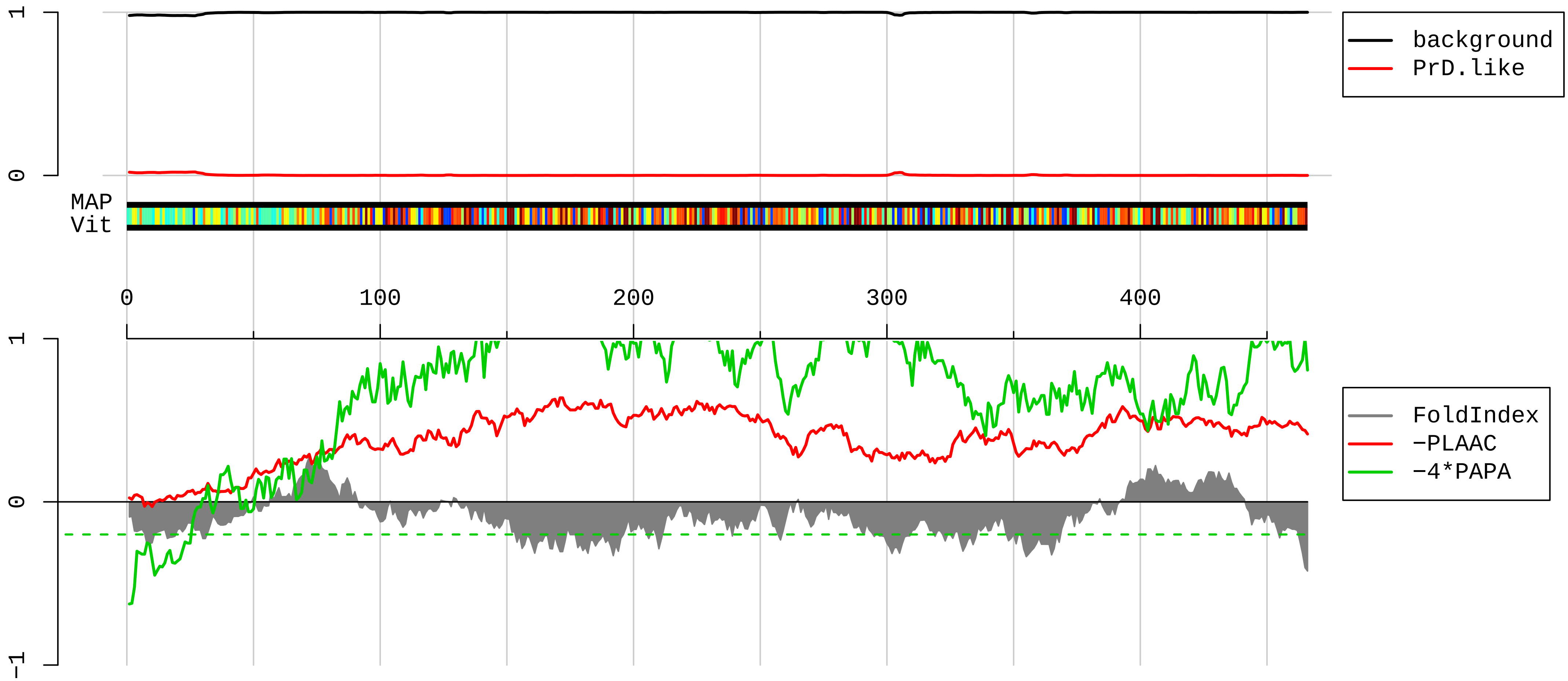


```
  1  MSTRSVSSSSYRRMFGGSGTSSRPSSNRSYVTTSTRTYSLGSALRPSTSRSLYSSSPGGAYVTRSSAVRLRSSVPGVRLLQDSVDFSLADAINTEFKNTR
101  TNEKVELQELNDRFANYIDKVRFLEQQNKILLAELEQLKGQGKSRLGDLYEEEMRELRRQVDQLTNDKARVEVERDNLAEDIMRLREKLQEEMLQREEAE
201  STLQSFRQDVDNASLARLDLERKVESLQEEIAFLKKLHDEEIQELQAQIQEQHVQIDVDVSKPDLTAALRDVRQQYESVAAKNLQEAEEWYKSKFADLSE
301  AANRNNDALRQAKQESNEYRRQVQSLTCEVDALKGTNESLERQMREMEENFALEAANYQDTIGRLQDEIQNMKEEMARHLREYQDLLNVKMALDIEIATY
401  RKLLEGEESRISLPLPTFSSLNLRETNLESLPLVDTHSKRTLLIKTVETRDGQVINETSQHHDDLE
```

**FMR1_MOUSE RecName: Full=Synaptic functional regulator FMR1; AltName: Full=Fragile X mental retardation protein 1 homolog; Short=FMRP; Short=Protein FMR−1; Short=mFmr1p−**

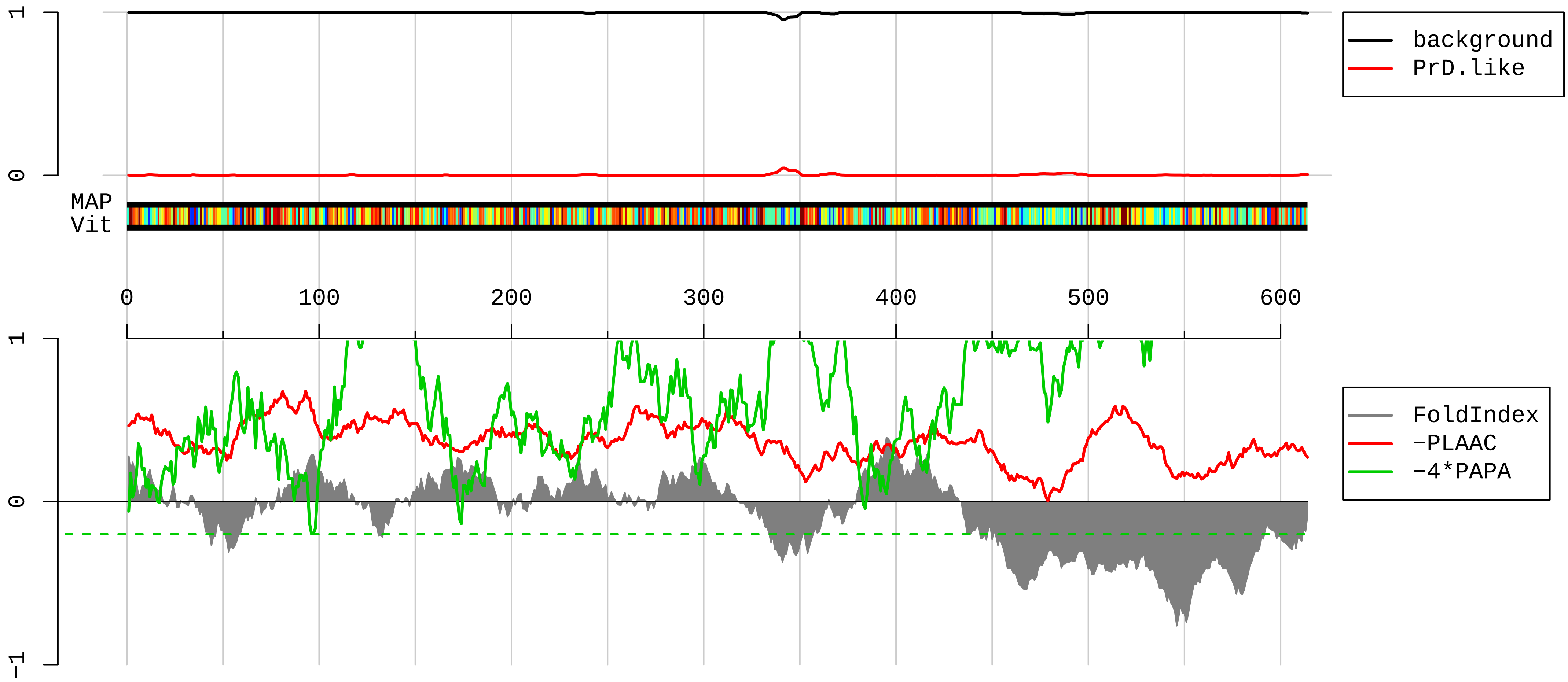


1 MEELVVEVRGSNGAFYKAFVKDVHEDSITVAFENNWQPERQIPFHDVRFPPPVGYNKDINESDEVEVYSRANEKEPCCWWLAKVRMIKGEFYVIEYAACD
101 ATYNEIVTIERLRSVNPNKPATKDTFHKIKLEVPEDLRQMCAKESAHKDFKKAVGAFSVTYDPENYQLVILSINEVTSKRAHMLIDMHFRSLRTKLSLIL
201 RNEEASKQLESSRQLASRFHEQFIVREDLMGLAIGTHGANIQQARKVPGVTAIDLDEDTCTFHIYGEDQDAVKKARSFLEFAEDVIQVPRNLVGKVIGKN
301 GKLIQEIVDKSGVVRVRIEAENEKSVPQEEEIMPPSSLPSNNSRVGPNSSEEKKHLDTKENTHFSQPNSTKVQRVLVVSSIVAGGPQKPEPKAWQGMVPF
401 VFVGTKDSIANATVLLDYHLNYLKEVDQLRLERLQIDEQLRQIGASSRPPPNRTDKEKGYVTDDGQGMGRGSRPYRNRGHGRRGPGYTSGTNSEASNASE
501 TESDHRDELSDWSLAPTEEERESFLRRGDGRRRRGGGRGQGGRGRGGGFKGNDDHSRTDNRPRNPREAKGRTADGSLQSASSEGSRLRTGKDRNQKKEKP
601 DSVDGLQPLVNGVP

**AAI72151.1 Scaper protein [Mus musculus]**

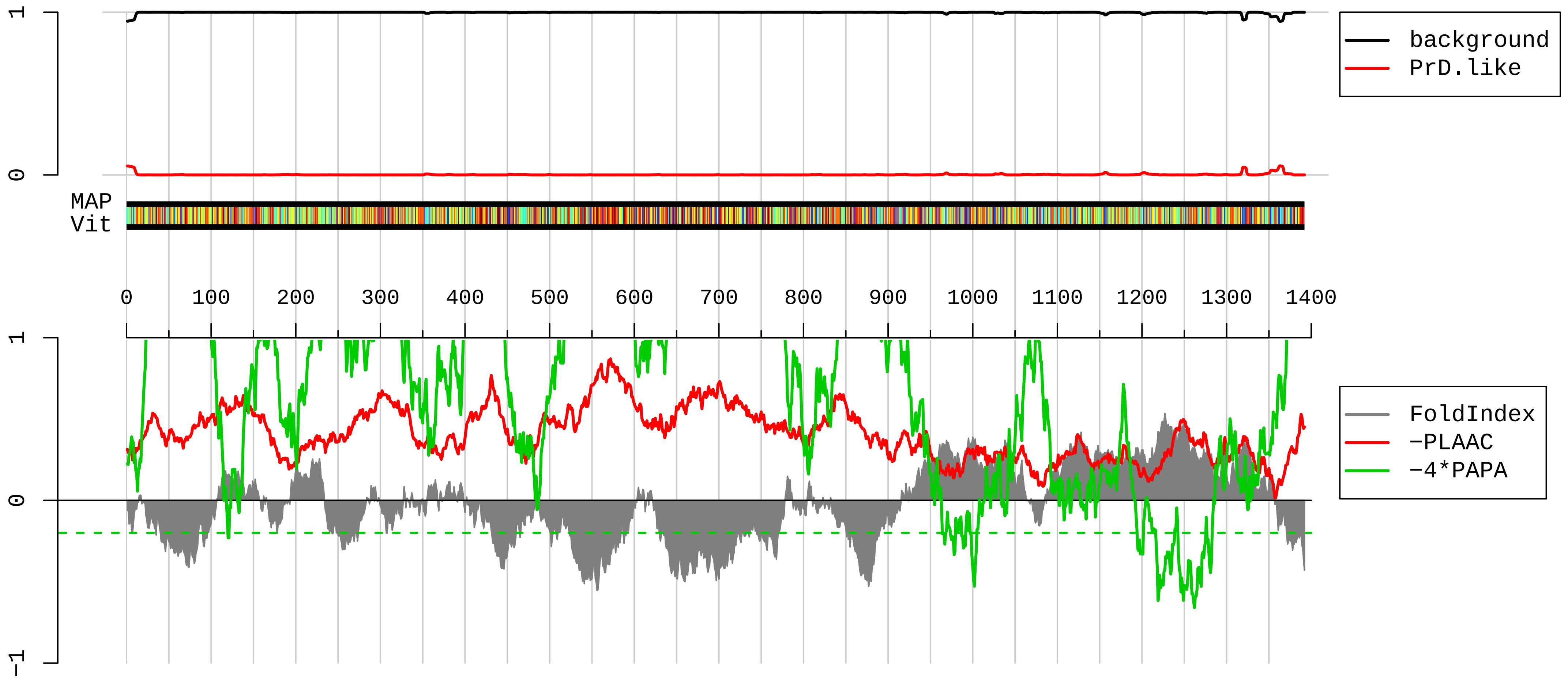


```
   1  MMASFQRSNSHDKVRRIVAEEGRTARNLIAWSVPLESKDDDGKPKCQTGGKSKRTIQGTHKTTKQNTAVECKITSTTGEKHFDKSPTKTRHPRKIDLRAR
 101  YWAFLFDNLRRAVDEIYVTCESDQSVVECKEVLMMLDYYVRDFKALIDWIQLQEKLEKTDAQSRPTSLAWEVKKMSPGRHVIQSPSTDRMNVTSTARRSL
 201  NFGSLPGAVTAPCLAPTGVSWADKVKAHHTGSAPPEATPAQSCAPMTVQKTSRKNERKDAEGWETVQRGRPVRSRSTAVIPKVSLVTETLRSKDDSDKEN
 301  ICLLPEESIQKGKLEDRCSTVESLSKDSLLHSSDHPLSERTQFTVCILDDVKNSGSSQDSSVHTSEIPAIDSDAVCVAPQNTTALPTDRVPAETTGMVTE
 401  MTDPSDISNSMAEVLAKKEELADRLEKANEEAIASAIAEEEQLTREIEAEENNDINIETDNDSDFSASMGSGSLSFCGVSLDWNDVLADYEARESWRQNT
 501  SWGDIVEEEPARLPGHGIHMHEKLSSPSRKRTIAESKKKYEEKHMKAQQLREKLREEKTLKLQKLLEREKDVRKWKEELLDQRRRMMEEKLLHAEFKREV
 601  QLQAIVKKAQEEEAKVNEIAFINTLEAQNKRHDVLSKLKEYEQRLNELQEERQRRQEEKQARDEAVQERKRALEAERQARVEELLMKRKEQEARIEQQRQ
 701  EKEKAREDAARERARDREERLAALTAAQQEAMEELQKKIQLKHDESIRRHMEQIEQRKEKAAELSSGRHASTDYAPKLTPYERKKQCSLCNVLIASEVYL
 801  FSHIKGKKHQQAVRENSSIQGRELSDEEVEHLSLKKYVVDIVIESAAPPEPVKDGEERQKNKKKAKKIKARMNIRAKEYENLVETKNSGSESPYKAKLQR
 901  LTKDLVKQLQVQDSGSWVNNKASALDRTLGEIARILEKENVPDQISFQVAGGLTALENILQVVTPATNMNTVSRIPPKSLCNAINVYSLACNNCSENCTD
1001  VLFSNKITFLMDLLIHQLTVYVPDENNAVLGRNTNKQVFEGLTTGLLRAGAVVFSCLIANRPDGNSRPATPKISTQEMKNKPSQGDAFNSRVQDLISYVV
1101  NMGLIDRLHGCFLSVQGPVDENPKMATFLQHAAGFLRGMCMLCFAVTGRSSSIFDNNHQDPTGLTAALQATDLAGVLHMLYCILFHGTISDAGTSPKDSY
1201  TQNTIQVAIQSLRFFNSFAILDLSAFQSVVGAEGLSLAFRHMASSLLGHCSQVSCESLLHEVIVCVGYFTVNHPDNQVIVQSGRHPTVLQKLCQLPFQYF
1301  SDPRLIKVLFPSLITACHNNHQNKLILEQEMSCVLLATFIQDFAQTPGHVDNQSSSLKGKFQNPGNYLELANRFPQQAWEEARQFFLKKEKK
```

**NP_038744.1 ras GTPase−activating protein−binding protein 1 [Mus musculus]**

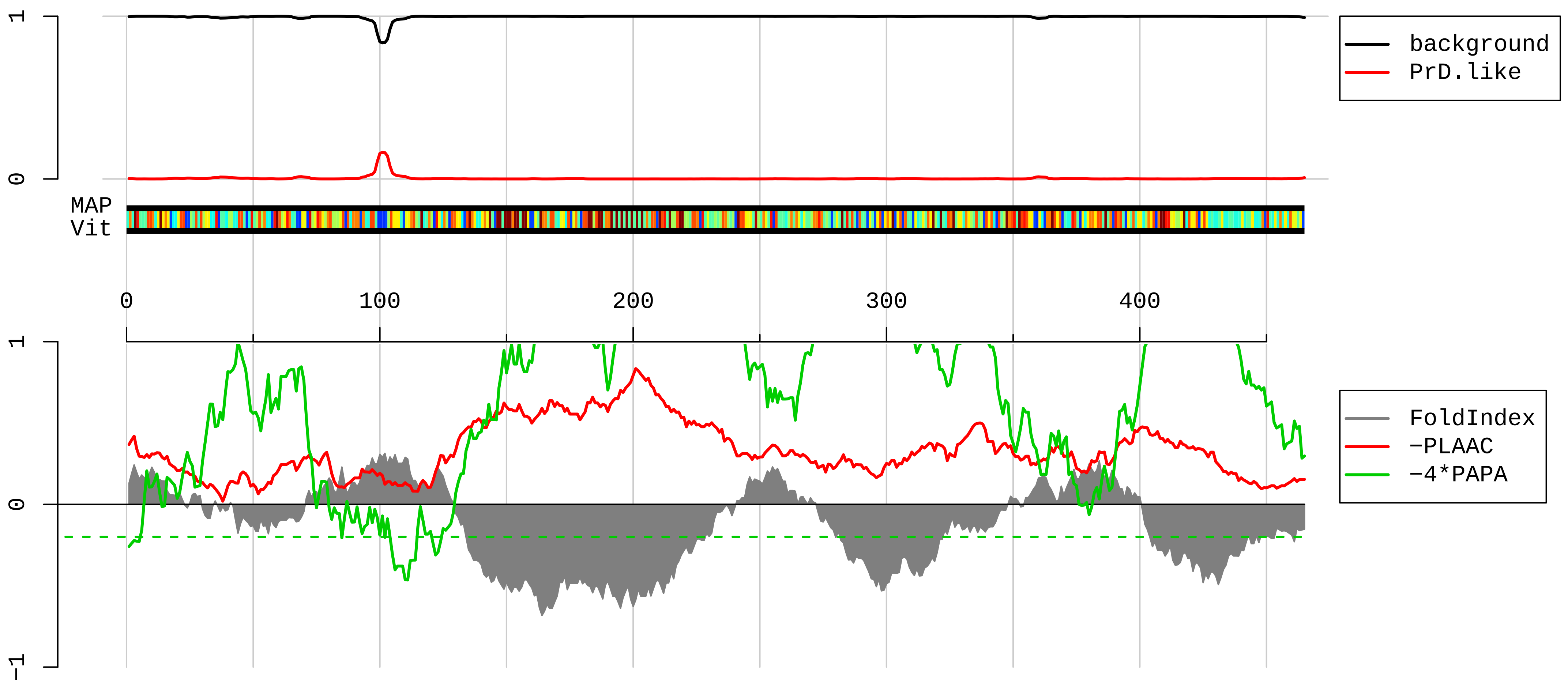


1 MVMEKPSPLLVGREFVRQYYTLLNQAPDMLHRFYGKNSSYAHGGLDSNGKPADAVYGQKEIHRKVMSQNFTNCHTKIRHVDAHATLNDGVVVQVMGLLSN
101 NNQALRRFMQTFVLAPEGSVANKFYVHNDIFRYQDEVFGGFVTEPQEESEEEVEEPEERQQTPEVVPDDSGTFYDQTVSNDLEEHLEEPVVEPEPEPEPE
201 PEPEPVSDIQEDKPEAALEEAAPDDVQKSTSPAPADVAPAQEDLRTFSWASVTSKNLPPSGAVPVTGTPPHVVKVPASQPRPESKPDSQIPPQRPQRDQR
301 VREQRINIPPQRGPRPIREAGEPGDVEPRRMVRHPDSHQLFIGNLPHEVDKSELKDFFQNFGNVVELRINSGGKLPNFGFVVFDDSEPVQKVLSNRPIMF
401 RGAVRLNVEEKKTRAAREGDRRDNRLRGPGGPRGGPSGGMRGPPRGGMVQKPGFGVGRGITTPRQ

#### NP_034617.1 ELAV-like protein 3 [Mus musculus]

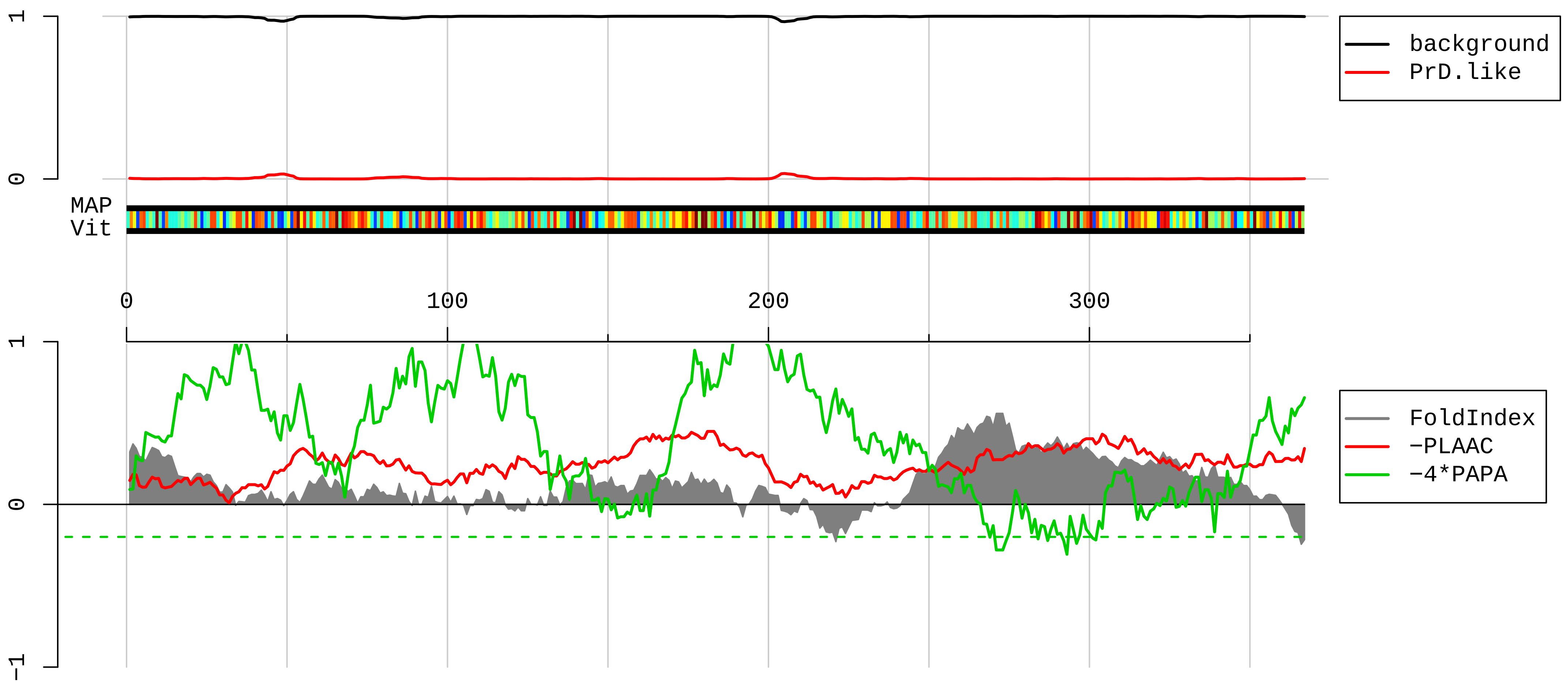


```
  1  MVTQILGAMESQVGGGPAGPALPNGPLLGTNGATDDSKTNLIVNYLPQNMTQDEFKSLFGSIGDIESCKLVRDKITGQSLGYGFVNYSDPNDADKAINTL
101  NGLKLQTKTIKVSYARPSSASIRDANLYVSGLPKTMSQKEMEQLFSQYGRIITSRILLDQATGVSRGVGFIRFDKRIEAEEAIKGLNGQKPLGAAEPITV
201  KFANNPSQKTGQALLTHLYQSSARRYAGPLHHQTQRFRLDNLLNMAYGVKSPLSLIARFSPIAIDGMSGLAGVGLSGGAAGAGWCIFVYNLSPEADESVL
301  WQLFGPFGAVTNVKVIRDFTTNKCKGFGFVTMTNYDEAAMAIASLNGYRLGERVLQVSFKTSKQHKA
```

**AAH81430.1 Ribosomal protein L27a [Mus musculus]**

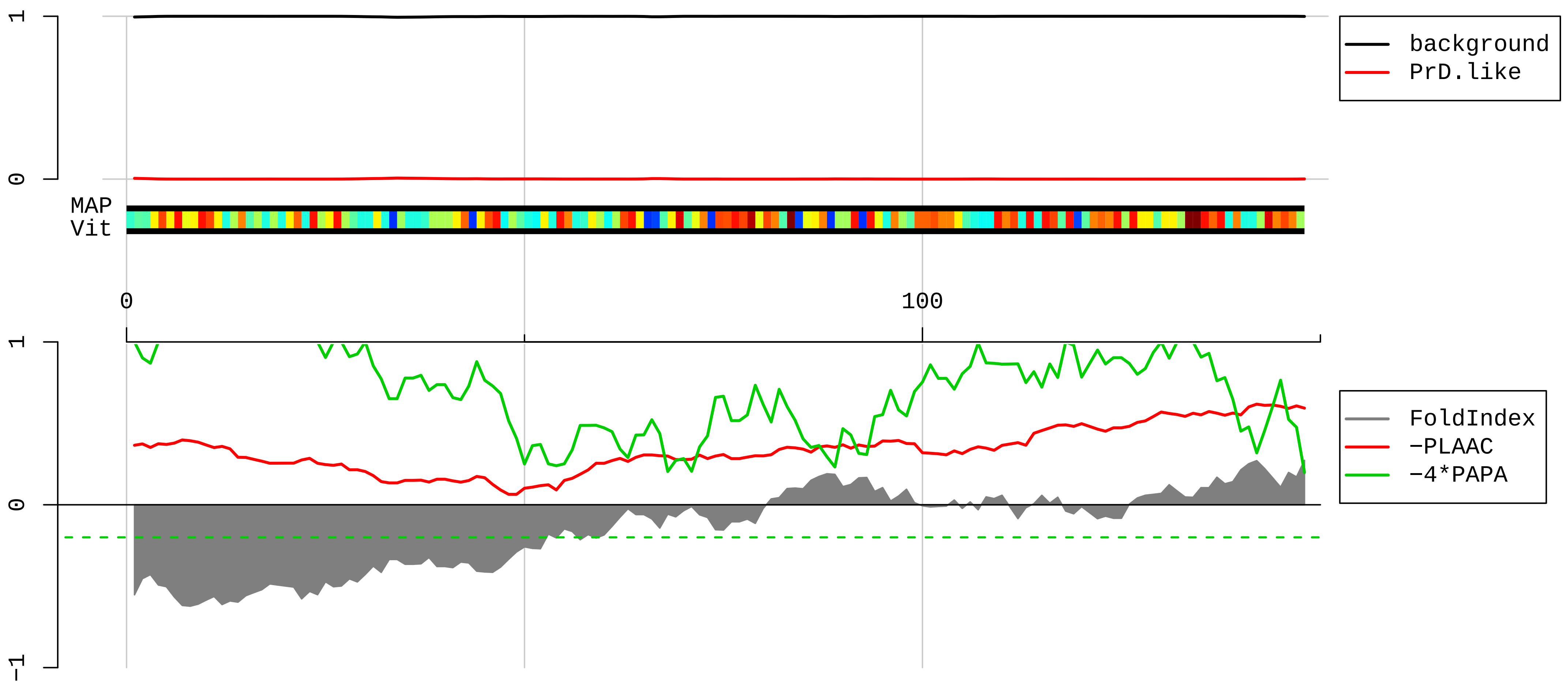


1 MPSRLRKTRKLRGHVSHGHGRIGKHRKHPGGRGNAGGMHHHRINFDKYHPGYFGKVGMRHYHLKRNQSFCPTVNLDKLWTLVSEQTRVNAAKNKTGVAPI

101 IDVVRSGYYKVLGKGKLPKQPVIVKAKFFSRRAEEKIKGVGGACVLVA

**CAA62383.1 FXR1 [Mus musculus]**

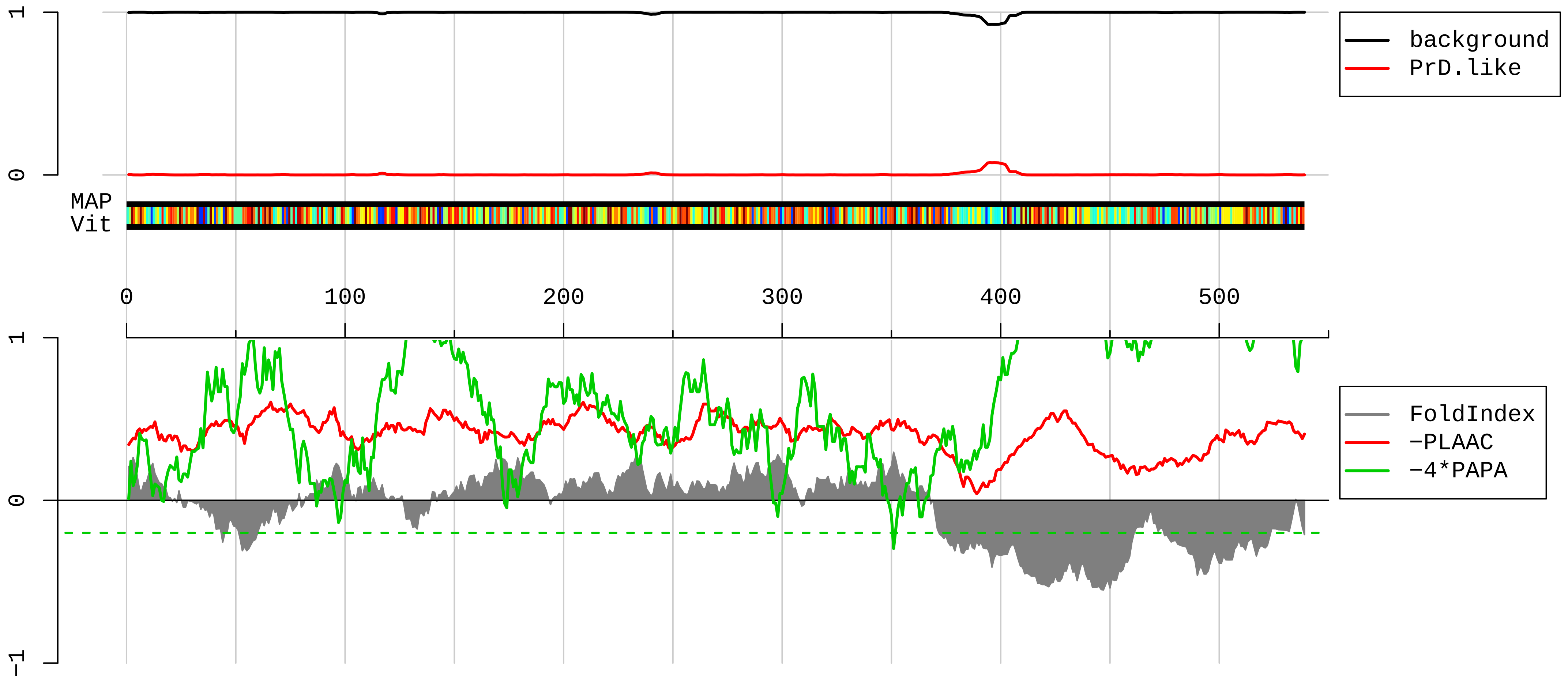


1 MAELTVEVRGSNGAFYKGFIKDVHEDSLTVVFENNWQPERQVPFNEVRLPPPPDIKKEISEGDEVEVYSRANDQEPCGWWLAKVRMMKGEFYVIEYAACD
101 ATYNEIVTFERLRPVNQNKTVKKNTFFKCTVDVPEDLREACANENAHKDFKKAVGACRIFYHPETTQLMILSASEATVKRVNILSDMHLRSIRTKLMLMS
201 RNEEATKHLECTKQLAAAFHEEFVVREDLMGLAIGTHGSNIQQARKVPGVTAIELDEDTGTFRIYGESAEAVKKARGFLEFVEDFIQVPRNLVGKVIGKN
301 GKVIQEIVDKSGVVRVRIEGDNENKLPREDGMVPFVFVGTKESIGNVQVLLEYHIAYLKEVEQLRMERLQIDEQLRQIGSRSYSGRGRGRRGPNYTSGYG
401 TNSELSNPSETESERKDELSDWSLAGEDDRETRHQRDSRRRPGGRGRSVSGGRGRGGPRGGKSSISSVLKDPDSNPYSLLDNTESDQTADTDASESHHST
501 NRRRRSRRRRTDEDAVLMDGLTESDTASVNENGLGKRCD

AAH16202.1 Dhx30 protein [Mus musculus]

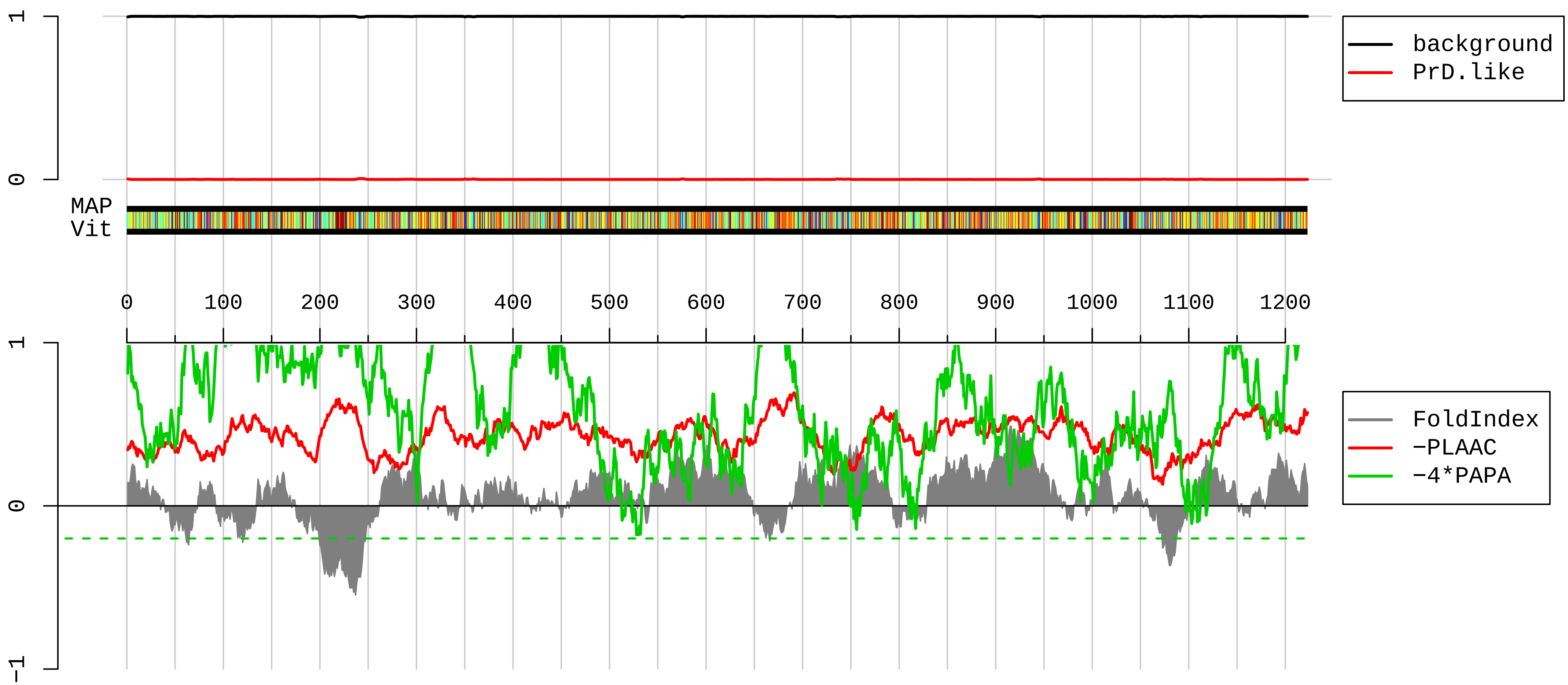


1 MAAARRLMALAAGVSPRLRPPDPLVASGRQGCSRGFSSSFVRSDGTQEAAEVESEVAPSEPGEGDGSMVNASRDLLKEFPQPKNLLNSVIGRALGISHAK
101 DKLVYVHTNGPKKKKVTLHIKWPKSVEVEGYGSKKIDAERQAAAAACQLFKGWGLLGPRNELFDAAKYRVLADRFGSPADSWWRPEPTMPPTSWRQLNPE
201 NIRPGGPAGLSRSLGREEEEDEEEELEEGTIDVTEFLSMTQQDSHNPLRDSRGGSFEMTDDDSAIRALTQFPLPKNLLAKVIQIATSSSTAKNLMQFHTV
301 GTKTKLATLTLLWPCPMTFVAKGRRKAEAENKAAALACKKLKSLGLVDRNNEPLTHAMYNLASLRELGETQRRPCTIQVPEPILRKIEAFLSHYPVDSSW
401 ISPELRLQSDDILPLGKDSGPLSDPITGKPYMPLSEAEEVRLSQSLLELWRRRGPIWQEAPQLPVDPHRDTILSAIEQHPVVVISGDTGCGKTTRIPQLL
501 LERYVTEGRGARCNVIITQPRRISAVSVAQRVSHELGPSLRRNVGFQVRLESKPPARGGALLFCTVGILLRKLQSNPSLEGVSHVIVDEVHERDVNTDFL
601 LILLKGLQRLNPALRLVLMSATGDNERFSRYFGGCPVIKVPGFMYPVKEHYLEDILAKLGKHQYPHRHRHHESEDECALDLDLVTDLVLHIDARGEPGGI
701 LCFLPGWQEIKGVQQRLQEALGMHESKYLILPVHSNIPMMDQKAIFQQPPLGVRKIVLATNIAETSITVNDIVHVVDSGLHKEERYDLKTKVSCLETVWV
801 SRANVIQRRGRAGRCQSGFAYHLFPRSRLEKMVPFQVPEILRTPLENLVLQAKIHMPEKTAVEFLSKAVDSPNIKAVDEAVILLQEIGVLDQREYLTTLG
901 QRLAHISTDPRLAKAIVLAAIFRCLHPLLVVVSCLTRDPFSSSLQNRAEVDKVKALLSHDSGSDHLAFVRAVAGWEEVLRWQDRTSRENYLEENLLYAPS
1001 LRFIHGLIKQFSENIYEAFLVGKPSDCTLPSAQCNEYSEEEELVKGVLMAGLYPNLIQVRQGKVTRQGKFKPNSVTYRTKSGNILLHKSTINREATRLRS
1101 RWLTYFMAVKSNGSVFVRDSSQVHPLAVLLLTDGDVHIRDDGRRATISLSDSDLLRLEGDSRTVRLLREFRRALGRMVERSLRSELAALPLSVQQEHGQL
1201 LALLAELLRGPCGSFDMRKTADD

**EDL15139.1 interleukin enhancer binding factor 2 [Mus musculus]**

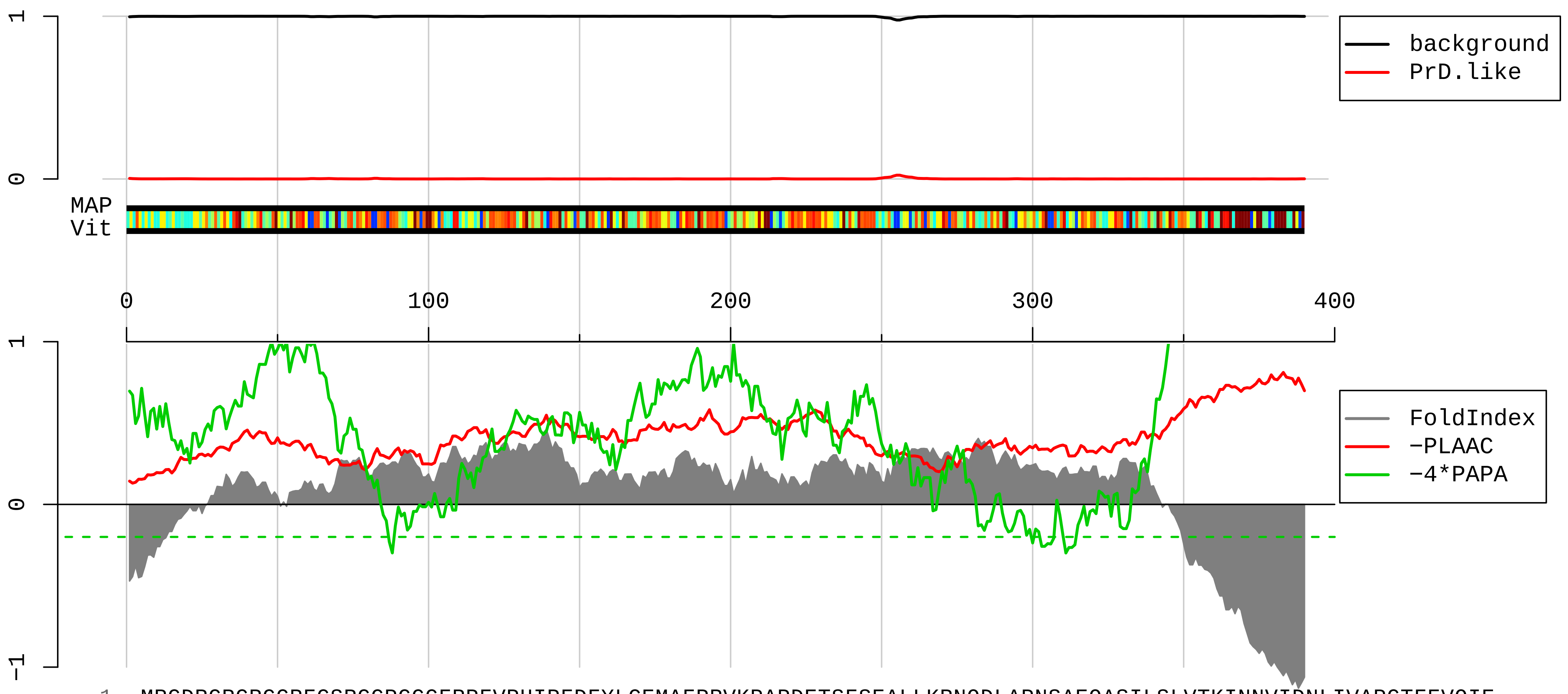


1 MRGDRGRGRGGRFGSRGGPG<u>GGFRP</u>FVPHIPFDFYLCEMAFPRVKPAPDETSFSEALLKRNQDLAPNSAEQASILSLVTKINNVIDNLIVAPGTFEVQIE
101 EVRQVGSYKKGTMTTGHNVADLVVILKILPTLEAVAALGNKVVESLRAQDPSEVLTMLTNETGFEISSSDATVKILITTVPPNLRKLDPELHLDIKVLQS
201 ALAAIRHARWFEENASQSTVKVLIRLLKDLRIRFPGFEPLTPWILDLLGHYAVMNNPTRQPLALNVAYRRCLQILAAGLFLPGSVGITDPCESGNFRVHT
301 VMTLEQQDMVCYTAQTLVRILSHGGFRKILGQEGDASYL<u>ASEISTWDGVIVTPSEKAYEKPPEKKEGEEE</u>EENTEEPPQGEEEESMETQE

**AAH96452.1 Ribosomal protein L7 [Mus musculus]**

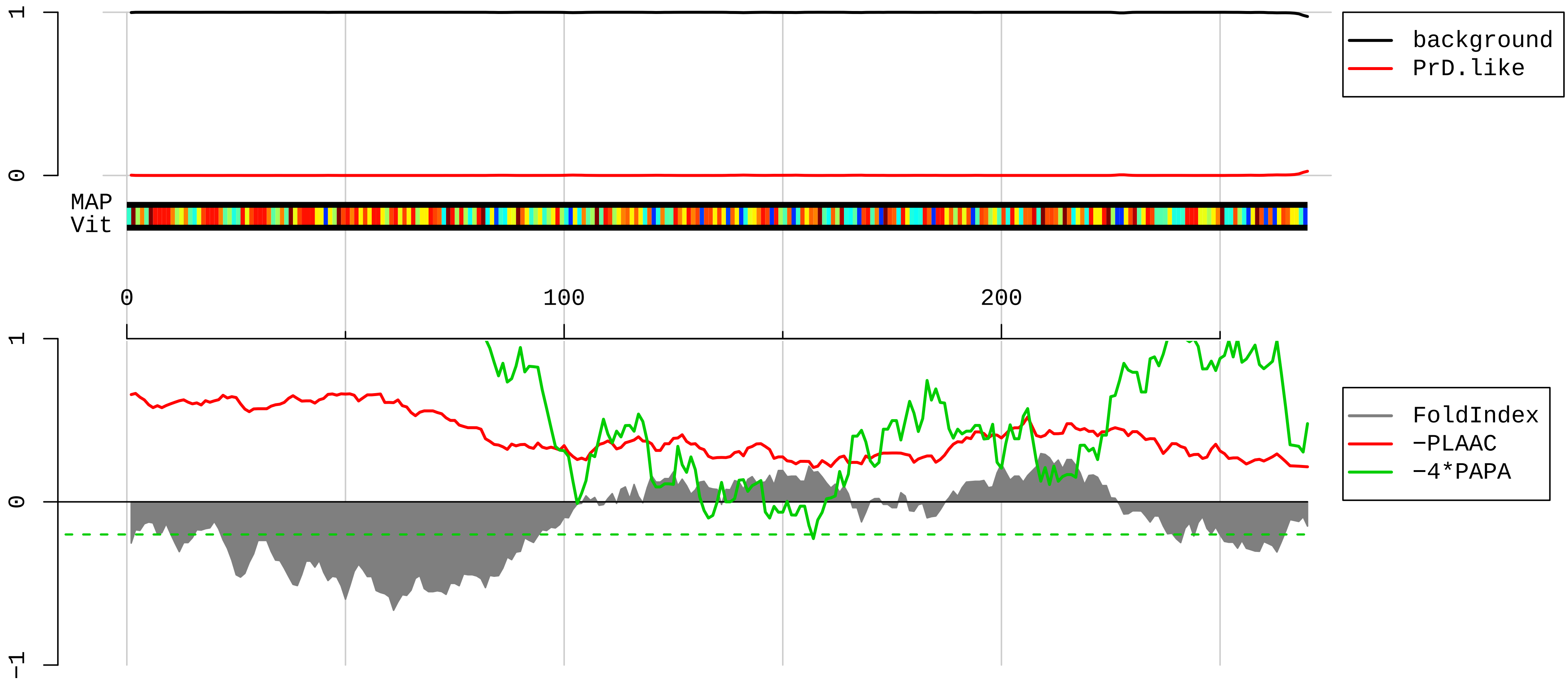


1 MEAVPEKKKKVATVPGTLKKKVPAGPKTLKKKVPAVPETLKKKRRNFAELKVKRLRKKFALKTLRKARRKLIYEKAKHYHKEYRQMYRTEIRMARMARKA
101 GNFYVPAEPKLAFVIRIRGINGVSPKVRKVLQLLRLRQIFNGTFVKLNKASINMLRIVEPYIAWGYPNLKSVNELIYKRGYGKINKKRIALTDNSLIARS
201 LGKFGIICMEDLIHEIYTVGKRFKEANNFLWPFKLSSPRGGMKKKTTHFVEGGDAGNREDQINRLIRRMN

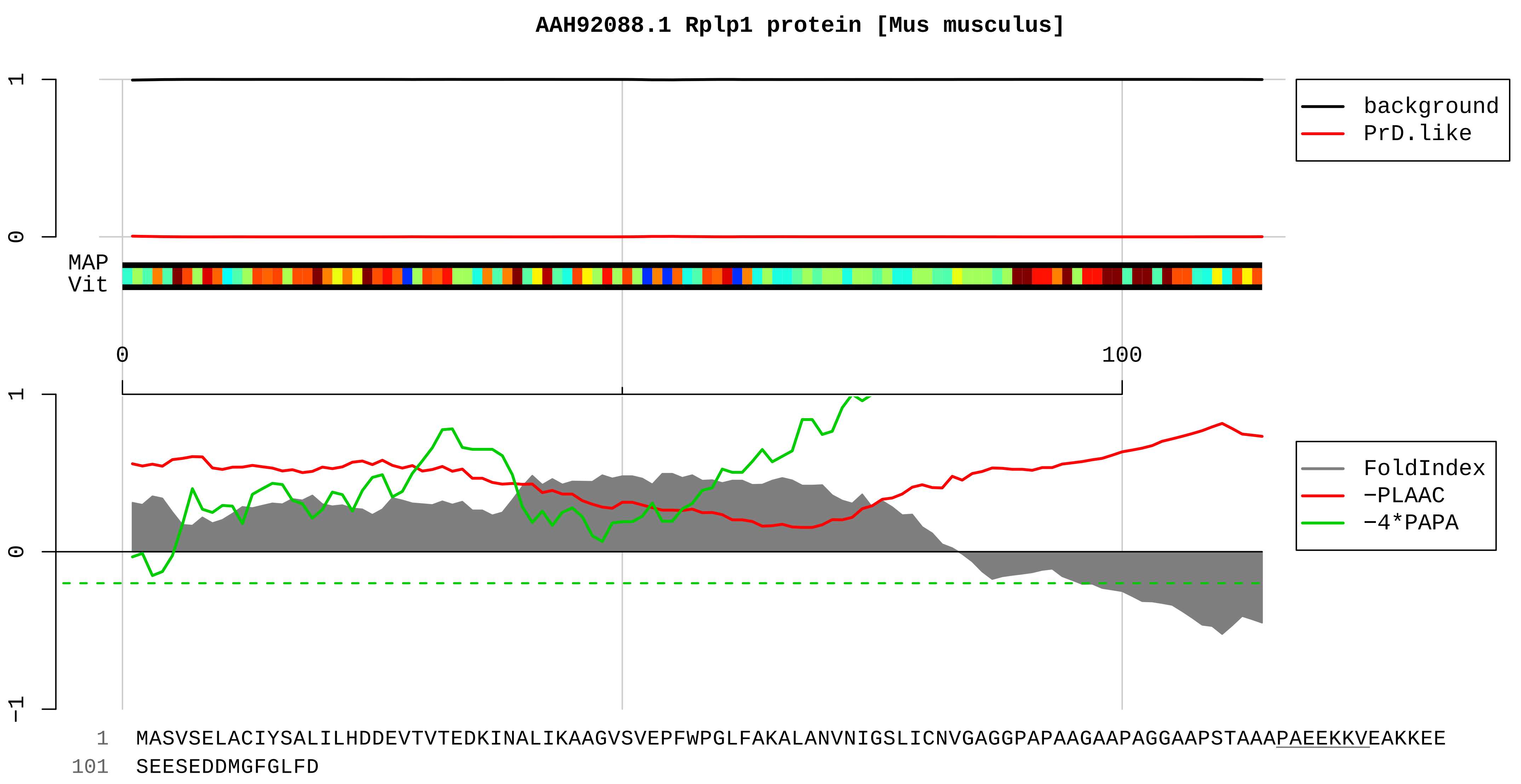
AAH92088.1 Rplp1 protein [Mus musculus]
1
0
background
PrD.like
MAP
Vit
0
100
1
0
−1
FoldIndex
−PLAAC
−4*PAPA
1 MASVSELACIYSALILHDDEVTVTEDKINALIKAAGVSVEPFWPGLFAKALANVNIGSLICNVGAGGPAPAAGAAPAGGAAPSTAAAPAEEKKVEAKKEE
101 SEESEDDMGFGLFD

**AAH85256.1 Tubulin, alpha 1A [Mus musculus]**

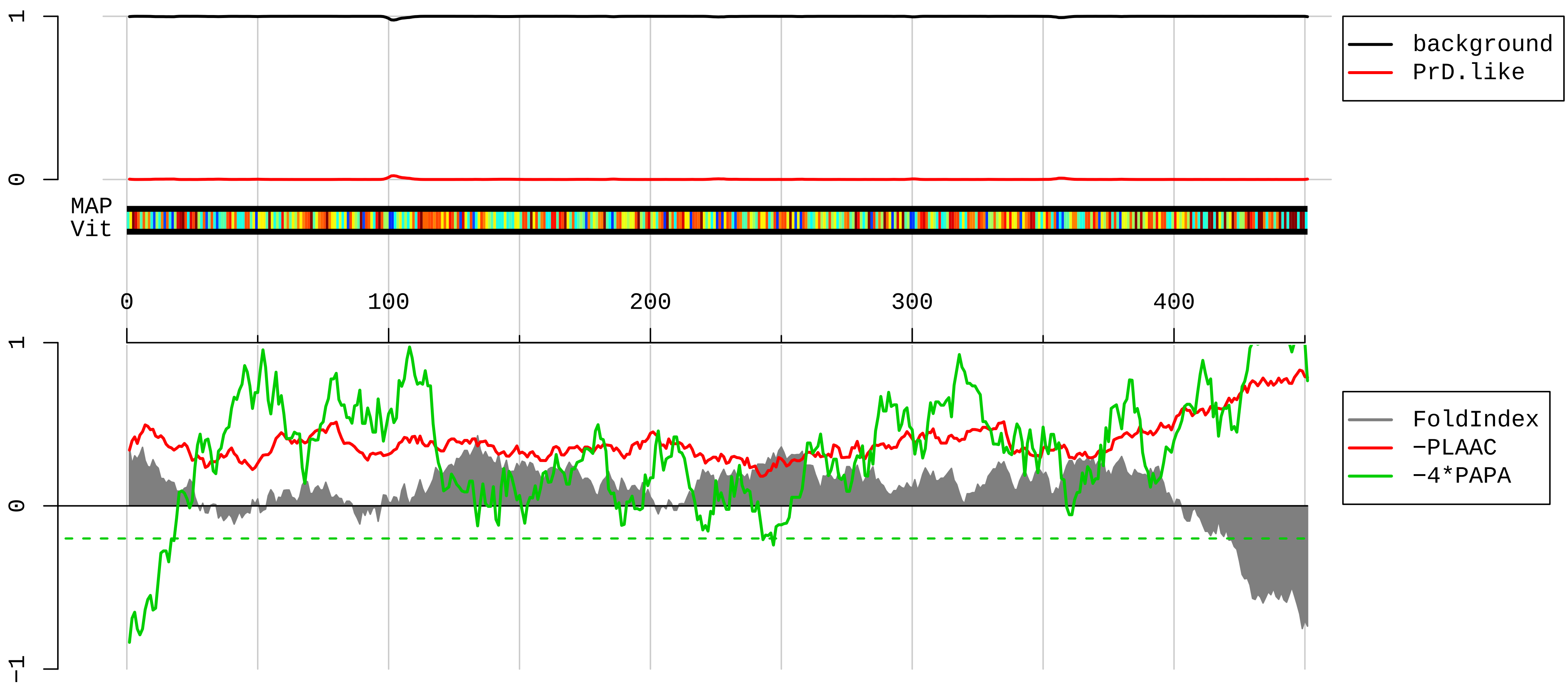


1 MRECISIHVGQAGVQIGNACWELYCLEHGIQPDGQMPSDKTIGGGDDSFNTFFSETGAGKHVPRAVFVDLEPTVIDEVRTGTYRQLFHPEQLITGKEDAA
101 NNYARGHYTIGKEIIDLVLDRIRKLADQCTGLQGFLVFHSFGGGTGSGFTSLLMERLSVDYGKKSKLEFSIYPAPQVSTAVVEPYNSILTTHTTLEHSDC
201 AFMVDNEAIYDICRRNLDIERPTYTNLNRLIGQIVSSITASLRFDGALNVDLTEFQTNLVPYPRIHFPLATYAPVISAEKAYHEQLSVAEITNACFEPAN
301 QMVKCDPRHGKYMACCLLYRGDVVPKDVNAAIATIKTKRTIQFVDWCPTGFKVGINYQPPTVVPGGDLAKVQRAVCMLSNTTAIAEAWARLDHKFDLMYA
401 KRAFVHWYVGEGMEEGEFSEAREDMAALEKDYEEVGVDSVEGEGEEEGEEY

**AAH86909.1 Ribosomal protein L32 [Mus musculus]**

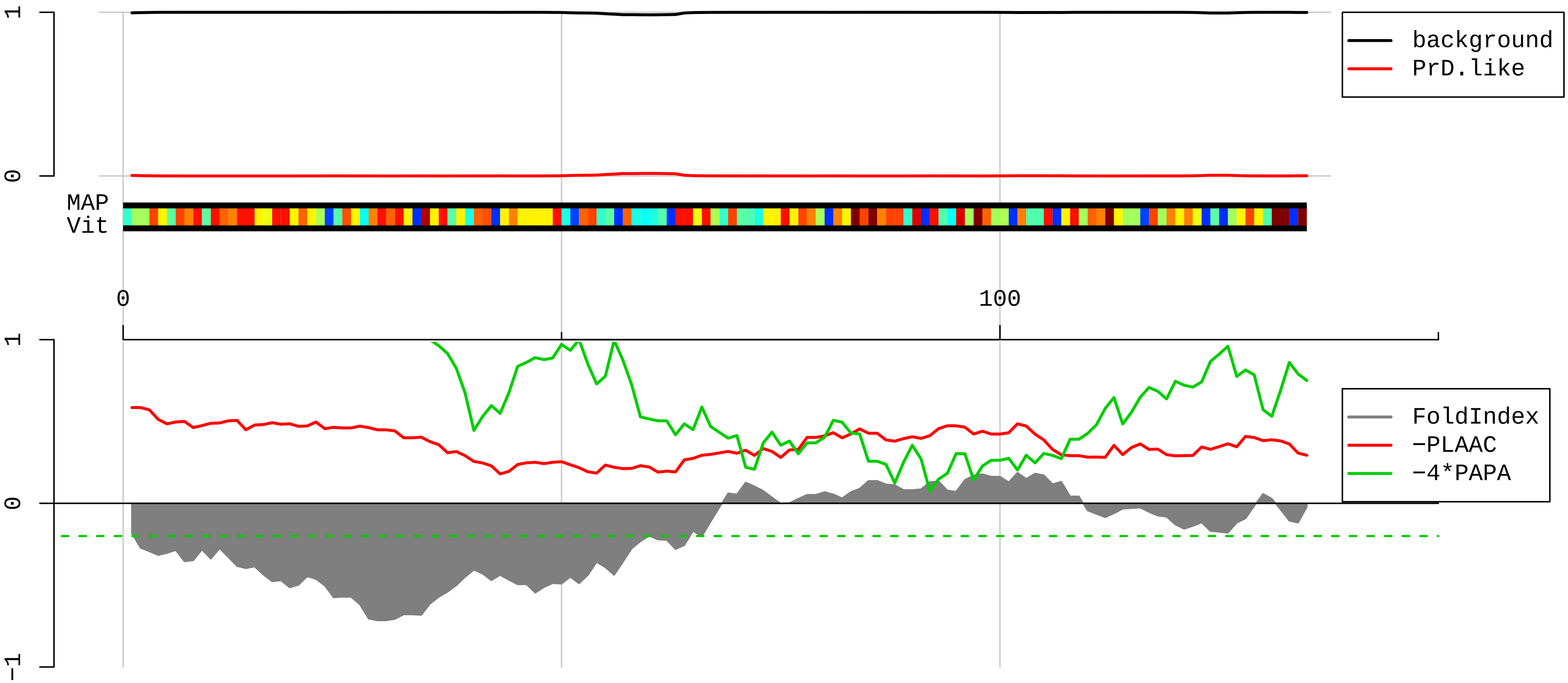


1 MAALRPLVKPKIVKKRTKKF<u>IRHQSDRYVKIKRNWRKPRGIDNRVRRRFKGQILMPNIGYGSNKKTKHMLPSGFRKF</u>LVHNVRELEVLLMCNKSYCAEIA
101 HNVSSKNRKAIVERAAQLAVRVTNPNARLRSEENE

**AAH31746.1 Ribosomal protein S9 [Mus musculus]**

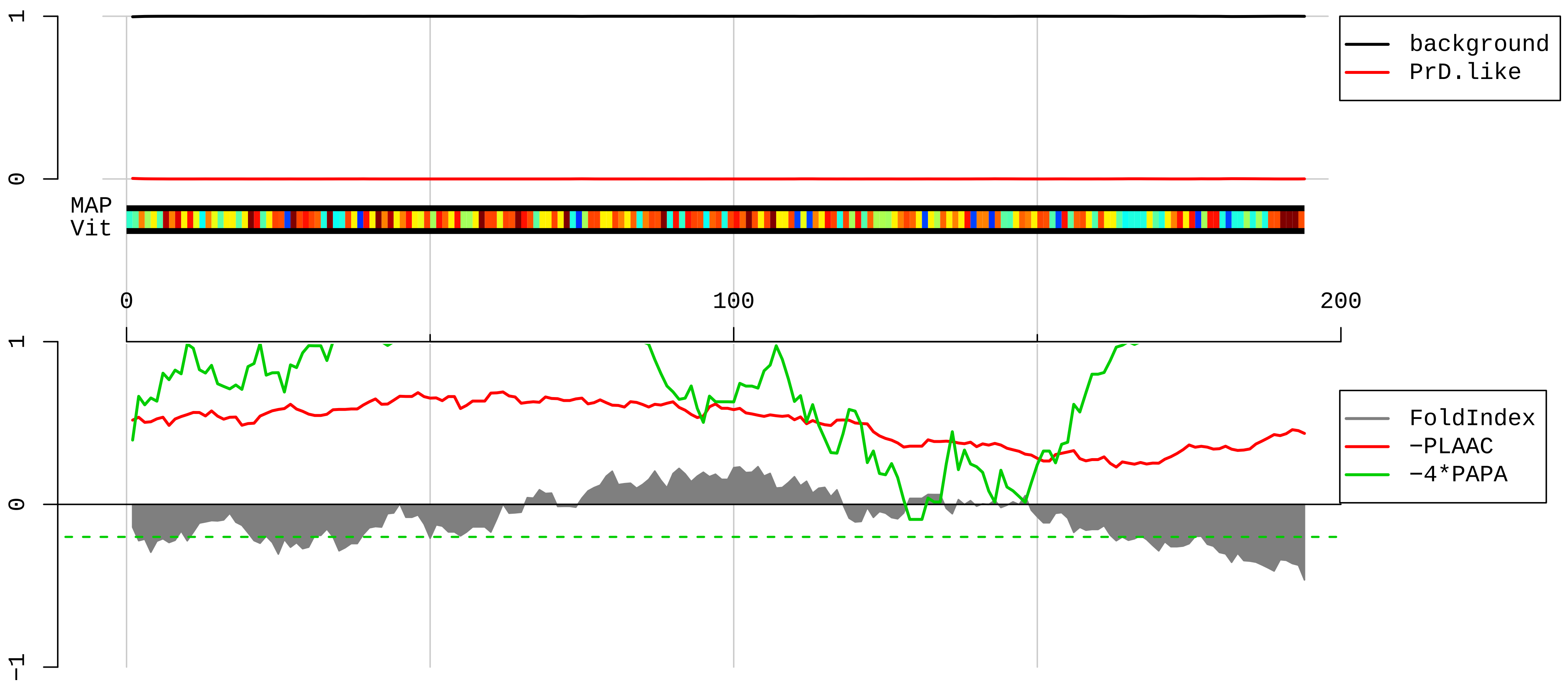


1 MPVARSWVCRKTYVTPRRPFEKSRLDQELKLIGEYGLRNKREVWRVKFTLAKIRKAARELLTLDEKDPRRLFEGNALLRRLVRIGVLDEGKMKLDYILGL
101 KIEDFLERRLQTQVFKLGLAKSIHHARVLIRQRHIRVRKQVVNIPSFIVRLDSQKPIDFSLRFPYGGGRPGRVKRKNAKKGQGGAGAGDDEEED

**AAI06116.1 Ribosomal protein S3A [Mus musculus]**

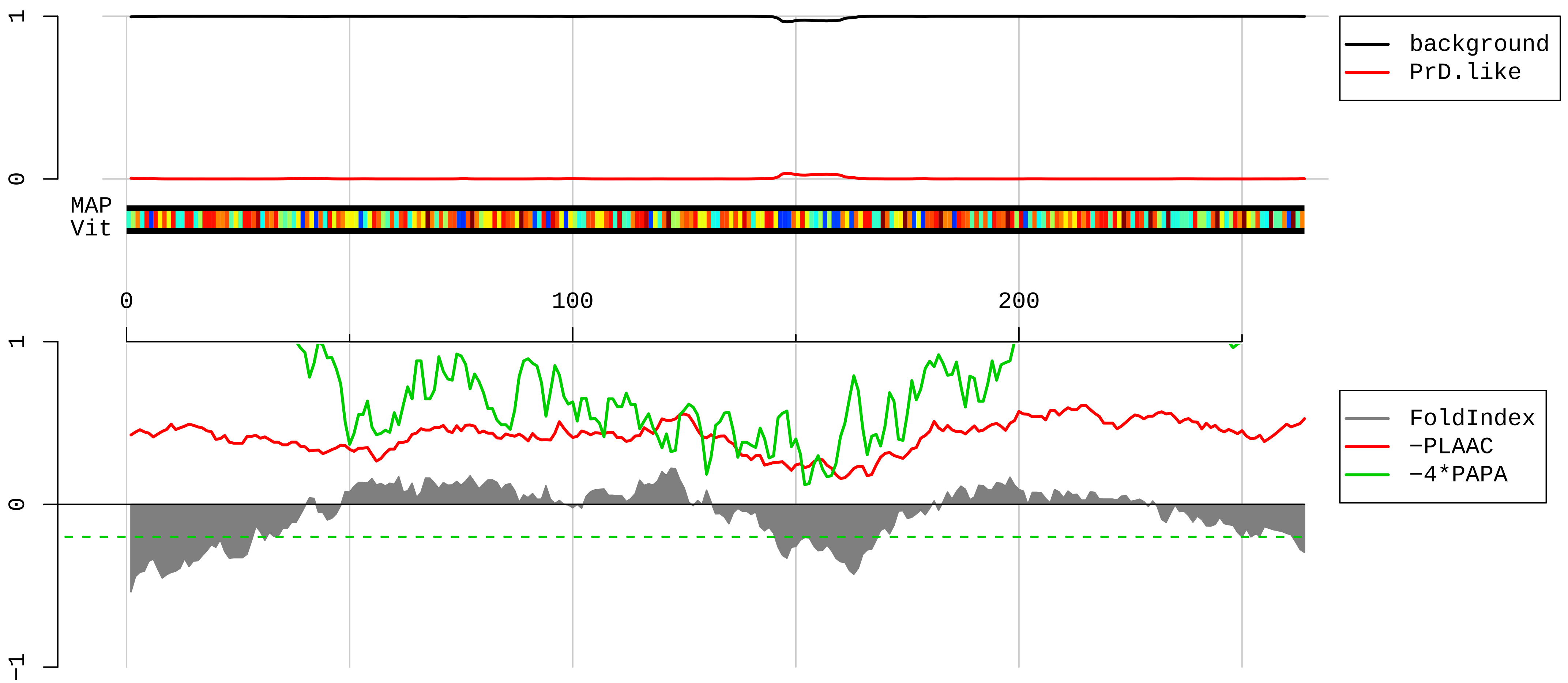


1 MAVGKNKRLTKGGKKGAKKKVVDPFSKKDWYDVKAPAMFNIRNIGKTLVTRTQGTKIASDGLKGRVFEVSLADLQNDEVAFRKFKLITEDVQGKNCLTNF
101 HGMDLTRDKMCSMVKKWQTMIEAHVDVKTTDGYLLRLFCVGFTKKRNNQIRKTSYAQHQQVRQIRKKMMEIMTREVQTNDLKEVVNKLIPDSIGKDIEKA
201 CQSIYPLHDVFVRKVKMLKKPKFELGKLMELHGEGGSSGKAAGDETGAKVERADGYEPPVQESV

#### AAH10763.1 Ribosomal protein S15 [Mus musculus]

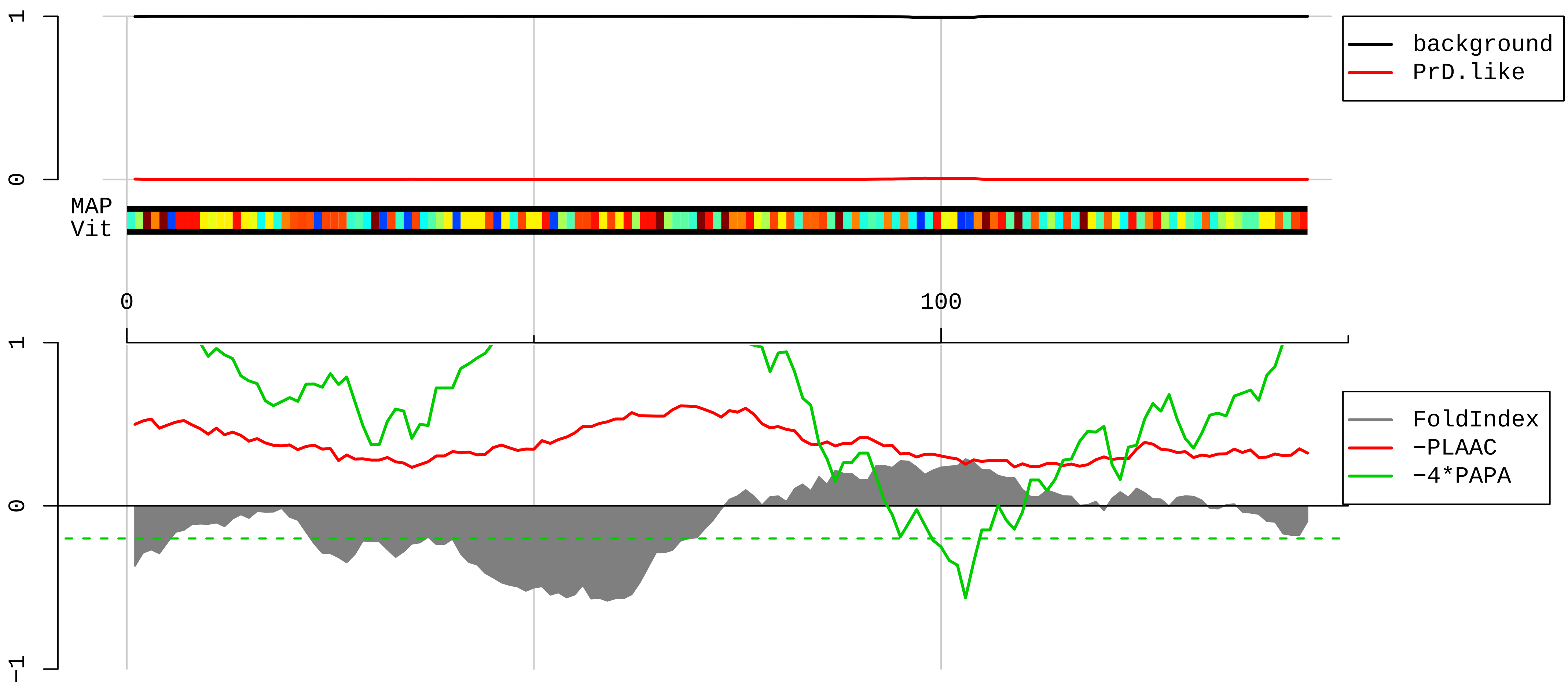


1 MAEVEQKKKRTFRKFTYRGVDLDQLLDMSYEQLMQLYSARQRRRLNRGLRRKQHSLLKRLRKAKKEAPPMEKPEVVKTHLRDMIILPEMVGSMVGVYNGK
101 TFNQVEIKPEMIGHYLGEFSITYKPVKHGRPGIGATHSSRFIPLK

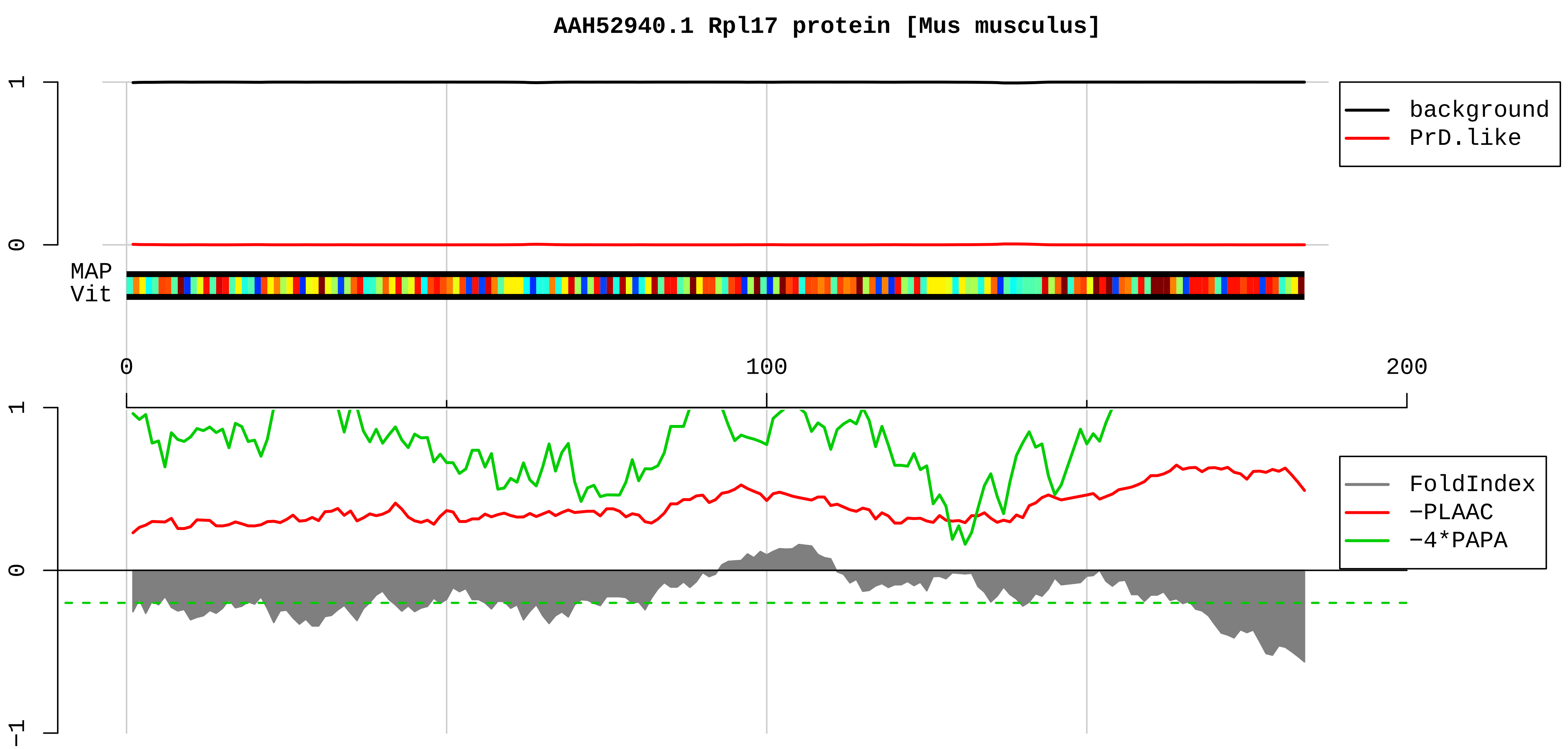


1 MVRYSLDPENPTKSCKSRGSNLRVHFKNTRETAQAIKGMHIRKATKYLKDVTLQKQCVPFRRYNGGVGRCAQAKQWGWTQGRWPKKSAEFLLHMLKNAES

101 NAELKGLDVDSLVIEHIQVNKAPKMRRRTYRAHGRINPYMSSPCHIEMILTEKEQIVPKPEEEVAQKKKISQKKLKKQKLMARE

**AAH81449.1 Ribosomal protein S14 [Mus musculus]**

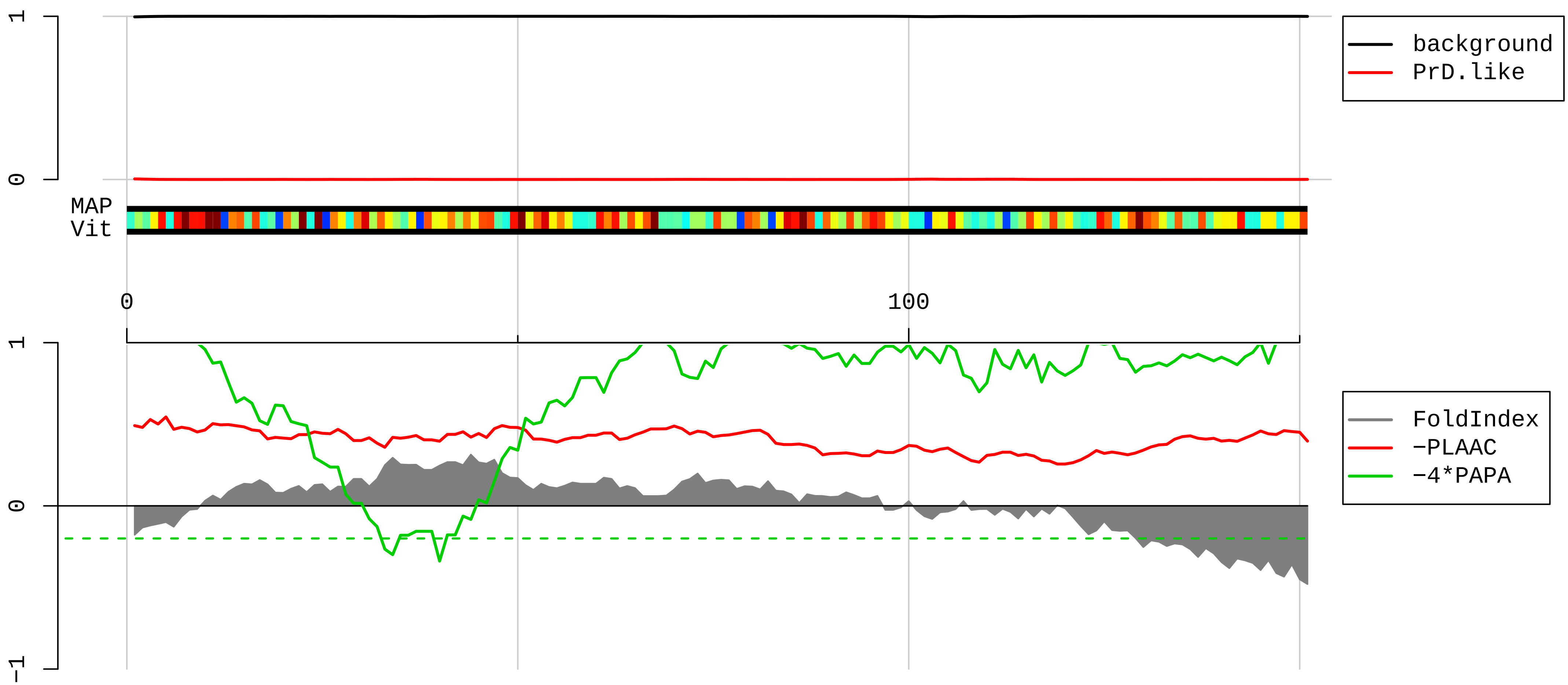


1 MAPRKGKEKKEEQVISLGPQVAEGENVFGVCHIFASFNDTFVHVTDLSGKETICRVTGGMKVKADRDESSPYAAMLAAQDVAQRCKELGITALHIKLRAT
101 GGNRTKTPGPGAQSALRALARSGMKIGRIEDVTPIPSDSTRRKGGRRGRRL

**AAI00595.1 Ribosomal protein S8 [Mus musculus]**

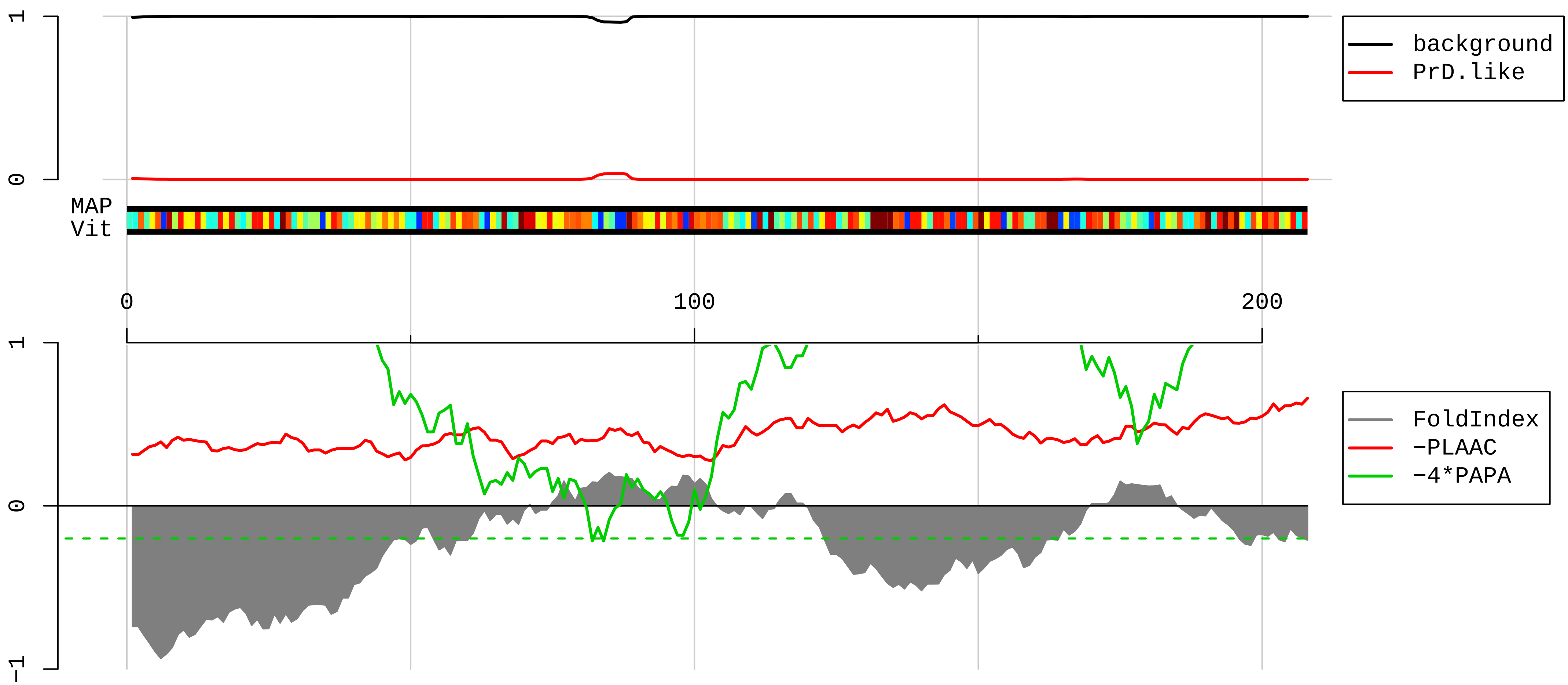


1 MGISRDNWHKRRKTGGKRKPYHKKRKYELGRPAANTKIGPRRIHTVRVRGGNKKYRALRLDVGNFSWGSECCTRKTRIIDVVYNASNNELVRTKTLVKNC
101 IVLIDSTPYRQWYESHYALPLGRKKGAKLTPEEEEILNKKRSKKIQKKYDERKKNAKISSLLEEQFQQGKLLACIASRPGQCGRADGYVLEGKELEFYLR
201 KIKARKGK

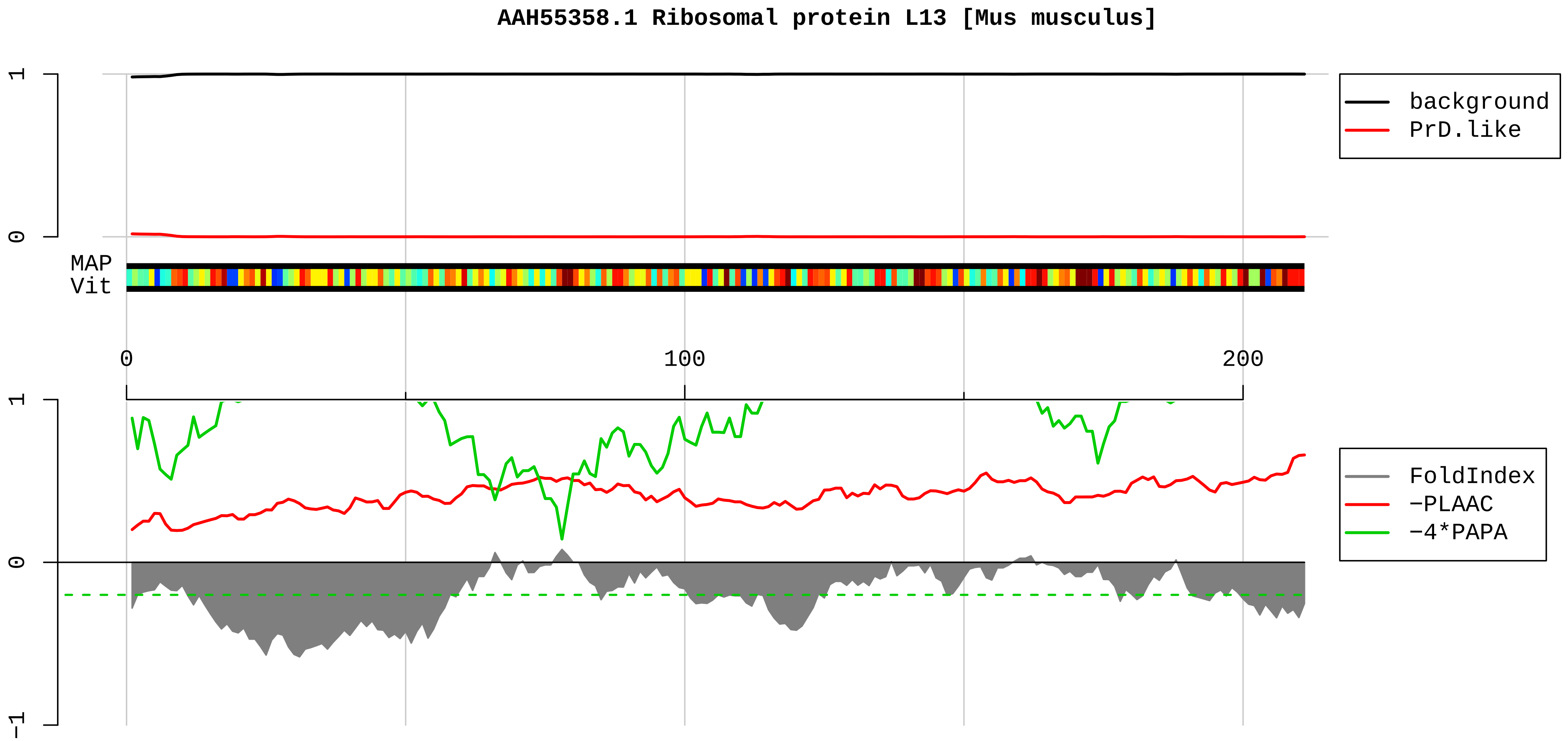


```
1    MAPSRNGMILKPHFHKDWQQRVDTWFNQPARKIRRRKARQAKARRIAPRPASGPIRPIVRCPTVRYHTKVRAGRGFSLEELRVAGIHKKVARTIGISVDP
101  RRRNKSTESLQANVQRLKEYRSKLILFPRKPSAPKKGDSSAEELKLATQLTGPVMPIRNVYKKEKARVITEEEKNFKAFASLRMARANARLFGIRAKRAK
201  EAAEQDVEKKK
```

**AAH12413.1 Ribosomal protein, large P2 [Mus musculus]**

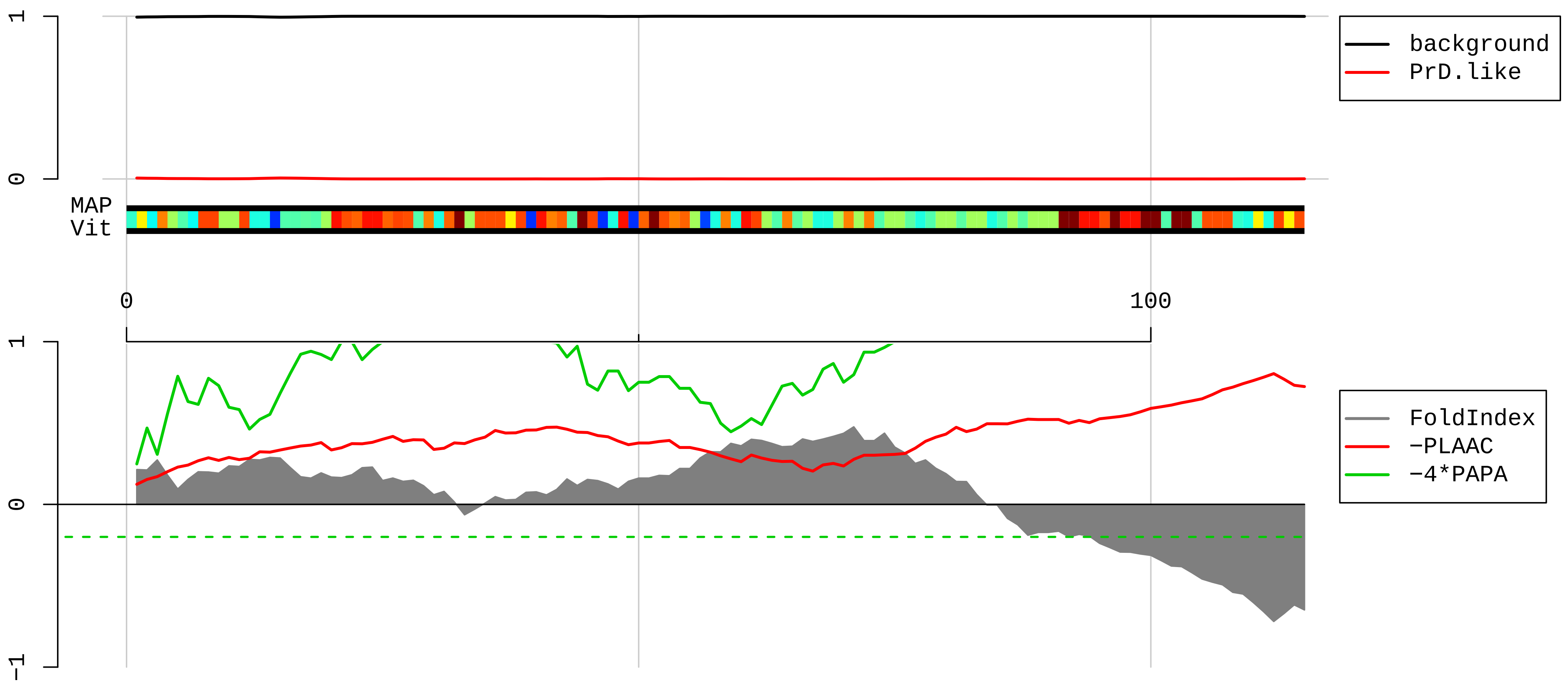


```
  1  MRYVASYLLAALGGNSSPSAKDIKKILDSVGIEADDDRLNKVISELNGKNIEDVIAQGVGKLASVPAGGAVAVSAAPGSAAPAAGSAPAAAEEKKDEKKE
101  ESEESDDDMGFGLFD
```

**AAI25641.1 Ribosomal protein L23a [Mus musculus]**

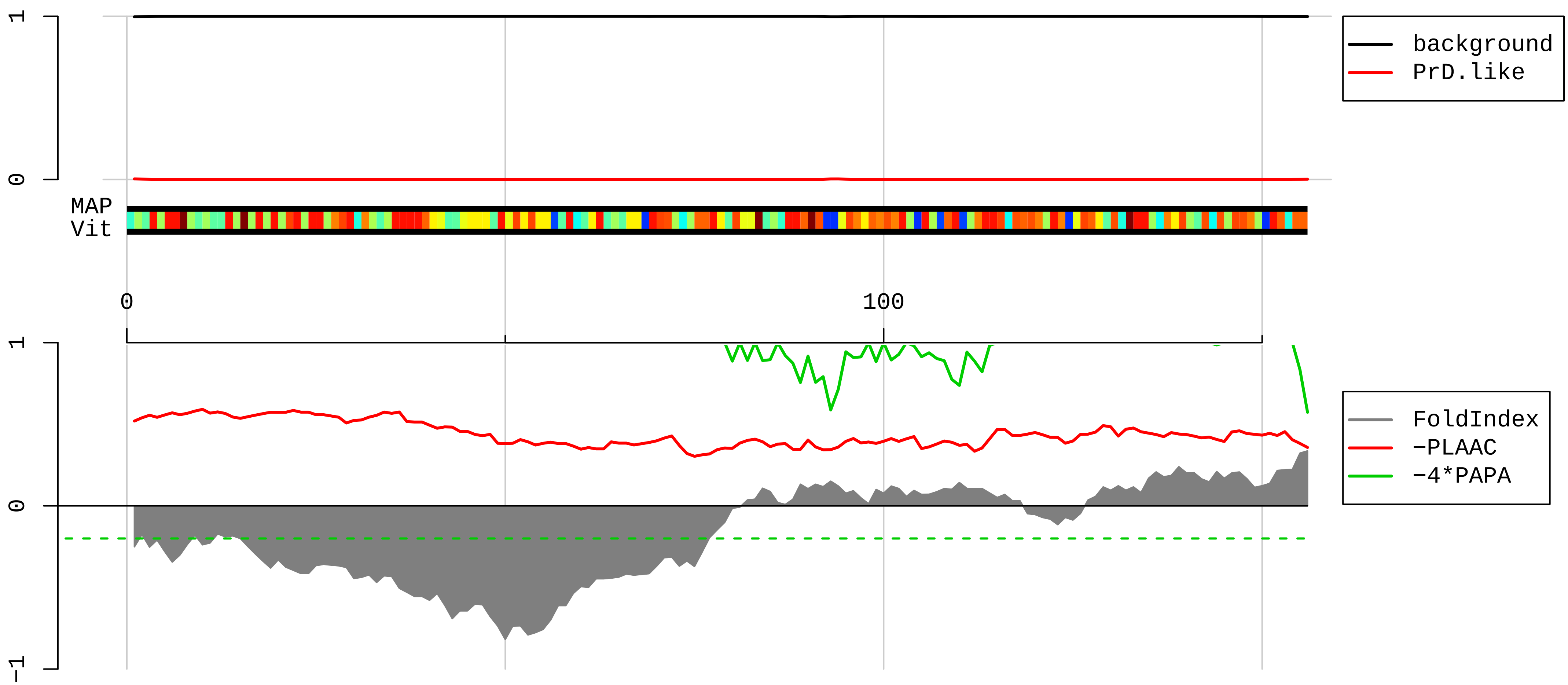


1 MAPKAKKEAPAPPKAEAKAKALKAKKAVLKGVHSHKKKKIRTSPTFRRPKTLRLRRQPKYPRKSAPRRNKLDHYAIIKFPLTTESAMKKIEDNNTLVFIV
101 DVKANKHQIKQAVKKLYDIDVAKVNTLIRPDGEKKAYVRLAPDYDALDVANKIGII

**AAH92050.1 Ribosomal protein S6 [Mus musculus]**

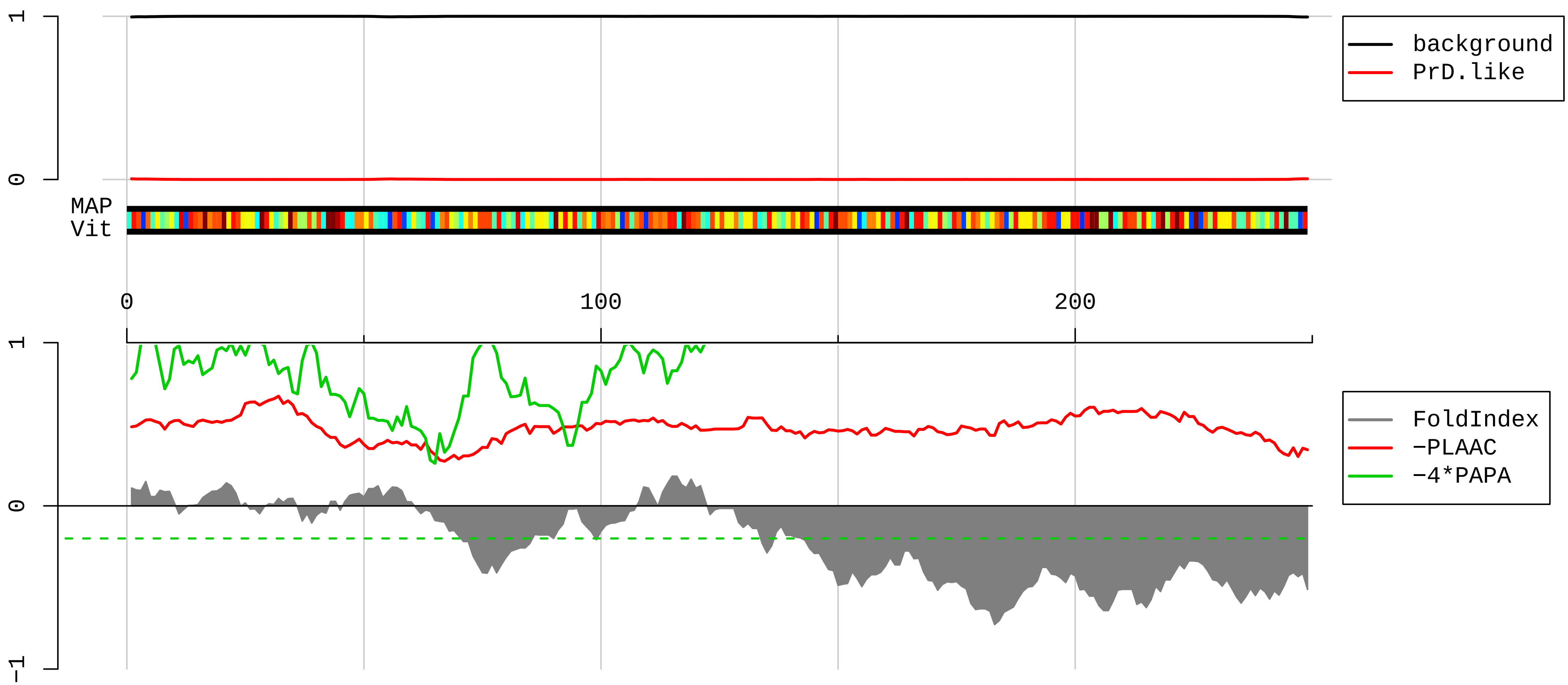


1 MKLNISFPATGCQKLIEVDDERKLRTFYEKRMATEVAADALGEEWKGYVVRISGGNDKQGFPMKQGVLTHGRVRLLLSKGHSCYRPRRTGERKRKSVRGC

101 IVDANLSVLNLVIVKKGEKDIPGLTDTTVPRRLGPKRASRIRKLFNLSKEDDVRQYVVRKPLNKEGKKPRTKAPKIQRLVTPRVLQHKRRRIALKKQRTK

201 KNKEEAAEYAKLLAKRMKEAKEKRQEQIAKRRRLSSLRASTSKSESSQK

**AAI25605.1 Ribosomal protein S13 [Mus musculus]**

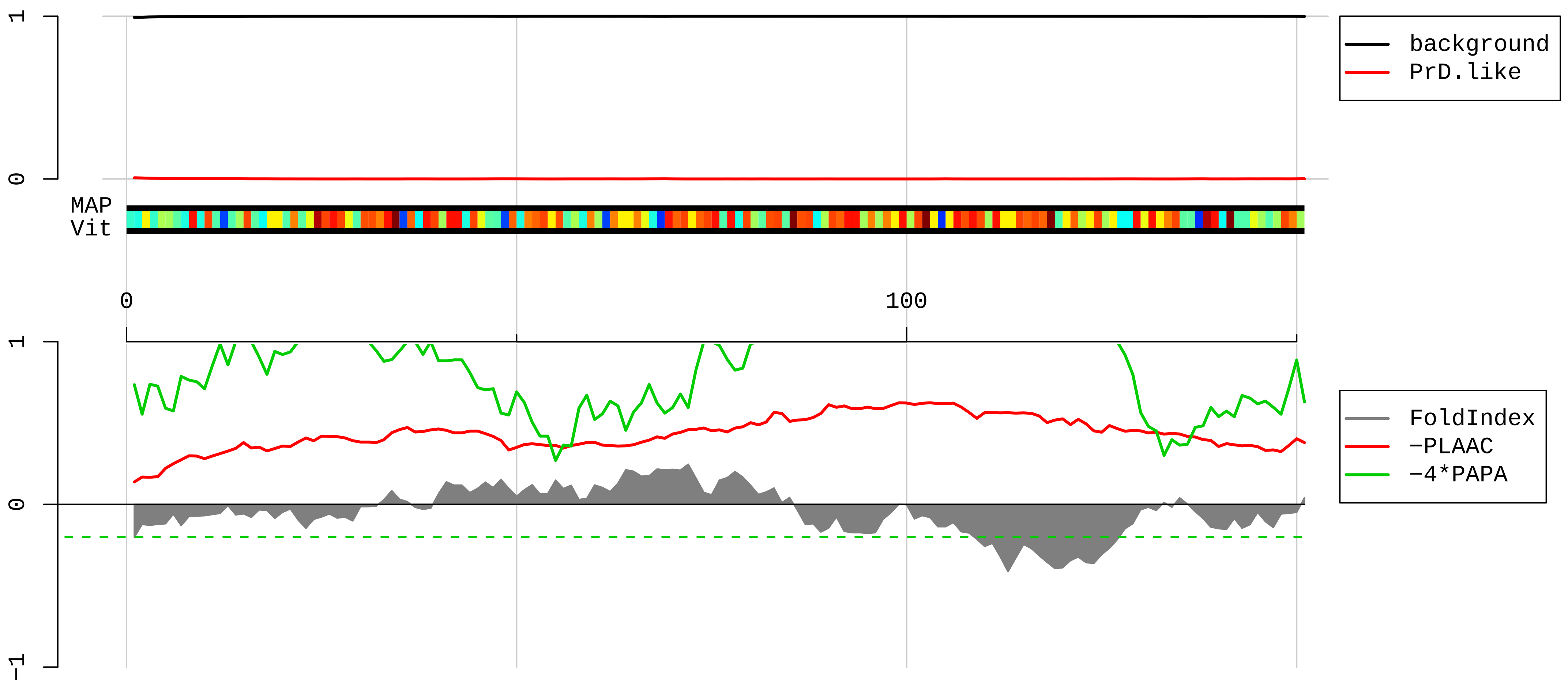


1 MGRMHAPGKGLSQSALPYRRSVPTWLKLTSDDVKEQIYKLAKKGLTPSQIGVILRDSHGVAQVRFVTGNKILRILKSKGLAPDLPEDLYHLIKKAVAVRK

101 HLERNRKDKDAKFRLILIESRIHRLARYYKTKRVLPPNWKYESSTASALVA

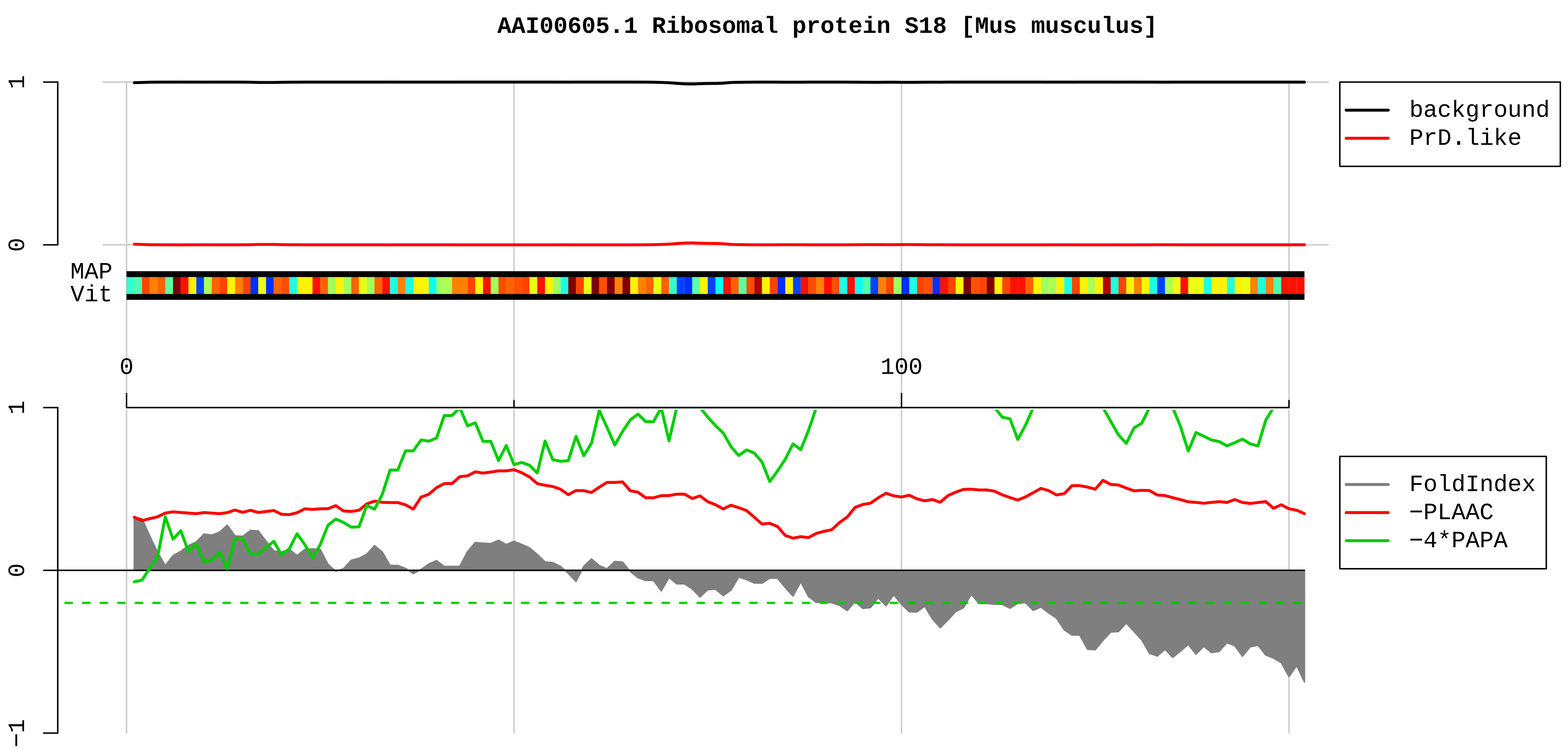


1 MSLVIPEKFQHILRVLNTNIDGRRKIAFAITAIKGVGRRYAHVVLRKADIDLTKRAGELTEDEVERVITIMQNPRQYKIPDWFLNRQKDVKDGKYSQVLA
101 NGLDNKLREDLERLKKIRAHRGLRHFWGLRVRGQHTKTTGRRGRTVGVSKKK

**AAH96392.1 Ribosomal protein S27A [Mus musculus]**

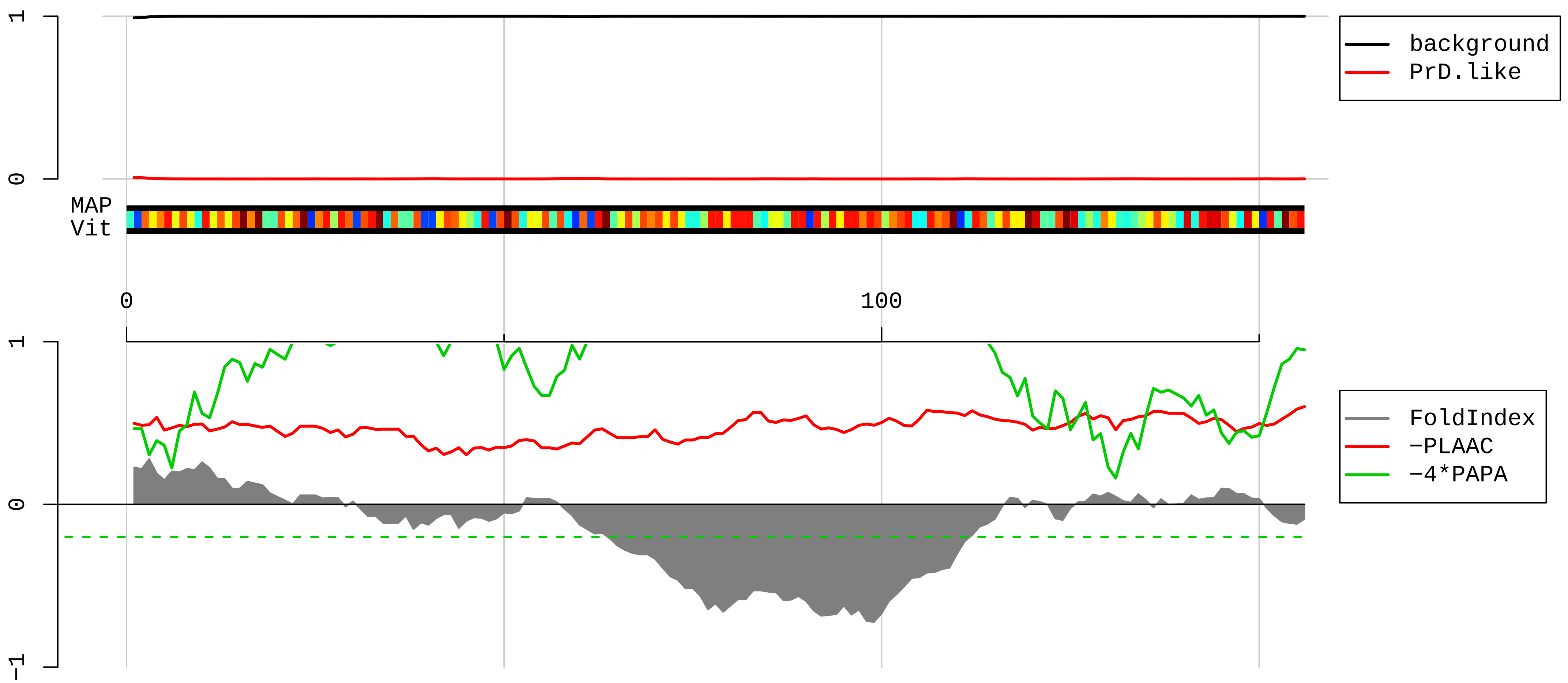


1 MQIFVKTLTGKTITLEVEPSDTIENVKA<u>KIQDKEGIPPDQQRLIFAGKQLEDGRTLSDYNIQKESTLHLVLRLRGGAKKRKKKSYTTPKKNKHKRKKVKL</u>

101 <u>AVLKYYKVDENGKISRLRRECPSDECGAGVF</u>MGSHFDRHYCGKCCLTYCFNKPEDK

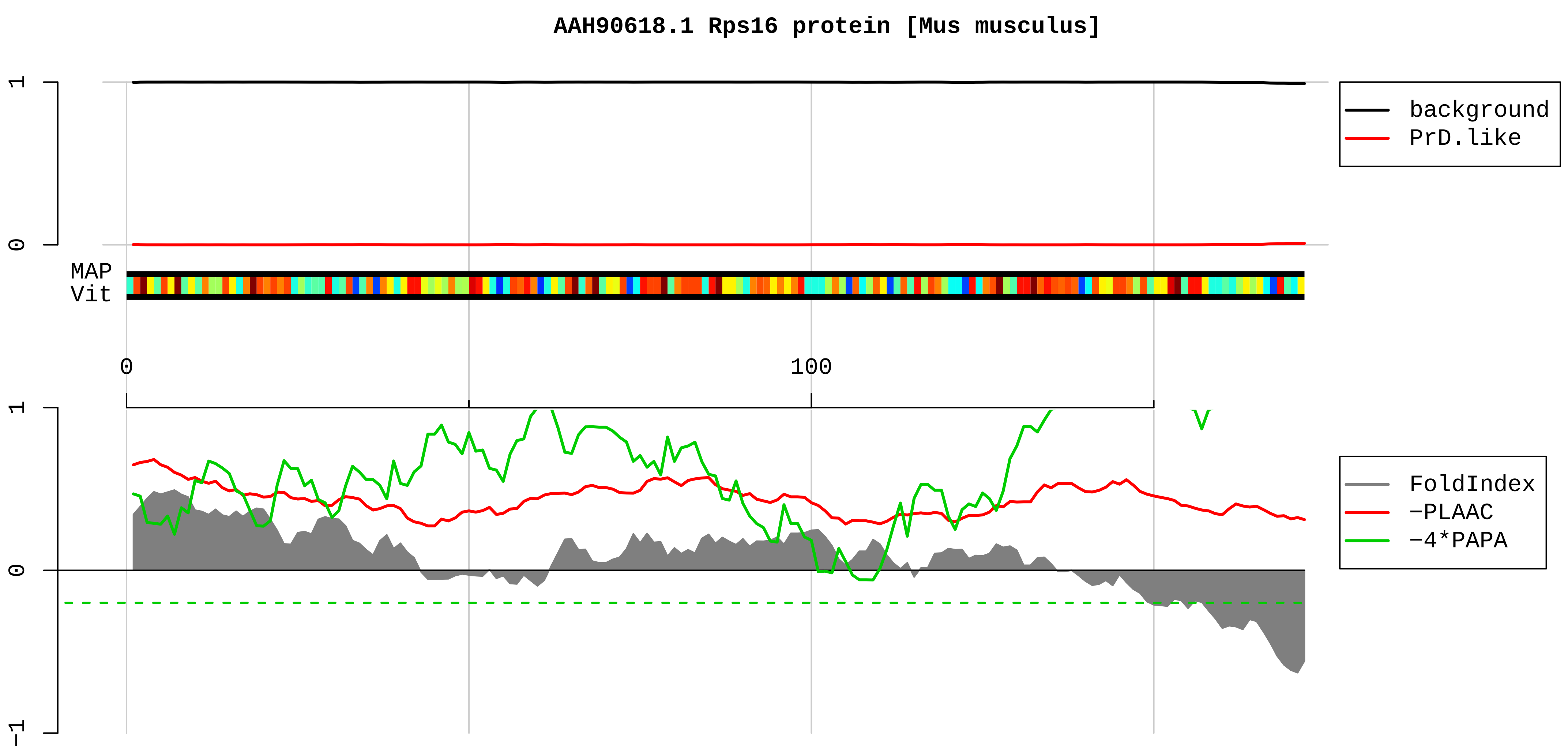


1 MLERPLREPFPVAALRGVELVLVLGAMPSKGPLQSVQVFGRKKTATAVAHCKRGNGLIKVNGRPLEMIEPRTLQYKLLEPVLLLGKERFAGVDIRVRVKG
101 GGHVAQIYAIRQSISKALVAYYQKYVDEASKKEIKDILIQYDRTLLVADPRRCESKKFGGPGARARYQKSYR

**AAH86914.1 Rpl36 protein [Mus musculus]**

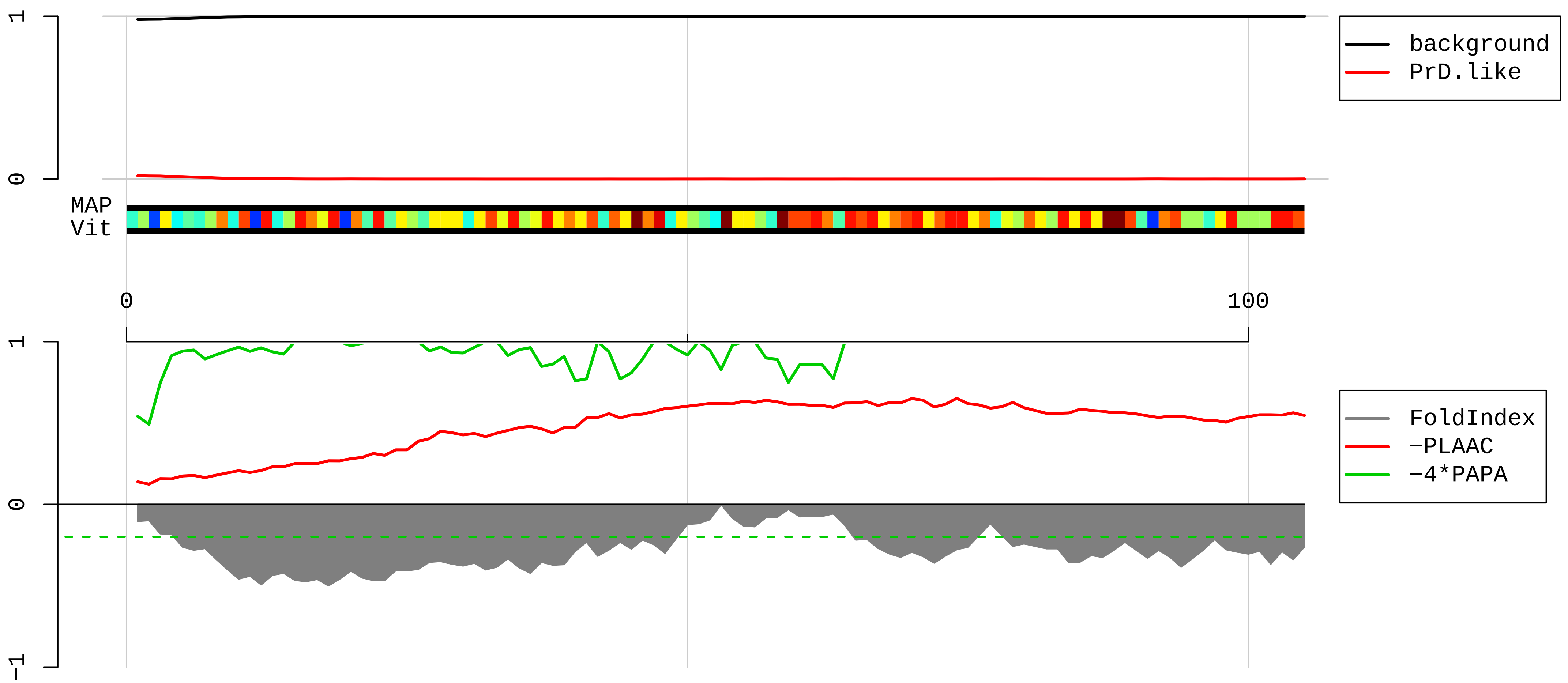


1 MAQRYPMAVGLNKGHKVTKNVSKPRHSRRRGRLTKHTKFVRDMIREVCGFAPYERRAMELLKVSKDKRVLKFIKKRVGTHIRAKRKREELSNVLAAMRKA
101 AAKKD

**CAJ18535.1 Npm1 [Mus musculus]**

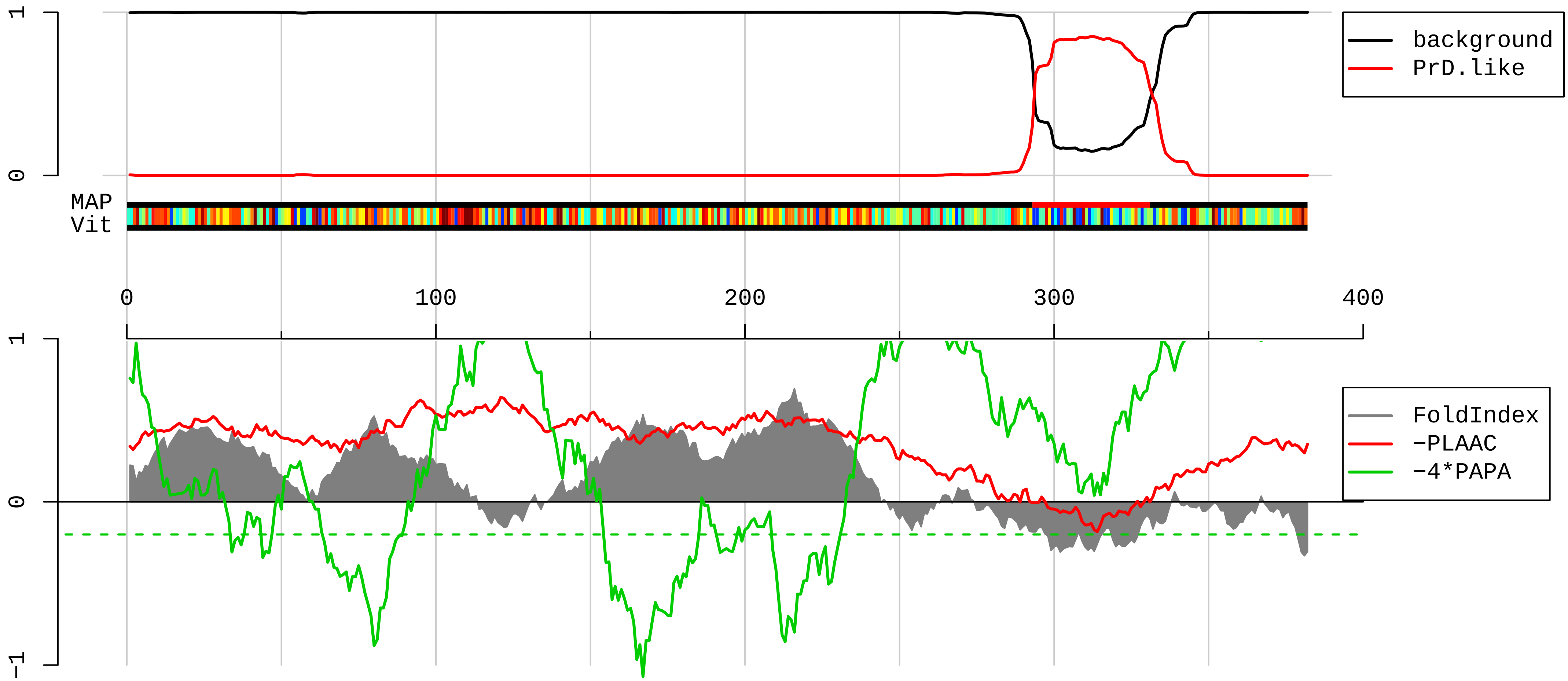


1 MGDWSALGKLLDKVQAYSTAGGKVWLSVLFIFRILLLGTAVESAWGDEQSAFRCNTQQPGCENVCYDKSFPISHVRFWVLQIIFVSVPTLLYLAHVFYVM
101 RKEEKLNKKEEELKVAQTDGVNVEMHLKQIEIKKFKYGIEEHGKVKMRGGLLRTYIISILFKSVFEVAFLLIQWYIYGFSLSAVYTCKRDPCPHQVDCFL
201 SRPTEKTIFIIFMLVVSLVSLALNIIELFYVFFKGVKDRVKGRSDPYHATTGPLSPSKDCGSPKYAYFNGCSSPTAPLSPMSPPGYKLVTGDRNNSSCRN
301 YNKQASEQNWANYSAEQNRMGQAGSTISNSHAQPFDFPDDSQNAKKVAAGHELQPLAIVDQRPSSRASSRASSRPRPDDLEI

**NP_033015.1 transcriptional activator protein Pur-alpha [Mus musculus]**

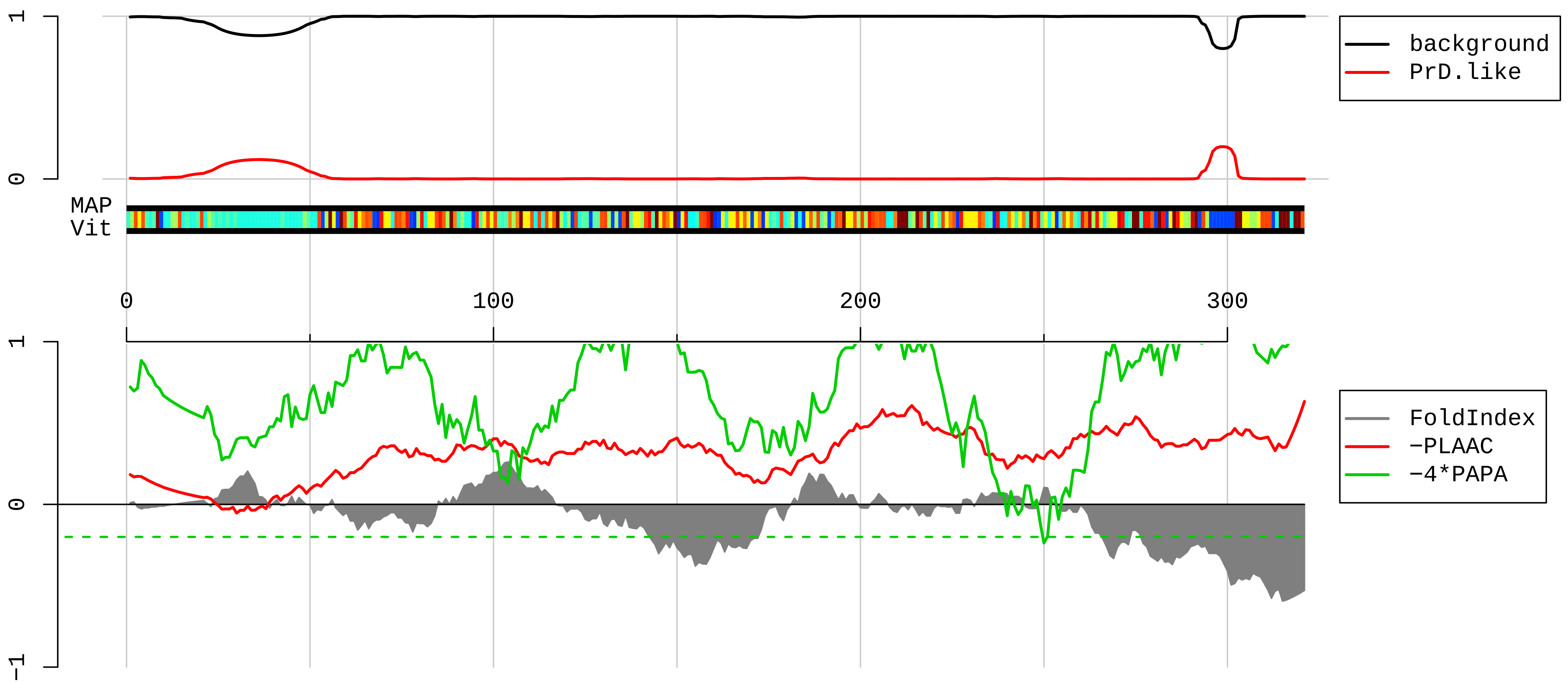


```
  1  MADRDSGSEQGGAALGSGGSLGHPGSGSGSGGGGGGGGGGGGSGGGGGAPGGLQHETQELASKRVDIQNKRFYLDVKQNAKGRFLKIAEVGAGGNKSRLT
101  LSMSVAVEFRDYLGDFIEHYAQLGPSQPPDLAQAQDEPRRALKSEFLVRENRKYYMDLKENQRGRFLRIRQTVNRGPGLGSTQGQTIALPAQGLIEFRDA
201  LAKLIDDYGVEEEPAELPEGTSLTVDNKRFFFDVGSNKYGVFMRVSEVKPTYRNSITVPYKVWAKFGHTFCKYSEEMKKIQEKQREKRAACEQLHQQQQQ
301  QQEETTAATLLLQGEEEGEED
```

**AAH94247.1 Ring finger protein 214 [Mus musculus]**

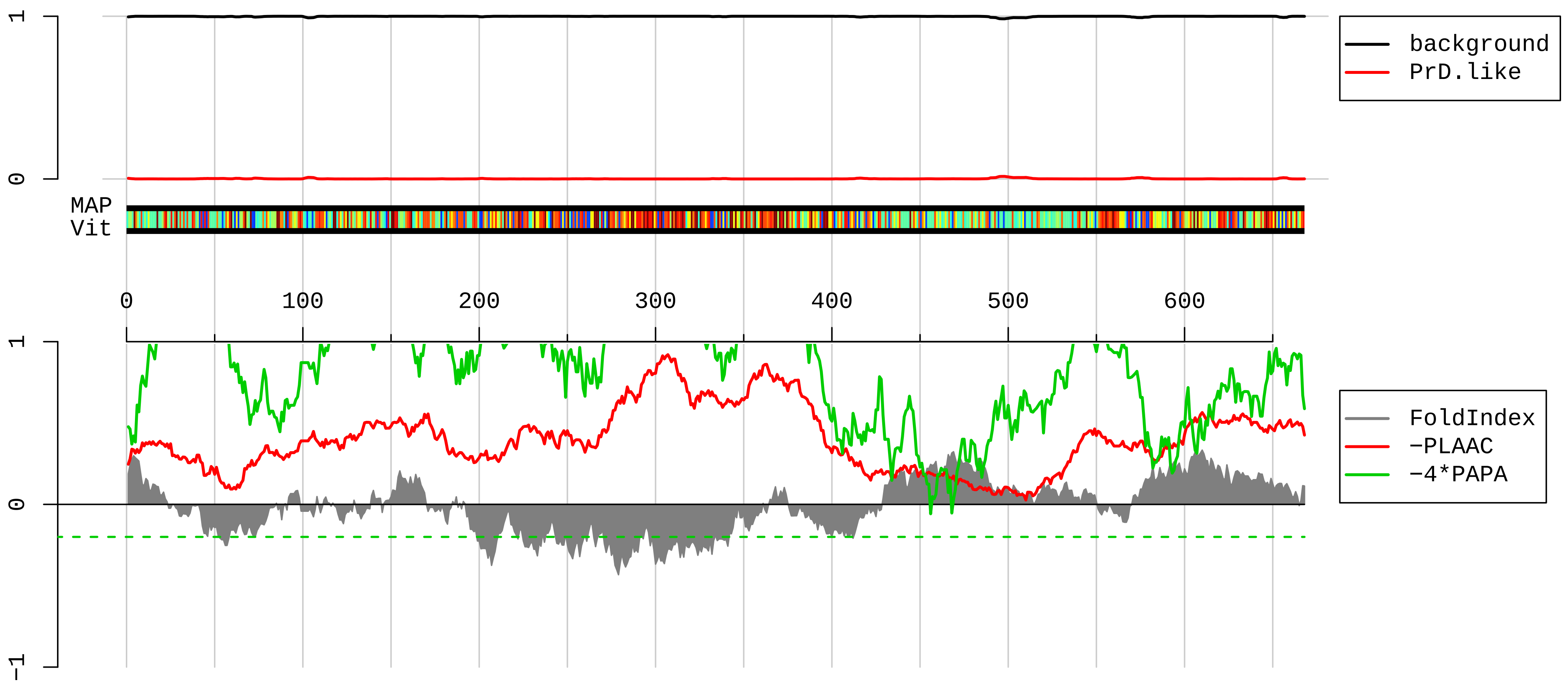


1 MAASEVAGLGAGTPSPSESSALCASKSDESLPDGLSPKDSAQKQKNLSPPSVSSQMITKESNRNAHLEHPEQNPGSSVGDTSAAHEEVVGENLVATALCL
101 SGNGSQSDLKDLTNPAGEEGDTSLRESLHPVTRSLKAGCHSKQLASGNCSEEKCPAASVLKEGSRDAGLDLLPVVPPANGVEGVRVDQDDDQDSSSLKLS
201 QNIAVQTDFKTADSEVNTDQDIEKNLDKMMTERTLLKERYQEVLDKQRQVESQLQVQLKQLQQRREEEMKNHQEILKAIQDVTIKREETKKKIEKEKKEF
301 LQKEQDLKAEIEKLCEKGRREVWEMELDRLKNQDGEINRNIMEETERAWKAEILSLESRKELLVLKLEEAEKEAELHLTYLKSTPPTLETVRSKQEWETR
401 LNGVRIMKKNVRDQFNSHIQLVRNGAKLSSLPQIPTPTLPPPPSEADFMLQVFQPSPSLTPRMPFSIGQVTMPMVMPSADPRSLSFPILNPALSQSSQPS
501 PPLPGSHGRNSPGLGSLVSPHGPHMPPAASIPPPPGLGGIKASSETPRPQPVDKLEKILEKLLTRFPQCNKAQMTNILQQIKTARTTMAGLTMEELIQLV
601 AARLAEHERVASSTQAPPTCKLCLMCQKLVQPSELHPMACTHALHKECIKFWAQTNTNDTCPFCPTLK

**AAQ87664.1 staufen [Mus musculus]**

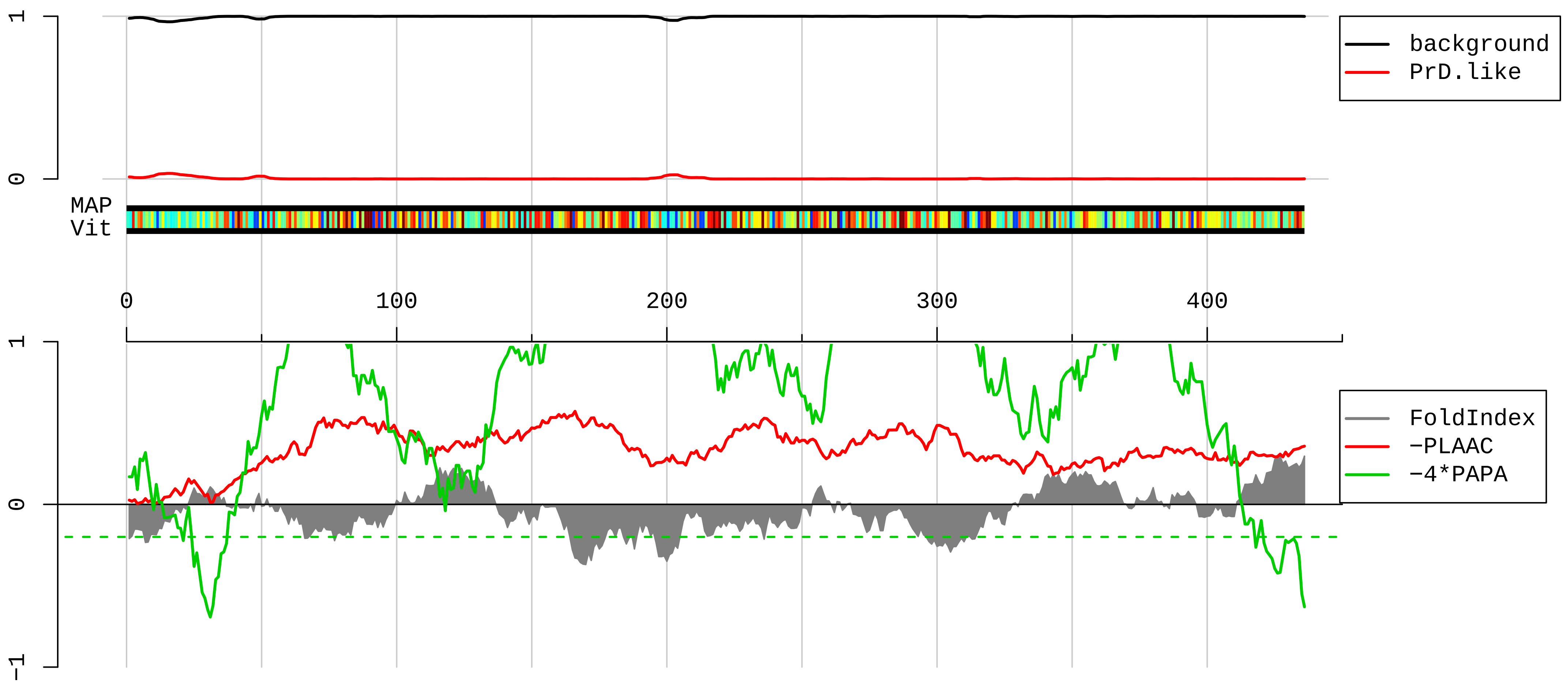


```
  1 MYKPVDPHSRMQSTYSYGMRGGAYPPRYFYPFPVPPLLYQVELSVGGQQFNGKGKMRPPVKHDAPARALRTLQSEPLPERLEVNGREAEEENLNKSEISQ
101 VFEIALKRNLPVNFESFLLTQVARESGPPHMKNVVTRVPVGEFVGEGEGKSKKISKKNAARAVLEQLRRLPPLPAVERVKPRIKKKSQPTCKLQTAPDYG
201 QGMNPISRLAQIQQAKKEKEPEYMLLTERGLPRRREFVMQVKVGHHTAEGVGTNKKVAKRNAAENMLEILGFKVPQAQPAKPALKSEEKTPVKKPGDGRK
301 VTFFEPSPGDENGTSNKDEEFRMPYLSHQQLPAGILPMVPEVAQAVGVSQGHHTKDFTRAAPNPAKATATYPMDLALDPLSNCTTFPEPRDSRLNTKIFP
401 RTTRTSVYLSSTAPHSRLSSVMASARMWSPVMIWLH
```

**AAH95922.1 Hnrpc protein [Mus musculus]**

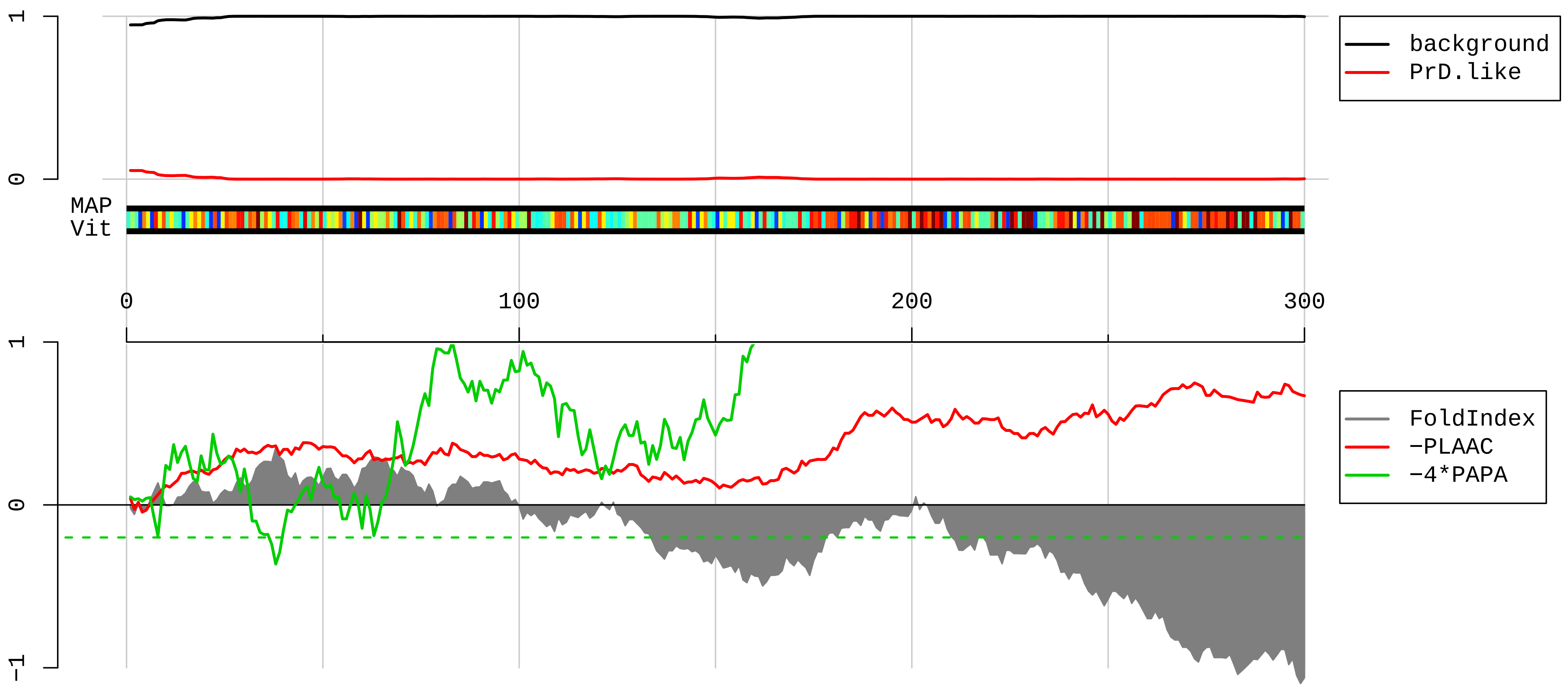


1 MASNVTNKTDPRSMNSRVFIGNLNTLVVKKSDVEAIFSKYGKIVGCSVHKGFAFVQYVNERNARAAVAGEDGRMIAGQVLDINLAAEPKVNRGKAGVKRS
101 AAEMY<u>GSSFDLDYDFQRDYYDRMYSYPARVPPPPPIARAVVPSKRQRVSGNTSRRGKSGFNSKSGQRGSSSKSGKLKGDDLQAIKKELTQIKQKVDSLLE</u>
201 <u>SLEKIEKEQSKQADLSFSSPVEMKNEKSEEEQSSASVKKDETNVKMESEAGADDSAEEGDLLDDDDNEDRGDDQLELKDD</u>EKEPEEGEDDRDSANGEDDS
301

**NP_032717.2 neurofilament medium polypeptide [Mus musculus]**

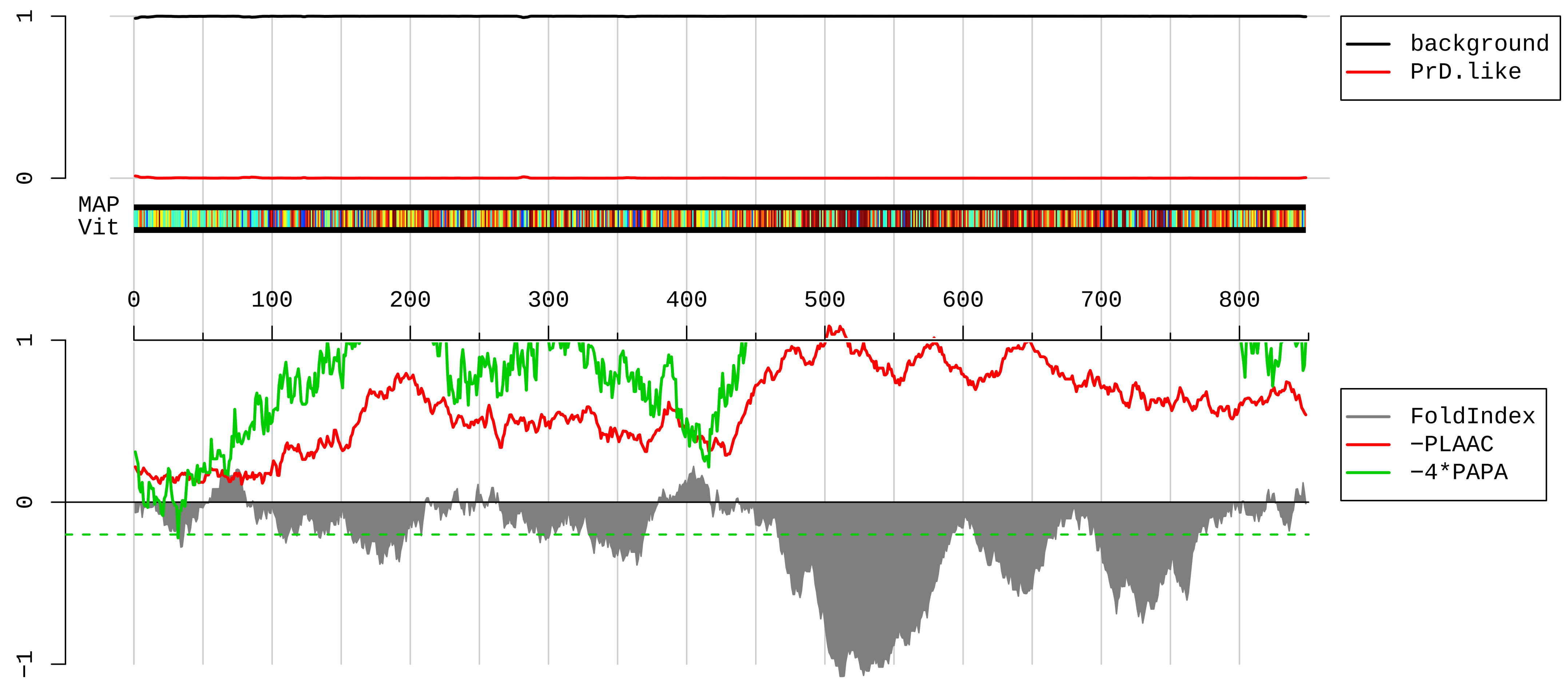


1 MSYTLDSLGNPSAYRRVTETRSSFSRVSGSPSSGFRSQSWSRGSPSTVSSSYKRSALAPRLAYSSAMLSSAESSLDFSQSSSLLNGGSGGDYKLSRSNEK
101 EQLQGLNDRFAGYIEKVHYLEQQNKEIEAEIQALRQKQASHAQLGDAYDQEIRELRATLEMVNHEKAQVQLDSDHLEEDIHRLKERFEEEARLRDDTEAA
201 IRALRKDIEESSMVKVELDKKVQSLQDEVAFLRSNHEEEVADLLAQIQASHITVERKDYLKTDISTALKEIRSQLECHSDQNMHQAEEWFKCRYAKLTEA
301 AEQNKEAIRSAKEEIAEYRRQLQSKSIELESVRGTKESLERQLSDIEERHNHDLSSYQDTIQQLENELRGTKWEMARHLREYQDLLNVKMALDIEIAAYR
401 KLLEGEETRFSTFSGSITGPLYTHRQPSVTISSKIQKTKVEAPKLKVQHKFVEEIIEETKVEDEKSEMEETLTAIAEELAASAKEEKEEAEEKEEEPEAE
501 KSPVKSPEAKEEEEEGEKEEEEEGQEEEEEEDEGVKSDQAEEGGSEKEGSSEKDEGEQEEEEGETEAEGEGEEAEAKEEKKIEGKVEEVAVKEEIKVEKP
601 EKAKSPMPKSPVEEVKPKPEAKAGKGEHKEEEKVEEEKKEVTKESPKEEKVEKKEEKPKDVADKKKAESPVKEKAVEEVITISKSVKVSLEKDTKEEKLQ
701 PQEKVKEKAEEEGGSEEEGSDRSPQESKKEDIAINGEVEGKEEEEQETQEKGSGREEEKGVVTNGLDVSPAEEKKGEDSSDDKVVVTKKVEKITSEGGDG
801 ATKYITKSVTVTQKVEEHEETFEEKLVSTKKVEKVTSHAIVKEVTQGD

**NP_035040.1 neurofilament light polypeptide [Mus musculus]**

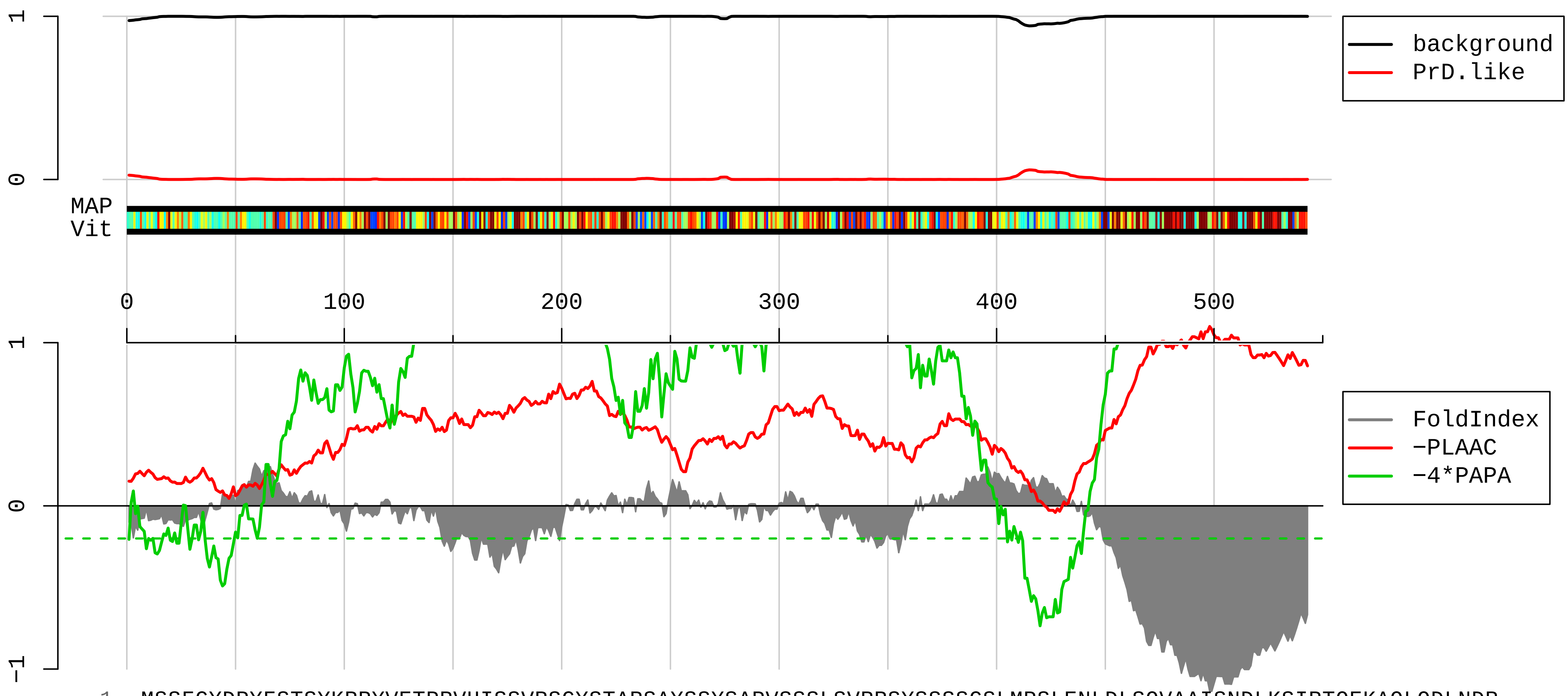


```
  1 MSSFGYDPYFSTSYKRRYVETPRVHISSVRSGYSTARSAYSSYSAPVSSSLSVRRSYSSSSGSLMPSLENLDLSQVAAISNDLKSIRTQEKAQLQDLNDR
101 FASFIERVHELEQQNKVLEAELLVLRQKHSEPSRFRALYEQEIRDLRLAAEDATNEKQALQGEREGLEETLRNLQARYEEEVLSREDAEGRLMEARKGAD
201 EAALARAELEKRIDSLMDEIAFLKKVHEEEIAELQAQIQYAQISVEMDVSSKPDLSAALKDIRAQYEKLAAKNMQNAEEWFKSRFTVLTESAAKNTDAVR
301 AAKDEVSESRRLLKAKTLEIEACRGMNEALEKQLQELEDKQNADISAMQDTINKLENELRSTKSEMARYLKEYQDLLNVKMALDIEIAAYRKLLEGEETR
401 LSFTSVGSITSGYSQSSQVFGRSAYSGLQSSSYLMSARSFPAYYTSHVQEEQTEVEETIEATKAEEAKDEPPSEGEAEEEEKEKEEGEEEEGAEEEEAAK
501 DESEDTKEEEEGGEGEEEDTKESEEEEKKEESAGEEQVAKKKD
```

**EDL40597.1 purine rich element binding protein B [Mus musculus]**

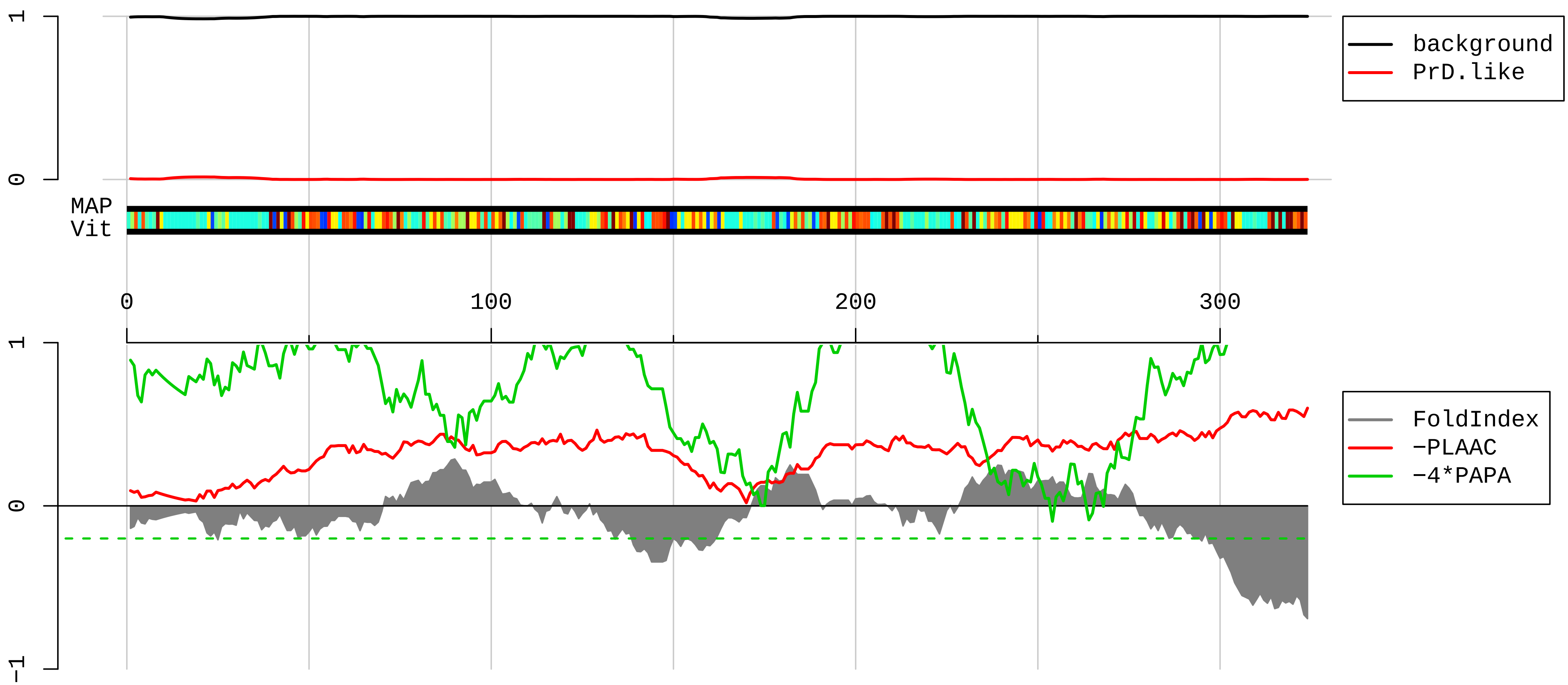


1 MADGDSGSERGGGGGGGGGGPGGFQPAPRGGGGGGGGGPGGEQETQELASKRLDIQNKRFYLDVKQNAKGRFLKIAEVGAGGSKSRLTLSMAVAAEFRDSLG
101 DFIEHYAQLGPSSPEQLAAGAEEGGGPRRALKSEFLVRENRKYYLDLKENQRGRFLRIRQTVNRGGGGFGGGPGPGGLQSGQTIALPAQGLIEFRDALAK
201 LIDDYGGDEDELAGGPGGGAGGPGGGLYGELPEGTSITVDSKRFFFDVGCNKYGVFLRVSEVKPSYRNAITVPFKAWGKFGGAFCRYADEMKEIQERQRD
301 KLYERRGGGSGGGDESEGEEVDED

**AAH07134.1 Ubiquitin specific peptidase 10 [Mus musculus]**

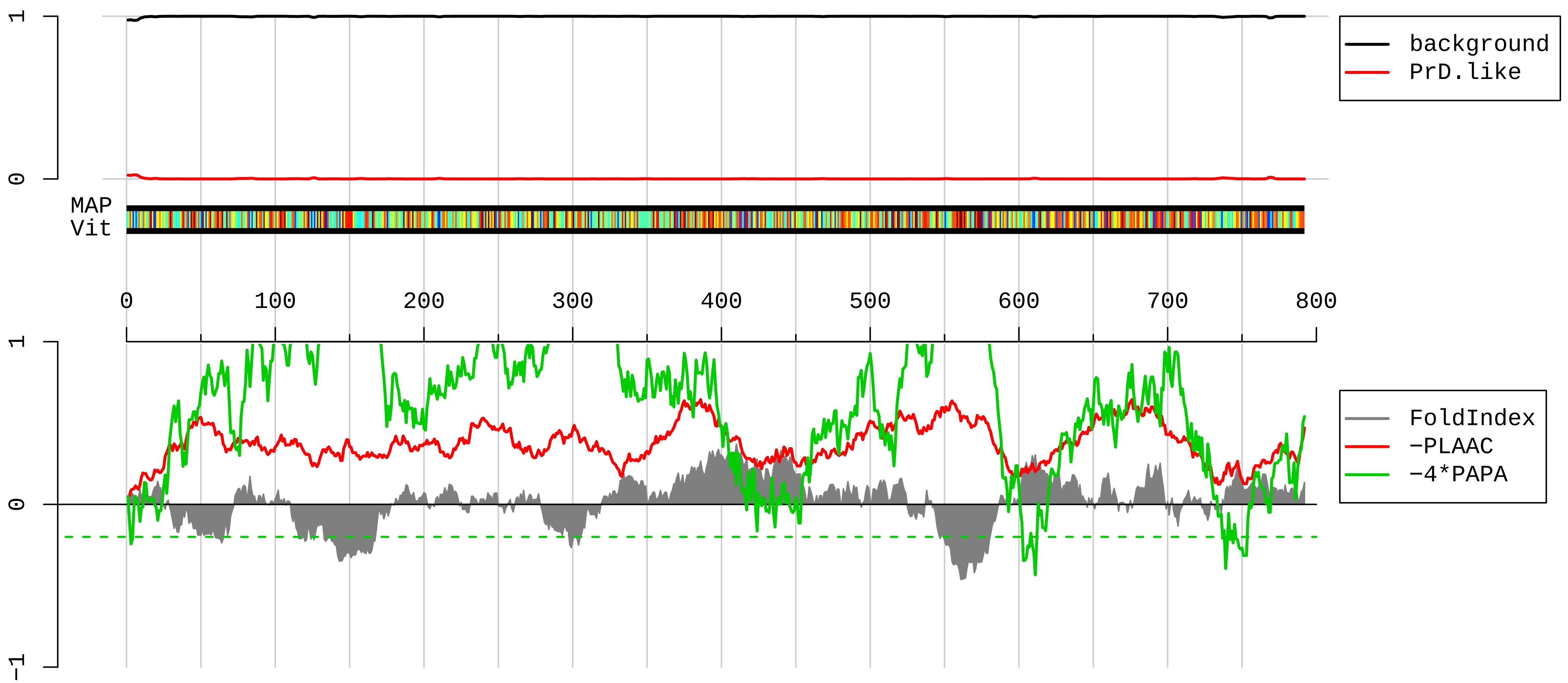


1 MALHNPQYIFGDFSPDEFNQFFVTPRSSVELPPYSGTLCSIQAEDELPDGQEHQRIEFGVDEVIEPSEGLPRTPSYSISSTLNPQAPEFILGCTTSKKIP
101 DAVEKDETYSSIDQYPASALALESNSNAEAETLENDSGAGGLGQRERKKKKKRPPGYYSYLKDGGEDSASPATLVNGHATSVGTSGEAVEDAEFMDVLPP
201 VMPRTCDSPQNPVDFISGPVPDSPFPRTLGGDARTAGLCEGCHEADFEQPCLPADSLLRTAGTQPYVGTDTTENFAVANGKILESPGEDTAASGAELHTD
301 EGADLDPAKPESQSPPAESALSASGAIPISQPAKSWASLFHDSKPSSSSPMAYVETKCSPPVPSPLASEKQMEVKEGLVPVSEDPVAIKIAELLETVTLI
401 HKPVSLQPRGLINKGNWCYINATLQALVACPPMYHLMKFIPLYSKVQRPCTSTPMIDSFVRLMNEFTNMPVPPKPRQALGDKIVRDIRPGAAFEPTYIYR
501 LLTVIKSSLSEKGRQEDAEEYLGFILNGLHEEMLSLKKLLSPTHEKHSVSNGPRSDLIEDEELEDTGKGSEDEWEQVGPKNKTSITRQADFVQTPITGIF
601 GGHIRSVVYQQSSKESATLQLFFTLQLDIQSDKIRTVQDALESLVARESVQGYTTKTKQEVEVSRRVTLEKLPPVLVLHLKRFVYEKTGGCQKLVKNIDY
701 PVDLEISRELLSPGIKNKNFKCQRTYRLFAVVYHHGNSATGGHYTTDVFQIGLNGWLRIDDQTVKVINQYQVVKPPADRTAYLLYYRRVDLL

**BAA25299.1 SRPK1 [Mus musculus]**

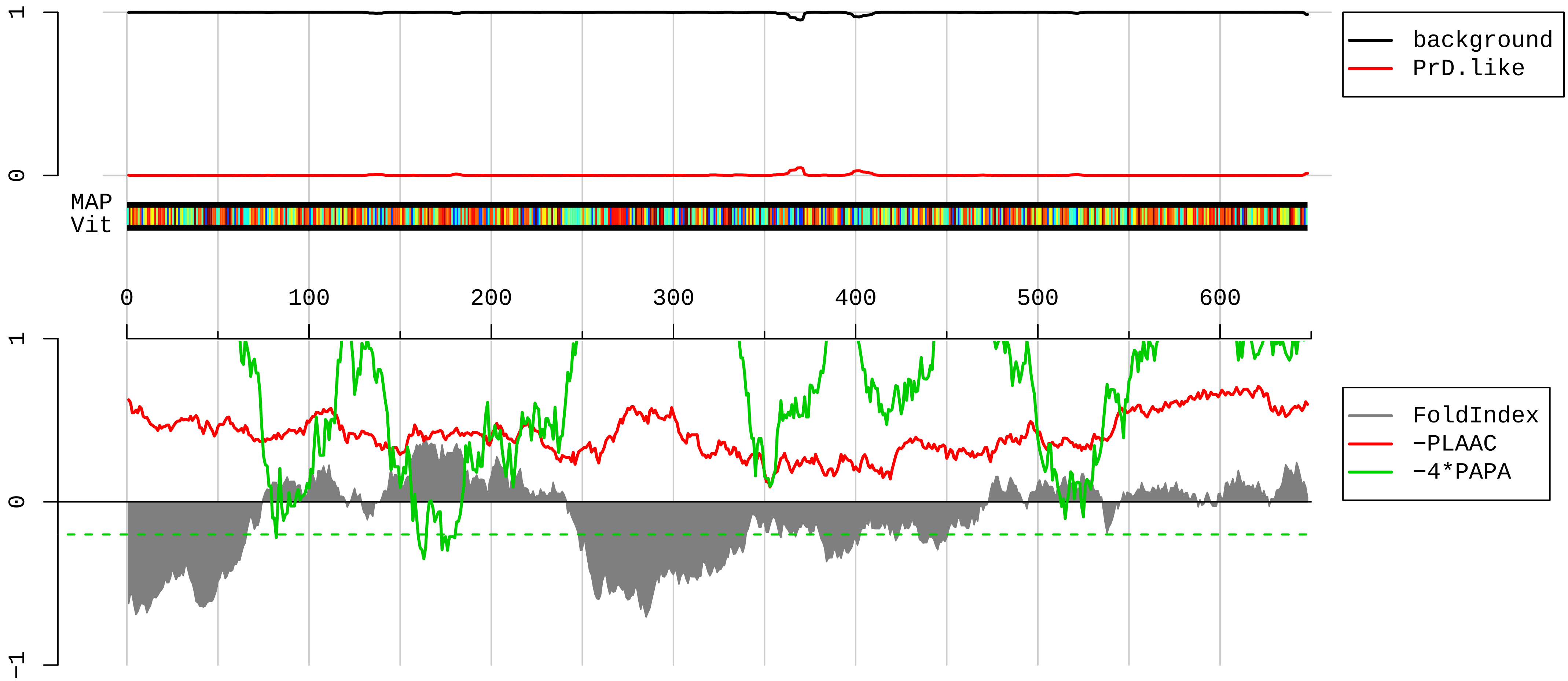


```
1    MERKVLALQARKKRTKAKKDKAQRKPETQHRGSAPHSESDIPEQEEEILGSDDDEQEDPNDYCKGGYHLVKIGDLFNGRYHVIRKLGWGHFSTVWLSWDI
101  QGKKFVAMKVVKSAEHYTETALDEIRLLKSVRNSDPNDPNGEMVVQLLDDFKISGVNGTHICMVFEVLGHHLLKWIIKSNYQGLPLPCVKKIIQQVLQGL
201  DYLHTKCRIIHTDIKPENILLSVNEQYIRRLAAEATEWQRSGAPPPSGSAVSTAPQPKPADKMSKNKKKKLKKKQKRQPELLEKRMQEIEEMEKESGPGQ
301  KRPNKQEESESPVDRPLTENPPNKMTQEKLEESNSIGQDQTLTERGGEGGAPEINCNGVIGVVNYPENSNNETLRHKEDLHNANDCDVHTLKQEPSFLNS
401  SNGDSSPSQDTDSCTPTASETMVCQSSAEQSLTRQDITQLEESIRADTPSGDEQEPNGALDSKGKFSAGNFLINPLEPKNAEKLQVKIADLGNACWVHKH
501  FTEDIQTRQYRSLEVLIGSGYNTPADIWSTACMAFELATGDYLFEPHSGEDYTRDEDHIALIIELLGKVPRKLIVAGKYSKEFFTKKGDLKHITKLKPWG
601  LLEVLVEKYEWPQEEAAGFTDFLLPMLELMPEKRATAAECLRHPWLNS
```

#### AAH54830.1 Actinin, alpha 1 [Mus musculus]

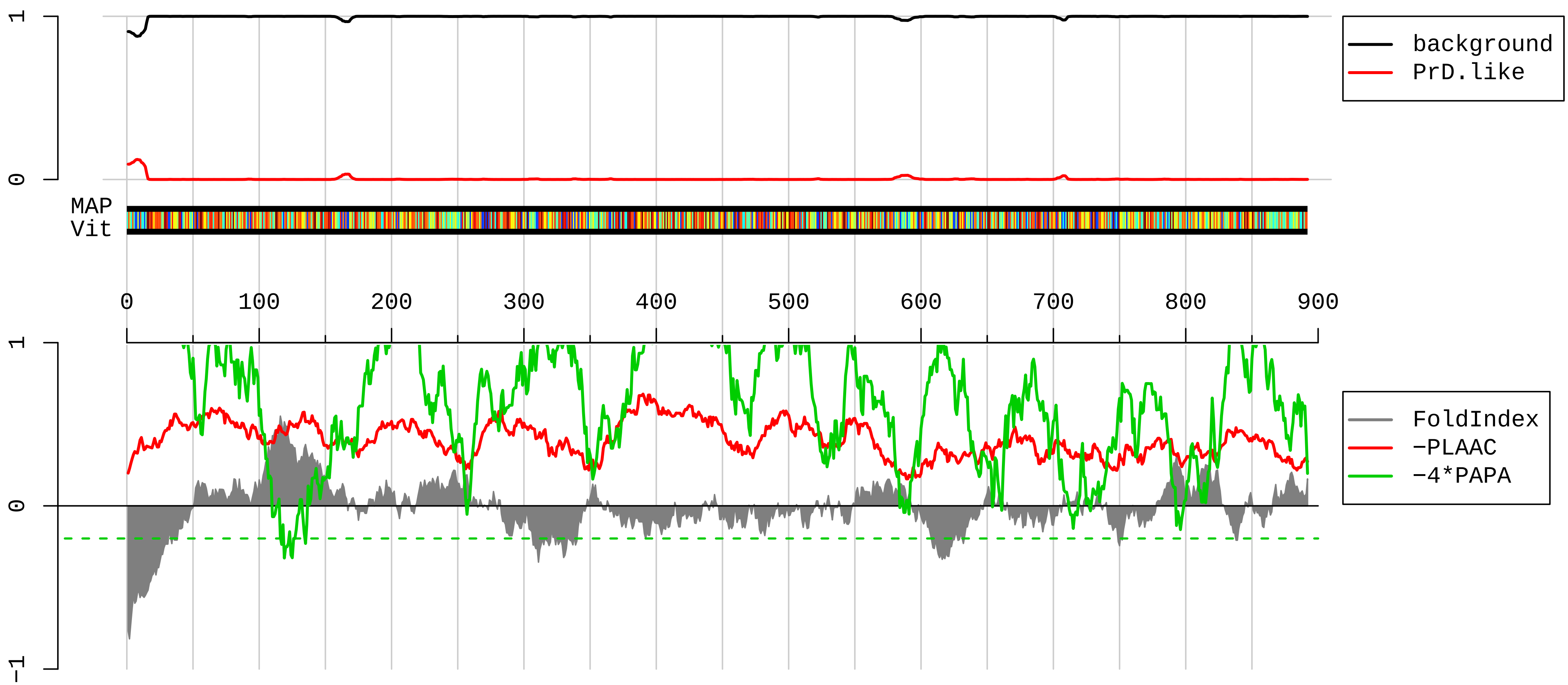


1 MDHYDSQQTNDYMQPEEDWDRDLLLDPAWEKQQRKTFTAWCNSHLRKAGTQIENIEEDFRDGLKLMLLLEVISGERLAKPERGKMRVHKISNVNKALDFI
101 ASKGVKLVSIGAEEIVDGNVKMTLGMIWTIILRFAIQDISVEETSAKEGLLLWCQRKTAPYKNVNIQNFHISWKDGLGFCALIHRHRPELIDYGKLRKDD
201 PLTNLNTAFDVAERFLDIPKMLDAEDIVGTARPDEKAIMTYVSSFYHAFSGAQKAETAANRICKVLAVNQENEQLMEDYEKLASDLLEWIRRTIPWLENR
301 VPENTMHAMQQKLEDFRDYRRLHKPPKVQEKCQLEINFNTLQTKLRLSNRPAFMPSEGRMVSDINNAWGCLEQAEKGYEEWLLNEIRRLERLDHLAEKFR
401 QKASIHEAWTDGKEAMLRQKDYETATLSEIKALLKKHEAFESDLAAHQDRVEQIAAIAQELNELDYYDSPSVNARCQKICDQWDNLGALTQKRREALERT
501 EKLLETIDQLYLEYAKRAAPFNNWMEGAMEDLQDTFIVHTIEEIQGLTTAHEQFKATLPDADKERLAILGIHNEVSKIVQTYHVNMAGTNPYTTITPQEI
601 NGKWDHVRQLVPRRDQALTEEHARQQHNERLRKQFGAQANVIGPWIQTKMEEIGRISIEMHGTLEDQLSHLRQYEKSIVNYKPKIDQLECDHQLIQEALI
701 FDNKHTNYNMEHIRVGWEQLLTTIARTINEVENQILTRDAKGISQEQMNEFRASFNHFDRDHSGTLGPEEFKACLISLGYDIGNDPQGEAEFARIMSIVD
801 PNRLGVVTFQAFIDFMSRETADTDTADQVMASFKILAGDKNYITEDELRRELPPDQAEYCIARMAPYAGPDSVPGALDYMSFSTALYGESDL

**AAH86896.1 Ribosomal protein L13A [Mus musculus]**

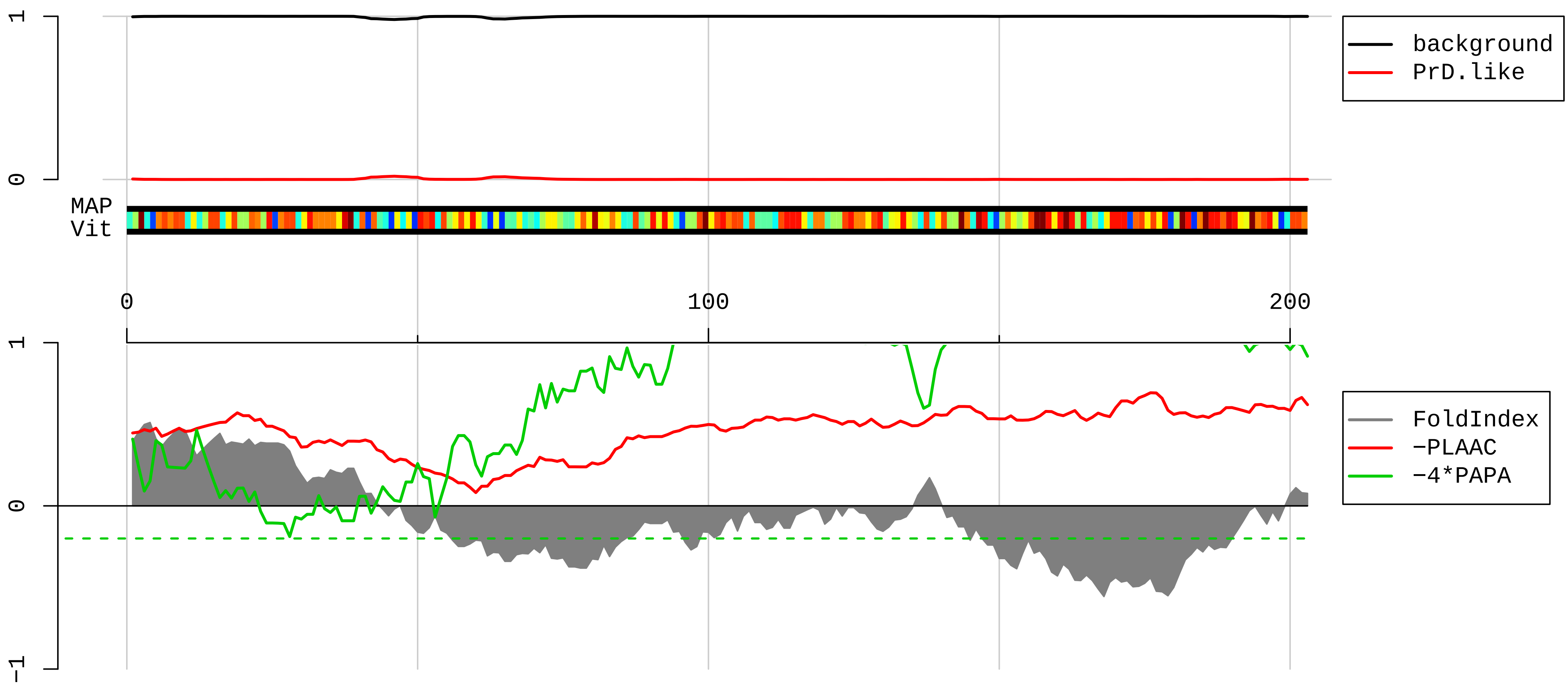


1 MAEGQVLVLDGRGHLLGRLAAIVAKQVLLGRKVVVVRCEGINISGNFYRNKLKYLAFLRKRMNTNPSRGPYHFRAPSRIFWRTVRGMLPHKTKRGQAALE
101 RLKVLDGIPPPYDKKKRMVVPAALKVVRLKPTRKFAYLGRLAHEVGWKYQAVTATLEEKRKEKAKMHYRKKKQILRLRKQAEKNVEKKICKFTEVLKTNG
201 LLV

**NP_075891.1 myosin regulatory light chain 12B [Mus musculus]**

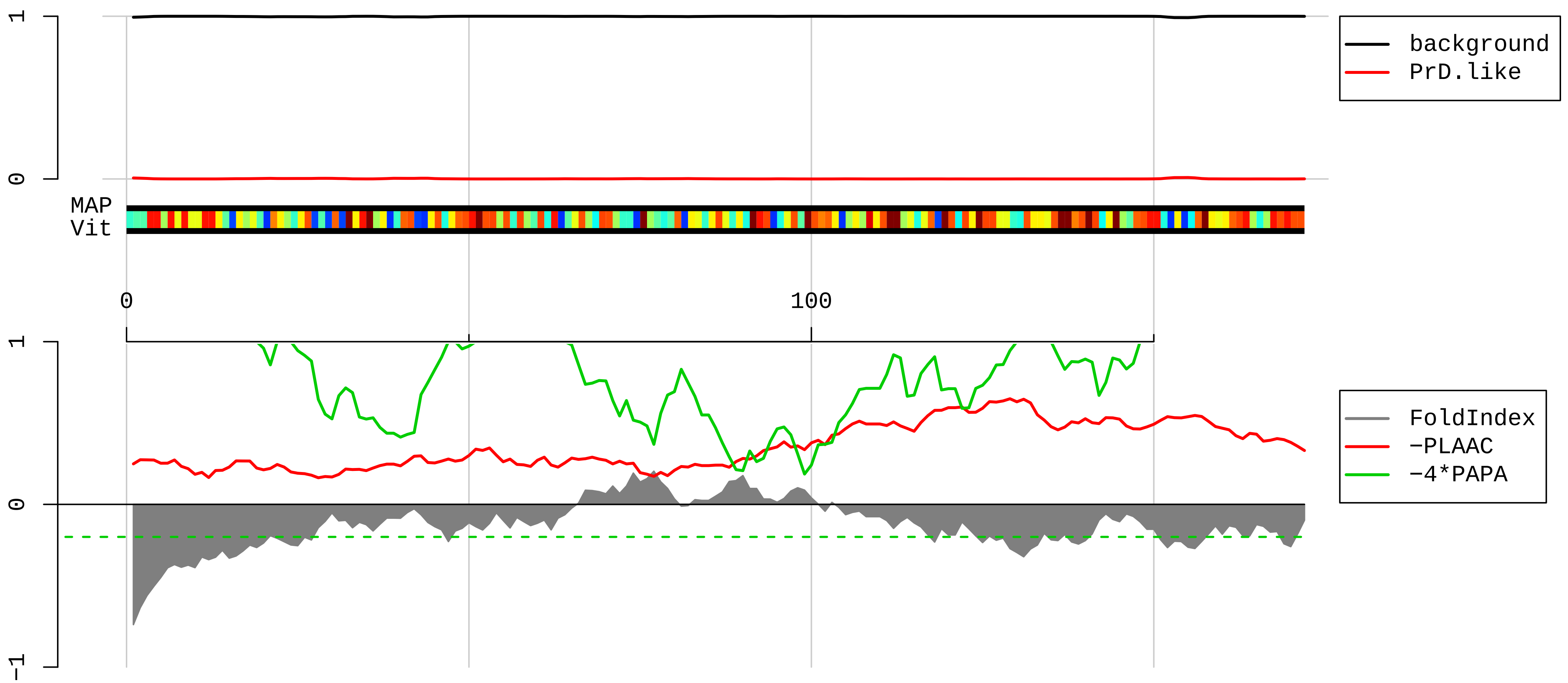


1 MSSKKAKTKTTKKRPQRATSNVFAMFDQSQIQEFKEAFNMIDQNRDGFIDKEDLHDMLASLGKNPTDAYLDAMMNEAPGPINFTMFLTMFGEKLNGTDPE
101 DVIRNAFACFDEEATGTIQEDYLRELLTTMGDRFTDEEVDELYREAPIDKKGNFNYIEFTRILKHGAKDKDD

**AAH30502.1 Serbp1 protein [Mus musculus]**

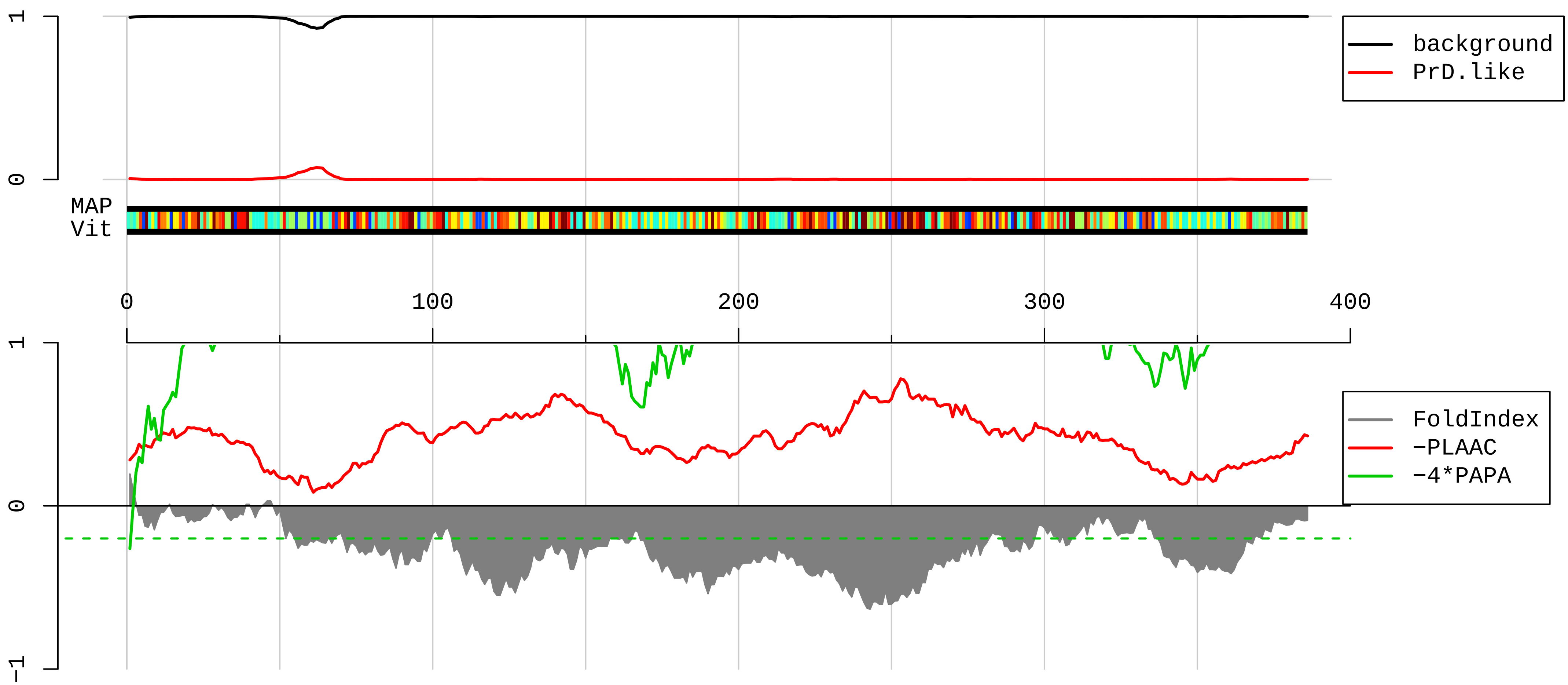


```
  1 MPGHLQEGFGCVVTNRFDQLFDDESDPFEVLKAAENKKKEAGGGGVGGPGAKSAAQAAAQTNSNAAGKQLRKESQKDRKNPLPPSVGVADKKEETQPPVA
101 LKKEGIRRVGRRPDQQLQGDGKLIDRRAERRPPRERRFEKPLEEKGEGGEFSVDRPIIERPIRGRGGLGRGRGGRGRGMGRGDGFDSRGKREFDRHSGSD
201 RSGLKHEDKRGGSGSHNWGTVKDELTDLDQSNVTEETPEGEEHPVADTENKENEVEEVKEEGPKEMTLDEWKAIQNKDRAKVEFNIRKPNEGADGQWKKG
301 FVLHKSKSEEAHAEDSVMDHHFRKPANDITSQLEINFGDLGRPGRGGRGGRGGRGRGGRPNRGSRTDKSSASAPDVDDPEAFPALA
```

**AAF65683.1 ribosomal protein S19 [Mus musculus]**

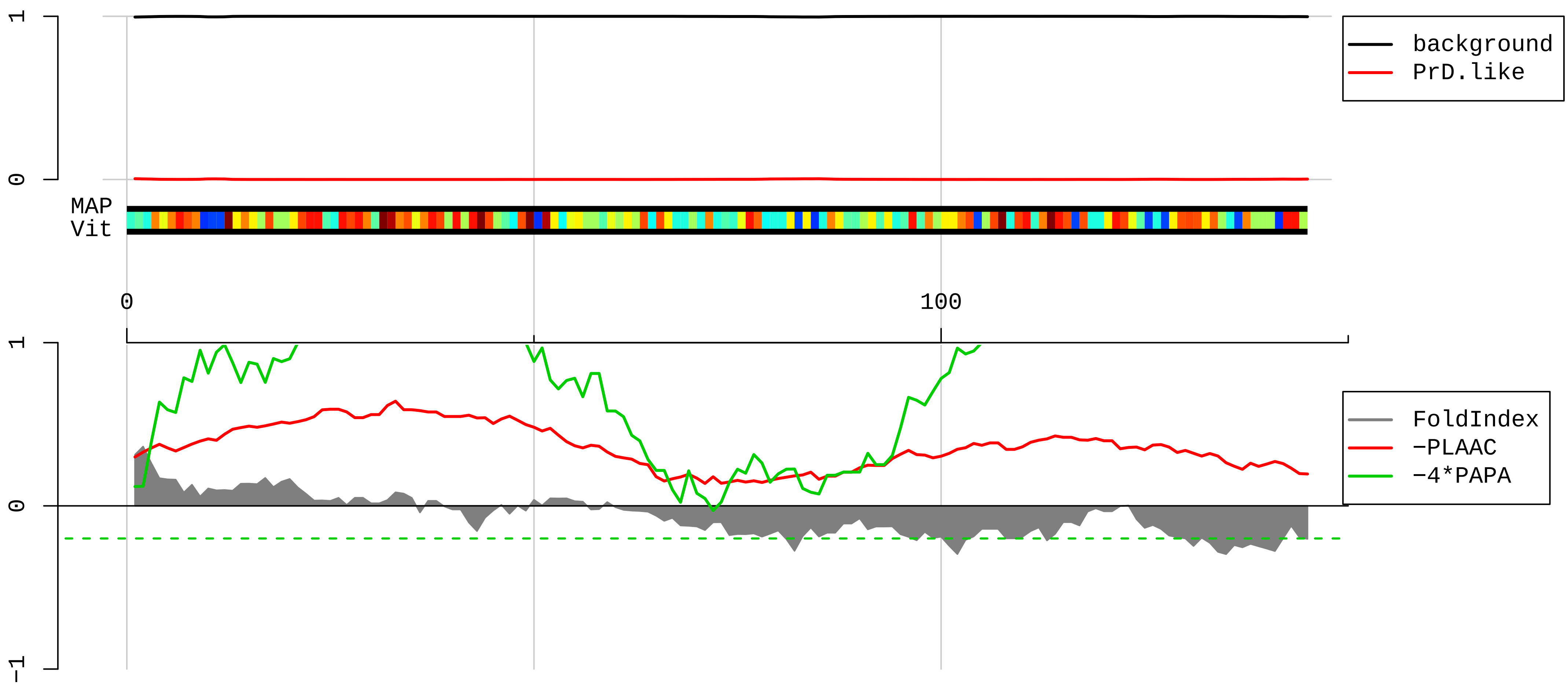


1 MPGVTVKDVNQQEFVRALAAFLKKSGKLKVPEWVDTVKLAKHKELAPYDENWFYTRAASTARHLYLRGGAGVGSMTKIYGGRQRNGVRPSHFSRGSKSVA

101 RRVLQALEGLKMVEKDQDGGRKLTPQGQRDLDRIAGQVAAANKKH

AAH06714.1 Drebrin 1 [Mus musculus]

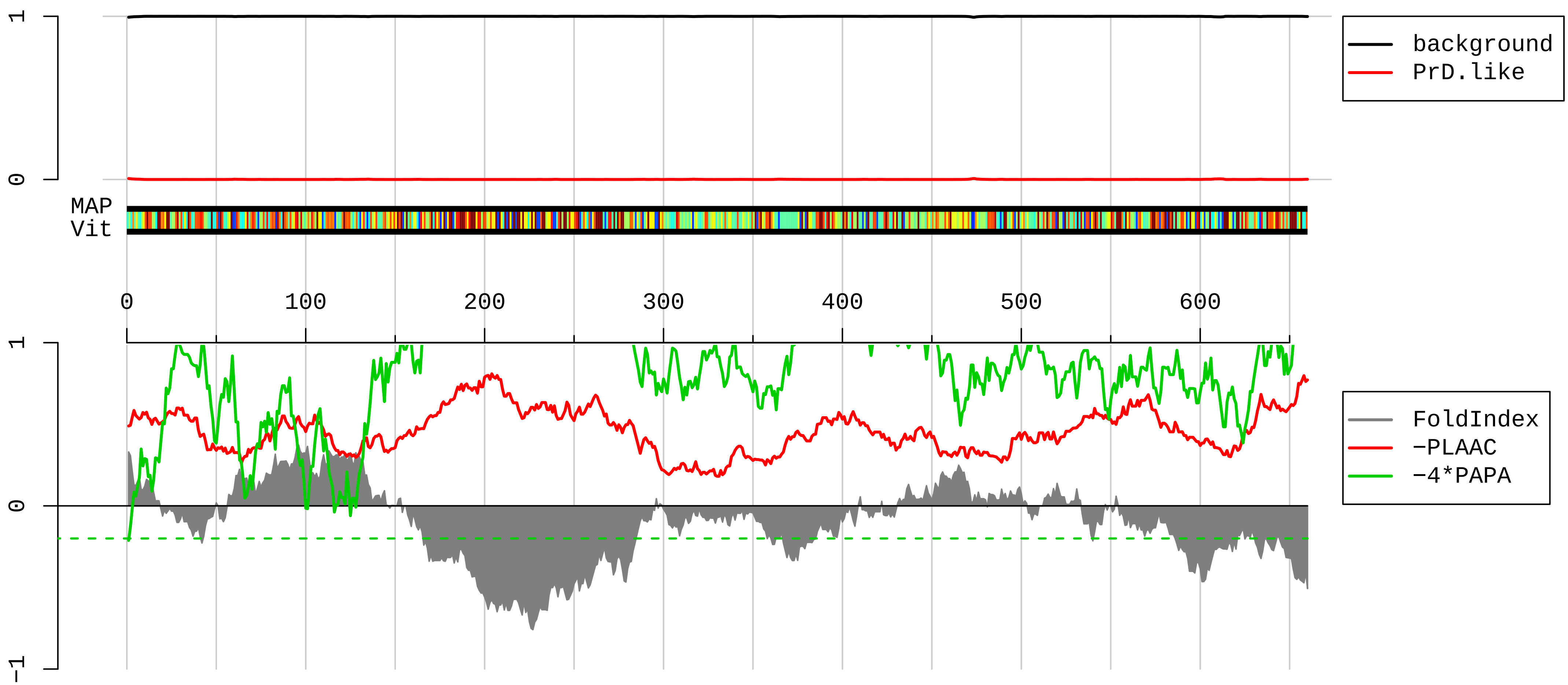


1 MAGVSFSGHRLELLAAYEEVIREESAADWALYTYEDGSDDLKLAASGEGGLQELSGHFENQKVMYGFCSVKDSQAALPKYVLINWVGEDVPDARKCACAS
101 HVAKVAEFFQGVDVIVNASSVEDIDAGAIGQRLSNGLARLSSPVLHRLRLREDENAEPVGTTYQKTDAAVEMKRINREQFWEQAKKEEELRKEEERKKAL
201 DARLRFEQERMEQERQEQEERERRYREREQQIEEHRRKQQSLEAEEAKRRLKEQSIFGDQRDEEEEESQMKKSESEVEEAAAIIAQRPDNPREFFRQQERV
301 ASASGGSCDAPAPAPFNHRPGSHLDSHRRMAPTPIPTRSPSDSSTASTPIAEQIERALDEVTSSQPPPPPPPPPPTQEAQETTPSLDEELSKEAKVTAAP
401 EVWAGCAAEPPQAQEPPLLQSSPLEDSMCTESPEQAALAAPAEPAASVTSVADVHAADTIETTTATTDTTIANNVTPAAASLIDLWPGNGEEASTLQAEP
501 RVPTPPSGAEASLAEVPLLNEAAQEPLPPVGEGCANLLNFDELPEPPATFCDPEEEVGETLAASQVLTMPSALEEVDQVLEQELEPEPHLLTNGETTQKE
601 GTQASEGYFSQSQEEEFAQSEEPCAKVPPPVFYNKPPEIDITCWDADPVPEEEEGFEGGD

**AAI31679.1 Dst protein, partial [Mus musculus]**

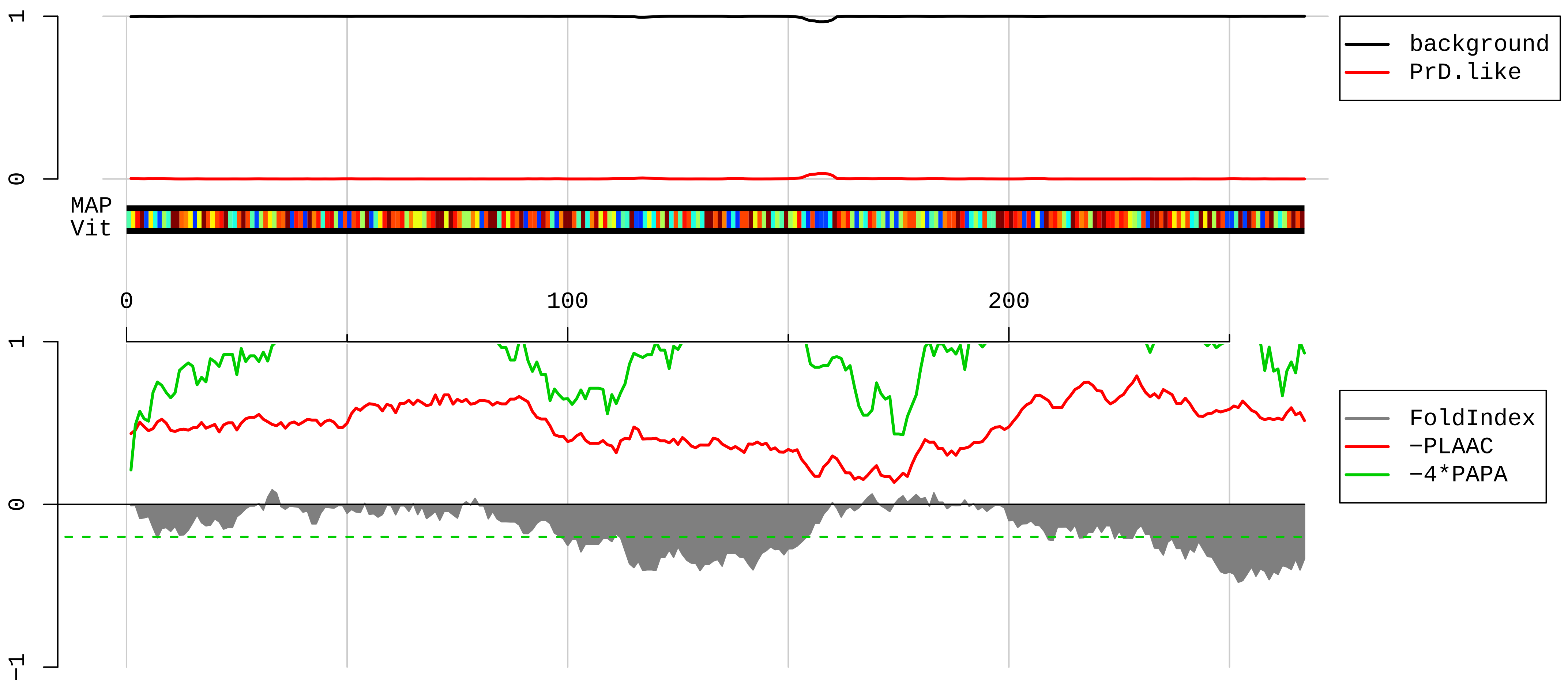


1 SFKEQRGQAMEEIVRNTELFLKESGDELSQADRALIEQKLNEVKMKCAQLNLKAEQSRKELDKAVTTALKEETEKVAAVRQLEESKTKIENLLNWLSNVE

101 EDSEGVWTKHTQPMEQNGTYLHEGDSKLGAGEEDEVNGNLLETDAEGHSEATKGNLNQQYEKVKAQHGKIMAQHQAVLLATQSAQVLLEKQGHYLSPEEK

201 EKLQKNTQELKVHYEKVLAECEKKVKLTHSLQEELEKFDTDYSEFEHWLQQSEQELANLEAGADELE

**AAH11311.1 Protein kinase, interferon inducible double stranded RNA dependent activator [Mus musculus]**

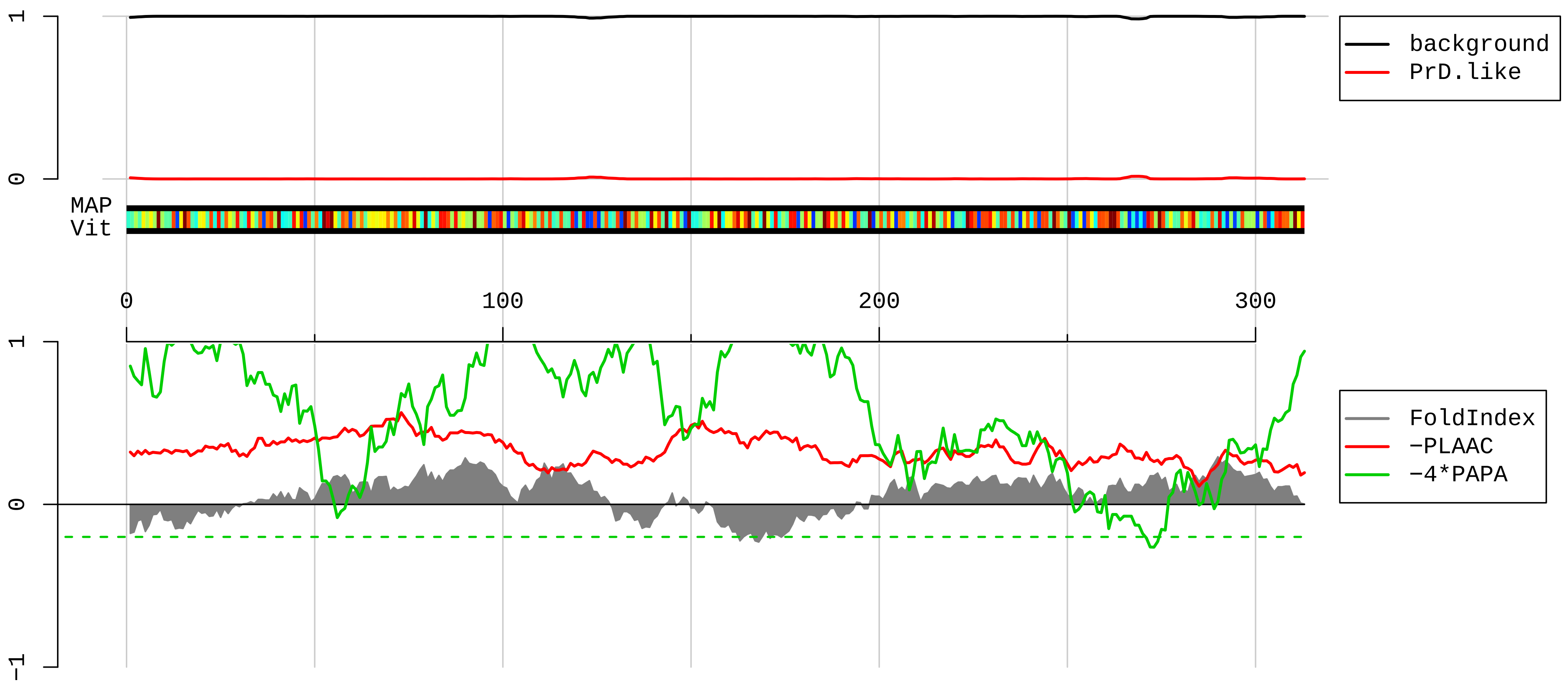


1 MSHSRHRAEAPPLQREDSGTFSLGKMITAKPGKTPIQVLHEYGMKTKNIPVYECERSDVQVHVPTFTFRVTVGDITCTGEGTSKKLAKHRAAEAAINILK
101 ANASICFAVPDPLMPDPSKQPKNQLNPIGSLQELAIHHGWRLPEYTLSQEGGPAHKREYTTICRLESFMETGKGASKKQAKRNAAEKFLAKFSNISPENH
201 ISLTNVVGHSLGCTWHSLRNSPGEKINLLKRSLLSLPNTDYIQLLSEIASEQGFNITYLDIEELSANGQYQCLAELSTSPITVCHGSGISCGNAQSDAAH
301 NALQYLKIIAERK

**AAH23767.1 Eif3b protein [Mus musculus]**

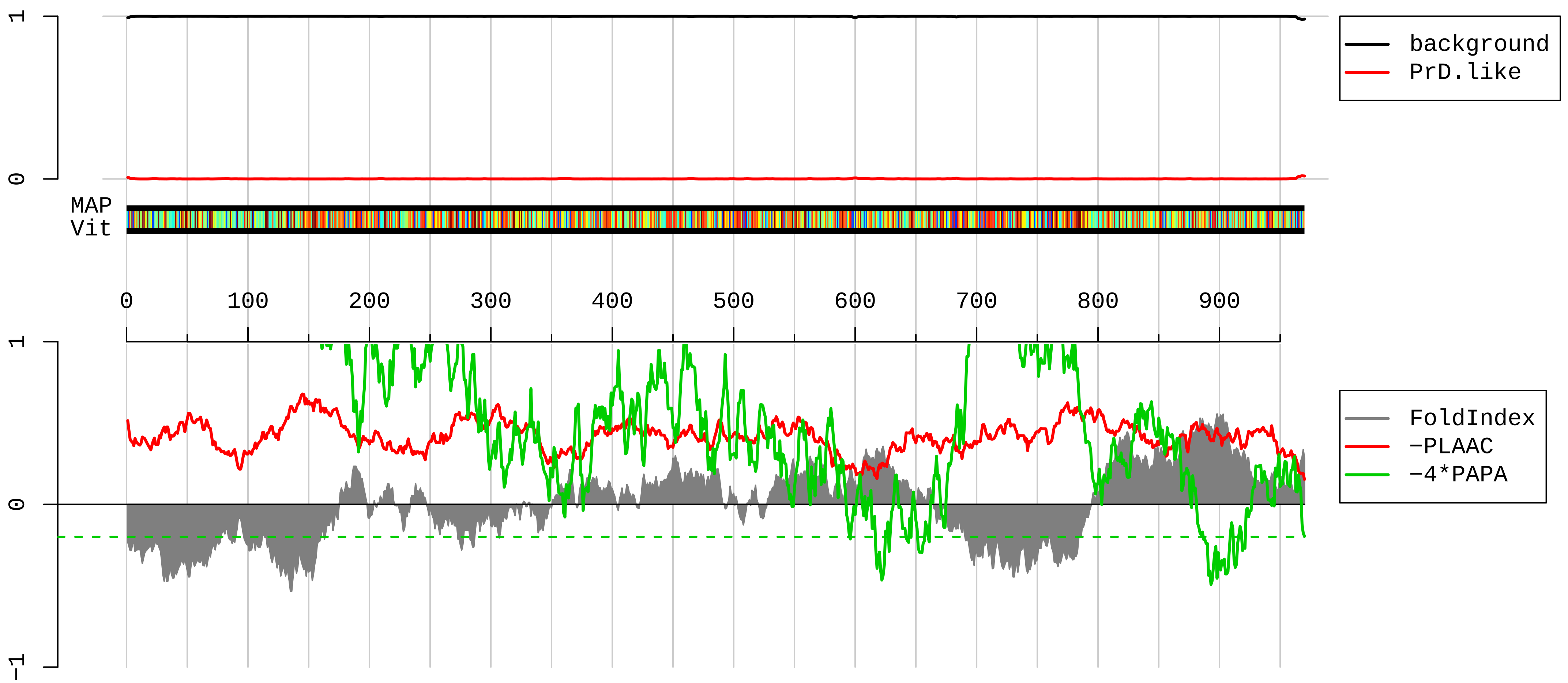


1 MQDAENVAVPEAAEERAEPARQQPASESPPTDEAAGSGGSEVGQTEDAEEDAEAGPEPEVRAKPAAQSEEETATSPAASPTPQSAERSPSQEPSAPGKAE
101 AVGEQARGHPSAGAEEEGGSDGSAAEAEPRALENGEADEPSFSDPEDFVDDVSEEELLGDVLKDRPQEADGIDSVIVVDNVPQVGPDRLEKLKNVIHKIF
201 SKFGKIINDYYPEEDGKTKGYIFLEYASPAHAVDAVKNADGYKLDKQHTFRVNLFTDFDKYMTISDEWDIPEKQPFKDLGNLRYWLEEAECRDQYSVIFE
301 SGDRTSIFWNDVKDPVSIEERARWTETYVRWSPKGTYLATFHQRGIALWGGDKFKQIQRFSHQGVQLIDFSPCERYLVTFSPLMDTQDDPQAIIIWDILT
401 GHKKRGFHCESSAHWPIFKWSHDGKFFARMTLDTLSIYETPSMGLLDKKSLKISGIKDFSWSPGGNIIAFWVPEDKDIPARVTLMQLPTRQEIRVRNLFN
501 VVDCKLHWQKNGDYLCVKVDRTPKGTQGVVTNFEIFRMREKQVPVDVVEMKETIIAFAWEPNGSKFAVLHGEAPRISVSFYHVKSNGKIELIKMFDKQQA
601 NTIFWSPQGQFVVLAGLRSMNGALAFVDTSDCTVMNIAEHYMASDVEWDPTGRYVVTSVSWWSHKVDNAYWLWTFQGRLLQKNNKDRFCQLLWRPRPPTL
701 LSQDQIKQIKKDLKKYSKIFEQKDRLSQSKASKELVERRRTMMEDFRQYRKMAQELYMKQKNERLELRGGVDTDELDSNVDDWEEETIEFFCHRGSHSSG
801 KSGVISALCVTVPCPGSEATLHGEPRAGLLPVLSCSGLRVHPGFFYPDTALCLQPQVHSGDTRASALTLPCLGFYQPCLHWSLTAVFLVAPLNSCCCPWG
901 FQSLHLQLSGVSIHGLAVPVGQCRVAWRPSGVCTPTTYLCQKLWRHPHLPPVQVVAASVQVGCGNKSNPV

**EDL41393.1 PRP19/PSO4 pre−mRNA processing factor 19 homolog (S. cerevisiae), isoform CRA_c [Mus musculus]**

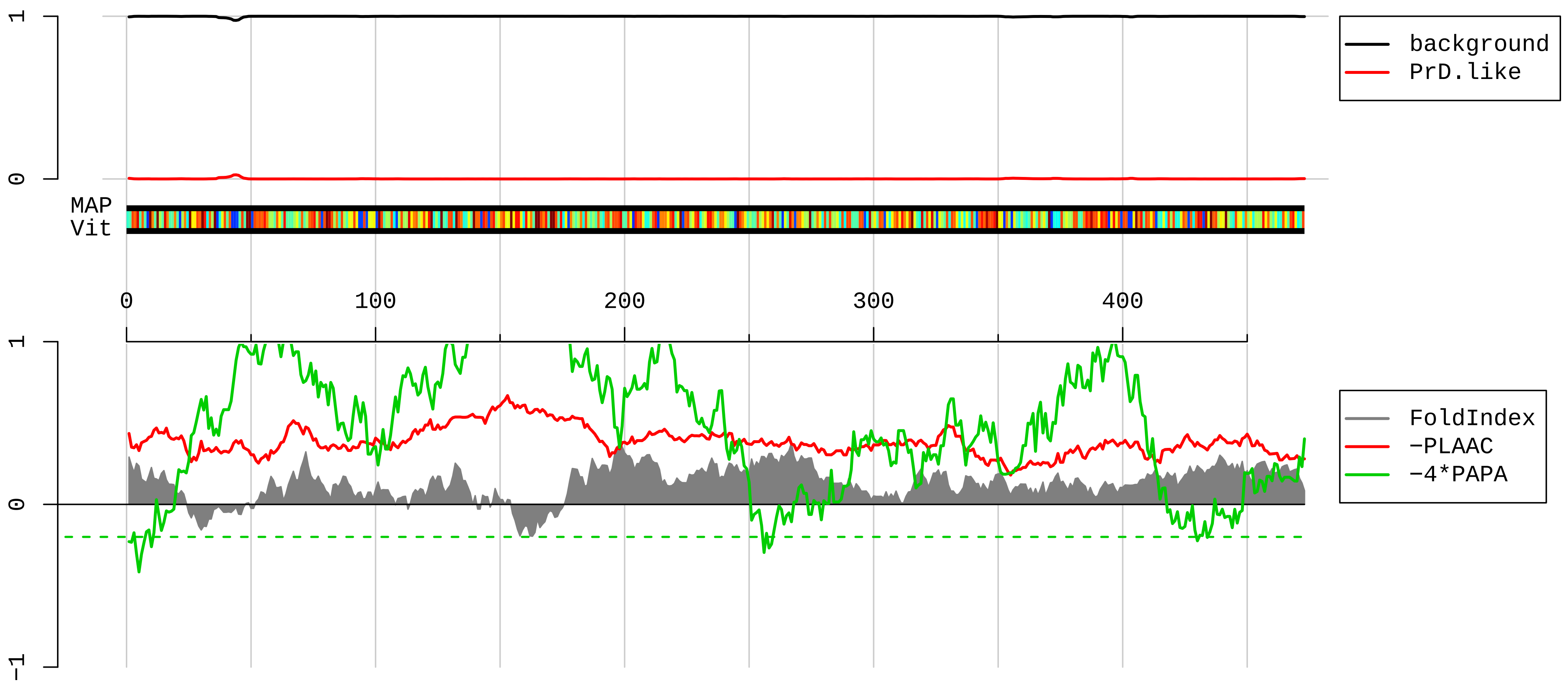


1 MSLICSISNEVPEHPCVSPVSNHVYERRLIEKYIAENGTDPINNQPLSEEQLIDIKVAHPIRPKPPSATSIPAILKALQDEWDAVMLHSFTLRQQLQTTR
101 QELSHALYQHDAACRVIARLTKEGAGEPMDLGELVGMTPEIIQKLQDKATVLTTERKKRGKTVPEELVKPEELSKYRQVASHVGLHSASIPGILALDLCP
201 SDTNKILTGGADKNVVVFDKSTEQILATLKGHTKKVTSVVFHPSQELVFSASPDATIRIWSVPNTSCVQVVRAHESAVTGLSLHATGDYLLSSSDDQYWA
301 FSDIQTGRVLTKVTDETSGCSLTCAQFHPDGLIFGTGTMDSQIKIWDLKERTNVANFPGHSGPITSIAFSENGYYLATAADDSSVKLWDLRKLKNFKTLQ
401 LDNNFEVKSLIFDQSGTYLALGGTDVQIYICKQWTEILHFTEHSGLTTGVAFGHHAKFIASTGMDRSLKFYSL

#### AAH58690.1 Ribosomal protein S5 [Mus musculus]

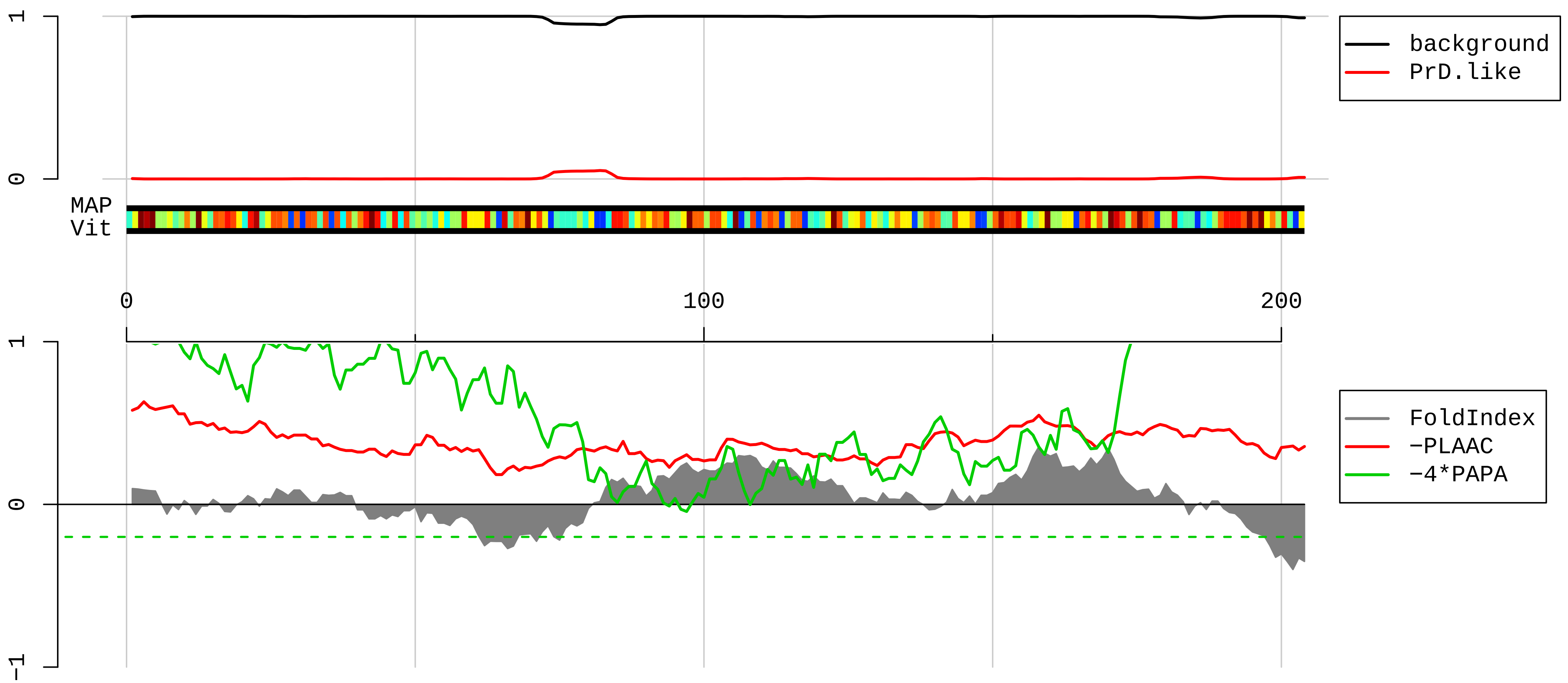


1 MTEWEAATPAVAETPDIKLFGKWSTDDVQINDISLQDYIAVKEKYAKYLPHSAGRYAAKRFRKAQCPIVERLTNSMMMHGRNNGKKLMTVRIVKHAFEII
101 HLLTGENPLQVLVNAIINSGPREDSTRIGRAGTVRRQAVDVSPLRRVNQAIWLLCTGAREAAFRNIKTIAECLADELINAAKGSSNSYAIKKKDELERVA
201 KSNR

**AAI06117.1 Ribosomal protein L21 [Mus musculus]**

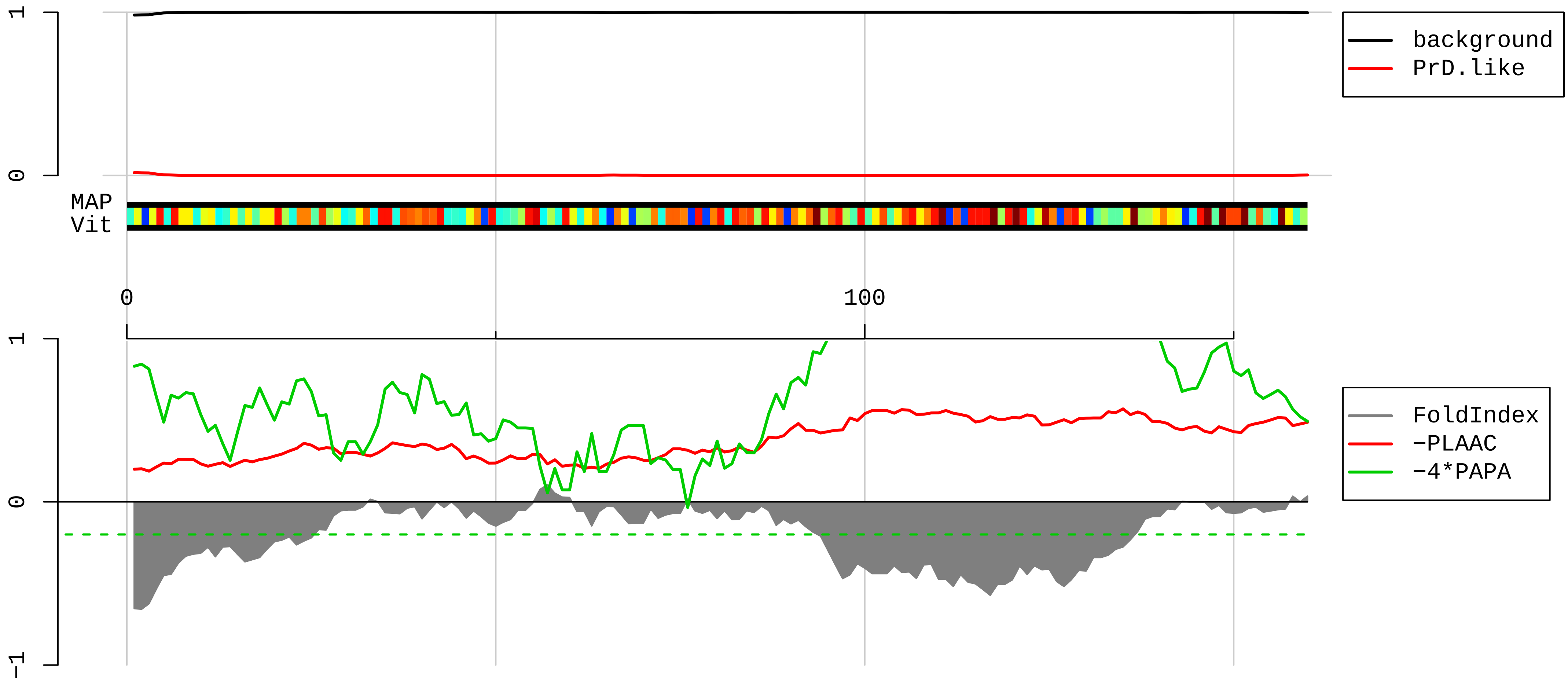


1 MTNTKGKRRGTRYMFSRPFRKHGVVPLATYMRIYKKGDIVDIKGMGTVQKGMPHKCYHGKTGRVYNVTQHAVGIIVNKQVKGKILAKRINVRIEHIKHSK
101 SRDSFLKRVKENDQKKKEAKEKGTWVQLKRQPAPPREAHFVRTNGKEPELLEPIPYEFMA

**AAH43017.1 Ribosomal protein L8 [Mus musculus]**

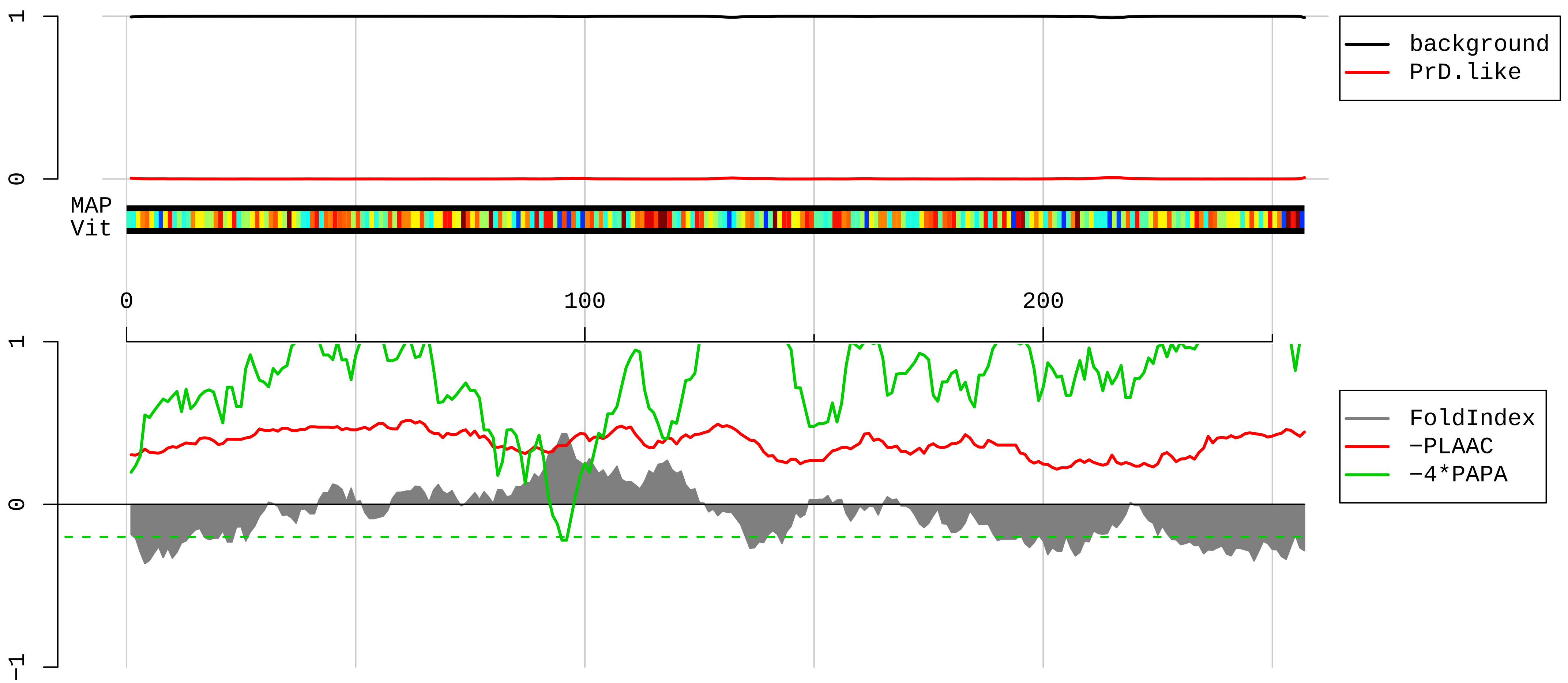


1 MGRVIRGQRKGAGSVFRAHVKHRKGAARLRAVDFAERHGYIKGIVKDIIHDPGRGAPLAKVVFRDPYRFKKRTELFIAAEGIHTGQFVYCGKKAQLNIGN
101 VLPVGTMPEGTIVCCLEEKPGDRGKLARASGNYATVISHNPETKKTRVKLPSGSKKVISSANRAVVGVVAGGGRIDKPILKAGRAYHKYKAKRNCWPRVR
201 GVAMNPVEHPFGGGNHQHIGKPSTIRRDAPAGRKVGLIAARRTGRLRGTKTVQEKEN

**AAH99481.1 Ribosomal protein L26 [Mus musculus]**

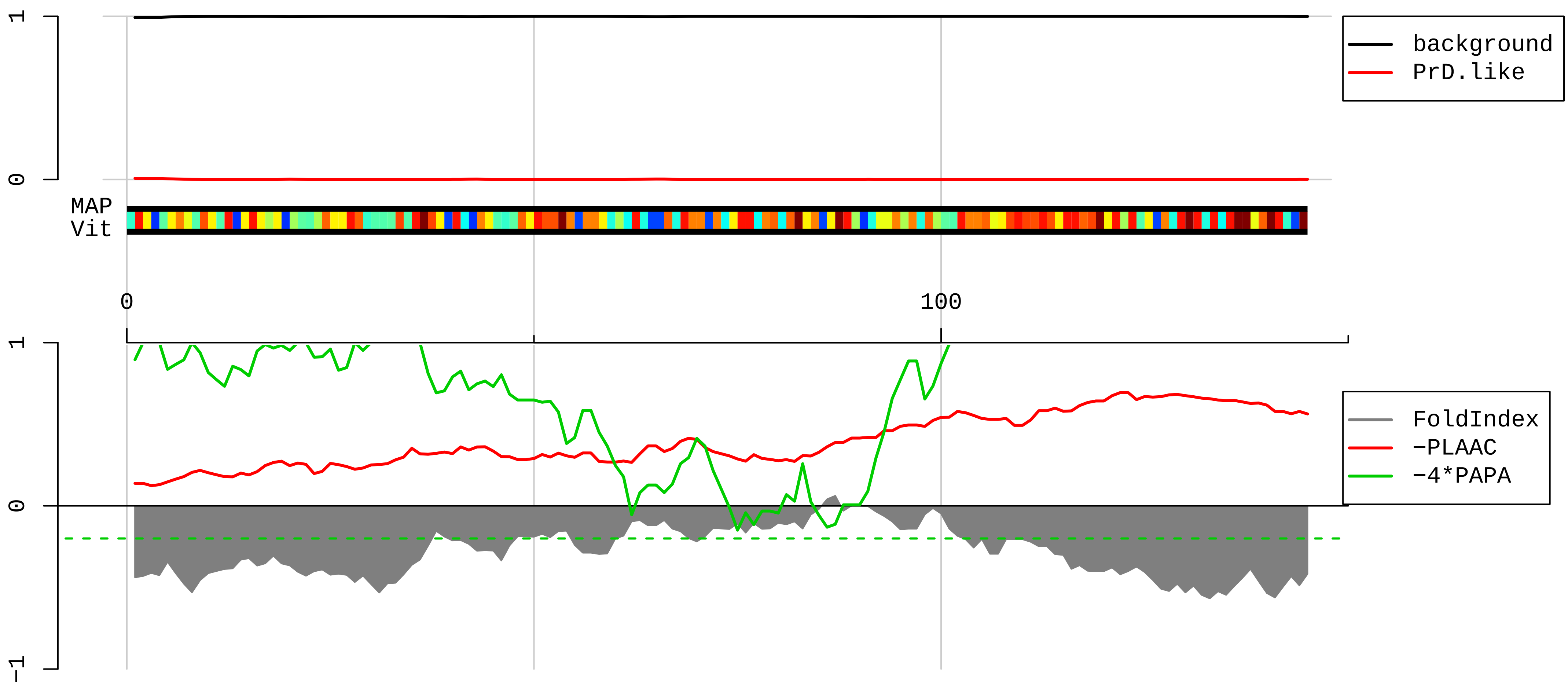


1 MKFNPFVTSDRSKNRKRHFNAPSHIRRKIMSSPLSKELRQKYNVRSMPIRKDDEVQVVRGHYKGQQIGKVVQVYRKKYVIYIERVQREKANGTTVHVGIH

101 PSKVVITRLKLDKDRKKILERKAKSRQVGKEKGKYKEETIEKMQE

**AAI17559.1 Ribosomal protein S2 [Mus musculus]**

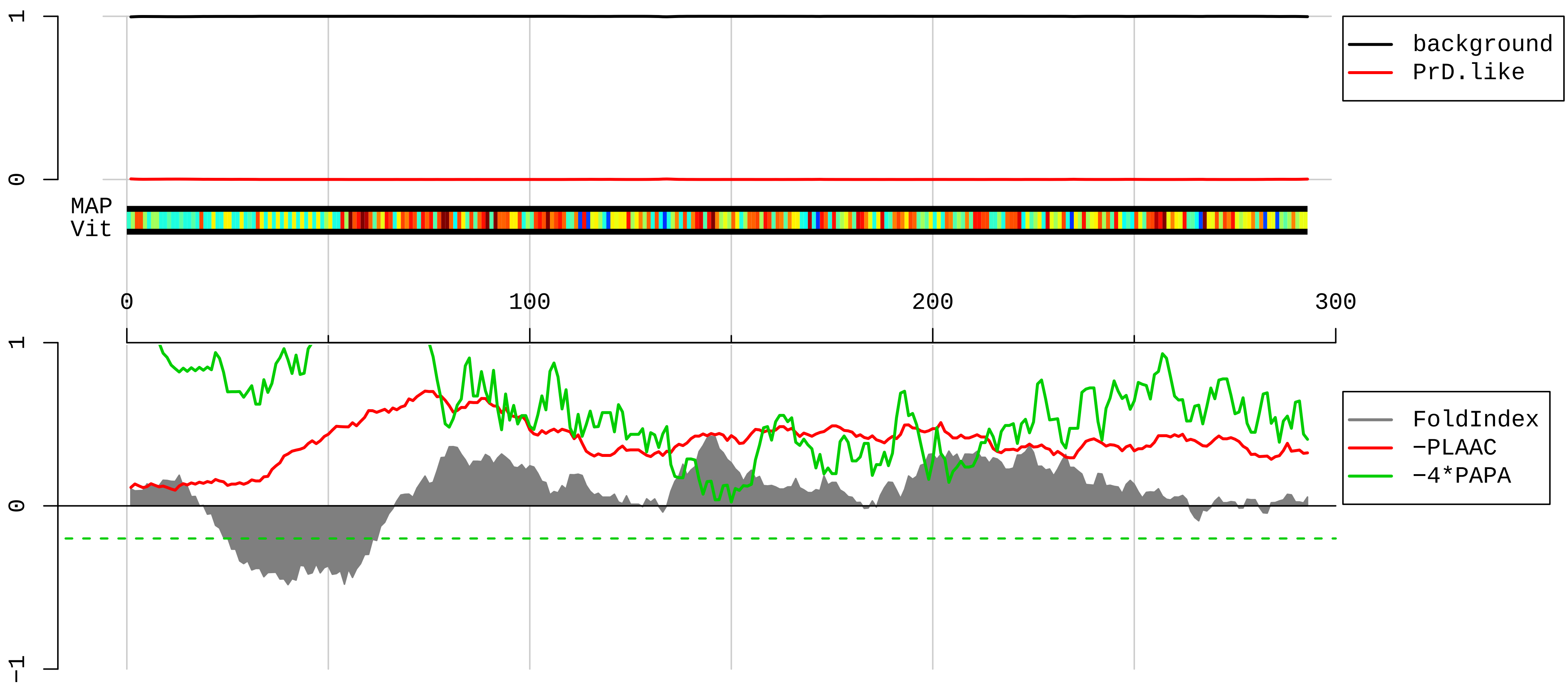


```
  1  MADDAGAAGGPGGPGGPGLGGRGGFRGGFGSGLRGRGRGRGRGRGRGRGARGGKAEDKEWIPVTKLGRLVKDMKIKSLEEIYLFSLPIKESEIIDFFLGA
101  SLKDEVLKIMPVQKQTRAGQRTRFKAFVAIGDYNGHVGLGVKCSKEVATAIRGAIILAKLSIVPVRRGYWGNKIGKPHTVPCKVTGRCGSVLVRLIPAPR
201  GTGIVSAPVPKKLLMMAGIDDCYTSARGCTATLGNFAKATFDAISKTYSYLTPDLWKETVFTKSPYQEFTDHLVKTHTRVSVQRTQAPAVATT
```

**AAH82284.1 Ribosomal protein L27 [Mus musculus]**

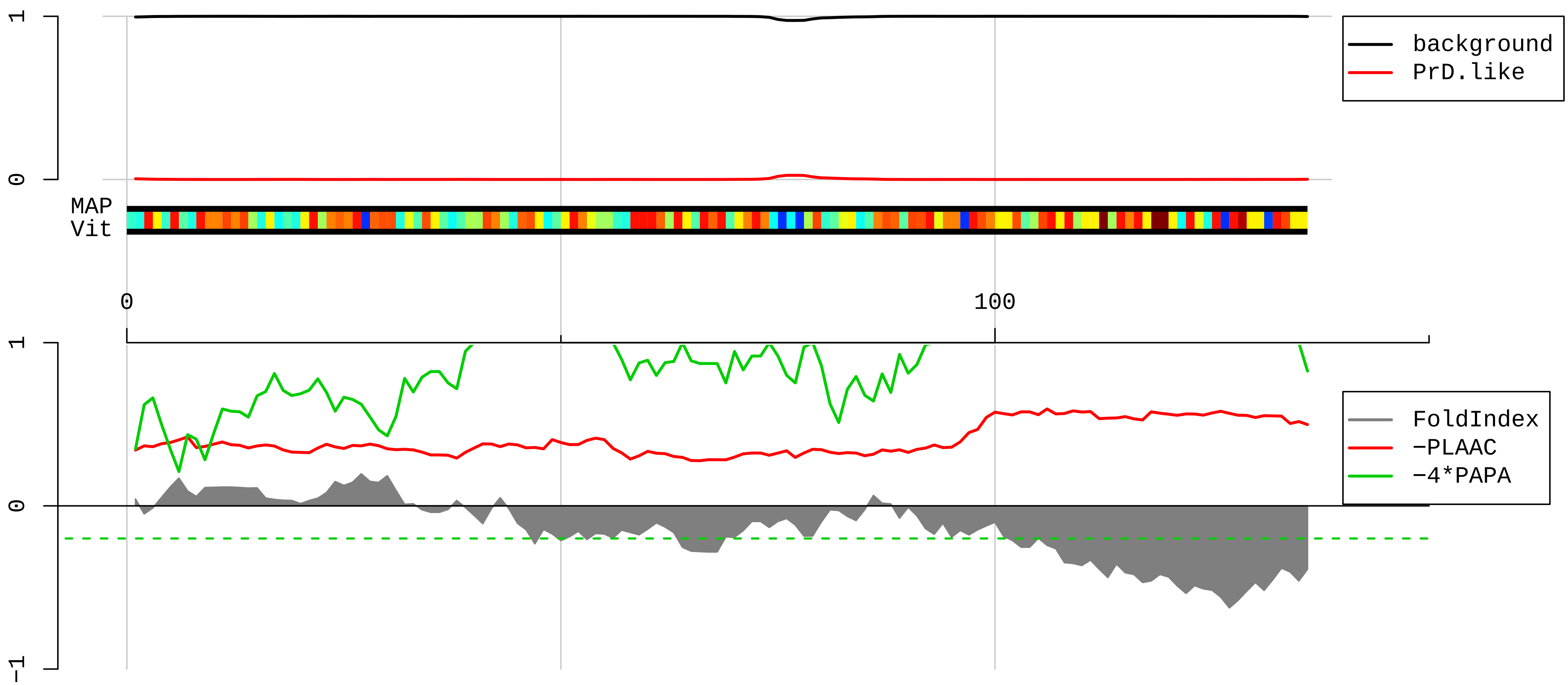


1 MGKFMKPGKVVLVLAGRYSGRKAVIVKNIDDGTSDRPYSHALVAGIDRYPRKVTAAMGKKKIAKRSKIKSFVKVYNYNHLMPTRYSVDIPLDKTVVNKDV
101 FRDPALKRKARREAKVKFEERYKTGKNKWFFQKLRF

**AAH69896.1 Ribosomal protein L11 [Mus musculus]**

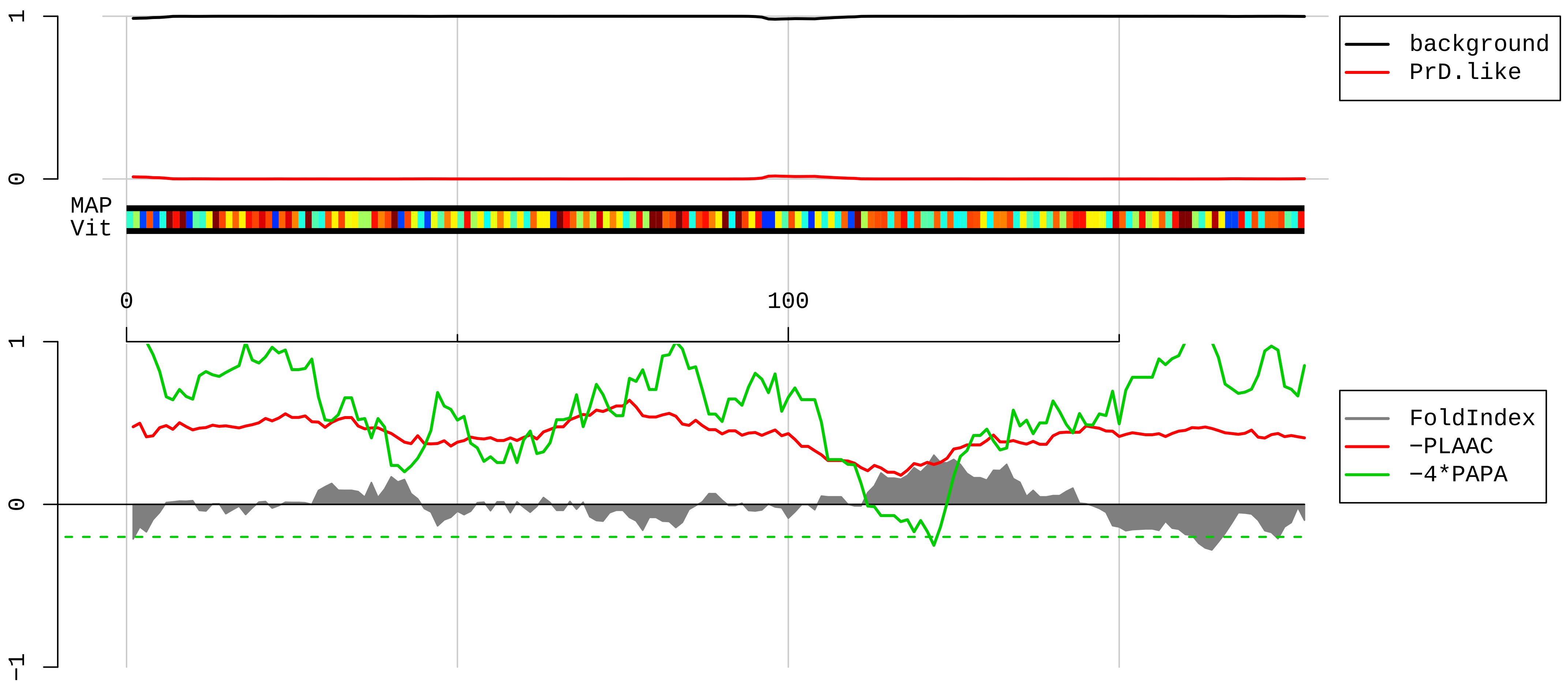


1 MAQDQGEKENPMRELRIRKLCLNICVGESGDRLTRAAKVLEQLTGQTPVFSKARYTVRSFGIRRNEKIAVHCTVRGAKAEEILEKGLKVREYELRKNNFS
101 DTGNFGFGIQEHIDLGIKYDPSIGIYGLDFYVVLGRPGFSIADKKRRTGCIGAKHRISKEEAMRWFQQKYDGIILPGK

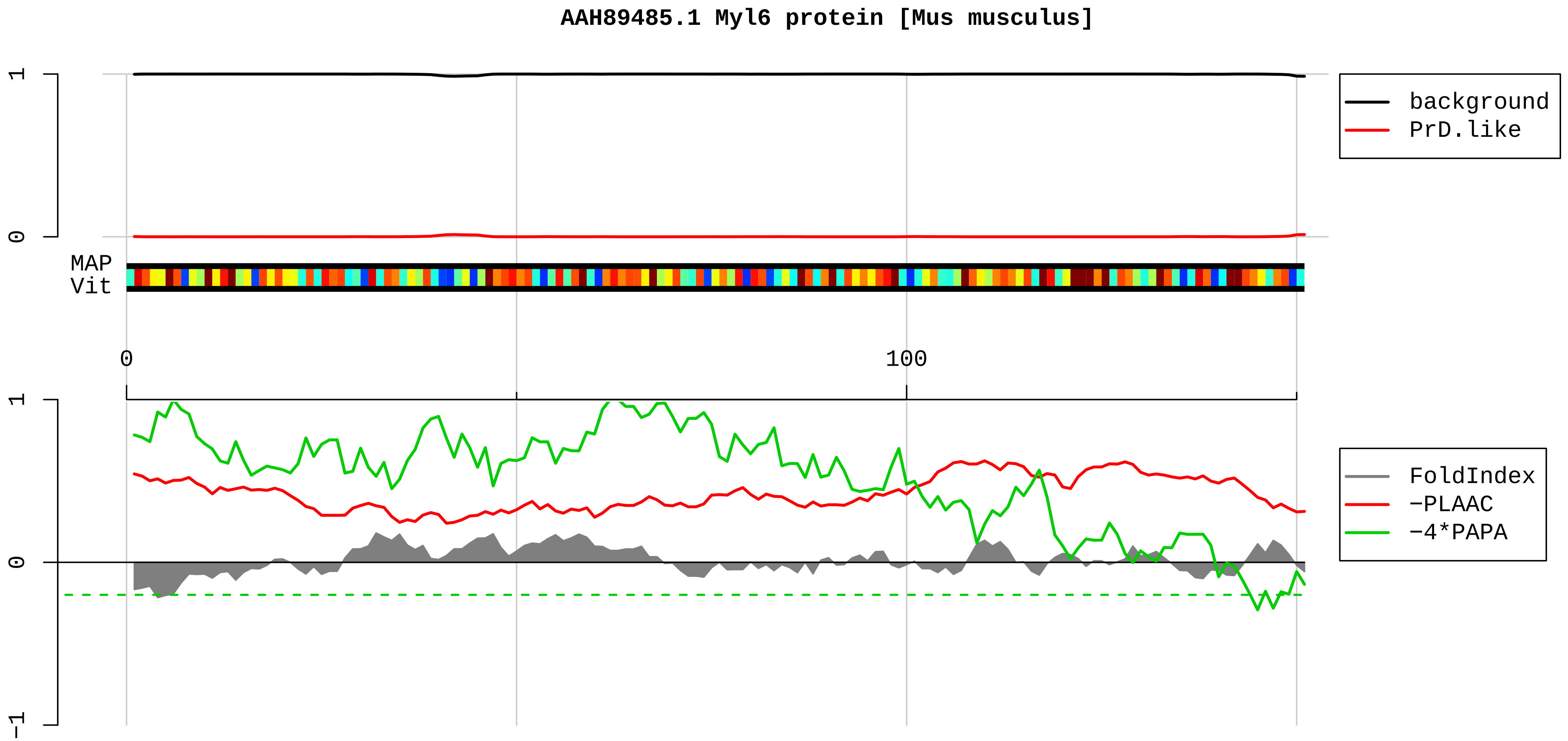


1 MCDFTEDQTAEFKEAFQLFDRTGDGKILYSQCGDVMRALGQNPTNAEVLKVLGNPKSDEMNVKVLDFEHFLPMLQTVAKNKDQGTYEDYVEGLRVFDKEG
101 NGTVMGAEIRHVLVTLGEKMTEEEVEMLVAGHEDSNGCINYEELVRMVLNG

**AAH89319.1 Ribosomal protein L9 [Mus musculus]**

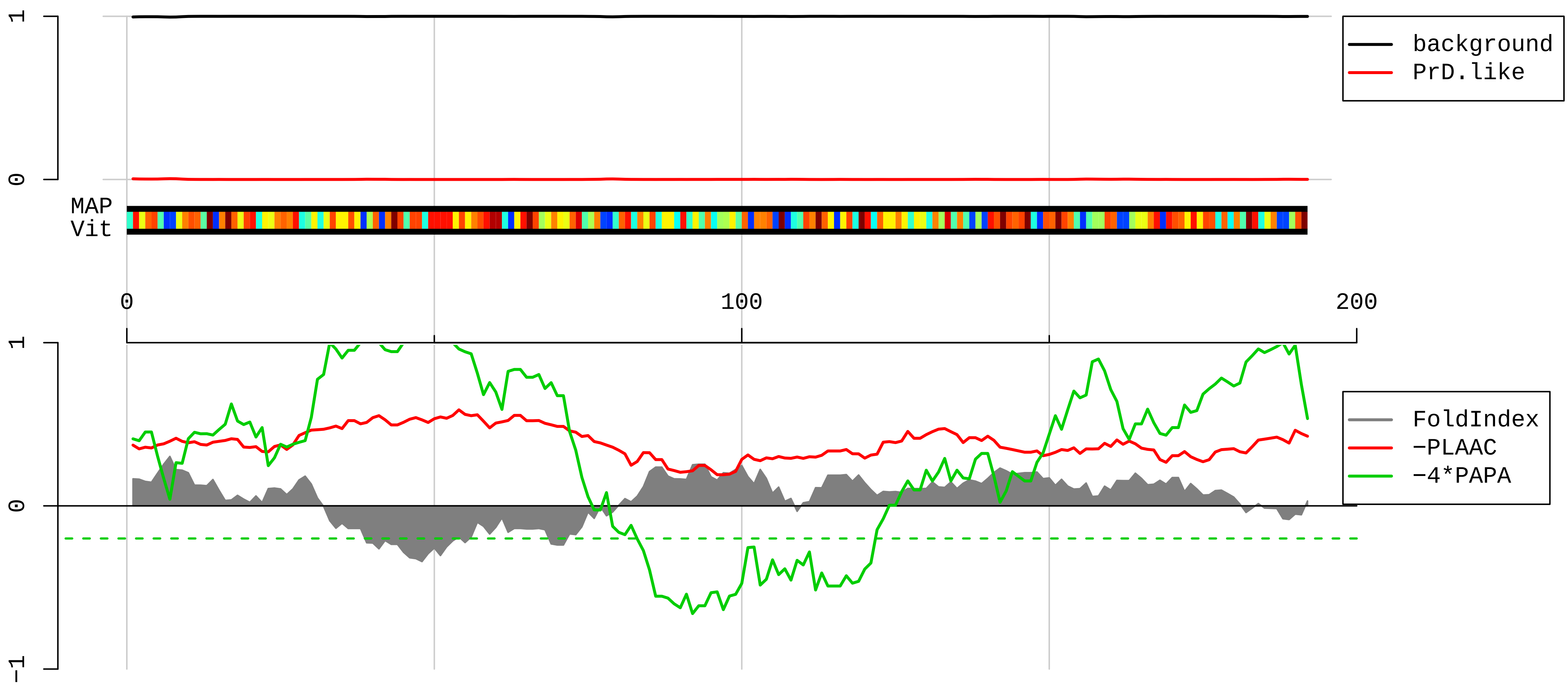


1 MKTILSNQTVDIPENVEITLKGRTVIVKGPRGTLRRDFNHINVELSLLGKKKKRLRVDKWWGNRKELATVRTICSHVQNMIKGVTLGFRYKMRSVYAHFP
101 INVVIQENGSLVEIRNFLGEKYIRRVRMRTGVACSVSQAQKDELILEGNDIELVSNSAALIQQATTVKNKDIRKFLDGIYVSEKGTVQQADE

#### AAH92008.1 Ribosomal protein L24 [Mus musculus]

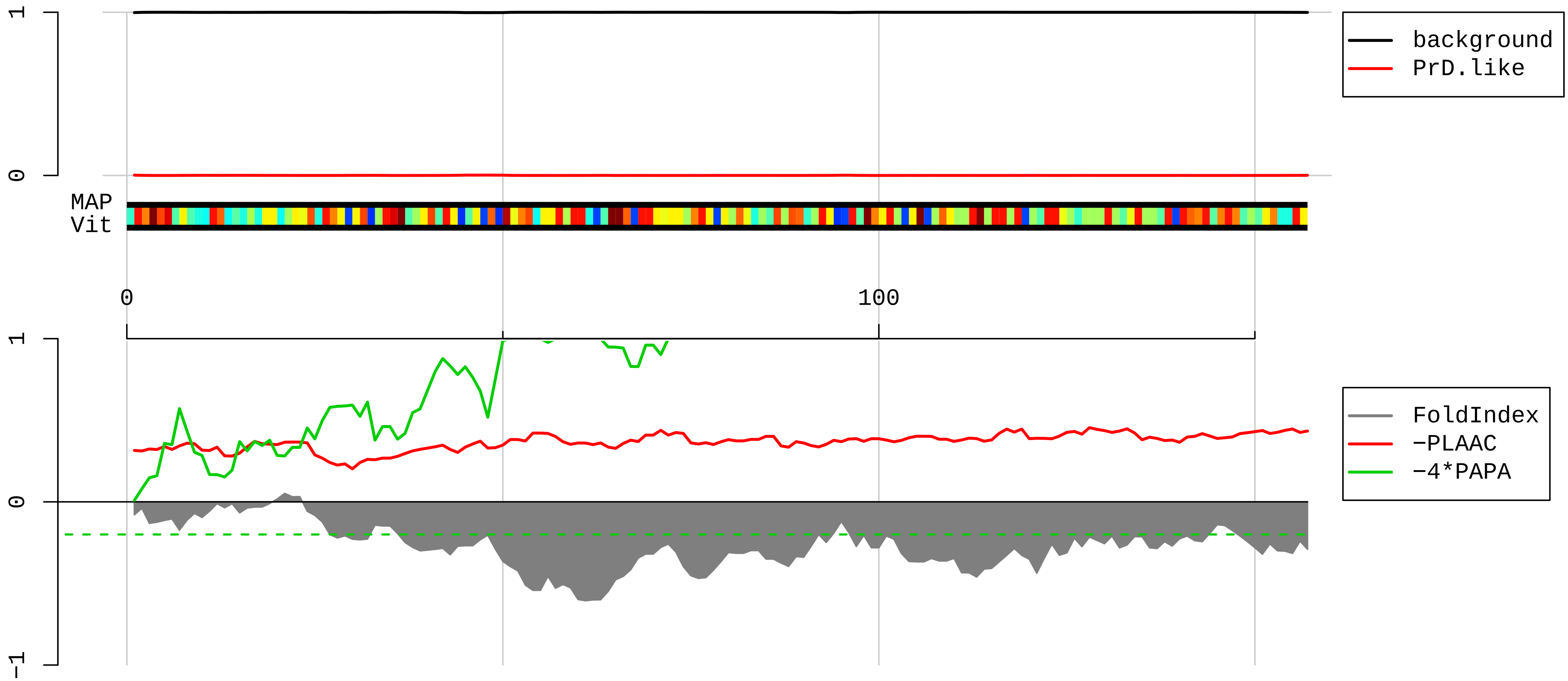


1 MKVELCSFSGYKIYPGHGRRYARTDGKVFQFLNAKCESAFLSKRNPRQINWTVLYRRKHKKGQSEEIQKKRTRRAVKFQRAITGASLADIMAKRNQKPEV
101 RKAQREQAIRAAKEAKKAKQASKKTAMAAAKAPTKAAPKQKIVKPVKVSAPRVGGKR

**AAI00336.1 Ribosomal protein L15 [Mus musculus]**

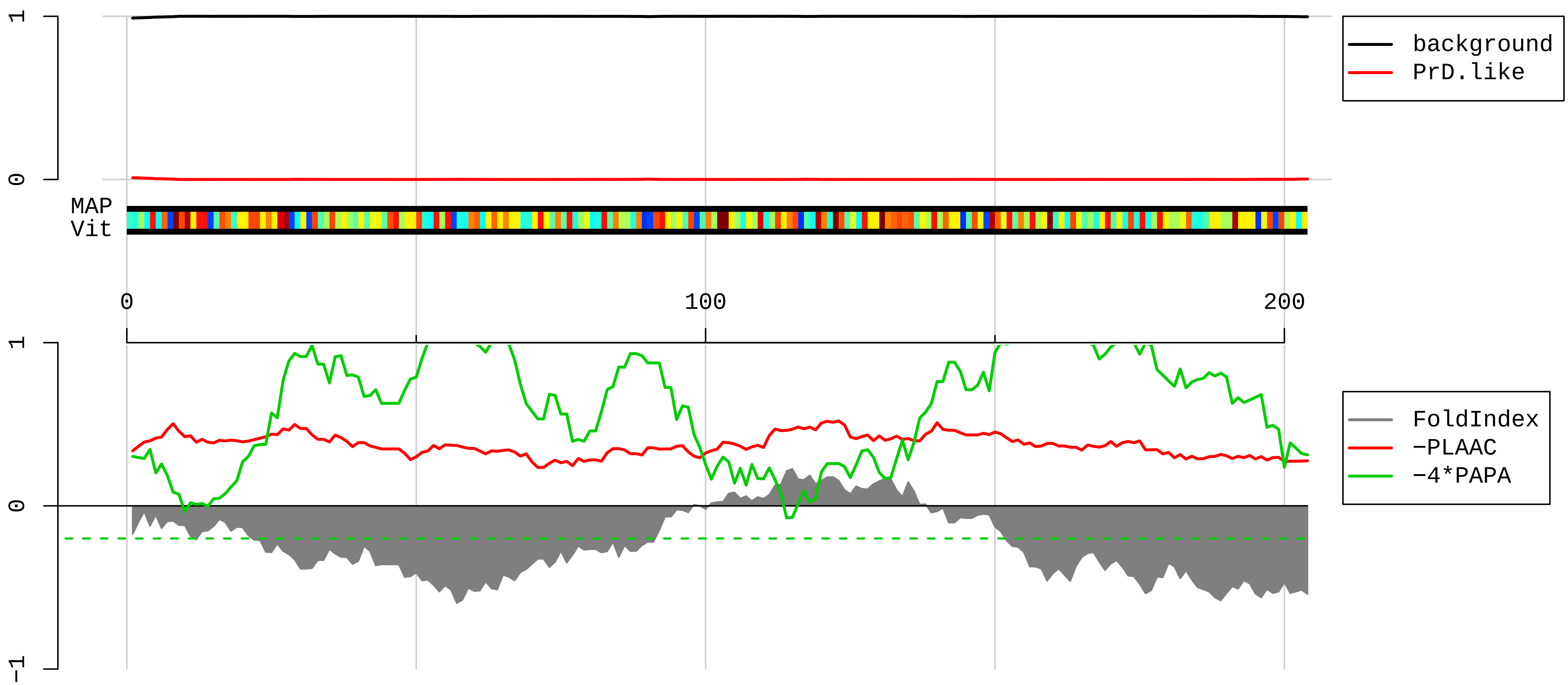


1 MGAYKYIQELWRKKQSDVMRFLLRVRCWQYRQLSALHRAPRPTRPDKARRLGYKAKQGYVIYRIRVRRGGRKRPVPKGATYGKPVHHGVNQLKFARSLQS
101 VAEERAGRHCGALRVLNSYWVGEDSTYKFFEVILIDPFHKAIRRNPDTQWITKPVHKHREMRGLTSAGRKSRGLGKGHKFHHTIGGSRRAAWRRRNTLQL
201 HRYR

**AAH89323.1 Ribosomal protein S10 [Mus musculus]**

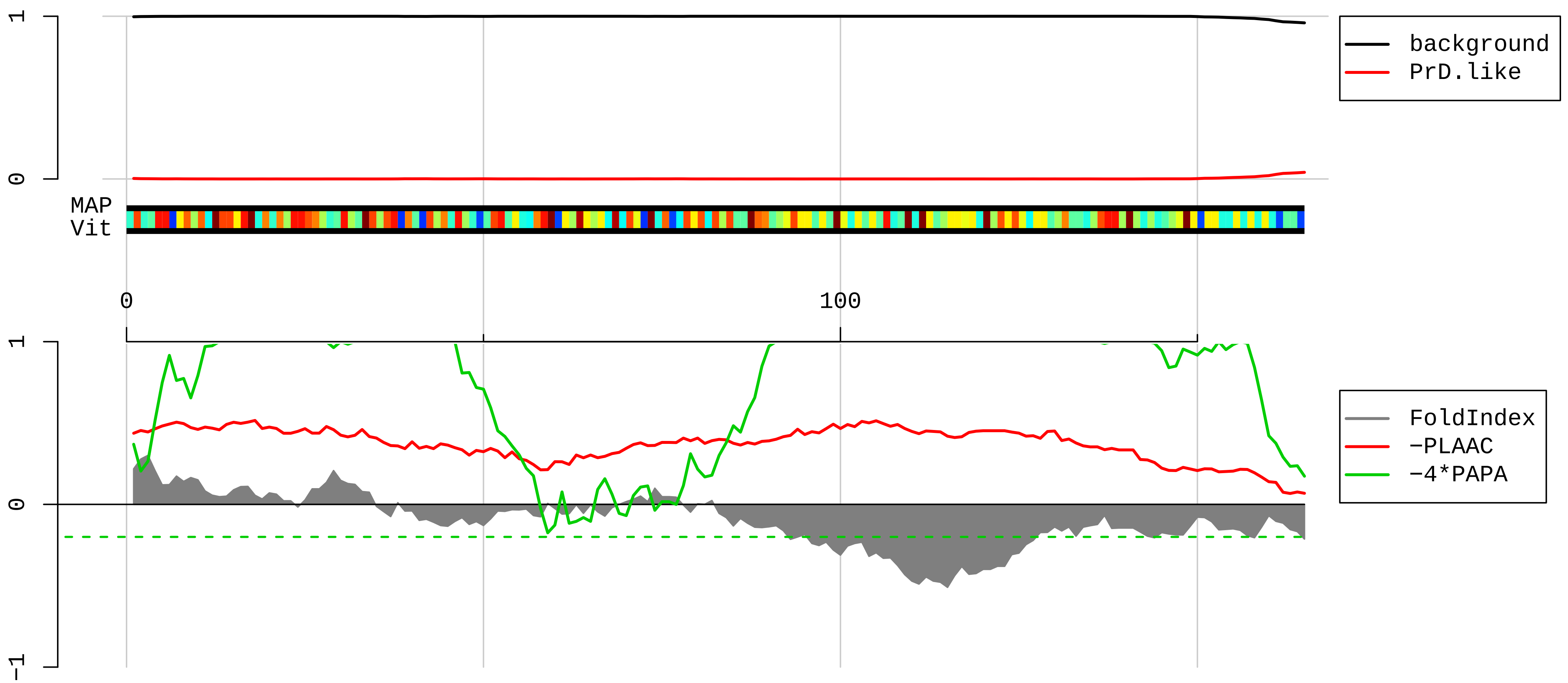


1 MLMPKKNRIAIYELLFKEGVMVAKKDVHMPKHPELADKNVPNLHVMKAMQSLKSRGYVKEQFAWRHFYWYLTNEGIQYLRDYLHLPPEIVPATLRRSRPE

101 TGRPRPKGPEGERPARFTRGEADRDTYRRSAVPPGADKKAEAGAGSATEFQFRGGFGRGRGQPPQ

**AAH81468.1 Ribosomal protein L18 [Mus musculus]**

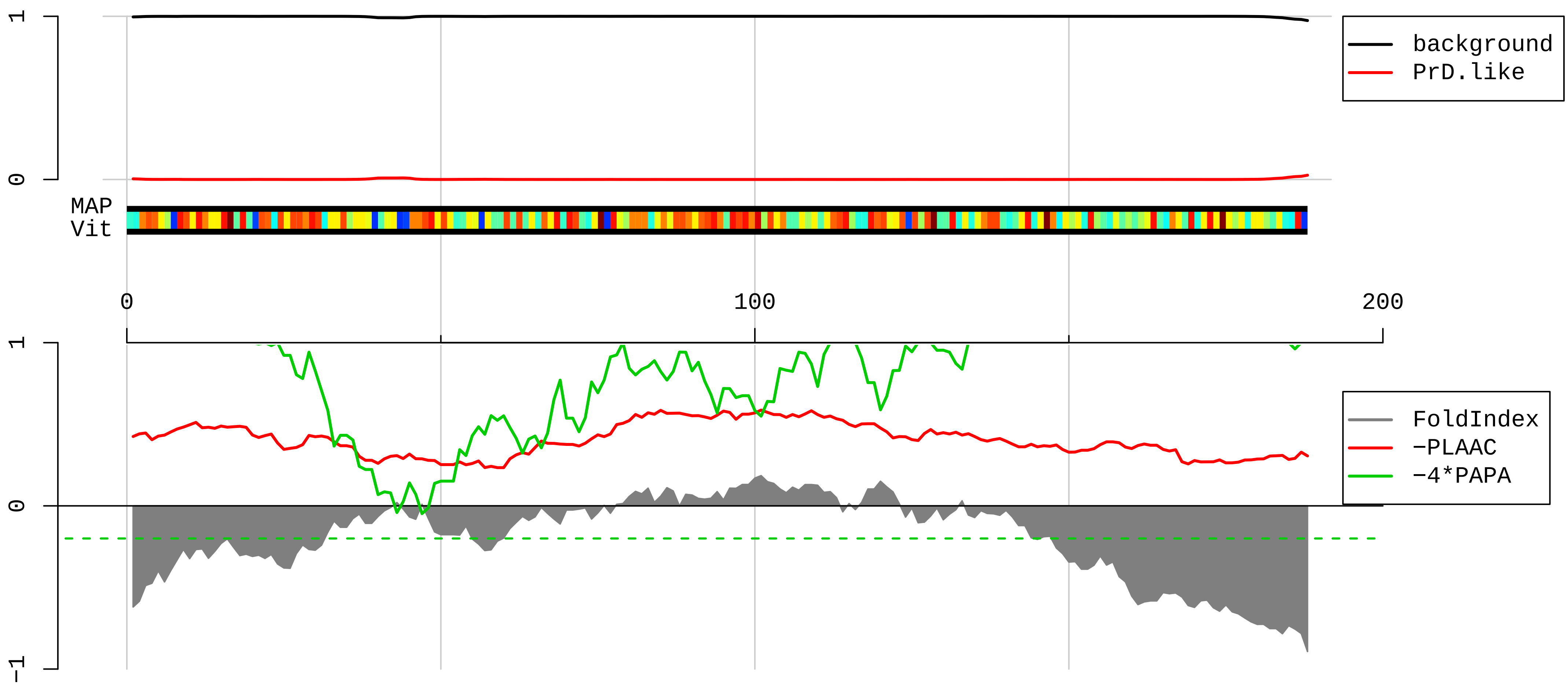


1 MGVDIRHNKDRKVRRKEPKSQDIYLRLLVKLYRFLARRTNSTFNQVVLKRLFMSRTNRPPLSLSRMIRKMKLPGRENKTAVVVGTVTDDVRILKVPKLKV
101 CALRVSSRARSRILKAGGKILTFDQLALESPKGRGTVLLSGPRKGREVYRHFGKAPGTPHSHTKPYVRSKGRKFERARGRRASRGYKN

**AAH12641.1 Ribosomal protein S11 [Mus musculus]**

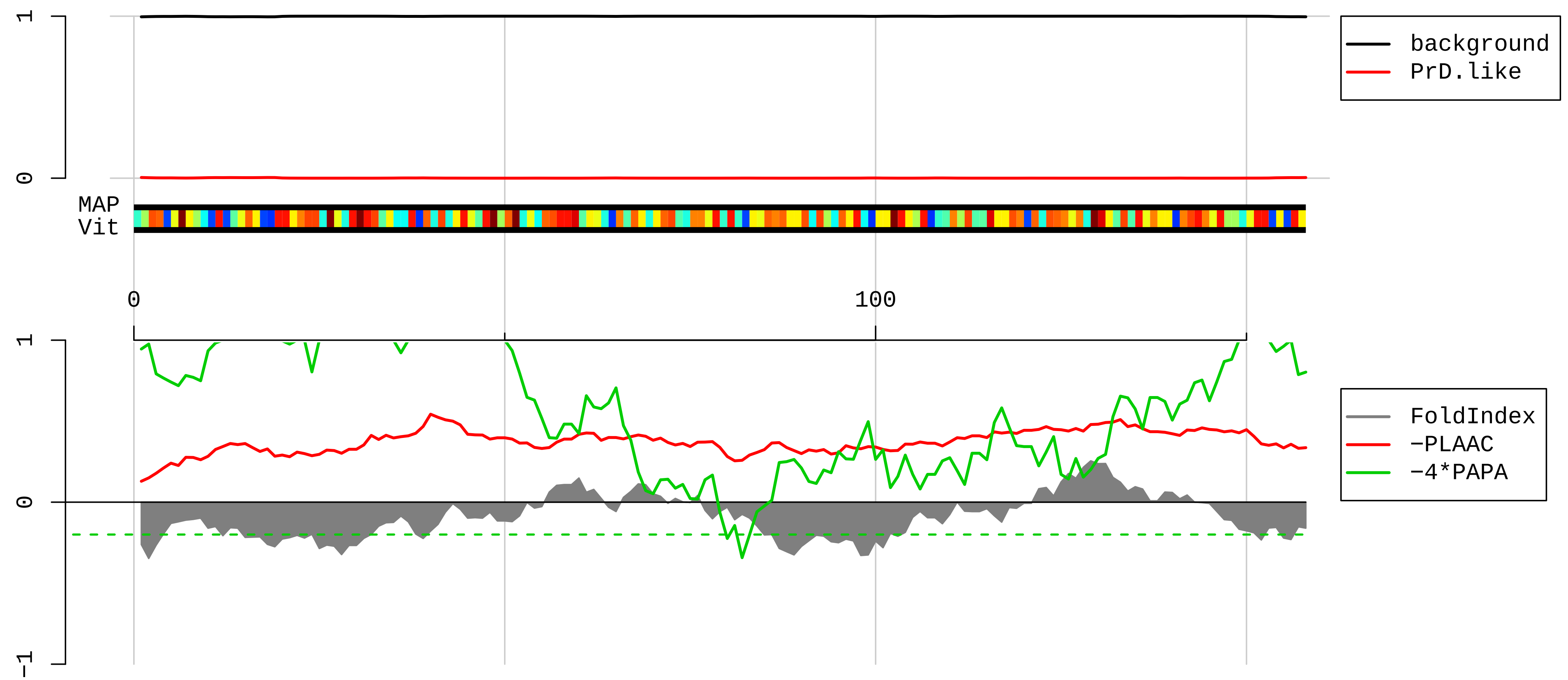


1 MADIQTERAYQKQPTIFQNKKRVLLGETGKEKLPRYYKNIGLGFKTPKEAIEGTYIDKKCPFTGNVSIRGRILSGVVTKMKMQRTIVIRRDYLHYIRKYN
101 RFEKRHKNMSVHLSPCFRDVQIGDIVTVGECRPLSKTVRFNVLKVTKAAGTKKQFQKF

AAH21344.1 Ribosomal protein L22 [Mus musculus]

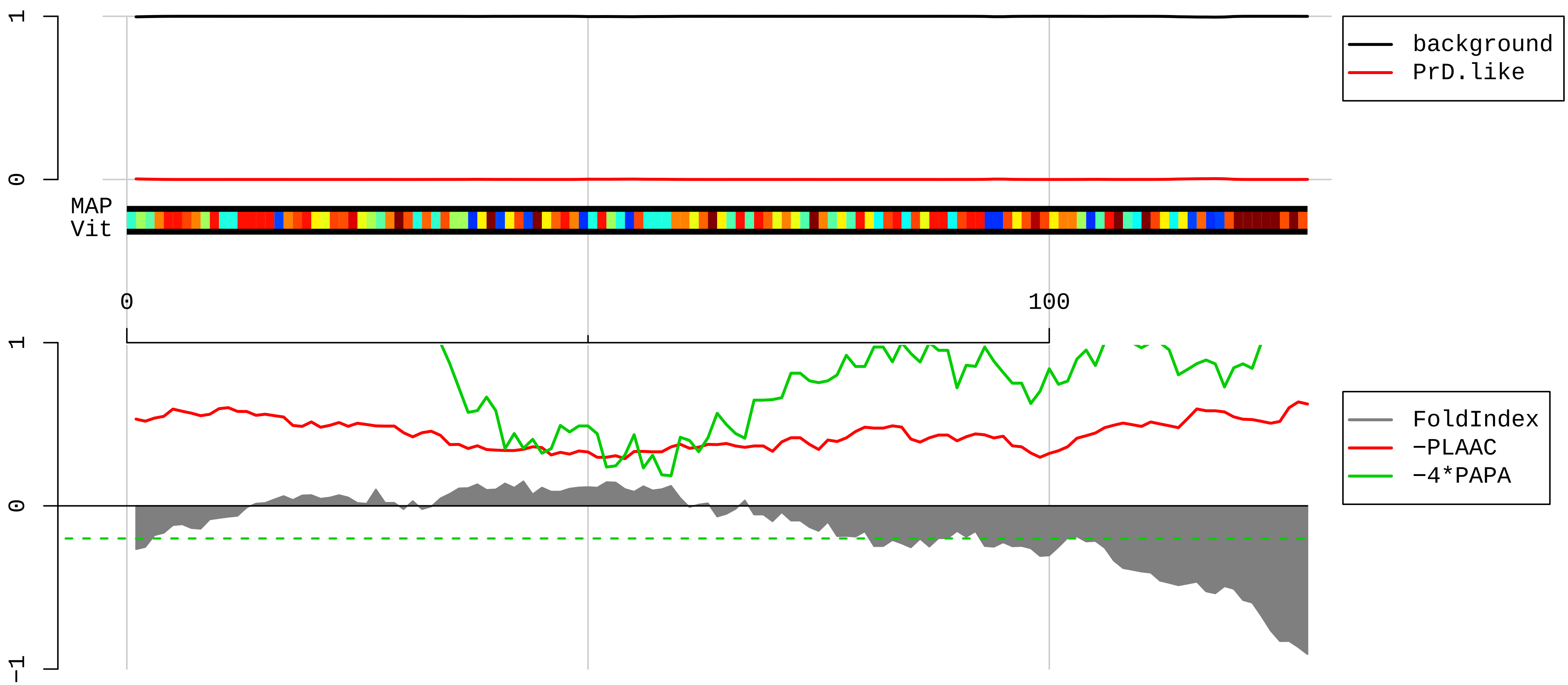


1 MAPVKKLVAKGGKKKKQVLKFTLDCTHPVEDGIMDAANFEQFLQERIKVNGKAGNLGGGVVTIERSKSKITVTSEVPFSKRYLKYLTKKYLKKNNLRDWL
101 RVVANSKESYELRYFQINQDEEEEEDED

**AAH99436.1 Ribosomal protein L31 [Mus musculus]**

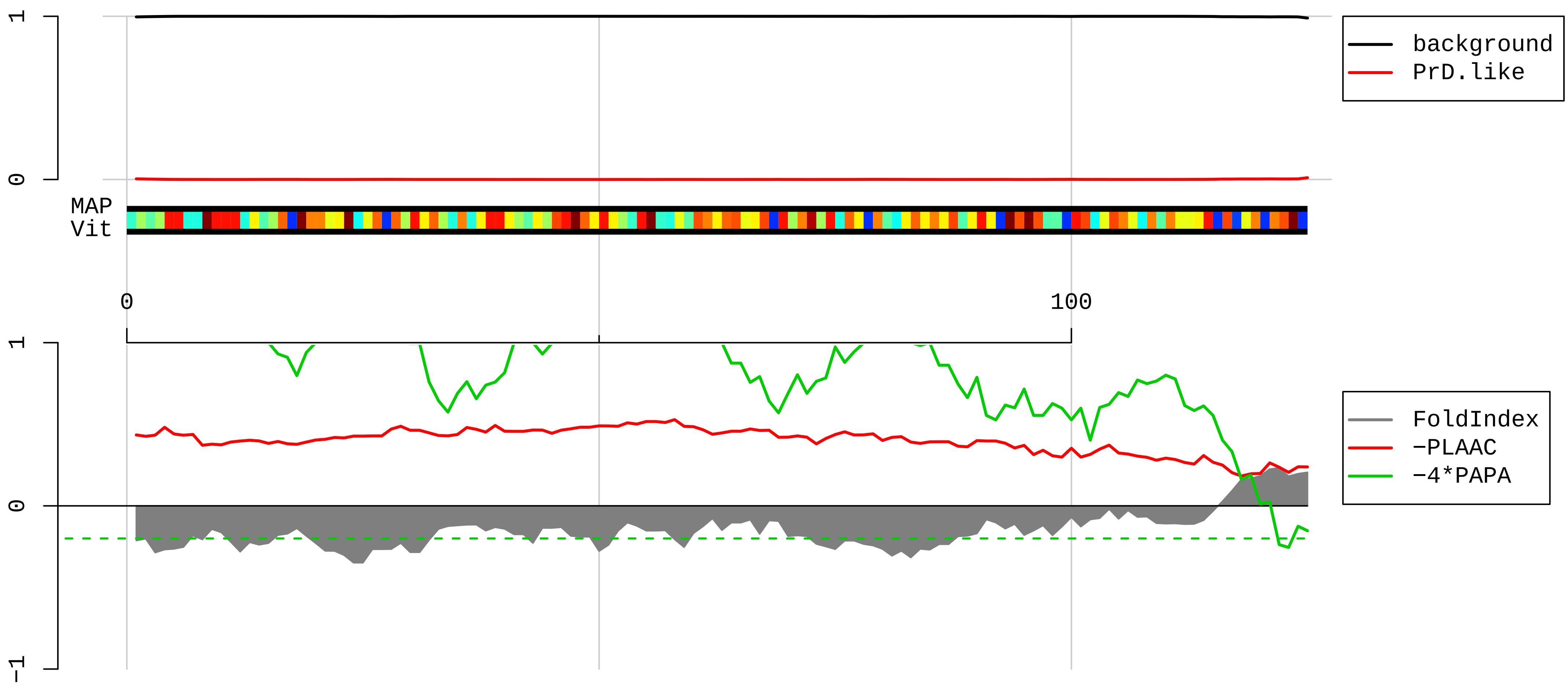


1 MAPAKKGGEKKKGRSAINEVVTREYTINIHKRIHGVGFKKRAPRALKEIRKFAMKEMGTPDVRIDTRLNKAVWAKGIRNVPYRIRVRLSRKRNEDEDSPN

101 KLYTLVTYVPVTTFKNLQTVNVDEN

**AAH91748.1 Ribosomal protein S24 [Mus musculus]**

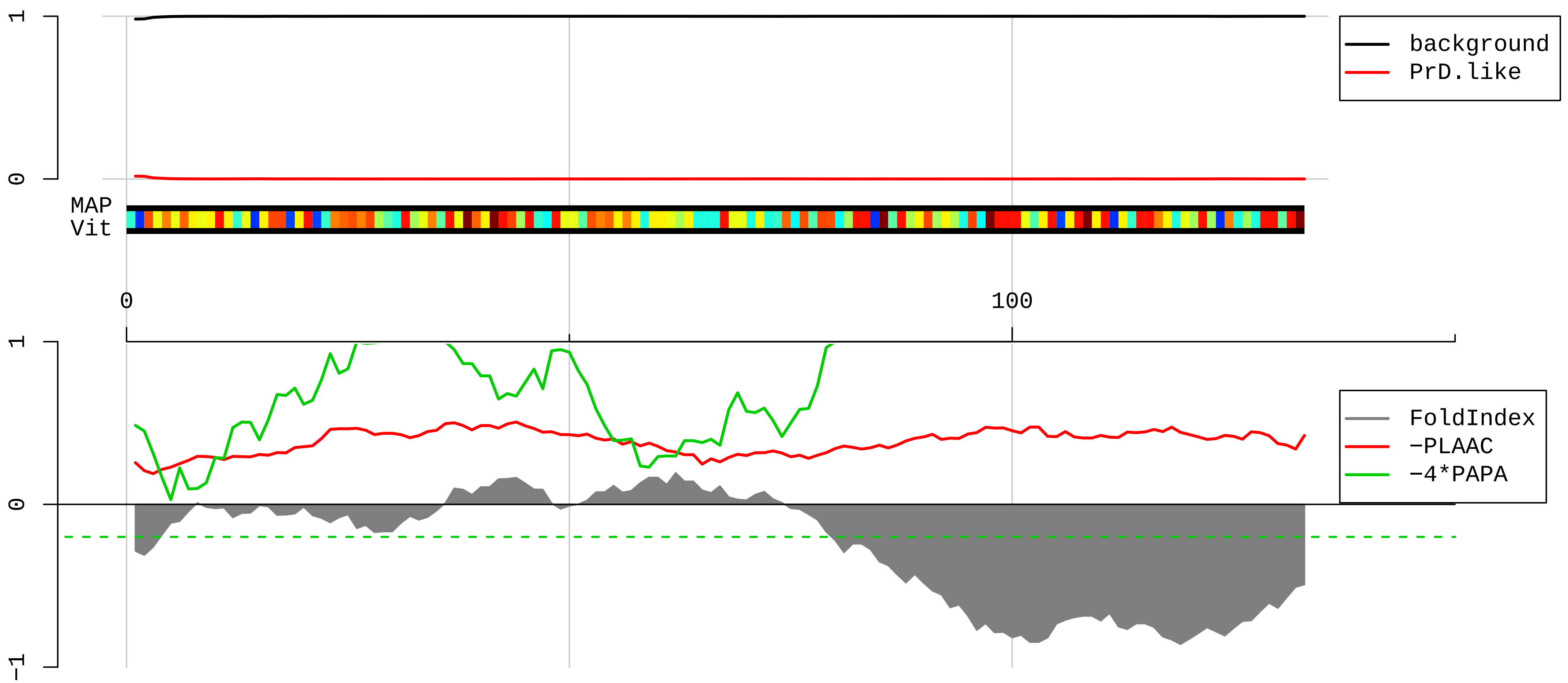


1 MNDTVTIRTRKFMTNRLLQRKQMVIDVLHPGKATVPKTEIREKLAKMYKTTPDVIFVFGFRTHFGGGKTTGFGMIYDSLDYAKKNEPKHRLARHGLYEKK

101 KTSRKQRKERKNRMKKVRGTAKANVGAGKKPKE

**AAH78418.1 Ribosomal protein S23 [Mus musculus]**

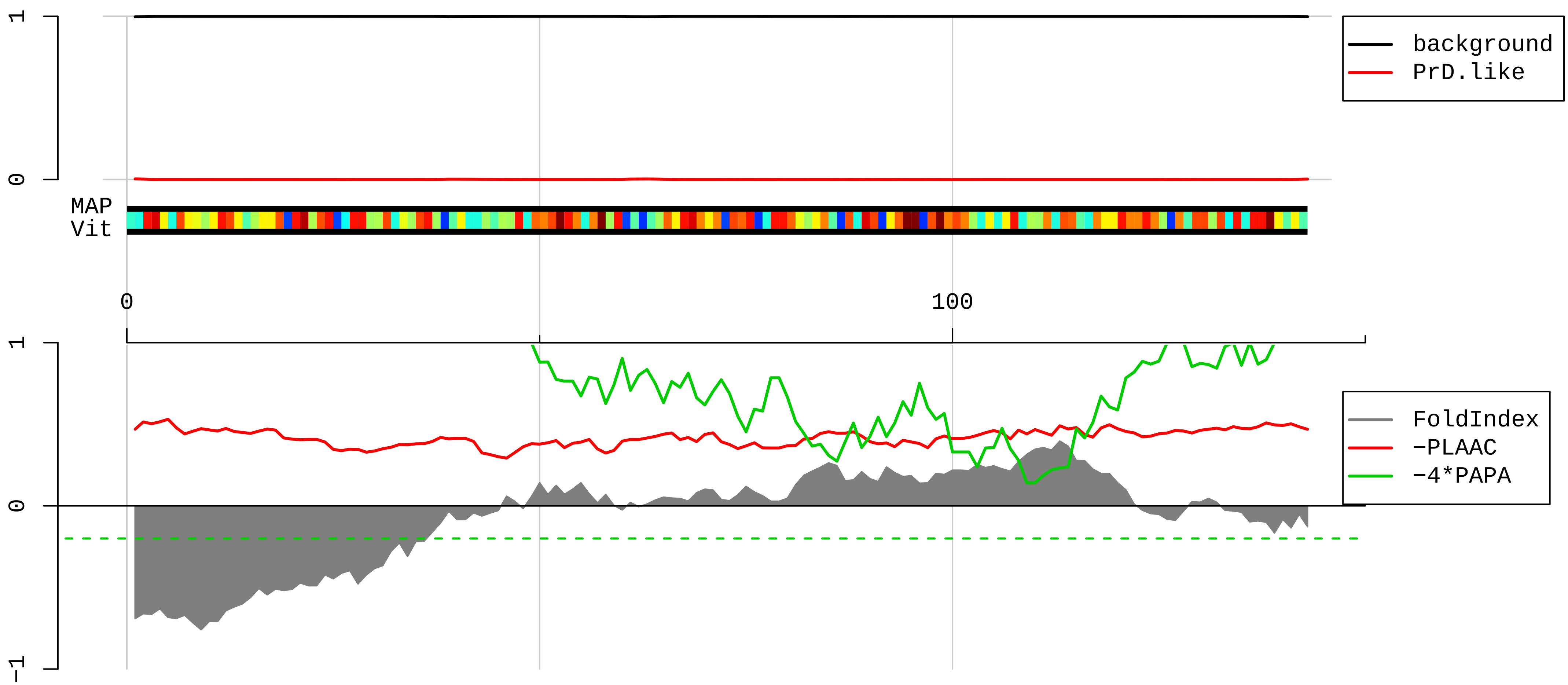


1 MGKCRGLRTARKLRSHRRDQKWHDKQYKKAHLGTALKANPFGGASHAKGIVLEKVGVEAKQPNSAIRKCVRVQLIKNGKKITAFVPNDGCLNFIEENDEV
101 LVAGFGRKGHAVGDIPGVRFKVVKVANVSLLALYKGKKERPRS

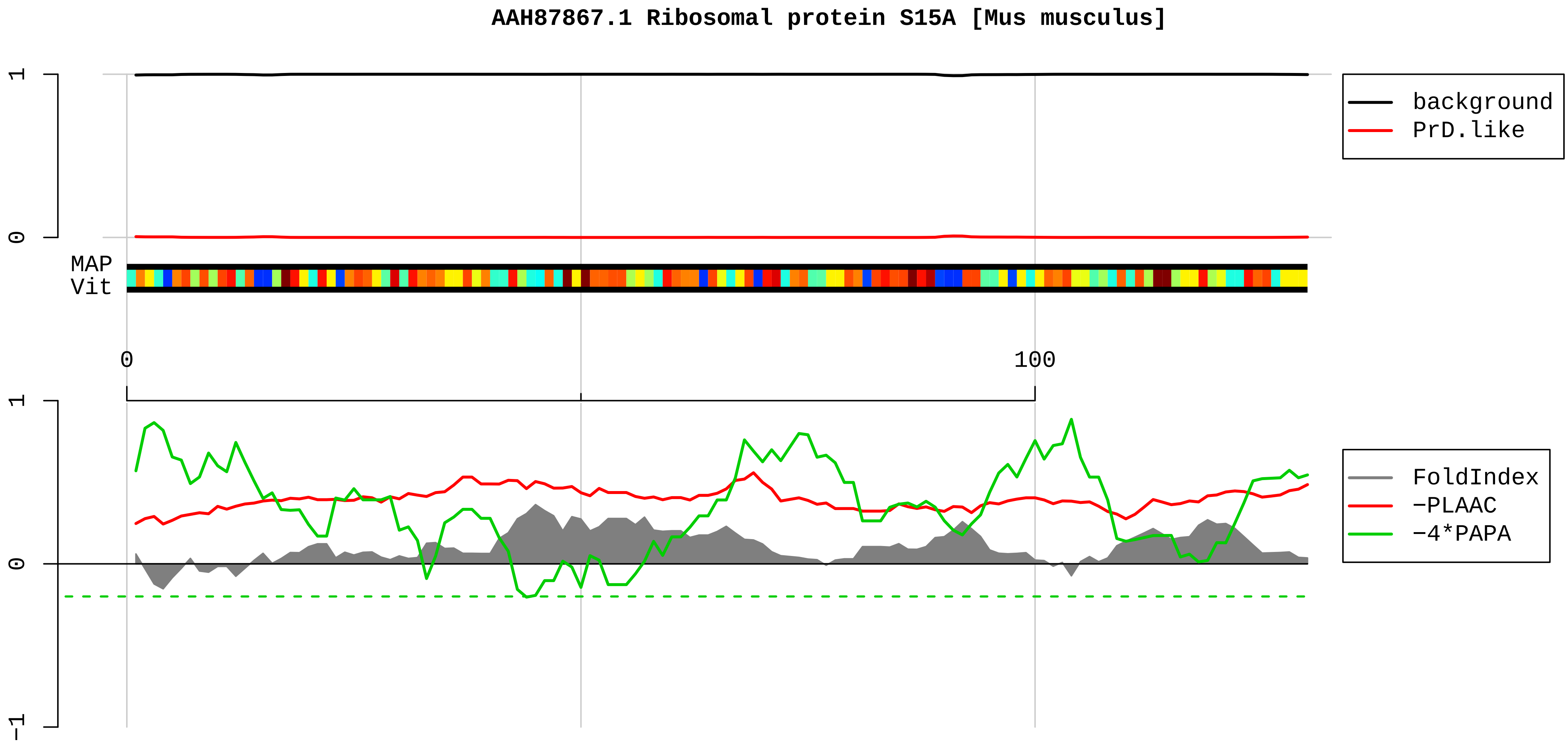


1 MVRMNVLADALKSINNAEKRGKRQVLIRPCSKVIVRFLTVMMKHGYIGEFEIIDDHRAGKIVVNLTGRLNKCGVISPRFDVQLKDLEKWQNNLLPSRQFG
101 FIVLTTSAGIMDHEEARRKHTGGKILGFFF

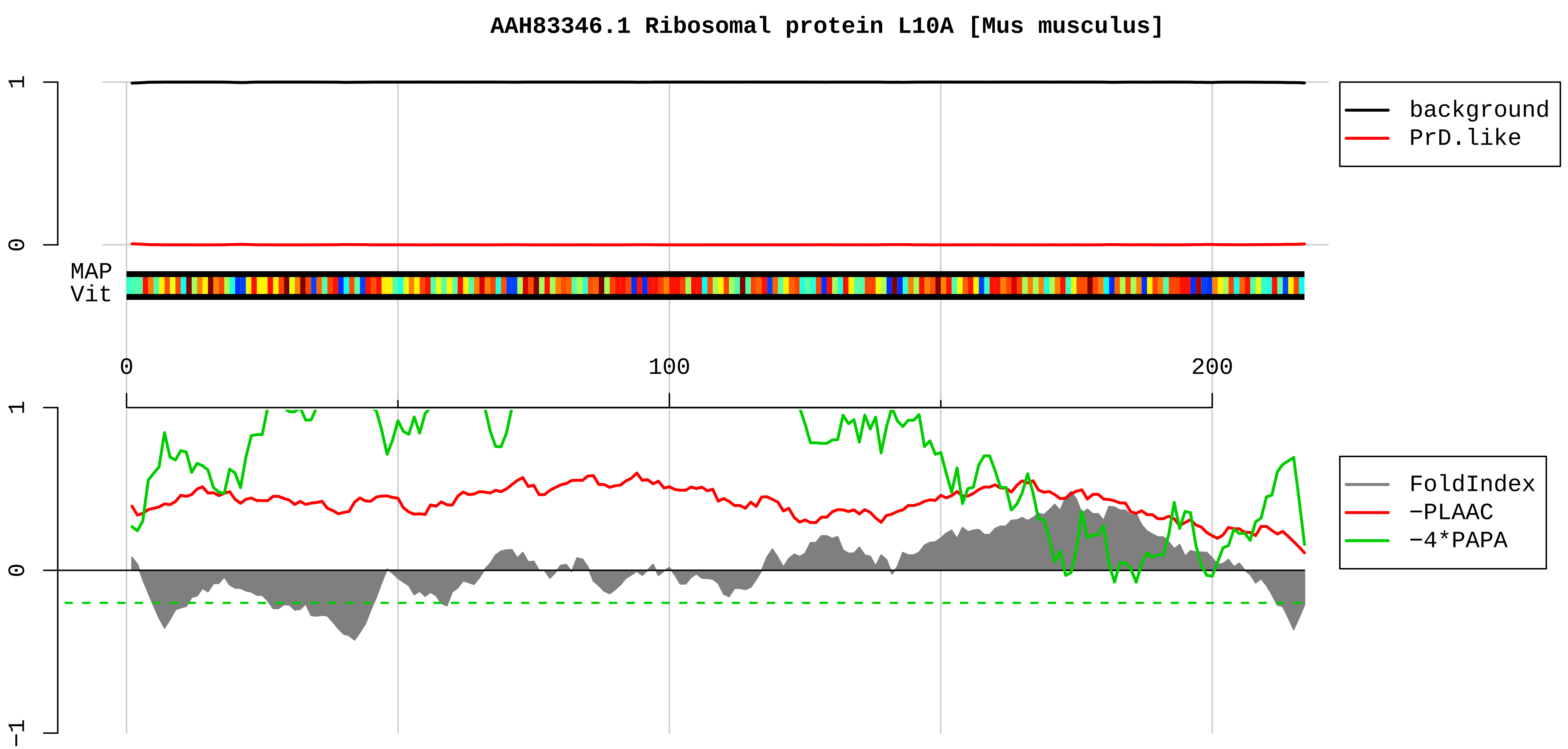


1 MSSKVSRDTLYEAVREVLHGNQRKRRKFLETVELQISLKNYDPQKDKRFSGTVRLKSTPRPKFSVCVLGDQQHCDEAKAVDIPHMDIEALKKLNKNKKLV
101 KKLAKKYDAFLASESLIKQIPRILGPGLNKAGKFPSLLTHNENMVAKVDEVKSTIKFQMKKVLCLAVAVGHVKMTDDELVYNIHLAVNFLVSLLKKNWQN
201 VRALYIKSTMGKPQRLY

#### AAH92005.1 Ribosomal protein S25 [Mus musculus]

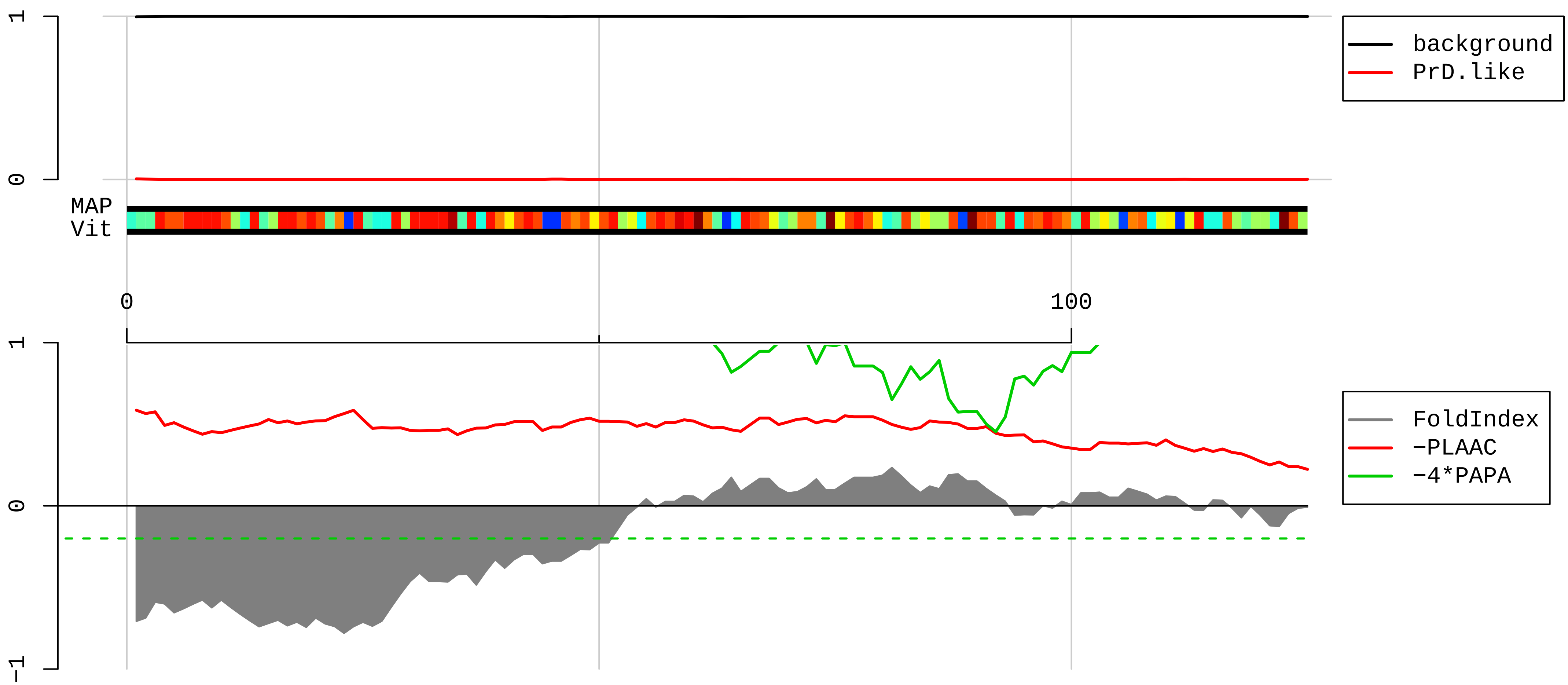


1 MPPKDDKKKKDAGKSAKKDKDPVNKSGGKAKKKKWSKGKVRDKLNNLVLFDKATYDKLCKEVPNYKLITPAVVSERLKIRGSLARAALQELLSKGLIKLV
101 SKHRAQVIYTRNTKGGDAPAAGEDA

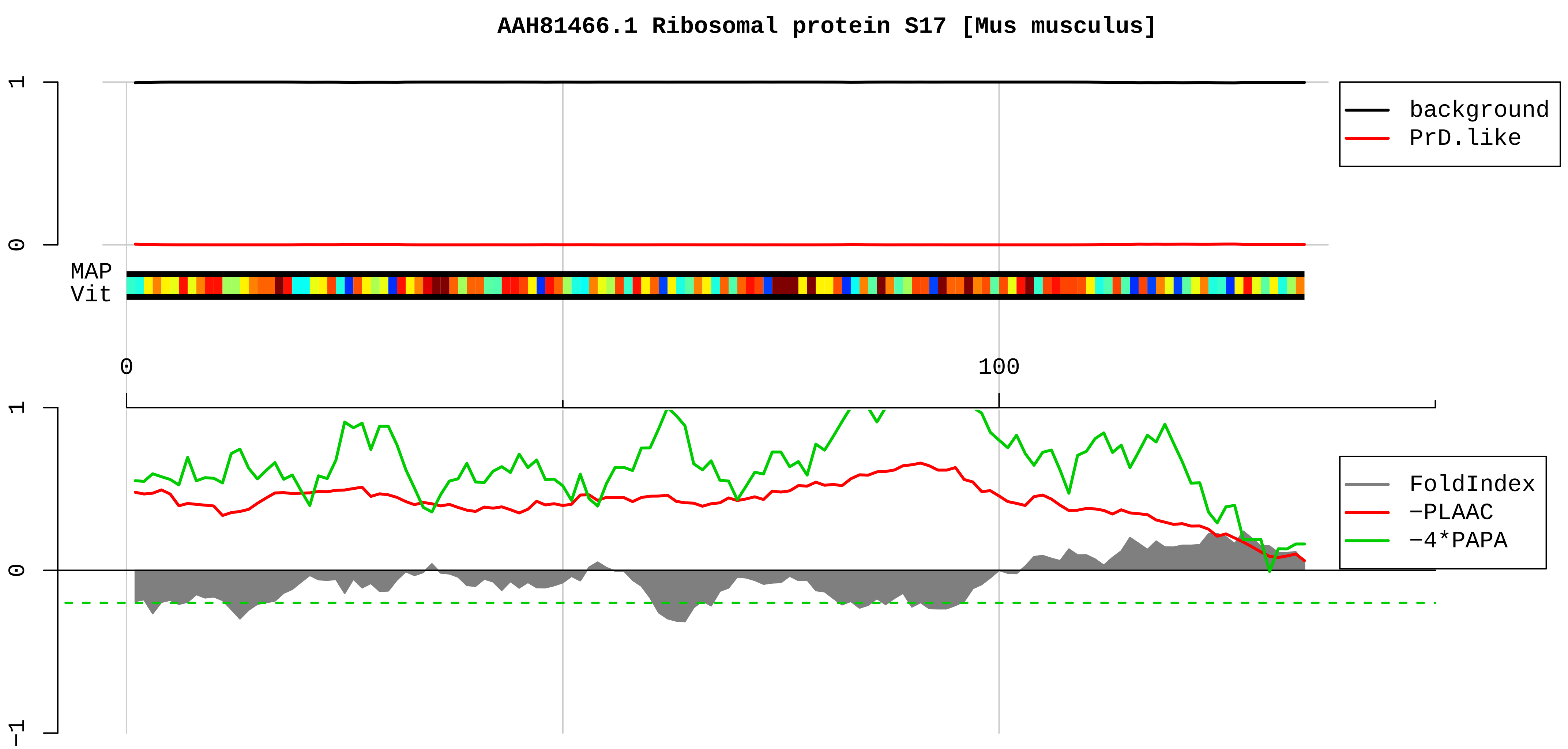


1 MGRVRTKTVKKAARVIIEKYYTRLGNDFHTNKRVCEEIAIIPSKKLRNKIAGYVTHLMKRIQRGPVRGISIKLQEEERERRDNYVPEVSALDQEIIEVDP
101 DTKEMLKLLDFGSLSNLQVTQPTVGMNFKTPRGAV

**AAI25657.1 Ribosomal protein L35 [Mus musculus]**

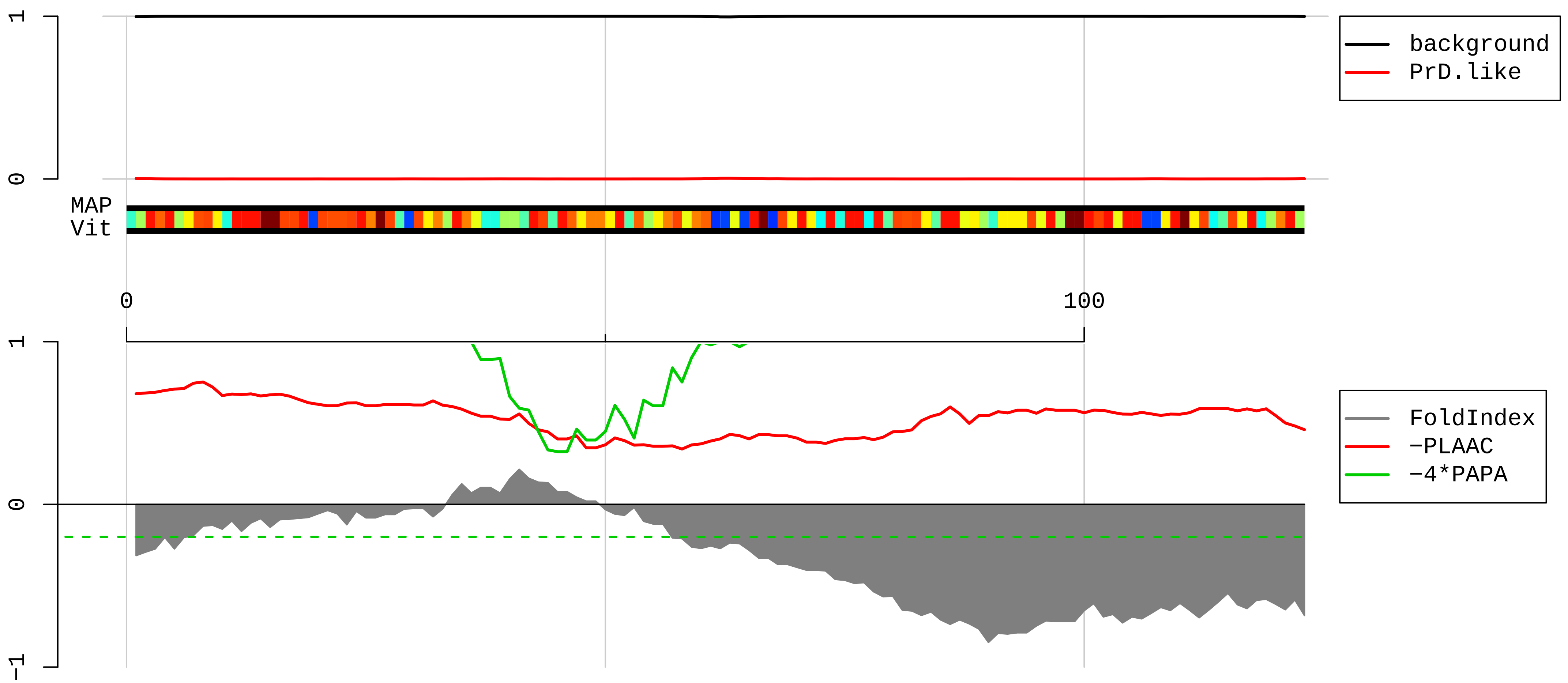


1 MAKIKARDLRGKKKEELLKQLDDLKVELSQLRVAKVTGGAASKLSKIRVVRKSIARVLTVINQTQKENLRKFYKGKKYKPLDLRPKKTRAMRRRLTKHEE
101 KLKTKKQQRKERLYPLRKYAVKA

**AAH02014.1 Ribosomal protein S7 [Mus musculus]**

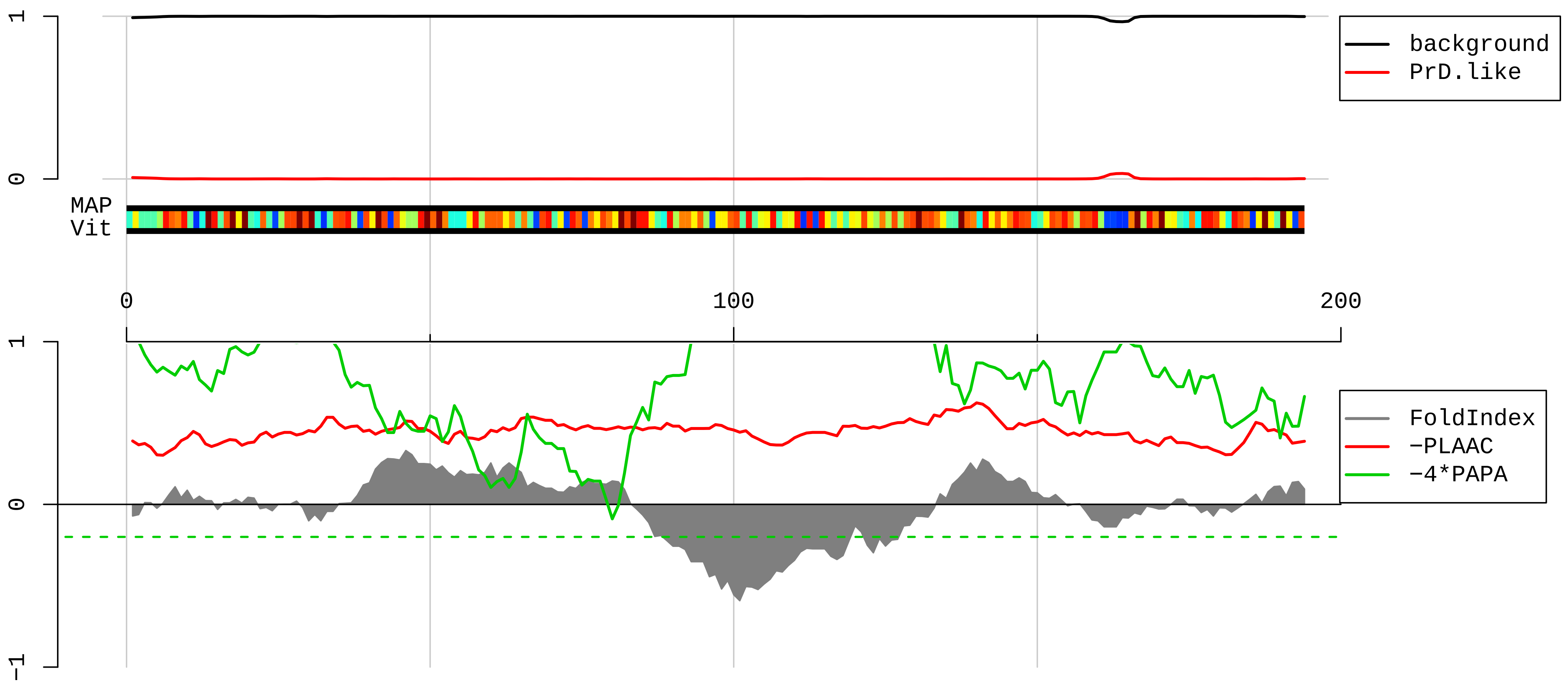


1 MFSSSAKIVKPNGEKPDEFESGISQALLELEMNSDLKAQLRELNITAAKEIEVGGGRKAIIIFVPVPQLKSFQKIQVRLVRELEKKFSGKHVVFIAQRRI
101 LPKPTRKSRTKNKQKRPRSRTLTAVHDAILEDLVFPSEIVGKRIRVKLDGSRLIKVHLDKAQQNNVEHKVETFSGVYKKLTGKDVNFEFPEFQL

#### AAI25645.1 Ribosomal protein S28 [Mus musculus]

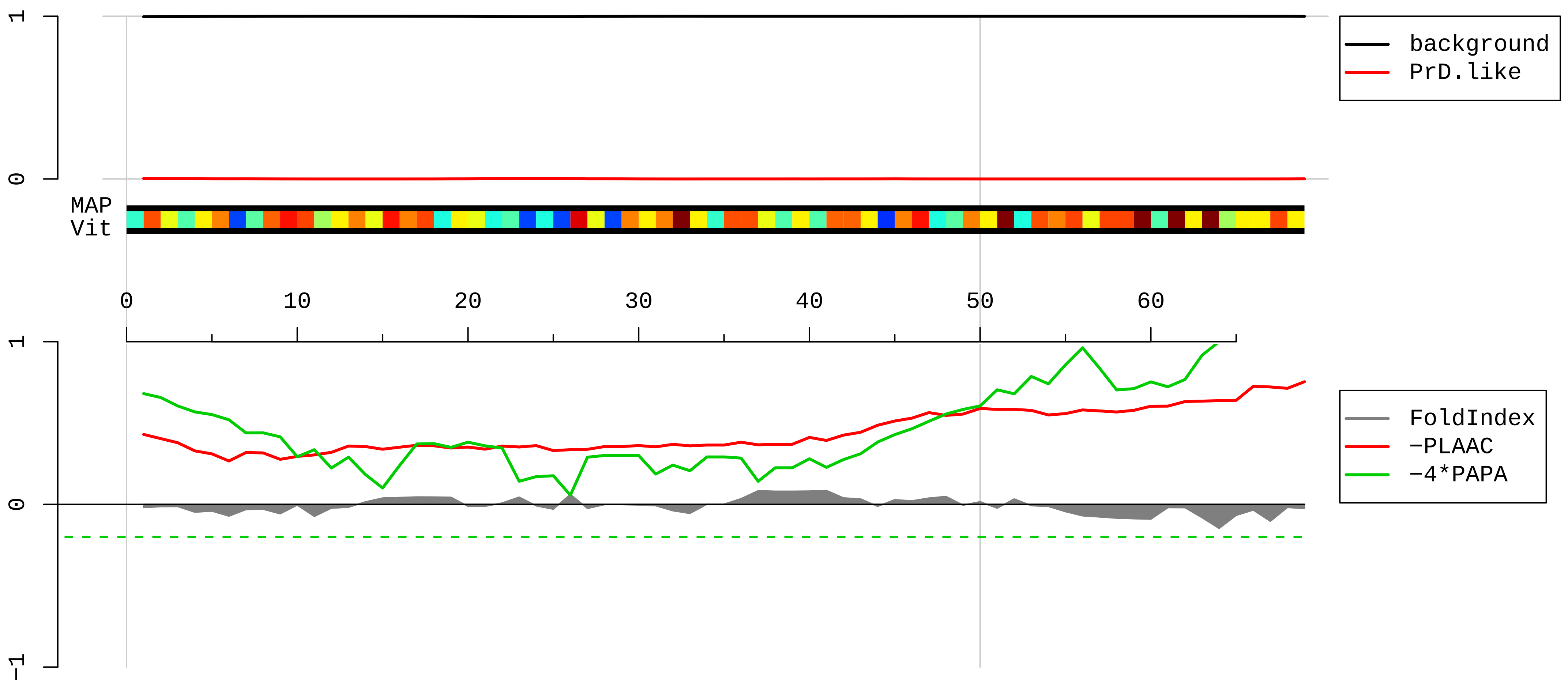


1 MDTSRVQPIKLARVTKVLGRTGSQGQCTQVRVEFMDDTSRSIIRNVKGPVREGDVLTLLESEREARRLR

**AAH11323.1 Ribosomal protein S20 [Mus musculus]**

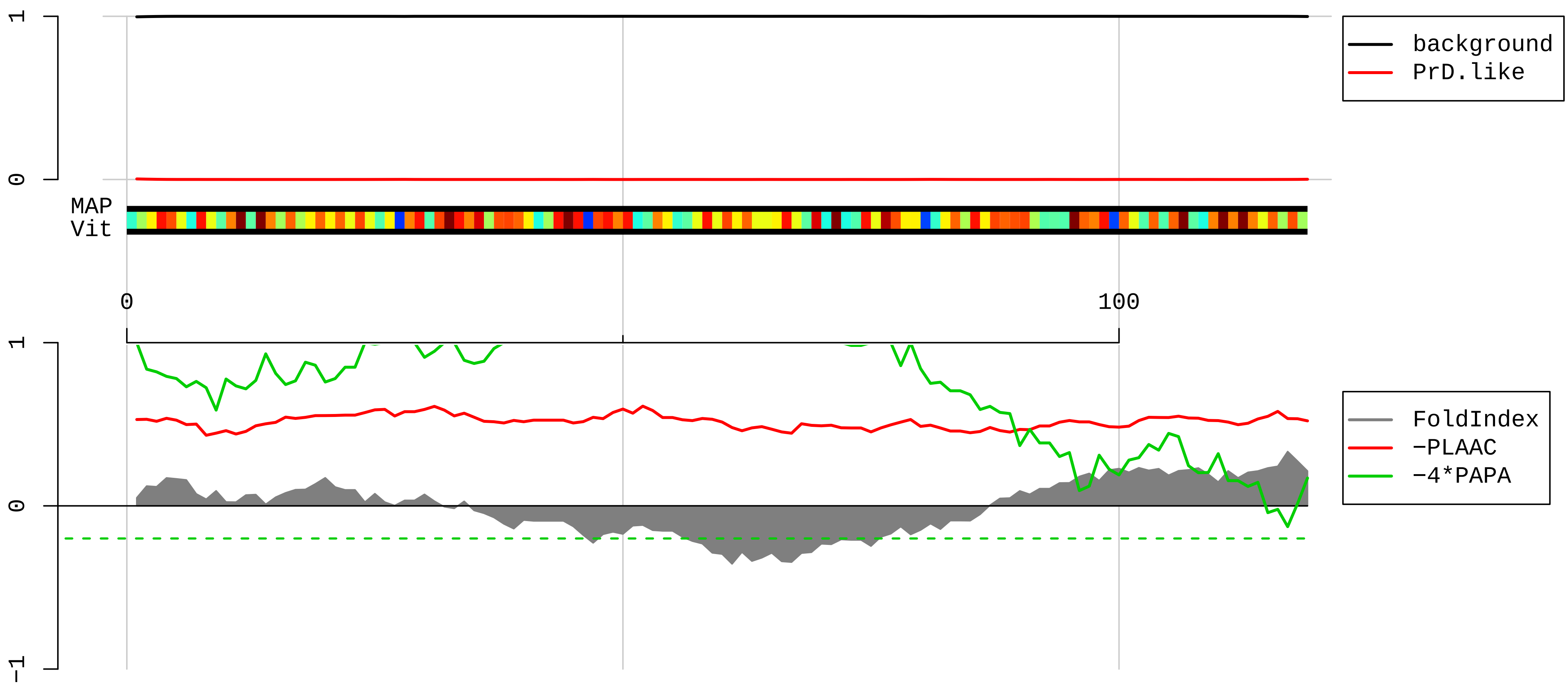


1 MAFKDTGKTPVEPEVAIHRIRITLTSRNVKSLEKVCADLIRGAKEKNLKVKGPVRMPTKTLRITTRKTPCGEGSKTWDRFQMRIHKRLIDLHSPSEIVKQ
101 ITSISIEPGVEVEVTIADA

**AAH99377.1 Ribosomal protein S12 [Mus musculus]**

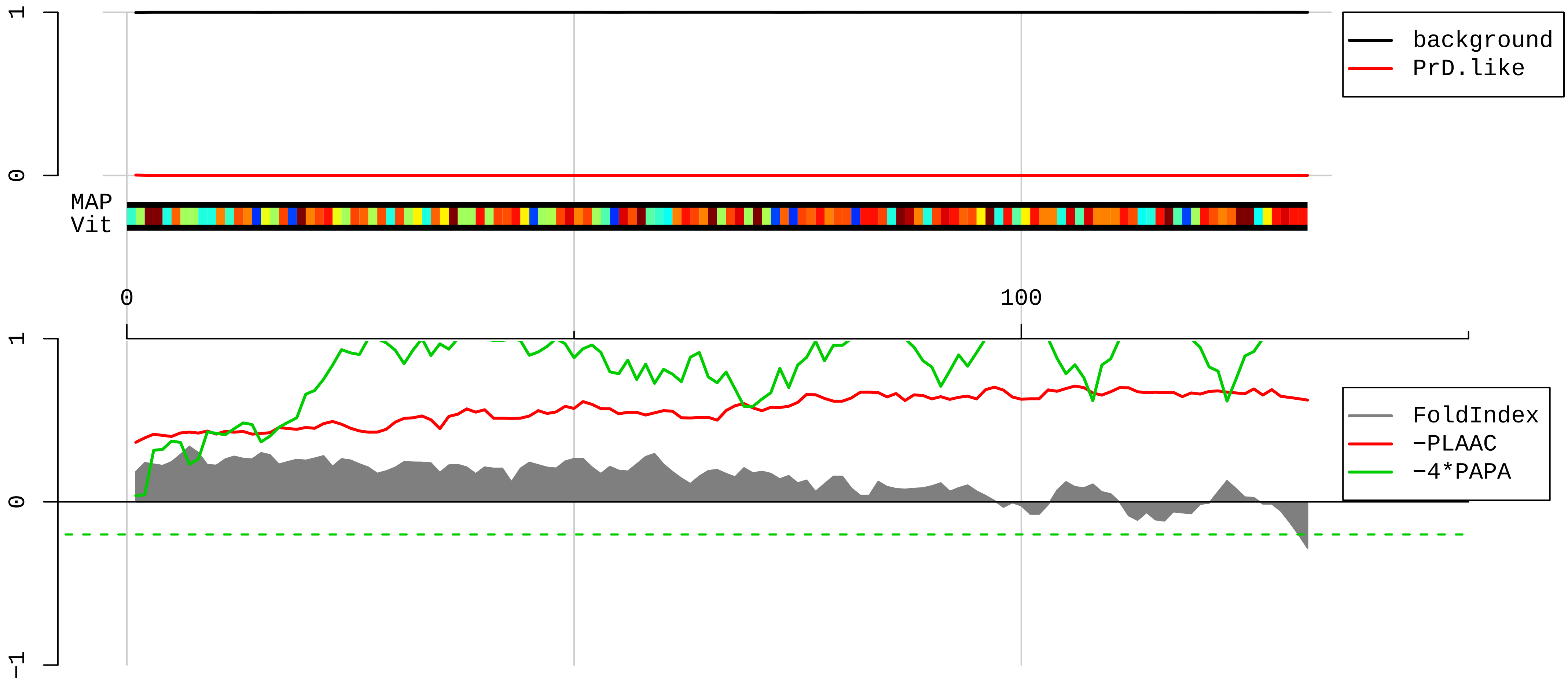


1 MAEEGIAAGGVMDVNTALQEVLKTALIHDGLARGIREAAKALDKRQAHLCVLASNCDEPMYVKLVEALCAEHQINLIKVDDNKKLGEWVGLCKIDREGKP

101 RKVVGCSCVVVKDYGKESQAKDVIEEYFKCKK

**AAH50910.1 Ubap2l protein [Mus musculus]**

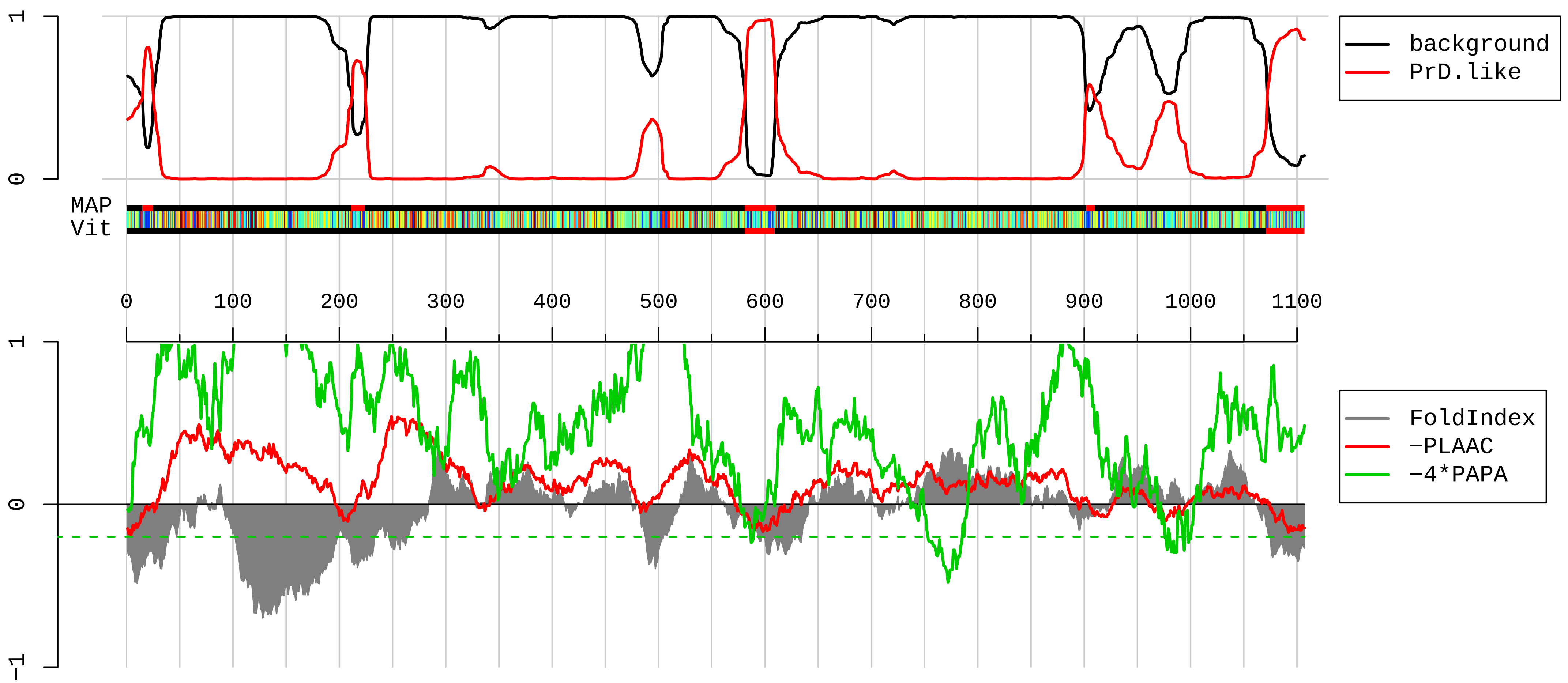


1 MMTSVGTNRARGNWEQPQNQNQTQHKQRPQATAEQIRLAQMISDHNDADFEEKVKQLIDITGKNQDECVIALHDCNGDVNRAINVLLEGNPDTHSWEMVG
101 KKKGVSGQKDGGQTESNEEGKENRDRDRDYSRRRGGPPRRGRGASRGREFRGQENGLDGTKSGGPSGRGTDRGRRGRGRGRGSSGRRGGRFSAQGMGTFN
201 PADYAEPANTDDNYGNSSGNTWNNTGHFEPDDGTRLDFIGVEGSNYPRKFETAPGAWRTATEEWGTEDWNEDLSETKIFTASNVSSVPLPAENVTITAGQ
301 RIDLAVLLGKTPSSMENDSSNLDPSQAPSLAQPLVFSNSKQNAISQPASGSTFSHHSMVSMLGKGFGDVGEAKGGSTTGSQFLEQFKTAQALAQLAAQHS
401 QSGSTTTSSWDMGSTTQSPSLVQYDLKSANDSTVHSPFTKRQAFTPSSTMMEVFLQEKPPAVATSTAAPPPPSSPLPSKSTSAPQMSPGSSDNQSSSPQP
501 AQQKLKQQKKKTSLTSKIPALAVEMPGSADISGLNLQFGALQFGSEPVLSDYESTPTTSASSSQAPSSLYTSTASESSSTVSSNQSQESGYQSGPIQSTT
601 YTSQNNAQGPLYEQRSTQTRRYPSSISSSPQKDLTQAKNGFSSVQATQLQTTQSVEGATGSAVKSESPSTSSIPSLNETVPAASLLTTANQHSSSLSGLS
701 HTEEIPNTTTTQHSSALSTQQNTLSSSTSSGRTSTSTLLHTSVESEANLHSSSSTFSTTSSTVSAPPPVVSVSSSLNSGSSLGLSLGSNSTVTASTRSSV
801 ATTSGKAPPNLPPGVPPLLPNPYIMAPGLLHAYPPQVYGYDDLQMLQTRFPLDYYSIPFPTPTTPLTGRDGSLASNPYSGDLTKFGRGDASSPAPATTLA
901 QPQQNQTQTHHTTQQTFLNPALPPGYSYTSLPYYTGVPGLPSTFQYGPAVFPVAPTSSKQHGVNVSVNASATPFQQPSGYGSHGYNTGVSVTSSNTGVPD
1001 ISGSVYSKTQQSFEKQGFHSGTPAASFNLPSALGSGGPINPATAAAYPPAPFMHILTPHQQPHSQILHHHLQQDGQTGSGQRSQTSSIPQKPQTNKSAYN
1101 SYSWGAN

**AAH65172.1 Hnrpm protein [Mus musculus]**

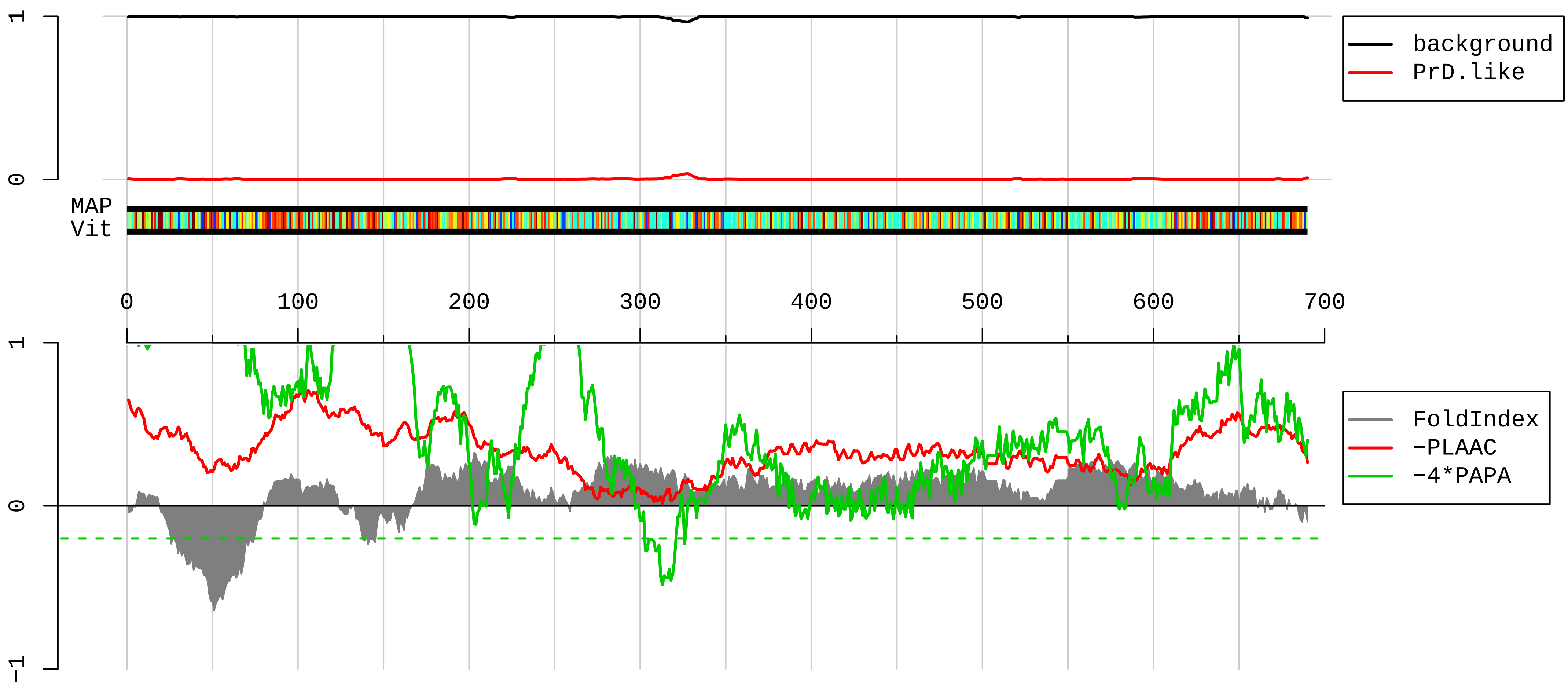


1 MAAGVEAAAEVAATEPKMEEESGAPCVPSGNGAPGPKGEERPTQNEKRKEKNIKRGGNRFEPYSNPTKRYRAFITNIPFDVKWQSLKDLVKEKVGEVTYV
101 ELLMDAEGKSRGCAVVEFKMEESMKKAAEVLNKHSLSGRPLKVKEDPDGEHARRAMQKAGRLGSTVFVANLDYKVGWKKLKEVFSMAGVVVRADILEDKD
201 GKSRGIGIVTFEQSIEAVQAISMFNGQLLFDRPMHVKMDERALPKGDFFPPERPQQLPHGLGGIGMGLGPGGQPIDANHLSKGIGMGNLGPAGMGMEGIG
301 FGINKIGGMEGPFGGGMENMGRFGSGMNMGRINEILSNALKRGEIIAKQGGGGAGGSVPGIERMGPGIDRISGAGMERMGAGLGHGMDRVGSEIERMGLV
401 MDRMGSVERMGSSIERMGPLGLDHMASSIERMGQTMERIGSGVERMGAGMGFGLERMAAPIDRVGQTIERMGSGVERMGPAIERMGLSMDRMVPTGMGAS
501 LERMGPVMDRMATGLERMGANNLERMGLERMGANSLERMGLERMGANSLERMGPAMGPALGAGIERMGLAMGGAGGASFDRAIEMERGNFGGSFAGSFGG
601 AGGHAPGVARKACQIFVRNLPFDFTWKMLKDKFNECGHVLYADIKMENGKSKGCGVVKFESPEVAERACRMMNGMKLSGREIDVRIDRNA

**NP_666312.1 eukaryotic translation initiation factor 3 subunit C [Mus musculus]**

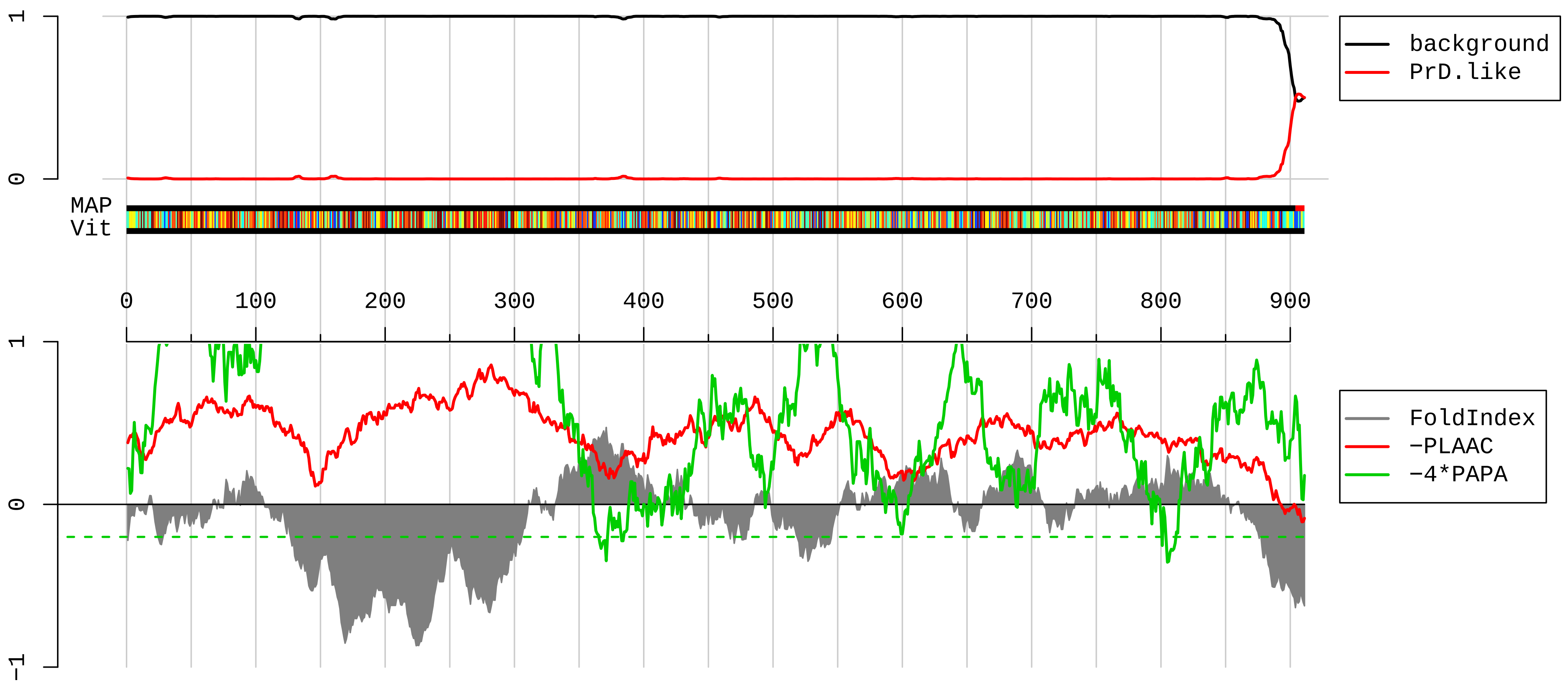


1 MSRFFTTGSDSESESSLSGEELVTKPVSGNYGKQPLLLSEDEEDTKRVVRSAKDKRFEELTNLIRTIRNAMKIRDVTKCLEEFELLGKAYGKAKSIVDKE
101 GVPRFYIRILADLEDYLNELWEDKEGKKKMNKNNAKALSTLRQKIRKYNRDFESHITNYKQNPEQSADEDAEKNEEDSEGSSDEDEDEDGVGNTTFLKKK
201 QESSGESRKFHKKMEDDDEDSEDSEDEEWDTSSTSSDSDSEEEEGKQTVLASKFLKKAPTTEEDKKAAEKKREDKAKKKHDRKSKRLDEEEEDNEGGEWE
301 RVRGGVPLVKEKPKMFAKGTEITHAVVIKKLNEILQVRGKKGTDRATQIELLQLLVQIAAENNLGVGVIVKIKFNIIASLYDYNPNLATYMKPEMWQMCL
401 DCINELMDTLVAHSNIFVGENILEESENLHNFDQPLRVRGCILTLVERMDEEFTKIMQNTDPHSQEYVEHLKDEAQVCAIIERVQRYLEEKGTTEEICQI
501 YLRRILHTYYKFDYKAHQRQLTPPEGSSKSEQDQAENEGEDSAVLMERLCKYIYAKDRTDRIRTCAILCHIYHHALHSRWYQARDLMLMSHLQDNIQHAD
601 PPVQILYNRTMVQLGICAFRQGLTKDAHNALLDIQSSGRAKELLGQGLLLRSLQERNQEQEKVERRRQVPFHLHINLELLECVYLVSAMLLEIPYMAAHE
701 SDARRRMISKQFHHQLRVGERQPLLGPPESMREHVVAASKAMKMGDWKTCHSFIINEKMNGKVWDLFPEADKVRTMLVRKIQEESLRTYLFTYSSVYDSI
801 SMETLSDMFELDLPTVHSIISKMIINEELMASLDQPTQTVVMHRTEPTAQQNLALQLAEKLGSLVENNERVFDHKQGTYGGYFRDQKDGYRKNEGYMRRG
901 GYRQQQSQTAY

**EDL40597.1 purine rich element binding protein B [Mus musculus]**

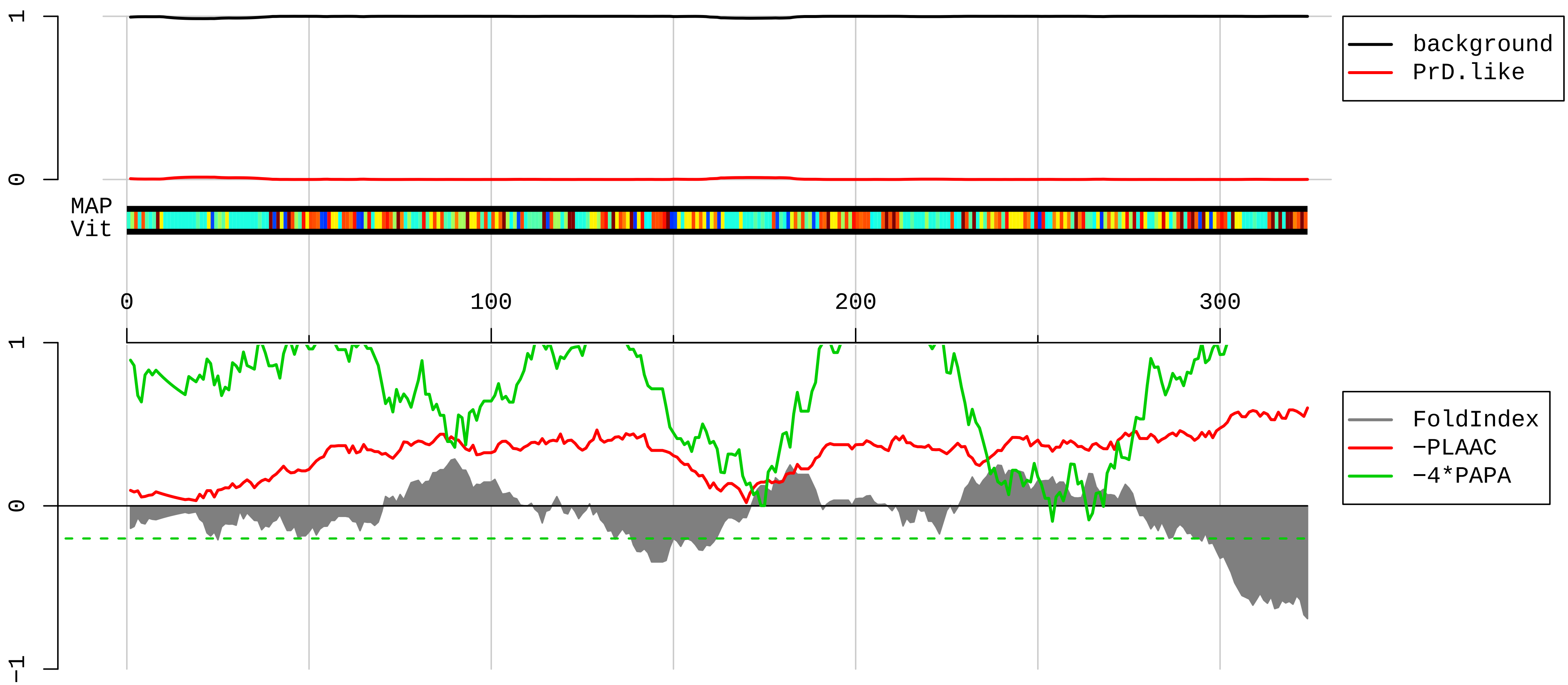


1 MADGDSGSERGGGGGGGGGPGGFQPAPRGGGGGGGGPGGEQETQELASKRLDIQNKRFYLDVKQNAKGRFLKIAEVGAGGSKSRLTLSMAVAAEFRDSLG
101 DFIEHYAQLGPSSPEQLAAGAEEGGGPRRALKSEFLVRENRKYYLDLKENQRGRFLRIRQTVNRGGGGFGGGPGPGGLQSGQTIALPAQGLIEFRDALAK
201 LIDDYGGDEDELAGGPGGGAGGPGGGLYGELPEGTSITVDSKRFFFDVGCNKYGVFLRVSEVKPSYRNAITVPFKAWGKFGGAFCRYADEMKEIQERQRD
301 KLYERRGGGSGGGDESEGEEVDED

AAI00600.1 Ribosomal protein L35A [Mus musculus]

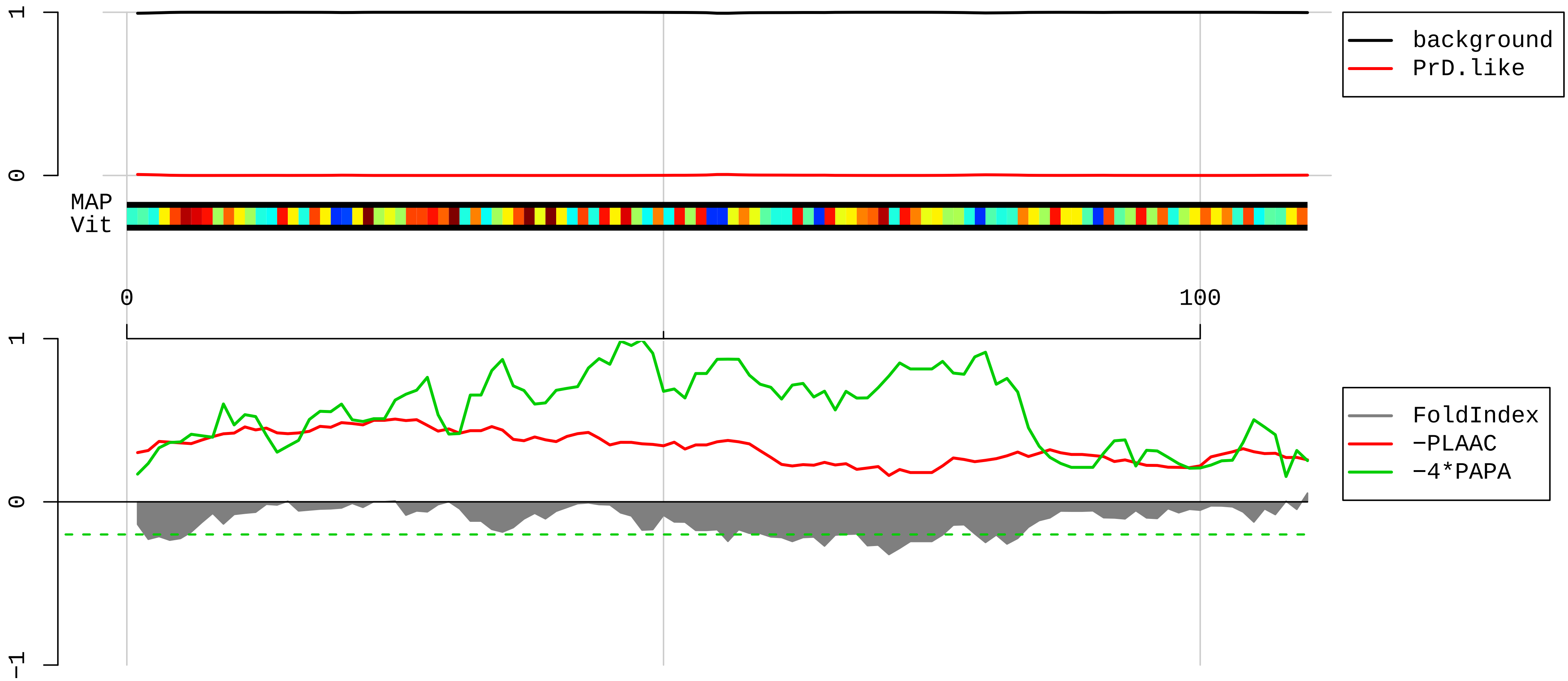


1 MSGRLWCKAIFAGYKRGLRNQREHTALLKIEGVYARDETEFYLGKRCAYVYKAKNNTVTPGGKPNKTRVIWGKVTRAHGNSGMVRAKFRSNLPAKAIGHR
101 IRVMLYPSRI

**AAH03820.1 Ssb protein [Mus musculus]**

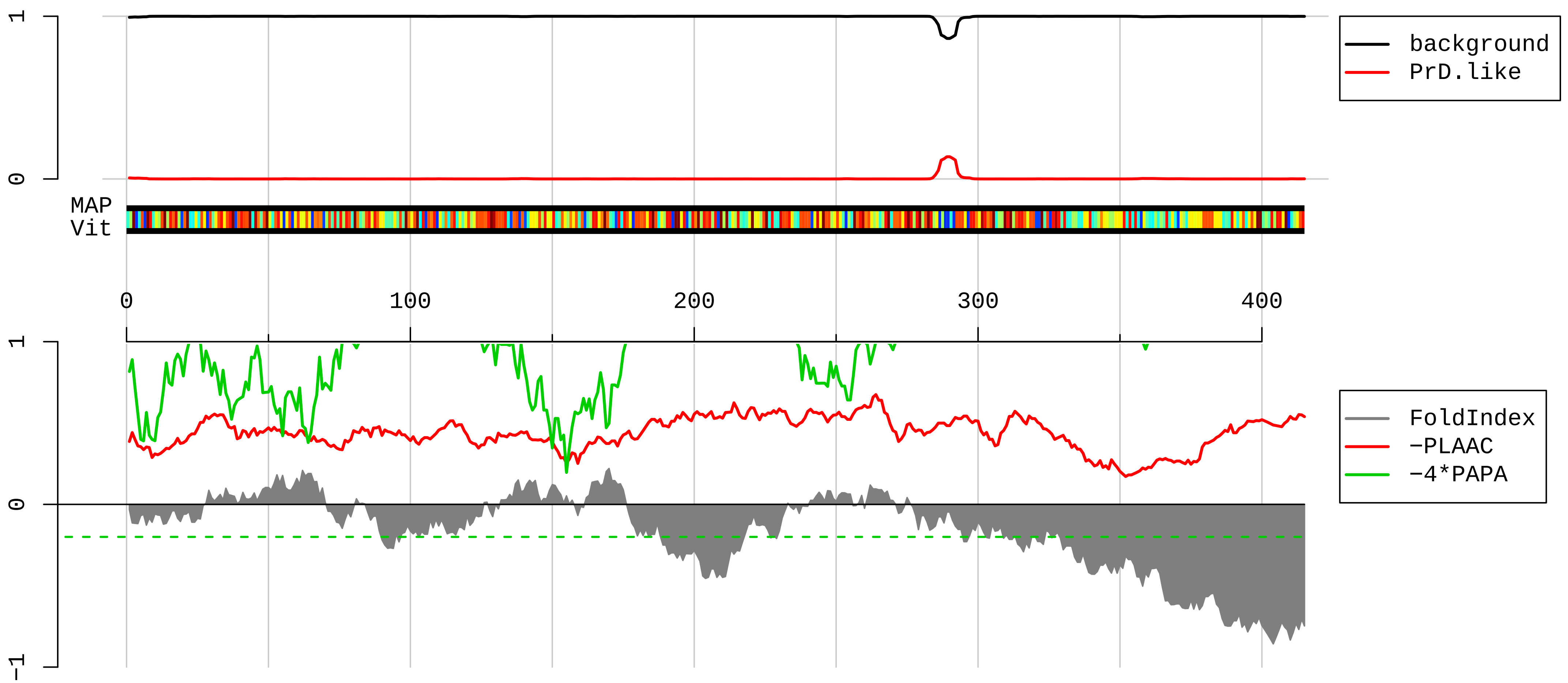


1 MAENGDNEKMTALEAKICHQIEYYFGDFNLPRDKFLKEQIKLDEGWVPLETMIKFNRLNRLTTDFNVIVQALSKSKAKLMEVSADKTKIRRSPSRPLPEV
101 TDEYKNDVKNRSVYIKGFPTDATLDDIKEWLDDKGQILNIQMRRTLHKTFKGSIFAVFDSIQSAKKFVEIPGQKYKDTNLLILFKEDYFAKKNEERKQSK
201 VEAKLKAKQEHEGRHKPGSTETRALEGKMGCLLKFSGDLDDQTCREDLHFLFSNHGEIKWVDFARGAKEGIILFKEKAKEALEKARNANNGNLLLRNKKV
301 TWKVLEGHAEKEALKKITDDQQESLNKWKSKGGHAGGRFKGSHVFTAARRFKGKGKGNRPGYAGAPKGRGQFHGRRTRFDDDDRRRGPMKRGRDGRDREE
401 PASKHKKRENGARDK

**AAH89334.1 Tyrosine 3−monooxygenase/tryptophan 5−monooxygenase activation protein, zeta polypeptide [Mus musculus]**

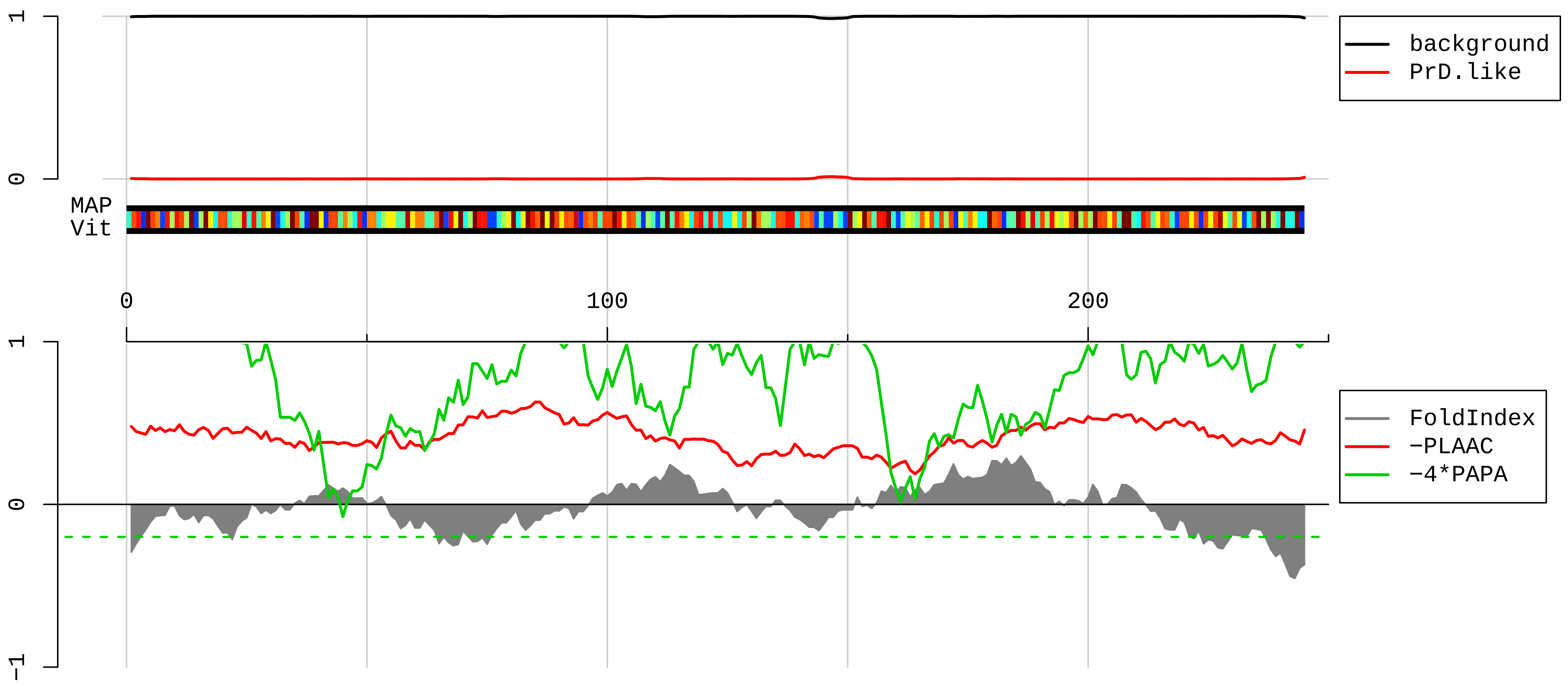


1 MDKNELVQKAKLAEQAERYDDMAACMKSVTEQGAELSNEERNLLSVAYKNVVGARRSSWRVVSSIEQKTEGAEKKQQMAREYREKIETELRDICNDVLSL
101 LEKFLIPNASQPESKVFYLKMKGDYYRYLAEVAAGDDKKGIVDQSQQAYQEAFEISKKEMQPTHPIRLGLALNFSVFYYEILNSPEKACSLAKTAFDEAI
201 AELDTLSEESYKDSTLIMQLLRDNLTLWTSDTQGDEAEAGEGGEN

**AAH24609.1 Rpl18a protein [Mus musculus]**

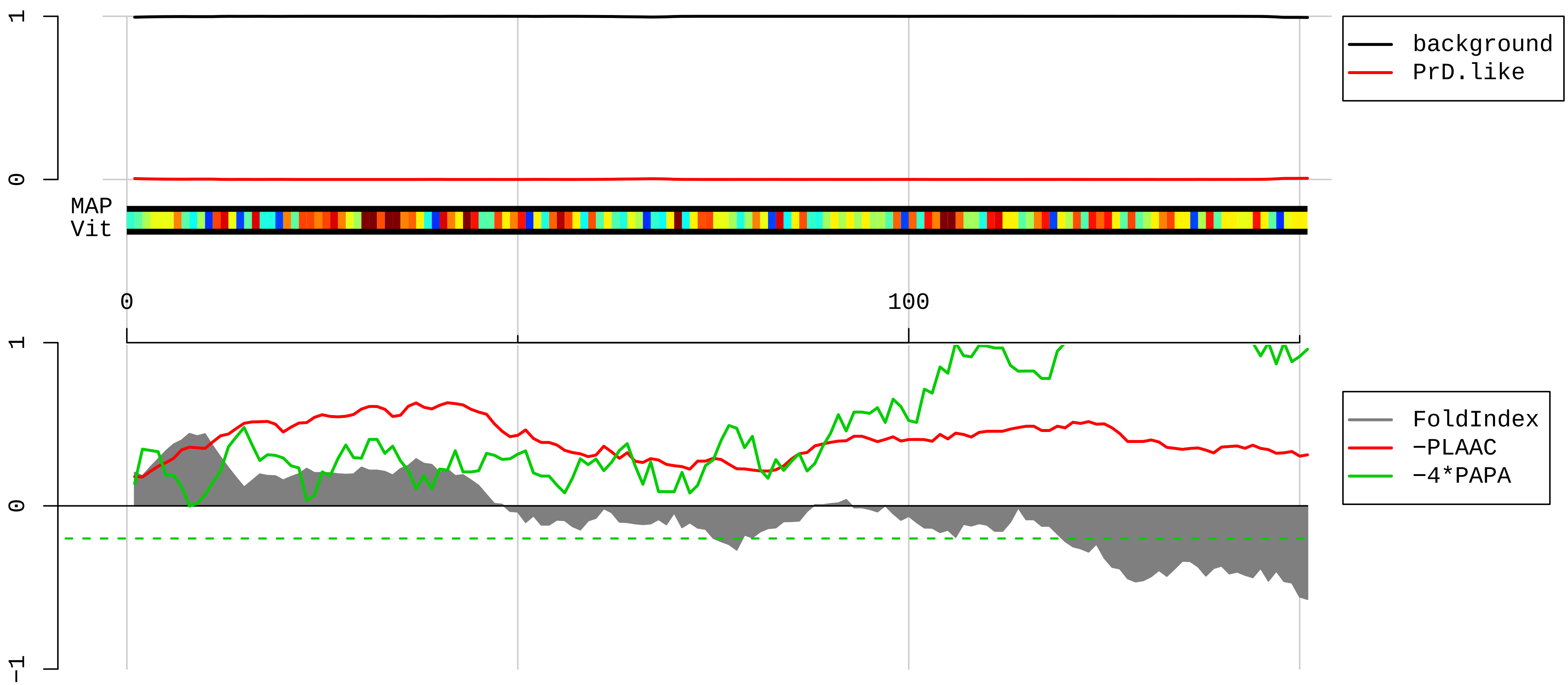


1 MPHTTTVPYANLCTQPCGGQVPLLVLCVTAEEDEEVIRGNCVFEKSPLRVKNFGIWLRYDSRSGTHNMYREYRDLTTAGAVTQCYRDMGARHRARAHSIQ

101 IMKVEEIAAGKCRRPAVKQFHDSKIKFPLPHRVLRRQHKPRFTTKRPNTFF

**AAH92539.1 Ribosomal protein L19 [Mus musculus]**

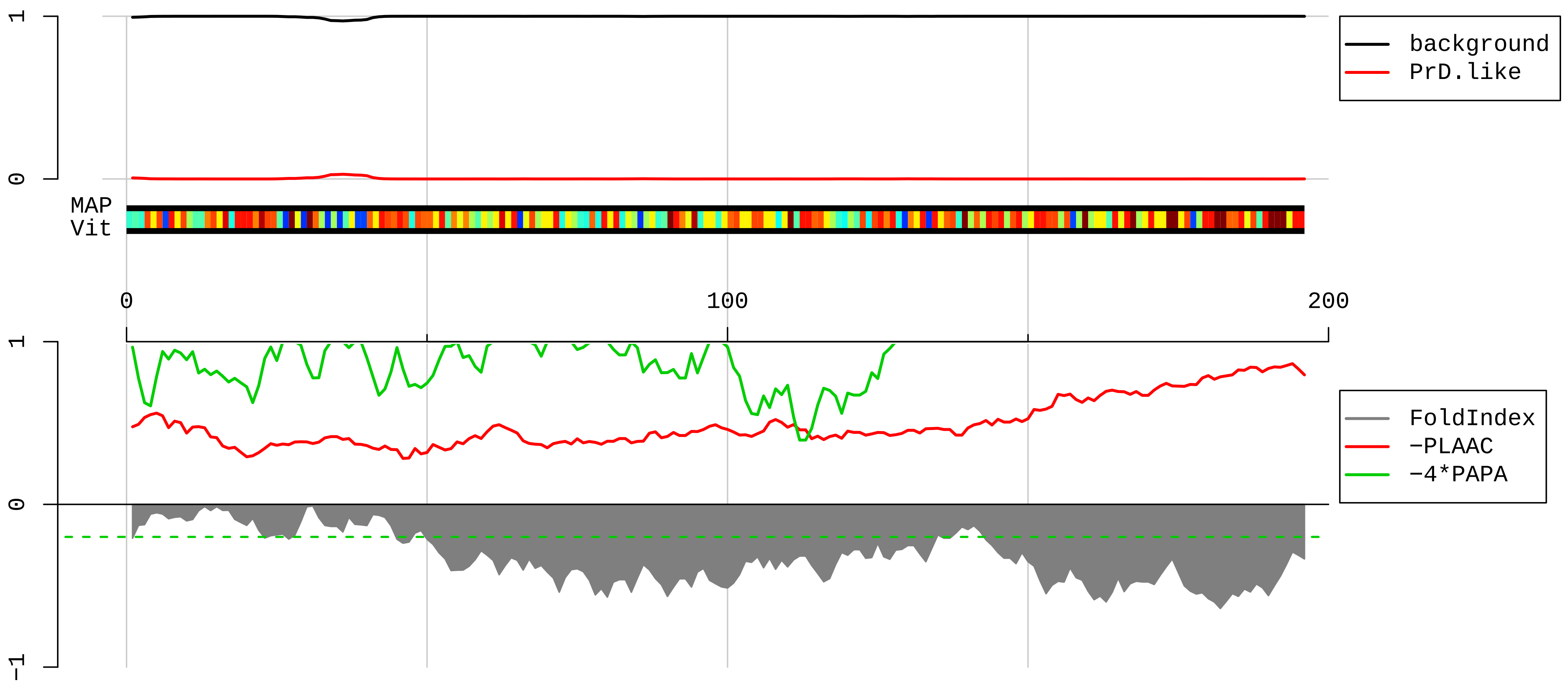


1 MSMLRLQKRLASSVLRCGKKKVWLDPNETNEIANANSRQQIRKLIKDGLIIRKPVTVHSRARCRKNTLARRKGRHMGIGKRKGTANARMPEKVTWMRRMR
101 ILRRLLRRYRESKKIDRHMYHSLYLKVKGNVFKNKRILMEHIHKLADKARKKLLADQAEARRSKTKEARKRREERLQAKKEEIIKTLSKEEETKK

**AAH58118.1 Ribosomal protein L34 [Mus musculus]**

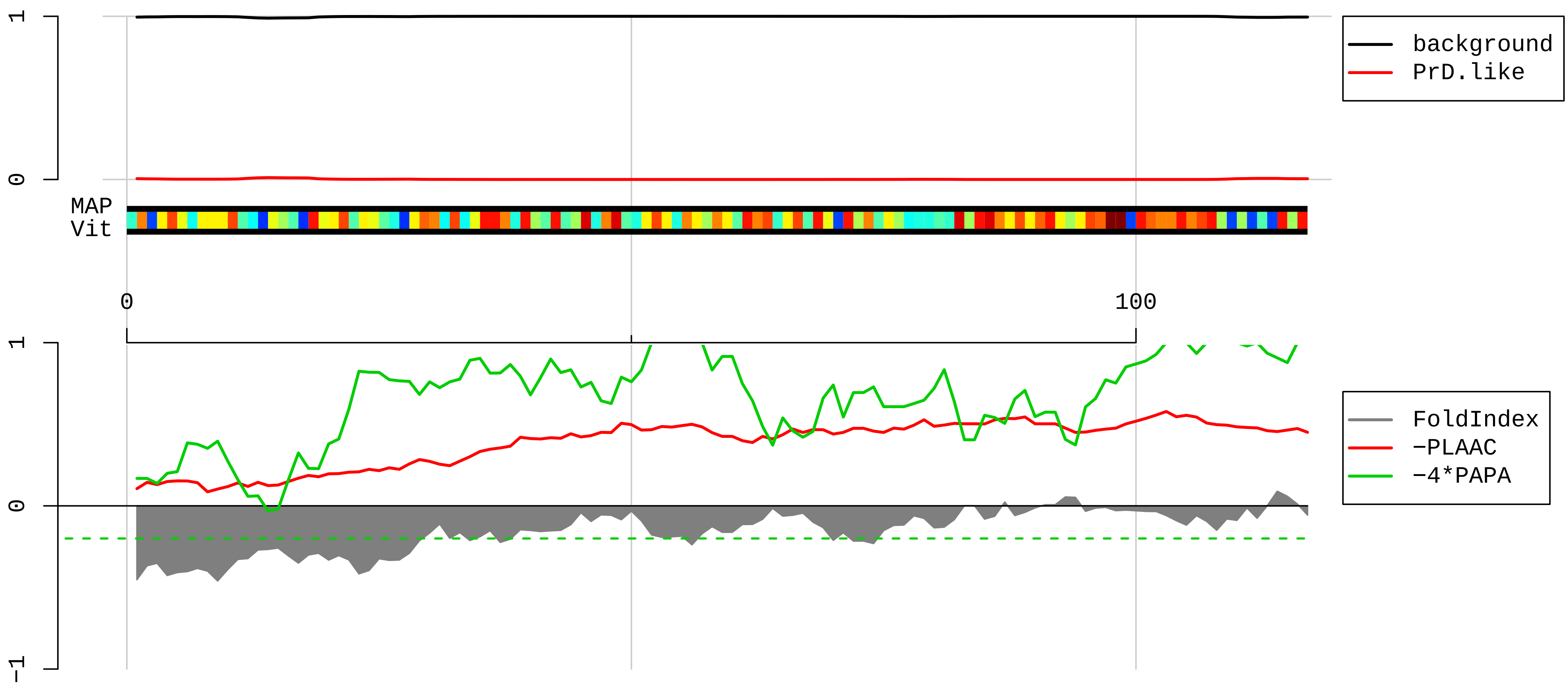


1 MVQRLTYRRRLSYNTASNKTRLSRTPGNRIVYLYTKKVGKAPKSACGVCPGRLRGVRAVRPKVLMRLSKTQKHVSRAYGGSMCAKCVRDRIKRAFLIEEQ
101 KIVVKVLKAQAQSQKAK

**AAI00457.1 Ribosomal protein S26 [Mus musculus]**

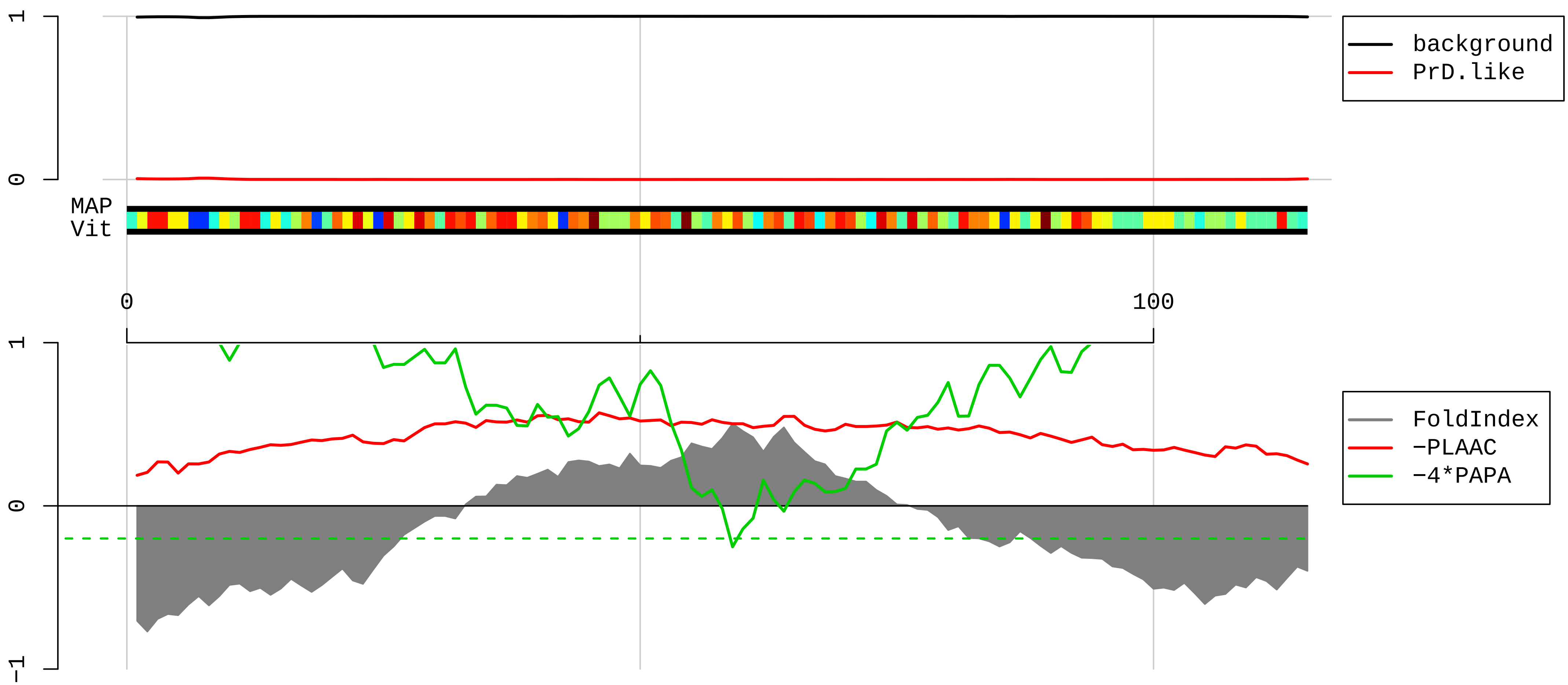


1 MTKKRRNNGRAKKGRGHVQPIRCTNCARCVPKDKAIKKFVIRNIVEAAAVRDISEASVFDAYVLPKLYVKLHYCVSCAIHSKVVRNRSREARKDRTPPPR

101 FRPAGAAPRPPPKPM

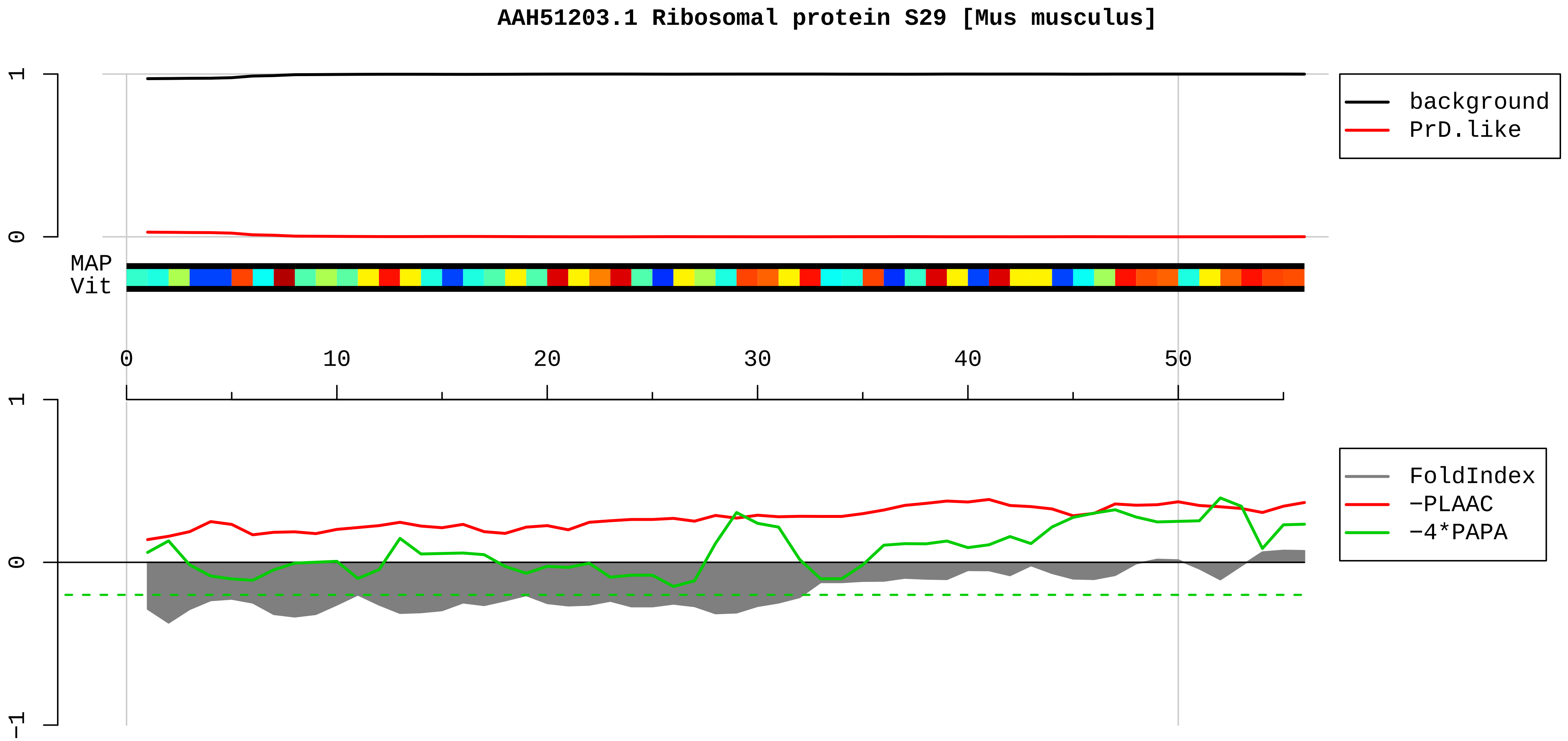


1 MGHQQLYWSHPRKFGQGSRSCRVCSNRHGLIRKYGLNMCRQCFRQYAKDIGFIKLD

**AAH23677.1 Microtubule-associated protein 7 domain containing 1 [Mus musculus]**

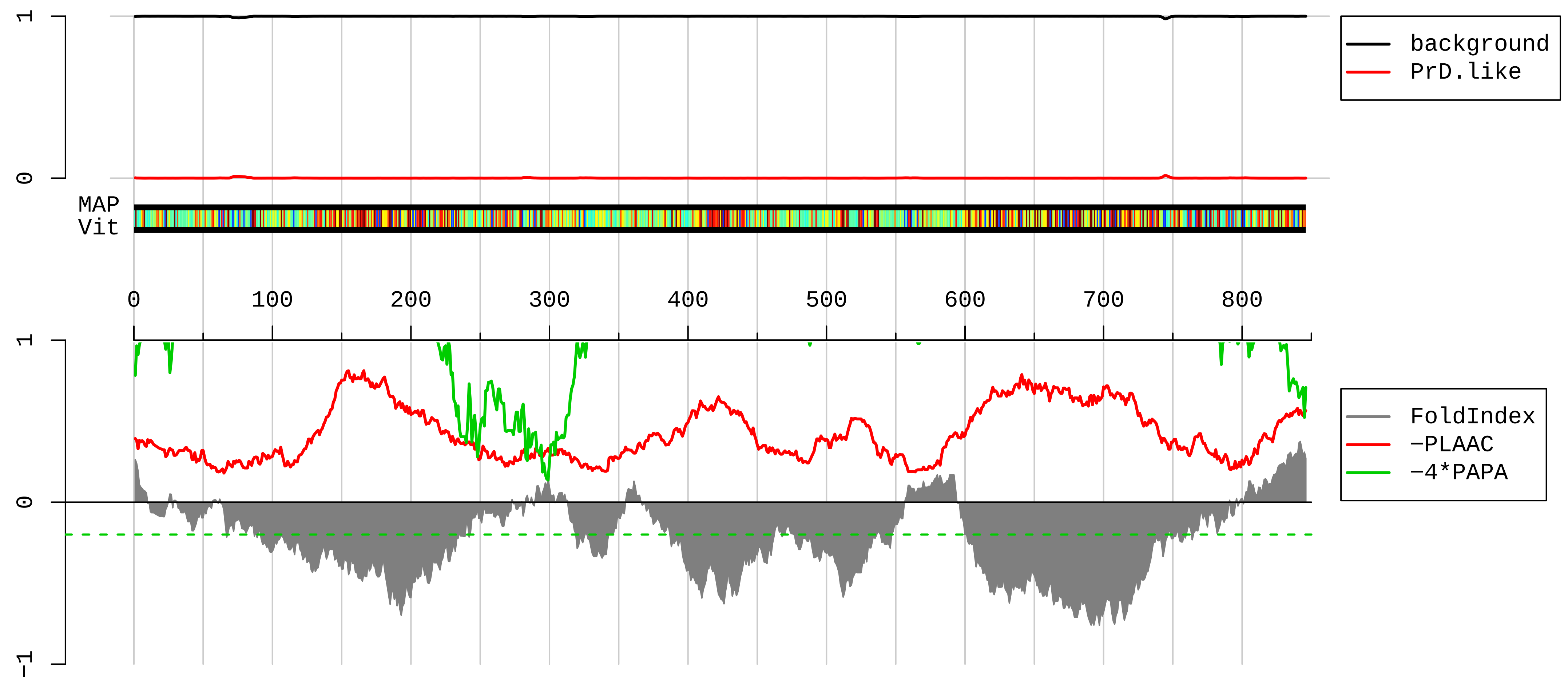


1 MESGPRVEPGPGAPAAVLARIPQEPRPSPEGDPSPPPPPTPMSALVPDTPPDTPPALKTATNPKQLPLEPGNPTGQISPQPAPPQEECPSSEAKSRGPTP
101 TATGPREAKPSRRSSQPSPTTVSAYDSPPAKQDVKKAGERHKLAKERREERAKYLAAKKAVWLEKEEKAKALREKQLQERRRRLEEQRLKAEQRRAALEE
201 RQRQKLEKNKERYEAAIQRSVKKTWAEIRQQRWSWAGALHHSSPGRKTSGSRCSVSAVNLPKHVDSIINKRLSKSSATLWNSPSRNRSLQLSAWESSIVD
301 RLMTPTLSFLARSRSAVTLPRNGRDQGRGSGPGRRPTRARAGASLAPGPHPDRTHPSAAVPVCPRSASASPLTPCSAPRSAHRCTPSGERPERRKPGAGG
401 SPALARRRLEATPVQKKEKKDKERENEKEKSALARERNLKKRQSLPASIRPRLSTGSELSPKSKARPSSPSTTWHRPASPCPSPGPGHALPPKPPSPRGT
501 TASPKGRVRRKEEAKESPSPSGPEDKNHRKSRAAEEKEPAAPASPAPSPVPSPTPAQPQKEQSSTQIPAETAVPAVPAAPTAPPTAAPSVTPSKPMAGTT
601 DREEATRLLAEKRRQAREQREREEQERKLQAERDKRMREEQLAREAEARAEREAEARRREEQEAREKAQAEQEEQERLQKQKEEAEARSREEAERQRQER
701 EKHFQKEEQERQERRKRLEEIMKRTRKSEAAETKKQDAKETAANNSGPDPVKAVETRPSGLQKDSMQKEELAPQEPQWSLPSKEMPGSLVNGLQPLPAHQ
801 ENGFSPKGTAGDKSLGRTAEGLLPFAEAEAFLKKAVVQPPQVTEVL

#### NP_032016.1 40S ribosomal protein S30 [Mus musculus]

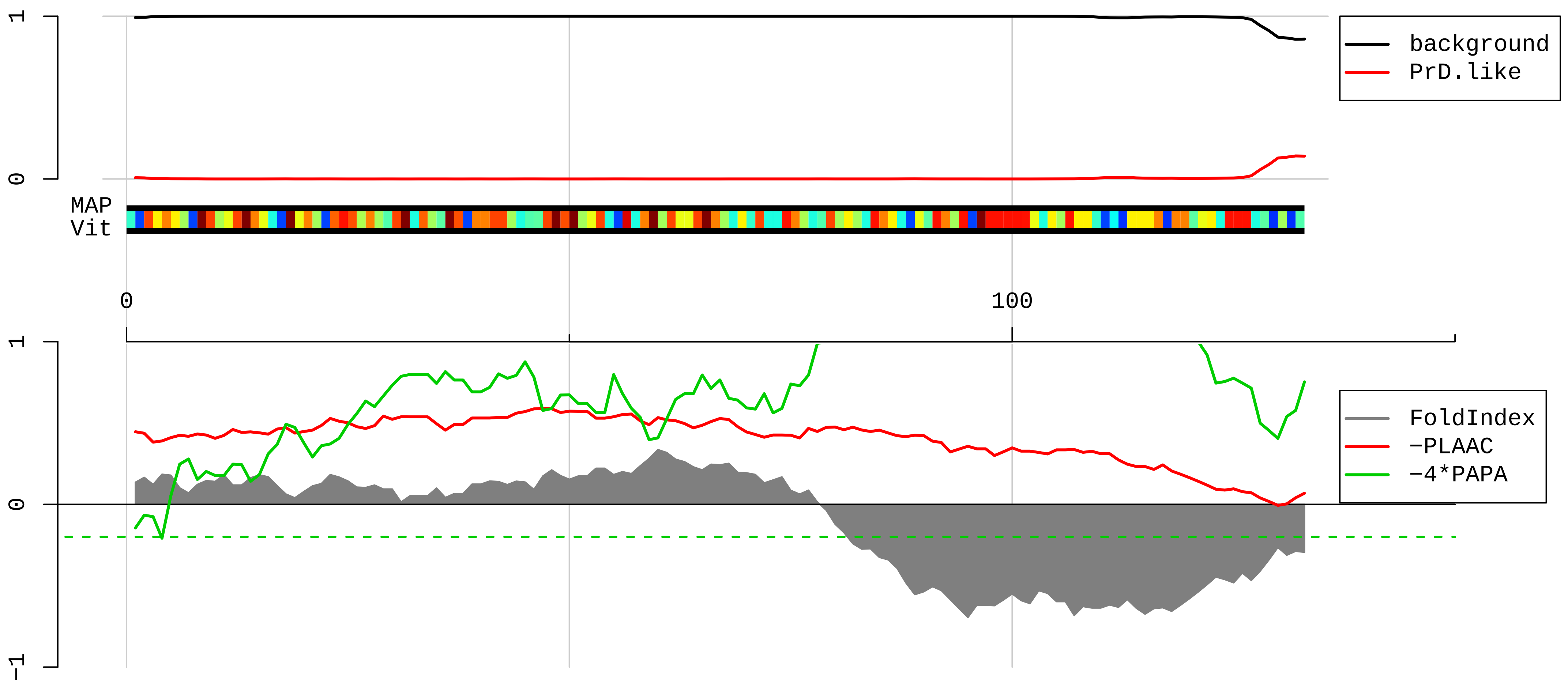


1 MQLFVRAQELHTLEVTGQETVAQIKDHVASLEGIAPEDQVVLLAGSPLEDEATLGQCGVEALTTLEVAGRMLGGKVHGSLARAGKVRGQTPKVAKQEKKK
101 KKTGRAKRRMQYNRRFVNVVPTFGKKKGPNANS
