## Supplementary datas for "m⁶A modification and prion-like domain proteins converge to dysregulate Neuronal RNA Granules in Alzheimer’s disease": Supplementary figures S1-S5.pdf

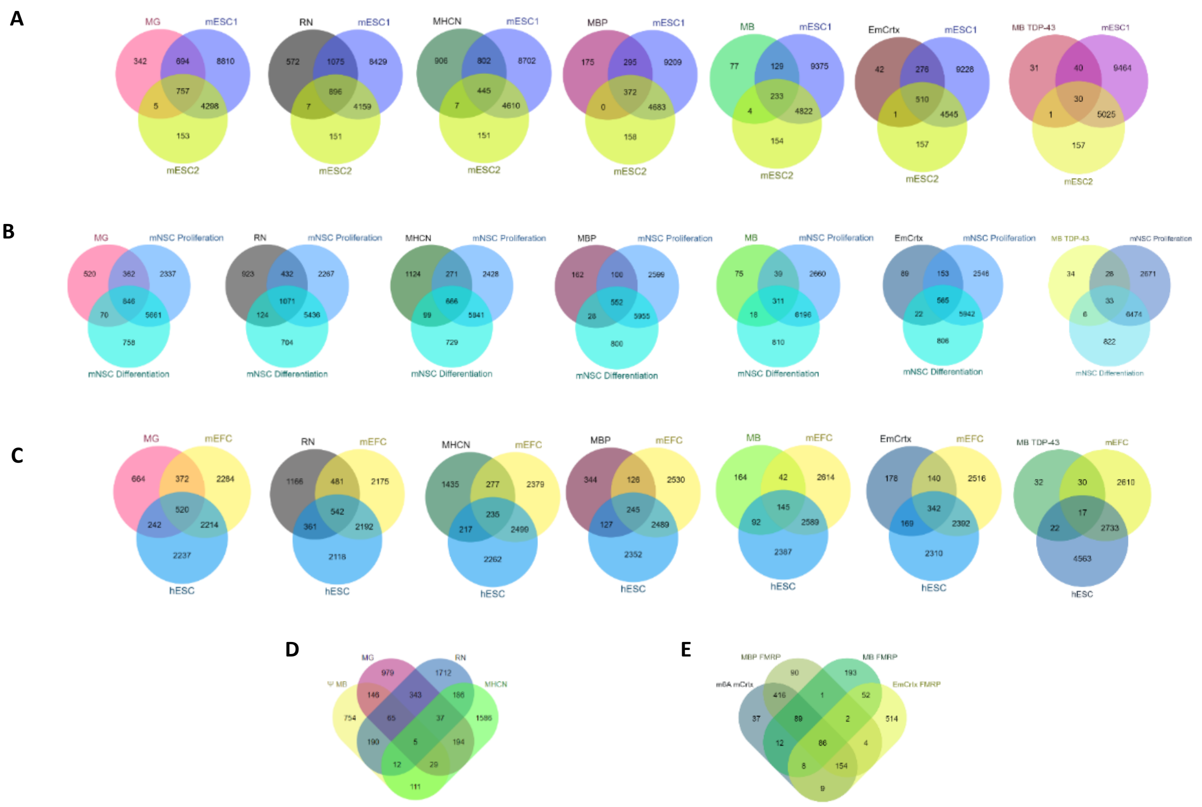


**S1. Venn diagrams obtained by overlapping mRNAs from different structures to estimate the percentage of m6A methylation.**

(**A**): overlap between the lists of mRNAs from mouse granules: (MG, n= 1798), mRNAs from rat dendritic/axonic synaptic neuropil (RN, n=2550), mRNAs from mouse hippocampal neurons (MHCN, n= 2160), mRNAs from adult mouse brain polyribosomes FMRP targets (MBP FMRP, n=842), mRNAs Associated with FMRP obtained from mouse brain (MB FMRP, n=443), mRNA FMRP targets from embryonic mouse cerebral cortex (EmCrtx FMRP, n=856), m6A methylation in TDP-43 RNAs target from mouse brain (MB TDP-43, n=101), with lists of m6A methylated mRNA obtained from**:** mouse embryonic stem cells (study1): (m6A mESC1, n=1455), and m6A methylated mRNAs from mouse embryonic stem cells (study 2): (mESC 2, n=5213). (**B**): overlap between the lists of mRNAs from MG, RN, MHCN, adult mouse brain polyribosomes FMRP targets (MBP FMRP), mRNAs associated with FMRP obtained from mouse brain (MB FMRP), mRNAs FMRP targets from embryonic mouse cerebral cortex (EmCrtx FMRP), and m6A methylated TDP-43 RNAs target from mouse brain (MB TDP-43), with lists of m6A methylated mRNAs from Mouse adult neuronal stem cells in proliferation and differentiation phases: (mNSC Proliferation, n=9206 and mNSC Differentiation, n=7336). (**C**): Overlap between the lists of mRNAs from MG, RN, MHCN, MBP FMRP, EmCrtx FMRP, MB TDP-43, with lists of m6A methylated mRNAs derived from mouse embryonic fibroblast cells (mEFC, n=15453) and m6A methylated mRNAs from human embryonic stem cells (hESC , n=7530). (**D**): Overlap between the lists of mRNAs from MG), RN, MHCN, and lists of RNAs with pseudouridine modification from mouse brain (Ψ MB, n= 1313). (**E**): overlap between the lists of mRNAs from MBP FMRP, MB FMRP, EmCrtx FMRP, and lists of mRNAs m6A methylated from mouse cerebral cortex (m6A mCrtx, n=811).

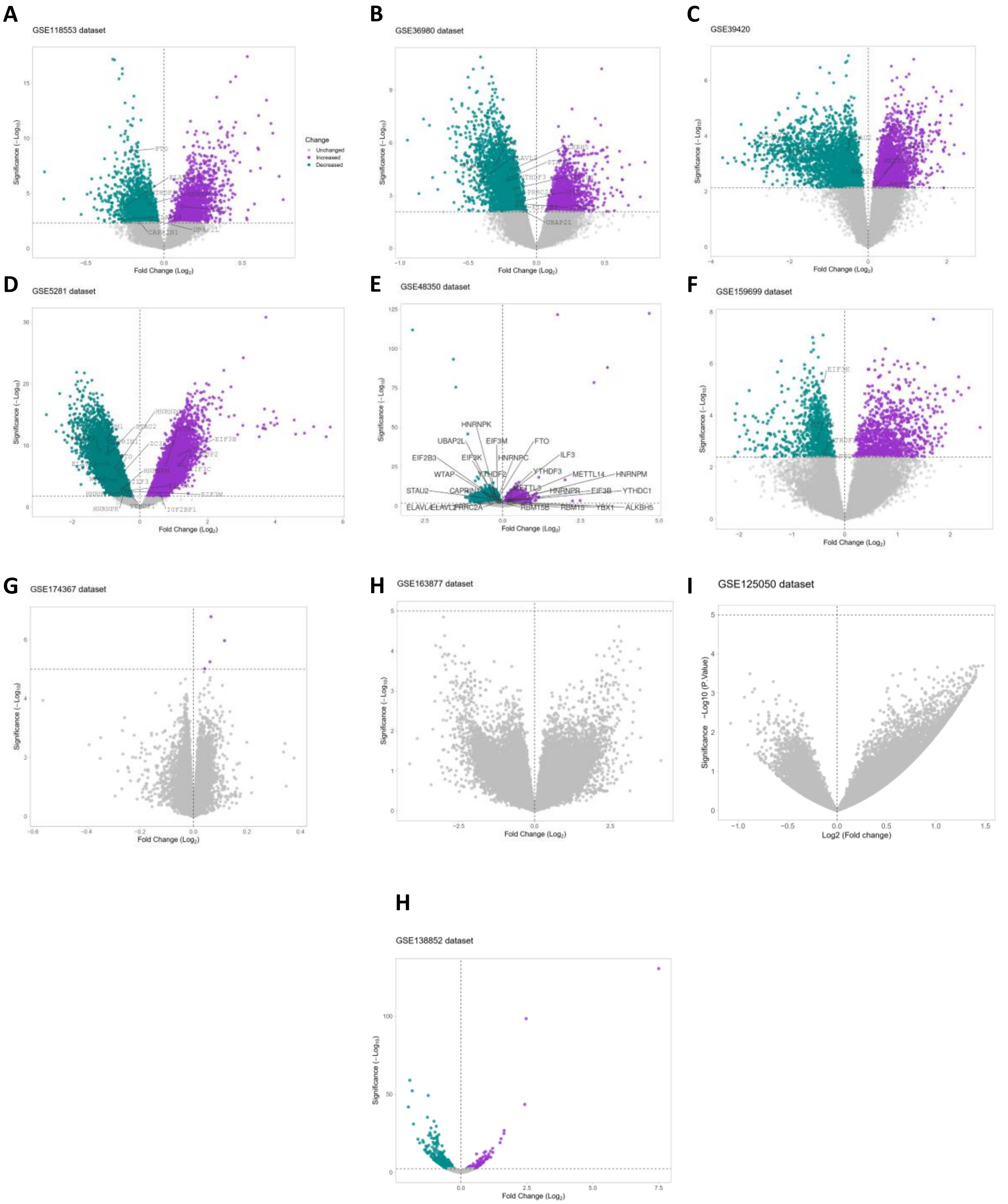


**Figure S2**: Volcanoplots describing differentially expressed genes in 10 human datasets with differentially expressed m6A methylation genes and proteins with PrLDs highlighted (LogFC=0, *P*≤0.05). A-E: Volcanoplots prepared from RNA-seq datasets. F-J: Volcanoplots prepared from microarrays datasets.

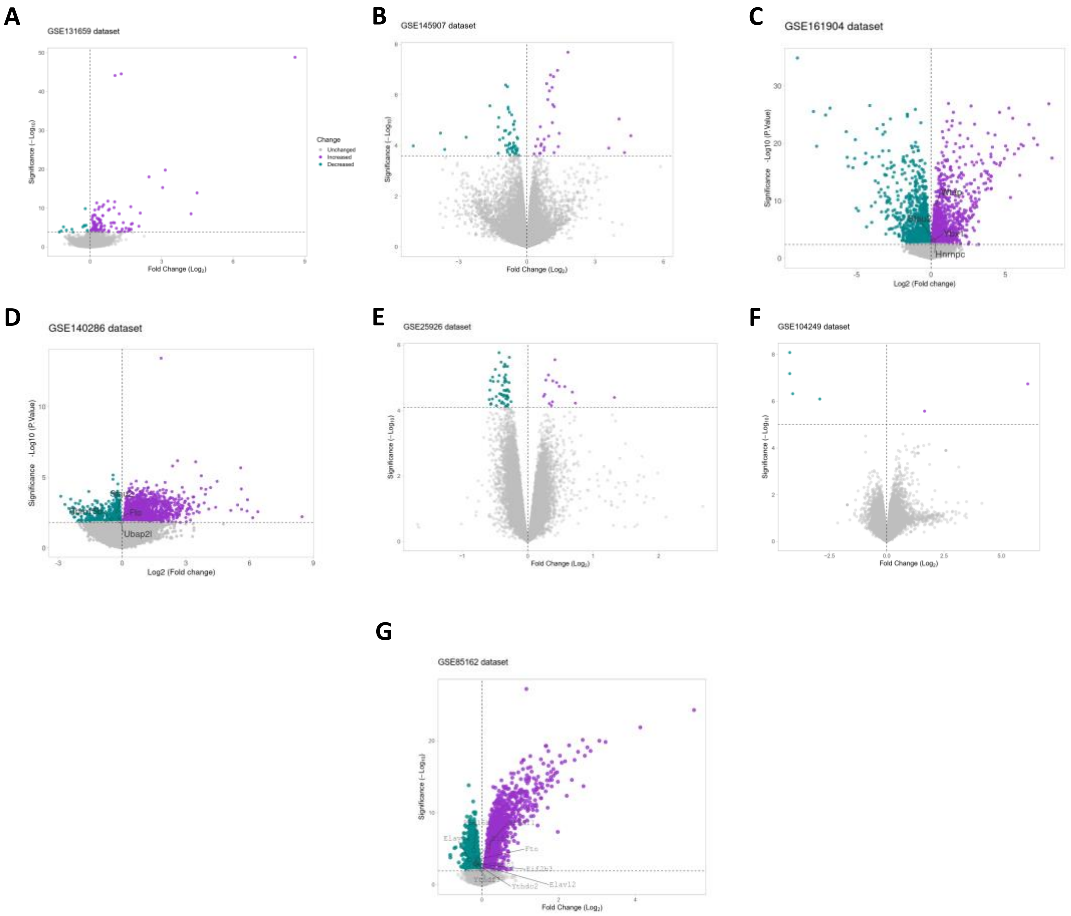


**Figure S3:** Volcanoplots describing differentially expressed genes in 7 mice datasets with differentially expressed m6A methylation genes and proteins with PrLD highlighted (LogFC=0, $P \leq 0.05$). A-D: Volcanoplots prepared from RNA-seq datasets. E-G: Volcanoplots prepared from microarrays datasets.

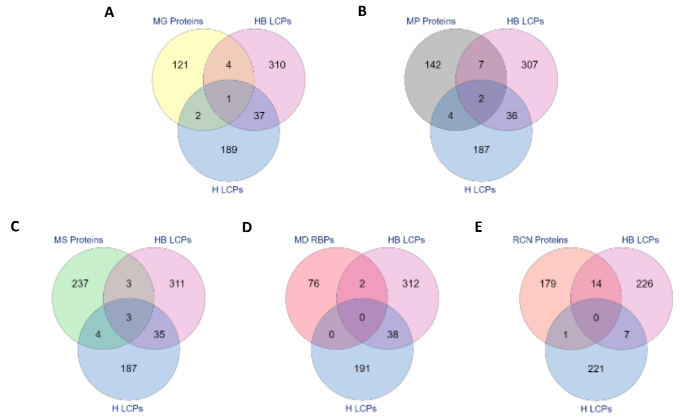


**Figure S4**: Venn diagrams obtained by overlapping lists of proteins from different structures to estimate the percentage of proteins with low complexity domains and prion-like domain. List of proteins from mouse granules (MG Proteins) (A), mouse polyribosomal proteins (MP Proteins) (B), mouse synaptosome proteins (MS Proteins) (C), mouse dendritic associated RNA-binding proteins (MD RBPs) (D), Rat cultured cortical neurons proteins (RCN Proteins) (E), with lists of human proteins with low complexity proteins derived from backbones (HB LCPs) and human proteins with prion-like domains detected by PLAAC (H LCPs).

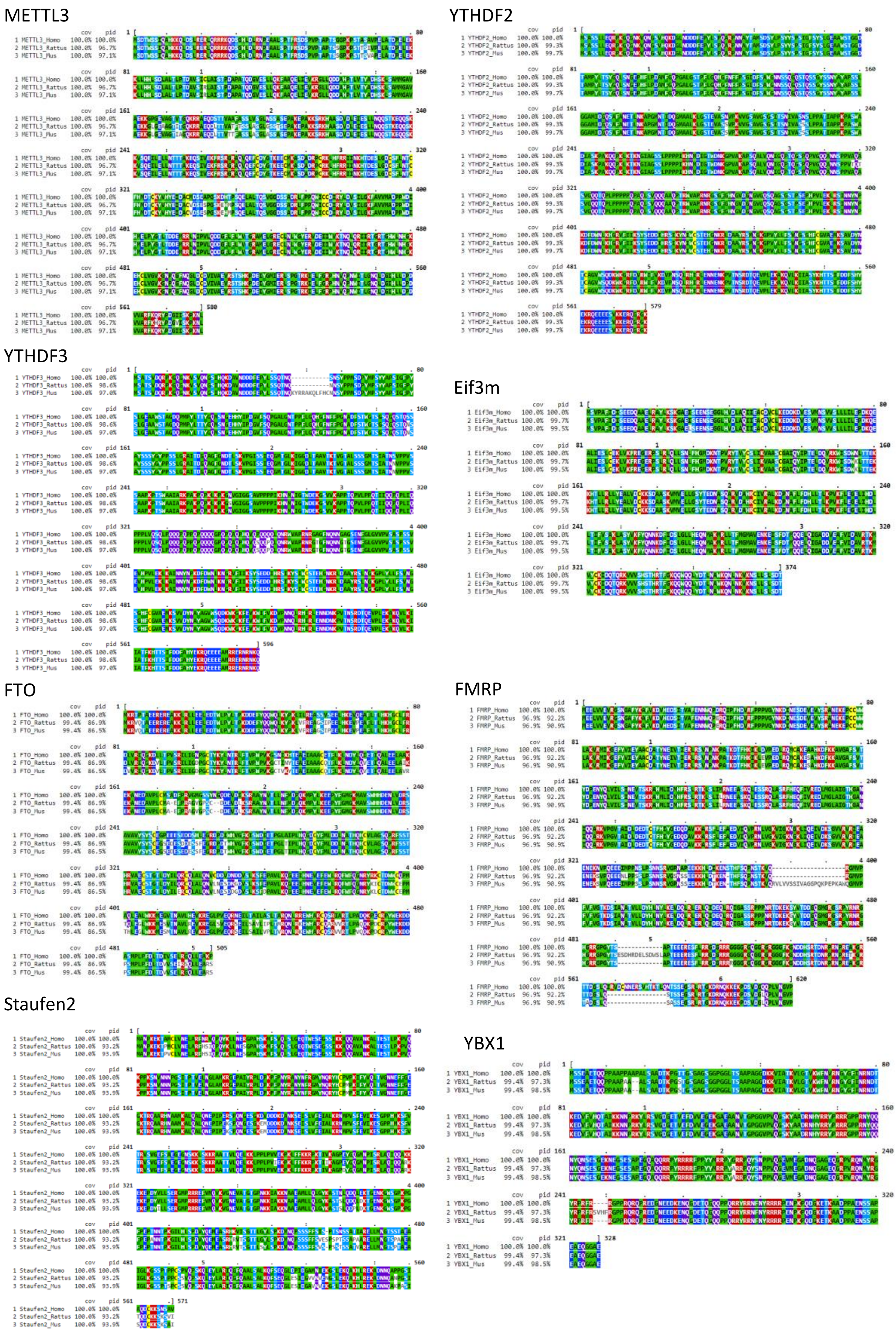


Figure S5. Multiple protein alignment of m6A target genes: Methyltransferase like 3 (Mettl3), fat mass and obesity-associated protein (Fto), YTH domain-containing family protein 2 (YTHDF2), YTH domain-containing family protein 3 (YTHDF3), double-stranded RNA-binding protein Staufen homolog 2 (Staufen 2), Fragile x mental retardation protein (FMRP), Y-box binding protein 1 (YBX1), eukaryotic translation initiation factor 3 subunit M (Eif3m), between three different species; Homo sapiens, Mus musculus, and Rattus norvegicus. The figure was created using MView: (https://www.ebi.ac.uk/Tools/msa/mview/). Coverage and percentage identity values are indicated in the “cov” and “pid” columns respectively
